# Diurnal Dynamics of the Arabidopsis Rosette Proteome and Phosphoproteome

**DOI:** 10.1101/2020.09.11.293779

**Authors:** R. Glen Uhrig, Sira Echevarría-Zomeño, Pascal Schlapfer, Jonas Grossmann, Bernd Roschitzki, Niklas Koerber, Fabio Fiorani, Wilhelm Gruissem

**Affiliations:** Institute of Molecular Plant Biology, Department of Biology, ETH Zurich, 8092 Zurich, Switzerland; Functional Genomics Center Zurich, University of Zurich, 8057 Zurich, Switzerland; Institute of Bio- and Geosciences, IBG-2: Plant Sciences, Forschungszentrum Jülich GmbH, 52425 Jülich, Germany; Department of Biological Sciences, University of Alberta, Edmonton, Alberta, Canada; Institute of Biotechnology, National Chung Hsing University, Taichung 40227, Taiwan

**Keywords:** *Arabidopsis thaliana*, diurnal cycle, proteome, phosphoproteome, quantitative proteomics

## Abstract

Plant growth depends on the diurnal regulation of cellular processes, but it is not well understood if and how transcriptional regulation controls diurnal fluctuations at the protein-level. Here we report a high-resolution *Arabidopsis thaliana* (Arabidopsis) leaf rosette proteome acquired over a 12 h light : 12 h dark diurnal cycle and the phosphoproteome immediately before and after the light-to-dark and dark-to-light transitions. We quantified nearly 5000 proteins and 800 phosphoproteins, of which 288 fluctuated in their abundance and 226 fluctuated in their phosphorylation status. Of the phosphoproteins, 60% were quantified for changes in protein abundance. This revealed six proteins involved in nitrogen and hormone metabolism that had concurrent changes in both protein abundance and phosphorylation status. The diurnal proteome and phosphoproteome changes involve proteins in key cellular processes, including protein translation, light perception, photosynthesis, metabolism and transport. The phosphoproteome at the light-dark transitions revealed the dynamics at phosphorylation sites in either anticipation of or response to a change in light regime. Phosphorylation site motif analyses implicate casein kinase II and calcium/calmodulin dependent kinases among the primary light-dark transition kinases. The comparative analysis of the diurnal proteome and diurnal and circadian transcriptome established how mRNA and protein accumulation intersect in leaves during the diurnal cycle of the plant.

**Summary Statement:** The manuscript provides quantitative information of diurnal changes in the accumulation and phosphorylation of proteins in *Arabidopsis thaliana* rosettes grown in a 12 h photoperiod. The highly resolved time-scale of the datasets offer new proteome-level insights for future targeted studies.

## INTRODUCTION

Plant growth and biomass production are direct functions of the diurnal cellular carbon balance, which is regulated by a combination of light responses and the circadian clock. Light responses are triggered by a change in regime (i.e., presence or absence of light), while the circadian clock is comprised of transcriptional regulators that operate in anticipation of a change (e.g. transition from light to dark) and whose activities span the 24 hour (h) photoperiod (Nohales & Kay, 2016; Oakenfull & Davis, 2017; Seluzicki, Burko, & Chory, 2017). The core clock transcriptional regulators include CCA1/LHY, PRR5, PRR7 and PRR9, which form the morning loop, and TOC1, ELF3, ELF4 and LUX, which form the evening loop (Flis et al., 2015; Staiger, Shin, Johansson, & Davis, 2013). More than 30% of all Arabidopsis genes are regulated by the circadian clock at the transcript level (Blasing et al., 2005; Covington, Maloof, Straume, Kay, & Harmer, 2008). However, less is known about how the resulting diurnal transcriptome relates to protein abundance (Abraham et al., 2016; Choudhary, Nomura, Wang, Nakagami, & Somers, 2015; Graf et al., 2017) and post-translational protein modifications (Choudhary et al., 2015; Uhrig, Schlapfer, Roschitzki, Hirsch-Hoffmann, & Gruissem, 2019), both of which may also affect protein function at light-dark transitions and throughout the diurnal cycle. Transcript and protein abundance changes are often disconnected because changes in transcript levels show no corresponding change in protein abundance (Baerenfaller et al., 2012; Seaton et al., 2018). For example, this disconnect was found in the circadian clock mutants CCA1/LHY, PRR7/PRR9, TOC1 and GI (Graf et al., 2017) and for the variability in the timing of peak transcript and protein levels (translational coincidence) as a function of the photoperiod-dependent coordination between transcriptome and proteome changes (Seaton et al., 2018). Variable delays between peak transcript and protein abundance have implicated post-transcriptional regulation (e.g. splicing), translational regulation (e.g. translation rate) as well as post-translational regulation (e.g. protein phosphorylation) as possible mechanisms to explain the temporal differences in RNA and protein abundance. More recent studies of plant protein-level regulation have also found extensive variability in protein turnover (Li et al., 2017; Seaton et al., 2018), which adds further regulatory complexity because conventional quantitative proteome workflows cannot easily account for protein turnover. Although transcript and protein synthesis, stability and turnover all contribute to the coordination of transcript and protein abundance, how these mechanisms are integrated is currently not well understood. Insights into this regulatory complexity requires protein-level time-course experimentation and, in particular, an understanding of protein post-translational modifications (PTMs). To address this, we undertook a large-scale quantitative proteomics approach to determine the extent of diurnal protein abundance and/or phosphorylation changes in Arabidopsis rosette proteins over a 24h photoperiod.

Reversible protein phosphorylation is the most abundant PTM in eukaryotes (Adam & Hunter, 2018; Rao, Thelen, & Miernyk, 2014). In non-photosynthetic eukaryotes phosphorylation is found to modulate more than 70% of all cellular processes (Olsen et al., 2006), including the circadian clock itself (Robles, Humphrey, & Mann, 2017). The extent of regulatory protein phosphorylation events is likely similar in land plants as they have significantly larger kinomes (Lehti-Shiu & Shiu, 2012; Zulawski, Schulze, Braginets, Hartmann & Schulze, 2014) compared to humans, which encode 518 protein kinases (Manning, Whyte, Martinez, Hunter, & Sudarsanam, 2002). Conversely, both plants and humans have an equally comparable number of protein phosphatases (Kerk, Templeton, & Moorhead, 2008). However, most protein phosphatases require association with regulatory subunits to achieve their specificity (Moorhead et al., 2008; Uhrig, Labandera, & Moorhead, 2013), suggesting similar complexity in how plants manage protein dephosphorylation through a likely expansion of protein phosphatase regulatory subunits.

In plants, diurnal protein phosphorylation is regulated either in response to light, by the circadian clock (Choudhary et al., 2015), or both (Uhrig et al., 2019), while the clock itself is regulated by phosphorylation (Kusakina & Dodd, 2012; Uehara et al., 2019). Recent studies of the circadian phosphoproteome combining the analysis of a free-running cycle and the circadian clock mutants *elf4* (Choudhary et al., 2015) or CCA1-OX over-expression (Krahmer et al., 2019) have revealed temporally modified phosphorylation sites related to casein kinase II (CKII) and sucrose non-fermenting kinase 1 (SnRK1). SnRKs are likely involved in the regulation of the circadian phosphoproteome because the transcription of genes encoding multiple SnRK and calcinuerin B-like (CBL) interacting kinases (CIPK) was mis-regulated in the Arabidopsis circadian clock mutants *cca1/lhy1, prr7prr9, toc1* and *gi201* mutants at end-of-day (ED) and end-of-night (EN) (Graf et al., 2017). Similarly, studies quantifying changes in the phosphoproteome at ED and EN in Arabidopsis rosette leaves, roots, flowers, siliques and seedlings have revealed a large number of diurnally changing phosphorylation events corresponding to diverse protein kinase motifs (Reiland et al., 2009; Uhrig et al., 2019).

Considering the marked physiological and metabolic changes occurring at the light-dark (L-D) and dark-light (D-L) transitions (Annunziata et al., 2018; Gibon et al., 2009; Usadel et al., 2008), we performed a quantitative phosphoproteome analysis of proteins that are phosphorylated immediately before and after the L-D and D-L transitions during a 12 h light : 12 h dark photoperiod and asked how these phosphorylation events intersect with changes in protein abundance. Together, our systems-level quantitative analysis of the Arabidopsis thaliana rosette proteome and phosphoproteome over a 24h photoperiod provides new insights into diurnal protein and phosphorylation regulation.

## MATERIALS AND METHODS

Arabidopsis Col-0 wild-type plants were grown at the Forschungszentrum Jülich (Germany) in an environmentally controlled chamber (GrowScreen Chamber; https://eppn2020.plant-phenotyping.eu/EPPN2020_installations#/tool/30; Barboza-Barquero et al., 2015) under a 12 h light:12 h dark photoperiod and controlled conditions as described in Baerenfaller *et al.* (2012), including air temperature of 21°C during the day and 20°C during the night, air humidity of 70%, and an incident light intensity of ~220 mmol/m^2^/s at the plant level. Whole rosettes were harvested at 31 days after sowing (DAS) prior to flowering. Four whole rosettes were pooled into one sample and 4 biological replicates were collected at each time point except for ZT1, 3, 5 9 and 23, which had only 3 biological replicas for proteome analysis and AL_10, which had only 3 replicas for phosphoproteome analysis. For total proteome analyses, samples were taken every 2 h during 24 h, starting at Zeitgeber time 1 (ZT1, i.e. 1 h after lights turned on). For protein phosphorylation analyses, samples were 30 min before, 10 min after and 30 min after the L-D and D-L transitions. Samples were snap-frozen in liquid N2 and stored at −80°C until protein extraction.

### Proteome Analysis

#### Extraction and digestion

Samples were randomized before processing to avoid batch effects. Frozen rosettes were ground under liquid N2. Proteins were extracted from 100 mg of frozen powder per sample by adding 150 μl of extraction buffer (30 mM Tris-HCl pH 8.0, 4% SDS). Tubes were incubated in a shaker (Eppendorf) at 4°C at 1400 rpm for 30 min. Samples were centrifuged at 16000 *g* and 4°C for 30 min and the supernatant was transferred to a new tube. Protein concentration was estimated based on Bradford (Bradford, 1976) using the Bio-Rad Protein Assay reagent. Subsequently, DTT was added to a final concentration of 50 mM and proteins were reduced for 30 min on ice. For digestion, 140 μg of proteins were processed following the FASP method (Wisniewski, Zougman, Nagaraj, & Mann, 2009). Peptides were desalted using SPE C18 columns (Finisterre) and dried down in a SpeedVac concentrator.

#### Peptide fractionation

To increase proteome coverage, peptide samples were fractionated by hydrophilic interaction chromatography (HILIC) on an Agilent 1200 series HPLC system with a YMC-Pack Polyamine II 250 x 3.0 mm size column with 5 μm particle size and 120 Å pore size. Samples were dissolved in 100 μl Buffer A (75% ACN, 8 mM KH2PO4, pH 4.5) and separated with Buffer B (5% ACN, 100 mM KH2PO4, pH 4.5) at a flow rate of 500 μl/min with the following gradient: 0-7.5 min, 0% B; 7.5-37.5 min, 0-50% B; 37.5-42.5 min, 50-100% B; 42.5-47.5 min, 100% B. Following the separation the column was washed with 100% buffer A and re-equilibrated for 60 min. For each sample, the 27 automatically collected fractions were pooled into five fractions that were subsequently dried down in a SpeedVac concentrator. Each sample was then dissolved in 200 μl of 3% ACN, 0.1% TFA, desalted on SPE C18 columns (Finisterre) and again dried in a SpeedVac concentrator.

#### LC-MS analysis

Mass spectrometry queues were arranged to process comparable fractions in the same batch, with sample order randomized within each batch. Peptide samples were dissolved in 20 μl 3% ACN, 0.1% FA and spiked with internal retention time (iRT) standards (Biognosys) for chromatography quality control. LC-MS/MS shotgun analyses were performed on a Thermo Orbitrap Fusion instrument coupled to an Eksigent NanoLC Ultra (Sciex). Samples were separated on a self-packed reverse-phase column (75 μm x 150 mm) with C18 material (ReproSil-Pur, C18, 120 Å, AQ, 1.9 μm, Dr. Maisch GmbH). The column was equilibrated with 100% solvent A (0.1% FA in water). Peptides were eluted using the following gradient of solvent B (0.1% FA in ACN) at a flow rate of 0.3 μl/min: 0-50 min: 3-25% B, 50-60 min: 25-35% B, 60-70 min: 35-97% B, 70-80 min: 97% B, 80-85 min: 2% B. Mass spectra were acquired in a data-dependent manner. All precursor signals were recorded in the Orbitrap using quadrupole transmission in the mass range of 300-1500 m/z. Spectra were recorded with a resolution of 120000 (FWHM) at 200 m/z, a target value of 4e5 and the maximum cycle time set to 3 s. Data dependent MS/MS were recorded in the linear ion trap using quadrupole isolation with a window of 1.6 Da and higher-energy collisional dissociation (HCD) fragmentation with 30% fragmentation energy. The ion trap was operated in rapid scan mode with a target value of 1E4 and a maximum injection time of 250 ms. Precursor signals were selected for fragmentation with a charge state from + 2 to + 7 and a signal intensity of at least 5e3. A dynamic exclusion list was used for 30 s and maximum parallelizing ion injections was activated. The mass spectrometry proteomics data were handled using the local laboratory information management system (LIMS) (Türker et al., 2010)

### Phosphoproteome Analysis

#### Extraction

Whole rosette tissue from each time point was harvested and ground under liquid N2. From each biological replicate 200 mg of ground leaf material was weighed out under liquid N2. In addition to each biological replicate, 200 mg of samples containing equal weighted parts of each biological replicate and time-point were created as a reference sample (gold-standard) for downstream dimethyl labeling. All proteins were extracted in a 250 μl solution of 50 mM HEPES pH 8.0, 6 M urea, 2 M thiourea, 100 mM NaCl, 10 mM EDTA, 2 mM NaOV, 5 mM NaF, 50 μg/mL PhosSTOP (Roche). Samples were shaken at room temperature for 30 min at 1000 *g* with vortexing every 10 min. Extracts were then brought to pH 8.0 using triethylammonium bicarbonate (TEAB). Protein extracts were then reduced for 30 min with 10 mM DTT, followed by alkylation with 30 mM iodoacetamide for 1 h. Extracts were clarified to separate soluble and insoluble fractions. The insoluble fraction was re-suspended in 300 μL 60:40 buffer containing 60% MeOH: 40% 50 mM TEAB pH 8.0 followed by shaking at 1000 rpm (Eppendorf tabletop) for 2.5 h. The protein concentration of the soluble fraction was then measured using the Bradford protein assay (Bradford, 1976). An amount of 1 mg of soluble protein from each sample was then diluted with 1 vol. of 50 mM TEAB and then water was added to a total volume of 1.2 ml and a final urea/thiourea concentration of 1.2 M. The soluble fraction was then digested for 20 h at 37°C using a 1:50 ratio of trypsin (Promega) to extracted protein while gently shaking. Each insoluble fraction was digested by 0.5 μg chymotrypsin and 1 μg trypsin at 37°C for 20 h shaking at 600 rpm (Eppendorf tabletop). Digestion reactions were stopped using TFA to a final concentration of 0.5%. The insoluble fractions were centrifuged for 10 min at 20000 *g* at room temperature and the supernatant removed. The supernatant was then dried and re-suspended in desalting buffer comprised of 3% ACN / 0.1% TFA. The soluble fraction and the supernatant from the insoluble fraction were desalted using SPE C18 columns (Finisterre) and dried in a SpeedVac concentrator.

#### Dimethyl labeling and phosphopeptide enrichment

Total peptide fractions from each experimental (light label) and gold-standard (heavy label) sample were labeled according to Boersema *et al.,* (Boersema, Raijmakers, Lemeer, Mohammed, & Heck, 2009). Heavy and light samples were then mixed 1:1 and desalted prior to phosphopeptide enrichment using TiO2. Phosphopeptide enrichment was performed using TiO2 heavy and light dimethyl-labelled phosphopeptides as previously described (Zhou et al., 2011).

#### LC-MS

Phosphorylated peptide samples were analyzed using a Q Exactive Orbitrap mass spectrometer (Thermo Scientific). Dissolved samples were injected using an Easy-nLC 1000 system (Thermo Scientific) and separated on a self-made reverse-phase column (75 μm x 150 mm) packed with C18 material (ReproSil-Pur, C18, 120 Å, AQ, 1.9 μm, Dr. Maisch GmbH). The column was equilibrated with 100% solvent A (0.1% formic acid (FA) in water). Peptides were eluted using the following gradient of solvent B (0.1% FA in ACN): 0-120 min, 0-35% B, 120-122 min, 35-95% B at a flow rate of 0.3 μl/min. High accuracy mass spectra were acquired in data-depended acquisition mode. All precursor signals were recorded in a mass range of 300-1700 m/z and a resolution of 70000 at 200 m/z. The maximum accumulation time for a target value of 3e6 was set to 120 ms. Up to 12 data dependent MS/MS were recorded using quadrupole isolation with a window of 2 Da and HCD fragmentation with 28% fragmentation energy. A target value of 1e6 was set for MS/MS using a maximum injection time of 250 ms and a resolution of 70000 at 200 m/z. Precursor signals were selected for fragmentation with charge states from +2 to +7 and a signal intensity of at least 1e5. All precursor signals selected for MS/MS were dynamically excluded for 30 s.

### Quantitative analysis and bioinformatics

#### Total proteome

Label-free precursor (MS1) intensity based quantification was performed using Progenesis QI for Proteomics (version 2.1, www.nonlinear.com) to quantify total proteome changes. Briefly, for each individual fraction, automatic alignment was reviewed and manually adjusted before normalization. From each Progenesis peptide ion (default sensitivity in peak picking) a maximum of the top five tandem mass spectra per peptide ion were exported as a Mascot generic file (*.mgf) using charge deconvolution and deisotoping option and a maximum number of 200 peaks per MS/MS. Searches were done in Mascot 2.4.1 (Matrix Science) against a decoyed (reversed) Arabidopsis protein database from TAIR (release TAIR10) concatenated with a collection of 261 known mass spectrometry contaminants. Precursor ion mass tolerance was set to 10 ppm and the fragment ion mass tolerance was set to 0.6 Da. The following search parameters were used: trypsin digestion (1 missed cleavage allowed), fixed modifications of carbamidomethyl modified cysteine and variable modifications of oxidation of methionine, deamidation of asparagine and glutamine, and acetylation of protein N terminal peptides. Mascot searches were imported into Scaffold 4.2.1 (Proteome Software). The following thresholds were applied: peptide FDR ≤ 5, protein FDR ≤ 10, 1 minimum peptide. Spectrum reports were imported again into Progenesis. After this, individual fraction analyses were combined into the full quantitative Progenesis experiment. From this, quantitative peptide values were exported for further processing. Only peptides that could be unambiguously assigned to a single protein (gene model annotation) were kept for quantification. A Hi-4 strategy (Grossmann et al., 2010) was applied to obtain protein quantitative values. Proteins with 2 or more peptides assigned were considered as quantifiable. Following these criteria, the final protein level FDR was estimated at 0.013.

#### Phosphoproteome

Quantification of changes in identified phosphopeptides was performed using MaxQuant (version 1.3.0.5) with default settings and the following modifications: fixed peptide modification by carbamidomethylation of cysteines and variable peptide modifications by phosphorylation of serine, threonine and tyrosine, and oxidation of methionine, and false discovery rate (FDR) tolerances of ≤ 0.05 (protein) and ≤ 0.01 (peptide). MaxQuant outputs were subsequently filtered for phosphopeptides with a phosphorylation site probability score ≥ 0.8 and presence in at least 2 of 3 (AL_10) or 2 of 4 biological replicates and 2 of 3 time-points for each light transition.

#### Data Analysis

Significant fluctuations in protein abundance and phosphopeptides were determined using an ANOVA analysis: total proteome (P value ≤ 0.05 and Fold-change (FC) ≥ 1.5) and phosphoproteome (P value ≤ 0.05). The significantly changing proteome was subjected to cluster analysis using GProX (Rigbolt, Vanselow, & Blagoev, 2011). Six clusters were generated in an unsupervised clustering manner based on the fuzzy c-means algorithm. Significantly changing proteins and phosphoproteins were subjected to gene set enrichment analysis (GSEA) using the SetRank algorithm relative to the identified proteome and phosphoproteome, respectively (Simillion, Liechti, Lischer, Ioannidis, & Bruggmann, 2017). Enrichment was calculated for all the available databases included in the SetRank R package. Only terms with a size ≥ 2 were considered (gene set size ≥ 2). For each protein cluster, a SetRank corrected P value ≤ 0.01 was applied as threshold. For phosphoproteins changing at the L-D or D-L transition, a SetRank corrected P value ≤ 0.01 and an FDR ≤ 0.05 were applied. To test for significantly non-changing proteins at the transitions to light, (i.e., at dawn, ZT23 to ZT1, and dusk, ZT11 to ZT13), a TOST equivalence test (equivalence R package) was applied with an ε = 0.4. Significance threshold was P value ≤ 0.05. The mass spectrometry proteomics data have been deposited to the ProteomeXchange Consortium via the PRIDE partner repository. Data are available via ProteomeXchange with identifier PXD007600.

#### Additional Analyses

To compare protein and mRNA profiles, mRNA data generated by the Alison Smith laboratory was obtained from the Diurnal database (http://diurnal.mocklerlab.org; Mockler et al., 2007). For this, we restricted the analysis to the information from LDHH_SM and LDHH_ST. Data was standardized to plot both protein and mRNA data in the same graph. Predicted subcellular localization of all changing proteins and phosphoproteins was performed using the consensus subcellular localization predictor SUBAcon (suba3.plantenergy.uwa.edu.au) (Tanz et al., 2013). String DB network analyses were undertaken using both proteome and phosphoproteome data. String DB analyses were performed in Cytoscape using the String DB plugin stringApp (Szklarczyk et al., 2017). A minimum correlation coefficient of 0.5 was used along with a second layer of 5 additional nodes to anchor each network to better infer network connectedness.

#### JTK Analyses

To compare diurnal protein fluctuations to free running circadian clock fluctuations published by Krahmer et al. (2019; dataset PXD009230 available at ProteomeXchange) we performed an equivalent analysis using the JTK cycle to identify proteins cycling with 22 or 24 h period (Hughes, Hogenesch, & Kornacker, 2010). The exact loading script JTK_analysis.zip is available upon request. JTK_cycle fits data of many entities (here protein abundances) to a cosine function model, and estimates a P value for the accuracy of the model for every protein permutation of the dataset (resulting in the ADJ.P values). Further it applies a Benjamini Hochberg correction for multiple testing resulting in q-values (resulting in the BH.Q values). The data was then used to produce Figure 3B, C and D. Proteins identified to fluctuate were normalized such that they fluctuate around a median of 0 with maximal amplitudes of 2. Transcriptome data from Diurnal DB (http://diurnal.mocklerlab.org; Mockler et al., 2007) was used to determine if the associated transcripts were also fluctuating, and if so, when. For this, we restricted the analysis to the information from LDHH_SM and LDHH_ST. To estimate a confidence interval for the relative expression or protein level errors, their relative levels were compared to the theoretical cosine function at the same timepoint. Based on all errors, irrespective of the exact timepoint, a 99% confidence interval was computed.

## RESULTS AND DISCUSSION

### Dynamics of the Arabidopsis diurnal proteome and phosphoproteome

Using proteotypic peptides, we performed a label-free quantitative proteomics analysis of the diurnal proteome. Here, we identified 7060 unique proteins, of which we were able to quantify 4762 proteins with two and more proteotypic peptides over the 24h time-course (Supplemental Figure 1; Table 1; Supplemental Table 1). Statistical analysis showed that 288 of these proteins were significantly changing in abundance (ANOVA P value ≤ 0.05, FC > 1.5); Table 1; Supplemental Table 2), suggesting that a portion (~6%) of the quantified proteome is dynamically regulated over the course of a day. Additionally, using a dimethyl-labeling approach, we identified a total of 2298 phosphopeptides (Supplemental Figure 1; Supplemental Table 3), of which 1776 had a phosphorylation site probability score ≥ 0.8. We were able to quantify 1056 of these phosphopeptides (present in at least 2 biological replicates and in 3 out of 3 time points for each transition; Table 1), which corresponded to a total of 1803 identified phosphorylation sites. Of these, 253 (14%) represented newly identified phosphorylation sites when compared to the compendium of 79,334 known phosphorylation sites (PhosPhat 4.0; Heazlewood et al., 2008) and a total of 271 phosphopeptides on 226 proteins (~26% of all quantified phosphopeptides) significantly changed in abundance (ANOVA P value ≤ 0.05) at either the D-L, L-D or both transitions (Table 1; Supplemental Table 4).

**Table 1:**
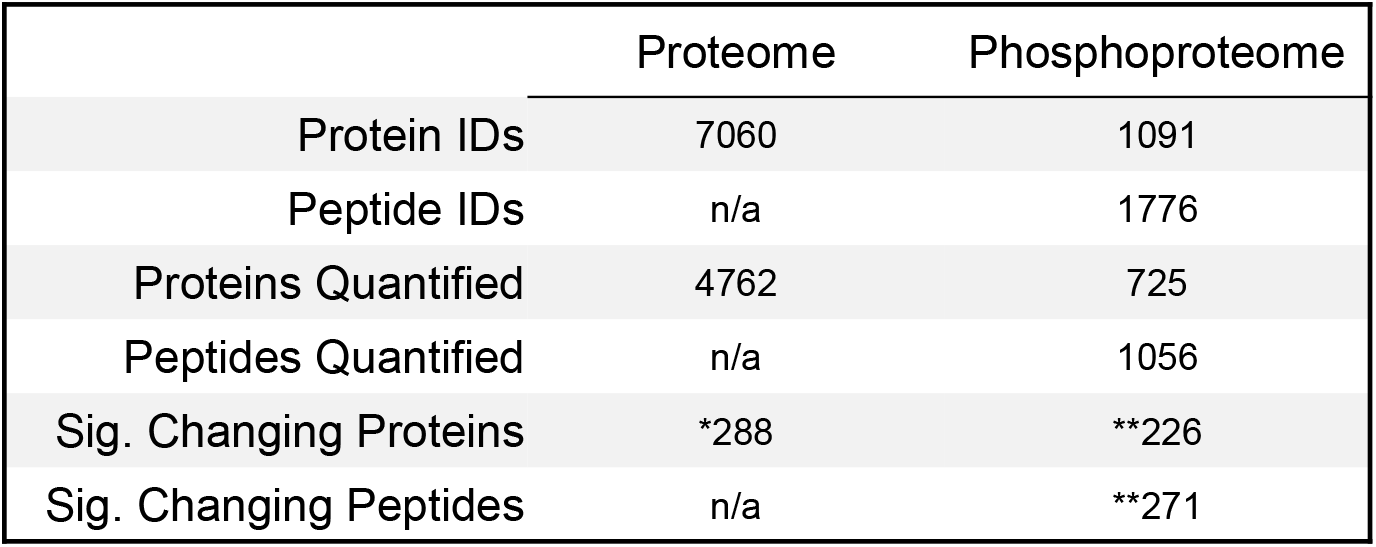
Proteome and phosphoproteome coverage. Summary of the identified, quantified and significantly changing diurnal proteins, phosphopeptides and phosphoproteins. Quantification confidence thresholds are shown for the proteome (proteins identified by ≥ 2 proteotypic peptides) and the phosphoproteome (site probability score ≥ 0.8) quantified in ≥ 3 biological replicates for each time point of the diurnal cycle and for each of the three time-points at the L-D and D-L transitions. The significance thresholds are shown for the proteome (FC ≥ 1.5; ANOVA P value ≤ 0.05) and the phosphoproteome (ANOVA P value ≤ 0.05). Application of proteome and phosphoproteome significance thresholds are denoted by a single (*) and double (**) asterisks, respectively.

### Most proteins with diurnal changes in abundance fluctuate independently of their transcript levels and belong to specific functional networks

To clarify which cellular and physiological processes possess protein abundance dynamics, we grouped all significantly changing proteins with similar accumulation profiles into clusters and then subjected these clusters to gene set enrichment analysis (GSEA). Not all clusters exhibited classic cosine dynamics, but instead exhibited complex profiles at specific times of day. Each of the resulting six clusters (CL1 – CL6) is enriched for proteins involved in specific processes (P value ≤ 0.01, gene set size ≥ 2) (Figure 1A-B; Supplemental Data 1 - 6). Cluster CL1 is enriched in proteins involved in RNA splicing that decrease before dawn, while CL2 is enriched in proteins that peak early in the light period and have roles in nitrogen metabolism, iron homeostasis, responses to gravity and chloroplast stroma protein import. CL5 contains proteins with peak abundance before dawn and lower abundance before dusk that have specific functions in aerobic respiration and proteasome complex formation, while proteins in CL3 have functions in membrane-related processes and ribosome biogenesis. The CL3 abundance profile is complex with a sharp minimum during the second half of the light period that is also found at the transcript level for selected proteins in this group. Cluster CL4 shows increasing levels during the first hours of the day, followed by a reduction until the end of the day, while levels are stable during night. CL4 is enriched for proteins involved in nitrogen metabolism and photosynthesis, which are required for light-dependent carbon assimilation to support growth. CL6 exhibits a similar pattern as CL4, but seems to be shifted by 4 to 6 hours so that the peak protein levels peak at dusk. CL6 is enriched for proteins involved in metabolic and RNA-related processes that indicate a systemic change in the plant cell environment.

**Figure 1:**
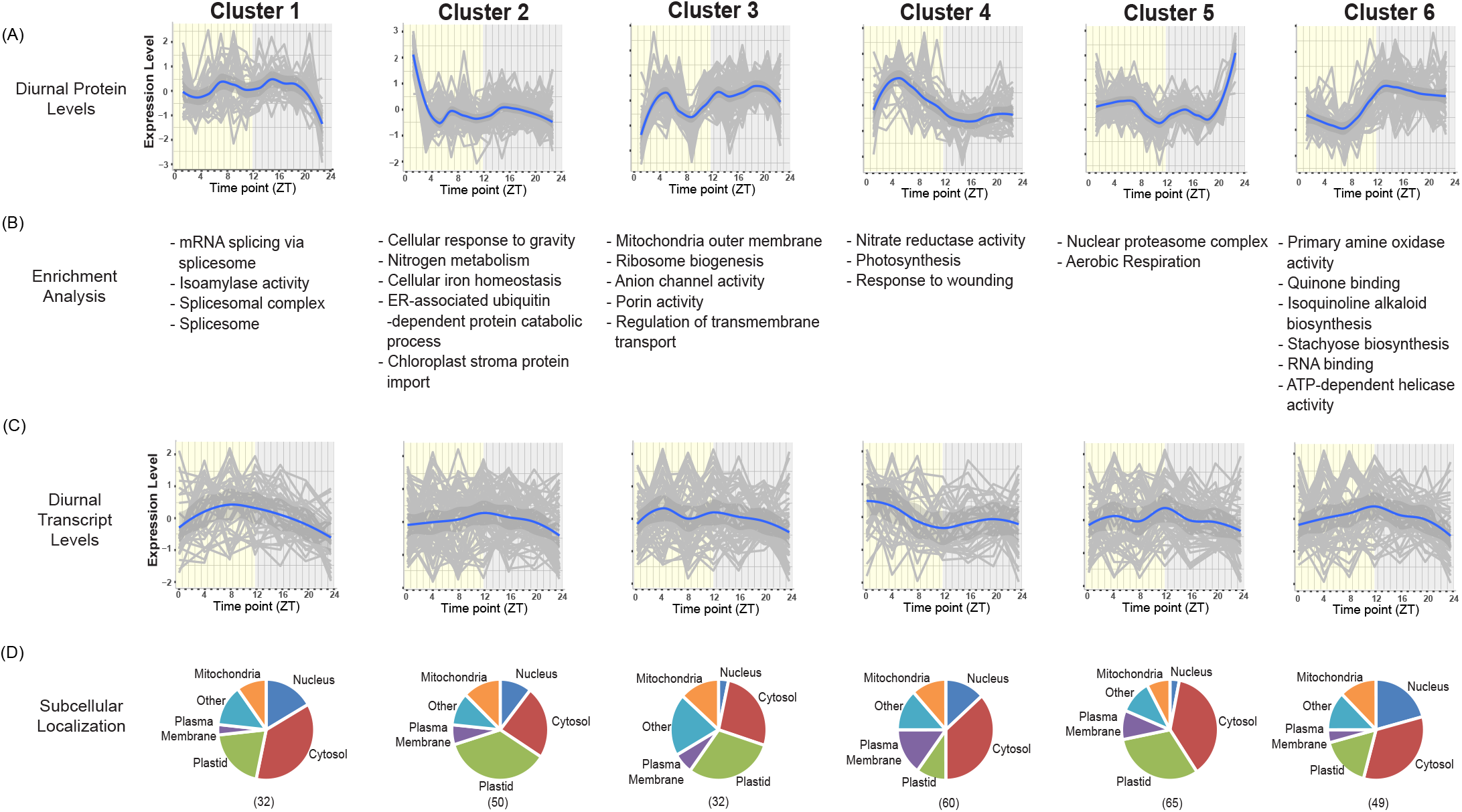
Analysis of the diurnal proteome: clustering, enrichment analysis and subcellular localization. (A) Significantly changing proteins (FC ≥ 1.5, ANOVA P value ≤ 0.05, ≥ 2 peptides) were subjected to an unsupervised clustering analysis (GProX; http://gprox.sourceforge.net) resolving 6 protein clusters. Y- and X-axis depict standardized expression level and harvest time (Zeitgeber time; ZT), respectively. Median expression is depicted in blue. (B) Term enrichment analysis of significantly changing proteins using SetRank (P value ≤ 0.01, size ≥ 2). (C) Standardized diurnal transcript expression level of each corresponding clustered protein (Log10). Median expression is depicted in blue. Transcript expression level was obtained from Diurnal DB (http://diurnal.mocklerlab.org/). (D) *In silico* subcellular localization analysis of significantly changing proteins using SUBAcon (SUBA3; http://suba3.plantenergy.uwa.edu.au). Bracketed numbers represent the number of proteins per cluster.

We then compared the proteins in CL1 to CL6 with their corresponding transcript expression profiles using transcriptome data generated from whole Arabidopsis rosettes grown and harvested in comparable conditions and at similar time-points (Figure 1C). This revealed that the dynamics of CL1 to CL6 protein changes are not strictly correlated with the diurnal abundance changes of their transcripts (Figure 1C; Supplemental Data 1-6), as has been found in multiple other studies (Baerenfaller et al., 2012; Abraham et al., 2016; Graf et al., 2017; Seaton et al., 2018). We then determined the subcellular compartmentalization of proteins in each cluster using the consensus localization predictor SUBAcon (SUBA3; http://suba3.plantenergy.uwa.edu.au; Figure 1D) (Tanz et al., 2013). Most clusters exhibited a similar distribution of localizations with the exception of CL4, which had an expanded complement of cytosolic and plasma membrane proteins coupled with a decrease in plastid-targeted proteins.

To determine connections between proteins with changing abundances, we next built functional association networks for each cluster using STRING-DB (http://string-db.org; Figure 2). STRING-DB scoring and Cytoscape visualization allowed us to estimate association confidence between protein nodes, while subcellular localization information resolved co-localized nodes. Second level nodes not found in our data were also included to anchor the network and help depict broader relationships between the significantly changing proteins. Although such anchoring nodes do not change themselves, abundance changes of their interaction partners may impact the behavior of these nodes. This analysis strategy resolved multiple protein hubs within each cluster that have variable degrees of interconnectedness to the depicted biological processes, with some processes complementing those enriched by GSEA (Figure 1B). Proteins with no known connections above the set association threshold were removed from the network. Using our STRING-DB analysis approach we defined network structures for proteins belonging to: RNA splicing (CL1) and processing (CL6; RNA helicases and binding proteins), chloroplast-related processes (CL4 and 5, light detection; CL1 and CL5, carbohydrate/starch metabolism; CL2, redox regulation), cell metabolism (CL4, nitrogen and fatty acid metabolism), secretion and intracellular transport (CL2), cell wall biosynthesis (CL5) as well as cytosolic (CL1, 3 and 5), mitochondrial (CL3) and plastidial (CL4 and 5) protein translation (Figure 2). Taken together, our GSEA and association network analyses provide new process- and protein-level information for when (time of day), where (subcellular compartment) and how (cellular processes) plants operate over a 24h photoperiod. This data is essential for a more precise understanding of molecular plant cell regulation.

**Figure 2:**
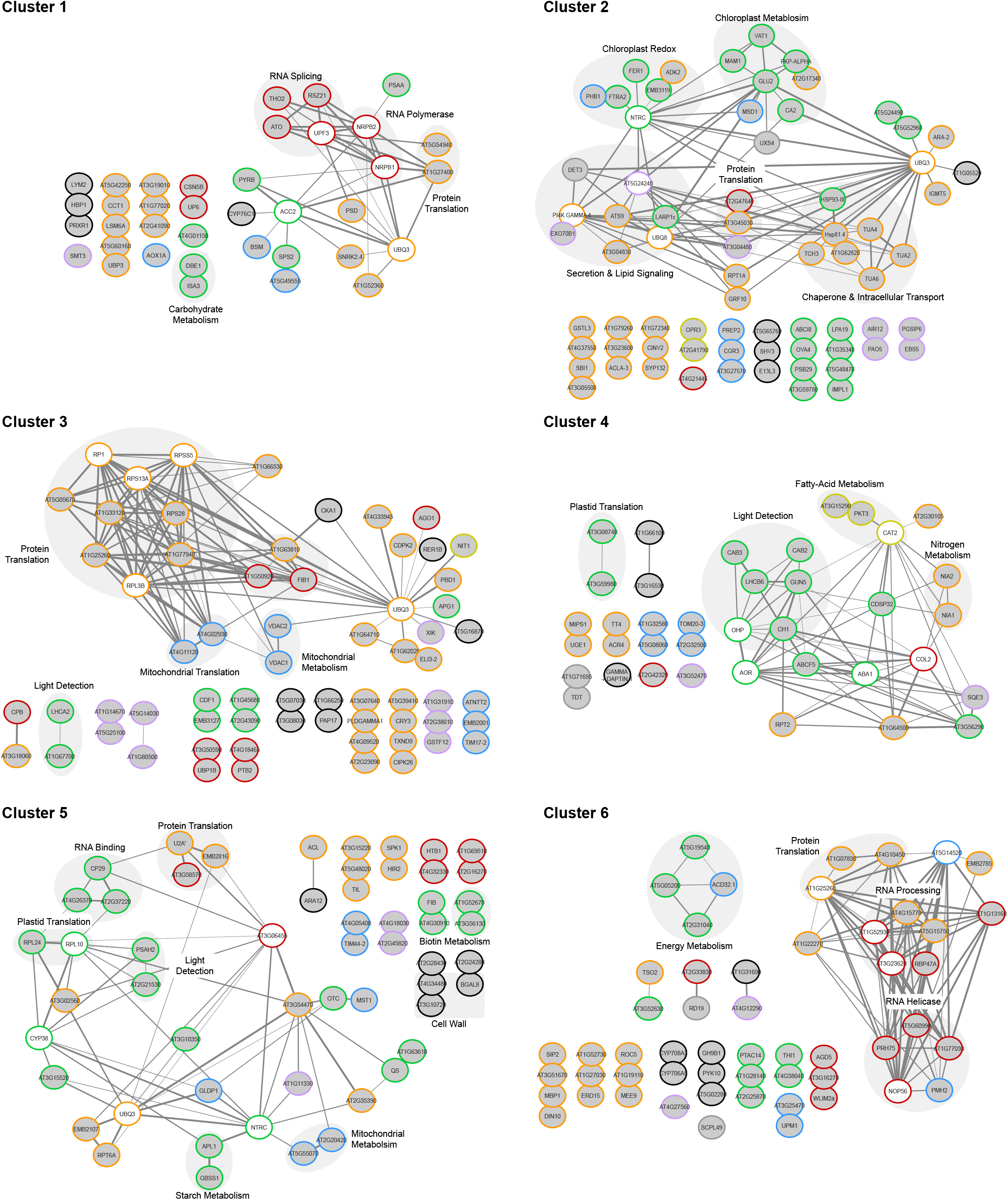
Interaction networks of the diurnal proteome. An association network analysis using STRING-DB (https://string-db.org/) of statistically significant diurnally changing proteins was performed using the generated unsupervised clusters shown in Figure 1. Edge thickness indicates confidence of the connection between two nodes (0.5 - 1.0). Changing proteins (grey circles) are labeled by either their primary gene annotation or Arabidopsis gene identifier (AGI). The colored outline of each node represents the *in silico* predicted subcellular localization of this protein (SUBAcon; suba3.plantenergy.uwa.edu.au). Nucleus (red), cytosol (orange), plastid (green), mitochondria (blue), plasma membrane (purple), peroxisome (dark yellow), endoplasmic reticulum/golgi/secreted (black) are depicted. A second layer of STRING-DB identified proteins (white nodes) not found in each respective significantly changing protein cluster was used to highlight the interconnectedness of proteins in the cluster. Multiple nodes encompassed by a labelled grey circle represent proteins involved in the same cellular process.

### The influence of the circadian clock on diurnal fluctuations of proteins is limited

To determine if the significant changes we measured in the diurnal proteome could be controlled by the circadian clock, we next compared our data to a quantitative proteomics dataset acquired under free-running (continuous light) conditions (Krahmer et al., 2019). Our dataset of 4762 quantified proteins contains 1800 of the 2038 proteins (88%) reported by Krahmer et al. (2019), allowing us to directly compare proteome results between studies (Supplemental Data 7). To avoid identification of differences based on the fact that the quantitative proteome analysis described above and the JTK_cycle analysis used by Krahmer et al. (2019) differ in their methods, we also performed a JTK_cycle analysis to identify proteins cycling with a 22 or 24 h period (Hughes et al., 2010). Unlike our previous analysis of diurnal proteome fluctuations, which identified 288 significantly changing proteins regardless of cycling preconditions, the JTK_cycle analysis approach aims to elucidate proteins exhibiting diurnal fluctuations in the form of a cosine function, and correspondingly evaluates how well these changes in abundance fit with this expected cosine behavior. JTK_cycle analysis estimates goodness of fit based on shuffling of protein values leading to a P value. It then uses a Benjamini Hochberg correction to correct for multiple testing. In accordance with the analysis approach of Krahmer et al. (2019), we identified a total of 147 fluctuating proteins prior to multiple testing correction, which is comparable to the 211 found to fluctuate under continuous light conditions by Krahmer et al. (2019). Upon correcting for multiple testing, our JTK_cycle analysis revealed a total of 21 proteins to exhibit a significant fluctuation in abundance, of which 3 demonstrated a similar pattern under continuous light conditions (Figure 3A). Using the statistically relevant proteins only, our study and Krahmer et al. (2019) find 3 proteins to fluctuate in both studies, one only in L-D conditions and 7 only in continuous light. The fact that of these 11 proteins only 10 have significant JTK-cycle fluctuations in continuous light (i.e., free-running condition), suggests that they are under circadian control, although additional proteome analysis of normal photoperiods prior to free-running conditions is needed to substantiate this possibility. Here, we find alpha-crystallin domain 32.1 (ACD32.1; AT1G06460) to fluctuate at the protein-level independent of the circadian clock. ACD32.1 was previously shown to be regulated diurnally at the transcript level in continuous light (Covington et al., 2008), but it did not fluctuate in the proteome data of Krahmer et al., 2019. ACD32.1 is a peroxisome-targeted chaperone protein (Pan et al., 2018) implicated in the suppression of protein aggregation (Ma, Haslbeck, Babujee, Jahn, & Reumann, 2006). We find ACD32.1 to peak in abundance immediately after dark, suggesting a potential need for peroxisomal protein stability in the dark to maintain peroxisome functions required for plant growth, including fatty acid oxidation (Pan et al., 2018).

**Figure 3:**
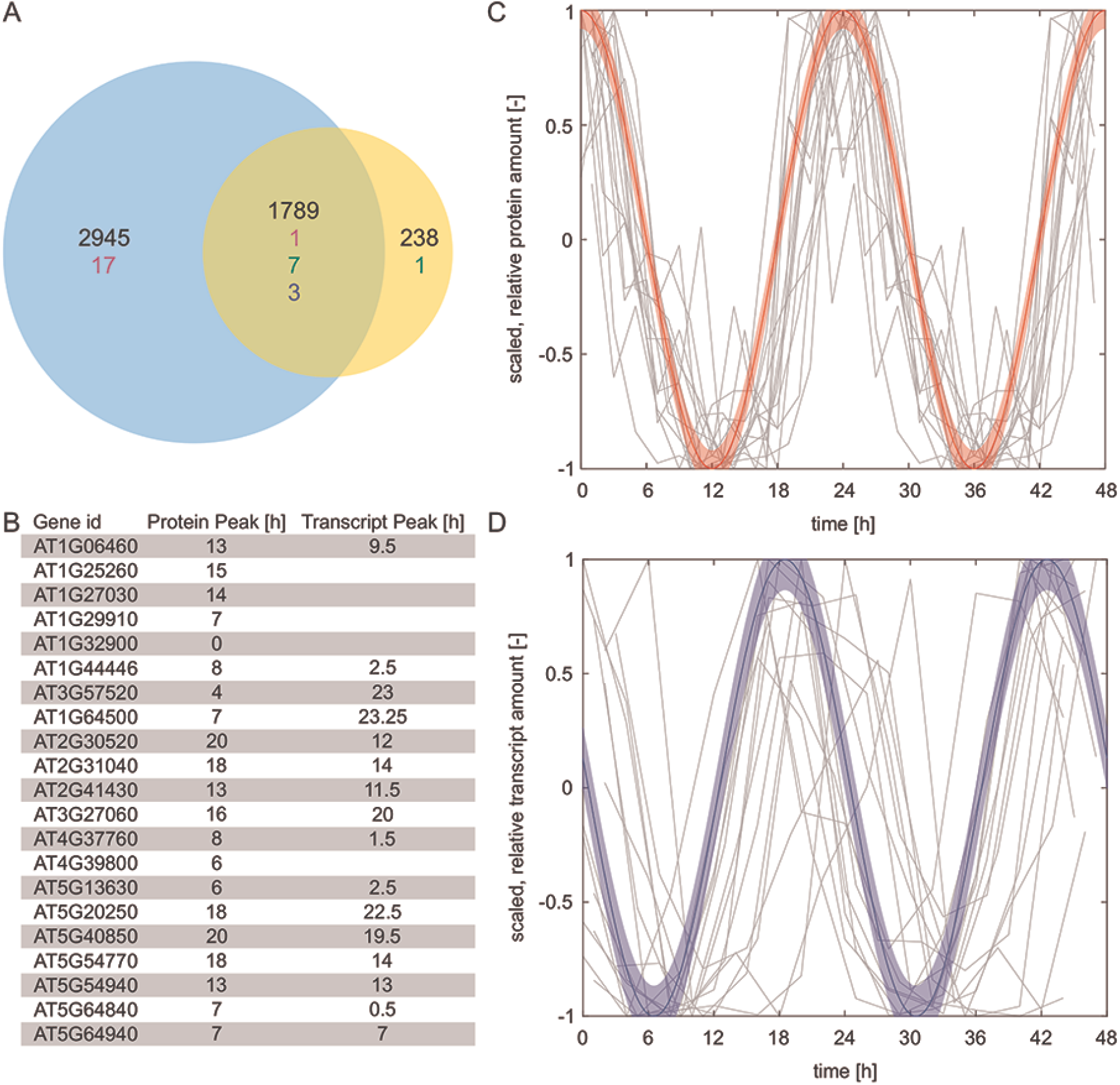
Comparative analysis of diurnal proteome to free-running circadian proteome (Krahmer *et al.,* 2019). (A) Number of proteins measured in this study (blue circle) and Krahmer *et al.* (2019) (orange circle). Number of stable proteins (black), fluctuating proteins in our study only (magenta), Krahmer *et al.* (2019) only (green) and both studies (blue). (B) Table of 21 proteins that show significant (B.Q) fluctuation using JTK with their respective peak time period for protein and transcript levels (Diurnal DB, http://diurnal.mocklerlab.org/). (C and D) Normalized (Median = 0, Amplitude of 2) protein levels of 15 proteins both fluctuating in protein and transcript levels (gray lines) shifted to peak at time zero for protein levels in (C) and transcript levels in (D). Protein data was plotted twice to visualize a 48 h timeframe. The theoretical cosine functions with associated 99% confidence interval for protein levels (C, red) and transcript levels (D, blue) are shifted by 5.5 h.

Given that the expression of many genes fluctuate at the transcript level, it is unexpected that such a low number of proteins exhibit rhythmic changes in protein abundance. For example, of the 22641 diurnal gene expression profiles stored in the Diurnal Database v2.0 (http://diurnal.mocklerlab.org/), 40.6% (9197) showed fluctuating transcripts in conditions that are comparable to ours (see Materials and Methods). Of our 4762 quantified proteins, gene expression profiles for 4468 were also present in the Diurnal Database and 2253 showed fluctuating transcript levels. Of our 4762 proteins we detected and quantified during the diurnal cycle, gene expression profiles for 4468 proteins were also found in the Diurnal Database and the transcripts for 2253 of these proteins had oscillating accumulation pattern. Of these oscillating transcripts, only 6.2% (140) had proteins that also showed peaks in abundance. For the remaining 2215 transcripts that did not oscillate, 4.0% (88) still had proteins with peaks in their abundance, indicating that there is no stringent relationship between transcript oscillation and protein peak abundance.

To see if the fluctuating proteins that we did find are potentially explained by fluctuating transcripts, we searched the Diurnal Database for the genes encoding the 21 proteins showing a significant JTK_cycle change in protein abundance (Figure 3A, magenta and blue) and found 18 of the genes. Of these 18, 15 were identified to possess diurnal changes in transcript abundance. For these 15 transcript-protein pairs, neither the protein nor the corresponding transcript levels were peaking at a specific Zeitgeber time (Figure 3B) and thus, these genes are likely regulated independently of each other. However, when comparing the patterns of individual pairs, normalizing for the transcript peak time, there was typically a median delay of 5.5 h between the peak transcript and peak protein (Figures 3C and D). Since such a shifted dependency of transcript and protein expression pattern is rare in our proteome dataset, its biological significance needs to be investigated further.

Together, while our diurnal proteome analysis revealed 288 proteins in different clusters that change in their abundance at different time intervals during the diurnal cycle, proteins for which their abundance changes follow a cosine function seem to be few when measured across the whole Arabidopsis rosette. The identification of only a single highly significant JTK-cycling protein in our diurnal proteome dataset is unexpected, but it is consistent with the limited fluctuations of proteins reported for measured proteomes of Arabidopsis wild-type and circadian clock mutants growing in free-running cycles of continuous light (Choudhary et al., 2015; Krahmer et al., 2019). This low number of cycling diurnal proteins could be a consequence of the stringency of the JTK_Cycle analysis, which only tests for periodical protein changes following a cosine function, similar to the oscillating fluctuations of a large number of mRNAs regulated by the circadian clock in Arabidopsis and animals (Doherty and Kay, 2010). Thus, in Arabidopsis rosettes the diurnal abundance of most measured proteins does not seem to be affected by the circadian clock or regulated in concert with oscillating mRNA levels, which has also been found in growing Arabidopsis leaves at fewer diurnal timepoints (Baerenfaller et al., 2012). Seedling proteins have turnover rates ranging from log2*k* –4 to –7 (Fan, Rendahl, Chen, Freund, Gray, Cohen & Hegeman, 2016) and the median degradation rate of proteins in growing Arabidopsis leaves is ~0.11d^−1^, but several proteins involved in protein synthesis, metabolic processes or photosynthesis have high degradation rates ranging from 0.6 up to 2.0 d^−1^ (Li, Nelson, Solheim, Whealan & Millar, 2017). Some of the fluctuating proteins in CL1–6 (Figure 1) that we identified in the diurnal proteome fall into categories of proteins with high degradation rates, including proteins in ribosome biogenesis in CL3 (Figure 1) that contribute to the replacement of the leaf cytosolic ribosome population (Salih, Duncan, Li, Troesch & Millar, 2020). For these proteins, oscillating mRNAs could contribute to the translational regulation of their changing accumulation (Missra, Ernest, Lohoff, Jia, Satterlee, Ke & von Arnim, 2015), also in case of the 15 mRNAs and proteins whose peaks are shifted by 5.5 h (Figures 3C and D). This does not exclude that oscillating mRNAs also contribute to the regulation of non-fluctuating proteins if the degradation and synthesis rates of these proteins are changing during the diurnal cycle. In Arabidopsis, there is increasing evidence of diurnal and photoperiodic dynamics of mRNA translation (Mills, Engatin and von Arnim, 2018; Seaton et al., 2018). If dynamic regulation of protein degradation and synthesis is coupled to differential ribosomal loading of oscillating mRNAs, this would result in stable diurnal protein levels. At present, our diurnal proteome dataset cannot distinguish between these scenarios, but it establishes an important framework for investigating the role of protein degradation and synthesis in circadian and diurnal protein level regulation in more detail.

### Analysis of light-dark transitions in a diurnal cycle reveal dynamic fluctuation in the Arabidopsis phosphoproteome

Protein phosphorylation is often associated with changing environmental conditions (Li et al., 2017; S. Zhang et al., 2019; Zhao et al., 2017). Therefore, we examined time-points before (30 min) and after (10 min, 30 min) the D-L and L-D transitions for changes in the phosphoproteome (Supplemental Figure 1). We identified 1776 phosphopeptides from 1091 proteins (phosphorylation site probability score ≥ 0.8) and quantified 1056 of these phosphopeptides from 725 proteins at the two light transitions (Table 1, Supplemental Table 3). We found that 176 phosphopeptides from 153 proteins at the D-L transition and 164 phosphopeptides from 144 proteins at the L-D transition had significant changes in abundance (Supplemental Figure 2 and 3; Supplemental Table 4). We then benchmarked the quality of our dataset by querying it for proteins known to be diurnally regulated by protein phosphorylation (Supplemental Table 5). This revealed phototropin 1 (PHOT1), phosphoenolpyruvate carboxylase (PEPC), nitrate reductase (NIA1 and NIA2) and CF1 ATP synthase. Phototropin 1 is phosphorylated in the light (Sullivan, Thomson, Kaiserli, & Christie, 2009; Sullivan, Thomson, Lamont, Jones, & Christie, 2008), while the NIA1, NIA2 and the CF1 ATP synthase beta-subunit are phosphorylated in the dark (Kanekatsu, Saito, Motohashi, & Hisabori, 1998; Lillo, Meyer, Lea, Provan, & Oltedal, 2004; G. Moorhead et al., 1999; Reiland et al., 2009). Our quantitation of NIA1 and 2 protein phosphorylation changes across time-points revealed that NIA2 was more rapidly dephosphorylated on Ser^534^ at the D-L transition than NIA1, potentially relating to regulatory differences between NIA1 and 2. Additionally, we found a new NIA2 phosphorylation site at Ser^63^ with opposing diurnal changes in phosphorylation at the same transition (Supplemental Figure 4). Both the rate of NIA1 and 2 phosphorylation as it relates to nitrate reduction and the new phosphorylation site require additional characterization that is beyond the scope of this study.

We next performed a GSEA of all significantly changing phosphoproteins (P value ≤ 0.01, FDR ≤ 0.05, gene set size ≥ 2) at each transition. Enriched biological processes at the D-L transition include phosphoproteins involved in light detection, nitrogen metabolism, cell wall-related processes and phosphorylation signaling, while phosphoproteins identified at the L-D transition are involved in light detection, vesicle-mediated transport, auxin signaling and nucleus organization (Table 2). We then generated a hierarchical heatmap of the phosphopeptides to identify clusters of proteins at each light transition with similar phosphorylation dynamics (Supplemental Figure 2 and 3). When compared to datasets of phosphorylated proteins previously identified in Arabidopsis growing under free-running cycle conditions (Choudhary et al., 2015; Krahmer et al., 2019), or at the ED and EN time-points of a 12-hour photoperiod (Reiland et al., 2009; Uhrig et al., 2019), our data reveals new proteins that have diurnal changes in their phosphorylation status and also novel information about the rate at which these phosphorylation events are occurring and disappearing (Supplemental Figure 2 and 3). For example, the L-D cluster I has phosphoproteins involved in nitrogen metabolism and the cell cycle (AD10 and AD30) and the L-D cluster III (BD30) has phosphoproteins involved in plastid organization (Supplemental Figure 2). In contrast, the D-L cluster II (AL10 and AL30) has phosphoproteins involved in central and carbohydrate metabolism (Supplemental Figure 3). Interestingly, parallel phosphorylation changes in L-D cluster I occur on proteins involved in nitrogen metabolism and the cell cycle. Nitrogen is acquired by plants primarily in the form of nitrate or ammonium, and is an essential macronutrient for plant growth. Nitrate signaling is linked to cell cycle progression through the TEOSINTE BRANCHED 1/ CYCLOIDEAPROLIFERATING CELL FACTOR 20 (TCP20) – NIN-LIKE PROTEIN 6/7 (NLP6/7) regulatory network. TCP20 positively regulates genes encoding proteins involved in nitrate assimilation and signaling and downregulates the expression of *CYCB1;1,* which encodes a key cell-cycle protein involved in the G2/M transition (Guan, 2017). Our data suggests that in addition to TCP20 transcriptional regulation, reversible protein phosphorylation may also play a role in this regulatory intersection between nitrate signaling and the cell cycle.

**Table 2:**
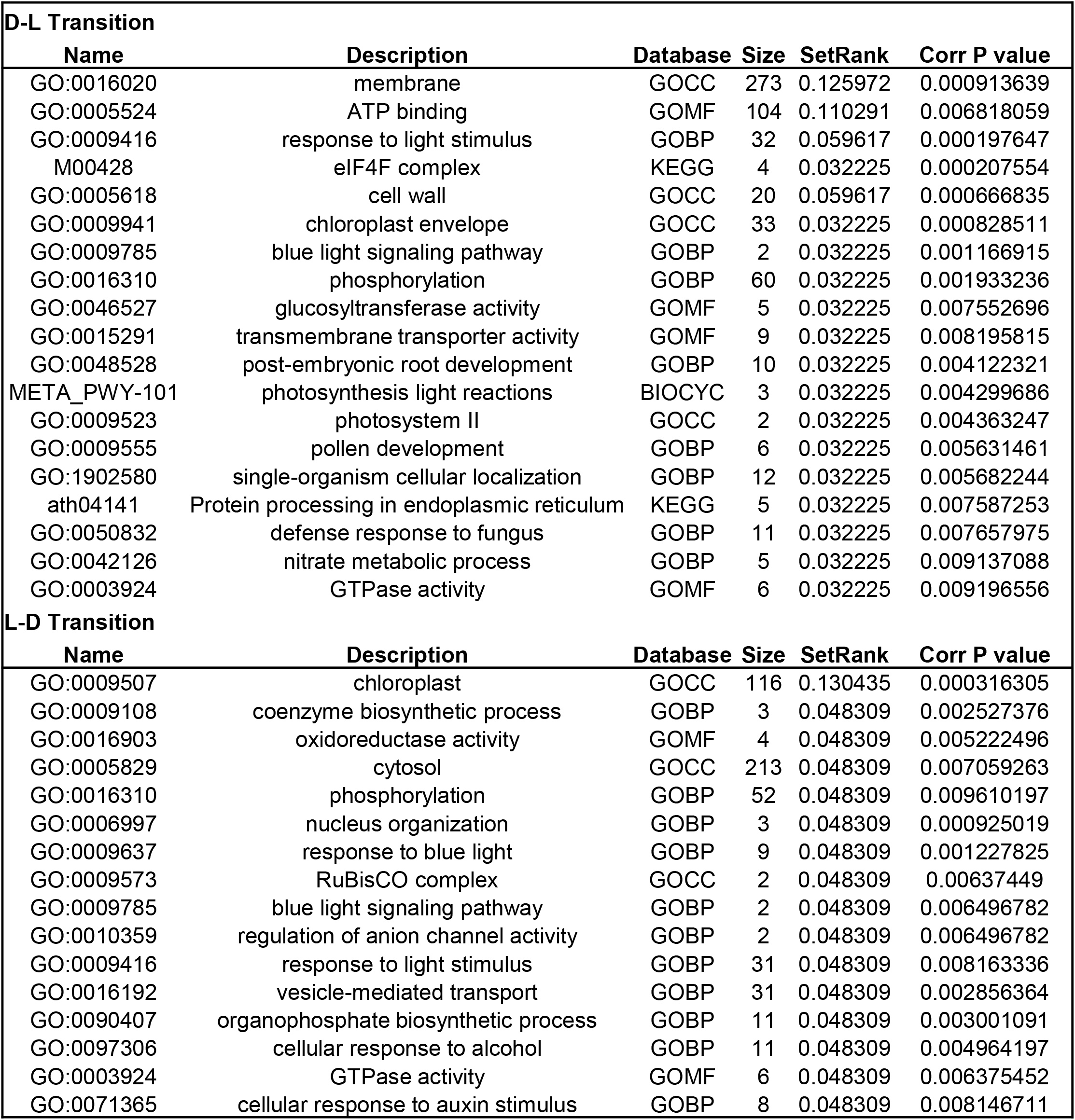
GSEA of significantly changing phosphoproteins at the D-L and L-D transition. GSEA was performed using SetRank (P value ≤ 0.01; FDR ≤ 0.05, minProt = 2).

Similar to our analysis of protein abundance changes, we built association networks using STRING-DB to complement the GSEA analysis of the phosphoproteome (Figure 4). Association networks were generated based on phosphopeptide quantification data and *in silico* subcellular localization information to examine relationships between the significantly changing phosphoproteins at both the D-L and L-D transitions. Most of the node clusters overlap between both the D-L (Figure 4A) and L-D (Figure 4B) networks, with larger clusters consisting of proteins involved in light detection and signaling, carbon and nitrogen metabolism, protein translation, hormone signaling, ion transport, cell wall related processes and protein phosphorylation. L-D transition-specific node clusters include RNA processing, transcription and secretion, and protein transport (Figure 4B). Similar to our proteome analyses, network association and GSEA analyses showed a high degree of overlap, indicating that the two approaches revealed the same cell processes in which proteins show differences in phosphorylation.

**Figure 4:**
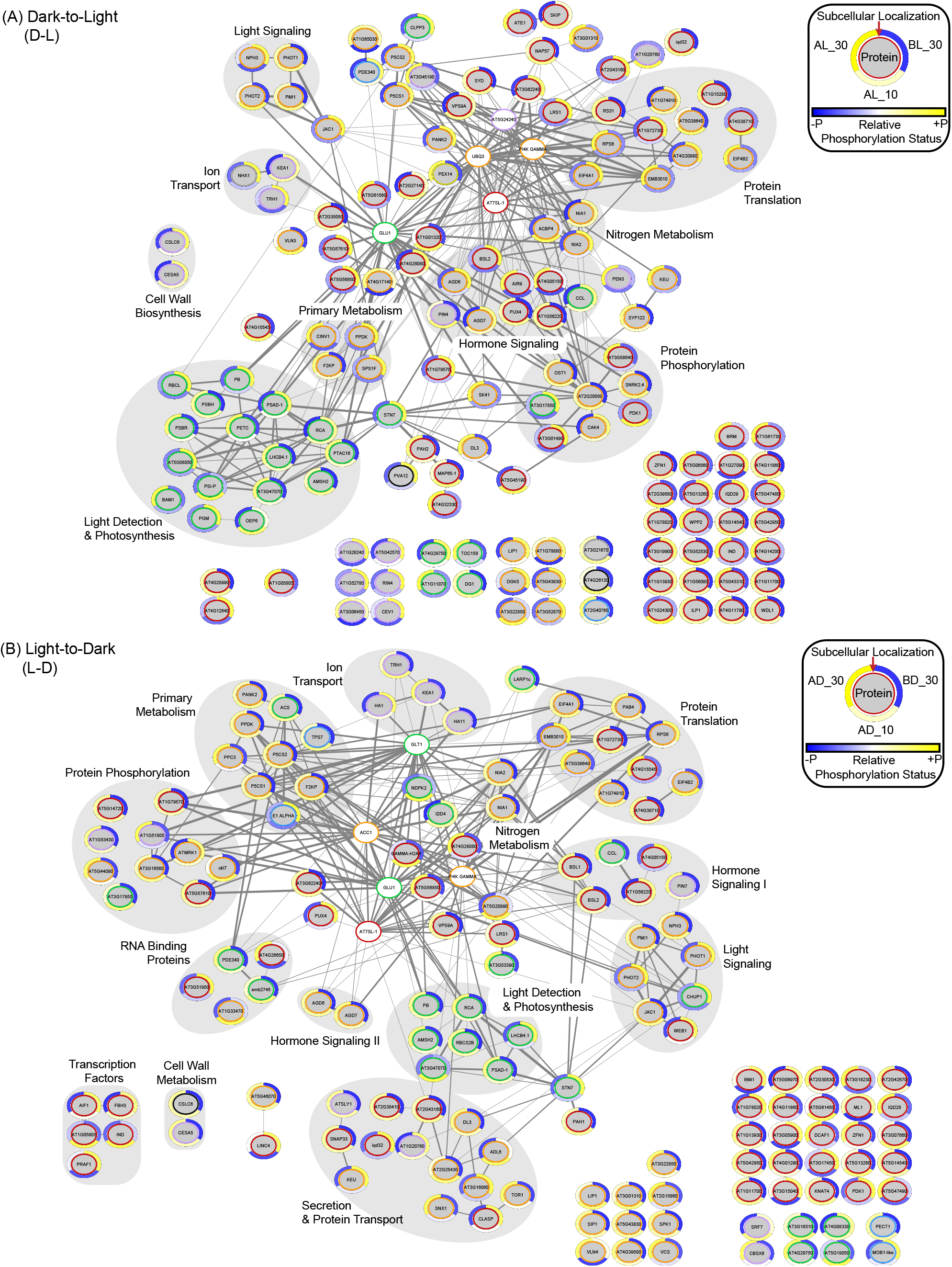
Interaction networks of the diurnal phosphoproteome at the D-L and L-D transitions. An association network analysis of statistically significant diurnally changing phosphorylated proteins was performed using the STRING-DB (ANOVA P value ≤ 0.05). Edge thickness indicates strength of the connection between two nodes (0.5 - 1.0). Phosphorylated proteins (grey circles) are labeled by either their primary gene annotation or Arabidopsis gene identifier (AGI). Outer circle around each node depicts the standardized relative log2 FC in phosphorylation status of this protein between time-points. The sliding scale of yellow to blue represents a relative increase and decrease in phosphorylation, respectively. The inner colored circles represent *in silico* predicted subcellular localization (SUBAcon; suba3.plantenergy.uwa.edu.au). Nucleus (red), cytosol (orange), plastid (green), mitochondria (blue), plasma membrane (purple), peroxisome (dark yellow), endoplasmic reticulum/golgi/secreted (black) are depicted. A second shell of 5 STING-DB proteins (white circles) not found in our dataset was used to highlight the interconnectedness of the network. Multiple nodes encompassed by a labelled grey circle represent proteins involved in the same cellular process.

Given this, we hypothesize that the significantly changing Arabidopsis proteome measured here over a 24 h photoperiod consists of proteins possessing key functions in each respective cellular process. As discussed above, protein abundance changes are generally not as widespread as transcriptome-level changes over a 24 h photoperiod (Baerenfaller et al., 2012; Graf et al., 2017; Seaton et al., 2018; Uhrig et al., 2019). Conversely, changes in protein phosphorylation can be dependent or independent of protein abundance fluctuations. To assess this, we compared our changing phosphoproteome to our changing proteome, and found that the majority of significantly changing diurnal phosphorylation events occur independent of protein abundance changes and therefore likely represent regulatory PTM events (Duby & Boutry, 2009; Le, Browning, & Gallie, 2000; Lillo et al., 2004; Muench, Zhang, & Dahodwala, 2012). Further research is required to elucidate the specific roles of these phosphorylation events. Based on these results, future investigations of which seemingly stable proteins / phosphoproteins and significantly changing phosphoproteins are in fact undergoing changes in their translation and turnover, but maintain their overall abundance (Li et al., 2017) are required to fully capture how the scale and dynamics of protein and PTM changes impact plant cell regulation.

### A small subset of the transition phosphoproteome has protein level changes

As the result of employing enrichment methods, one major question in phosphoproteomics is how the quantified phosphorylation changes relate to changes in protein abundance. To examine this, we performed an integrated analysis of the significantly changing proteome and phosphoproteome to determine if and how phosphorylation and protein abundance changes are related. Of the 226 proteins exhibiting a significant change in phosphorylation (Table 1), 60% (136 proteins) were quantified in our proteome data (Supplemental Table 6). These results are not unexpected because of the phosphopeptide enrichment strategy and indicate that 40% of the phosphorylated proteins in our phosphoproteome dataset are of lower abundance and not amongst the 4762 total quantified proteins. Further assessment of significantly changing phosphoproteins relative to the quantified proteome at the light transitions found that 25% (L-D) and 7.1% (D-L) of the changing phosphoproteins were not significantly changing at the protein level (TOST P value ≤ 0.05, ε = 0.4).

We then directly compared the significantly changing phosphoproteome and proteome to identify proteins exhibiting a change in both diurnal protein abundance and phosphorylation status. We found that a total of six phosphoproteins (totaling 2.1% of all 288 proteins significantly changing in protein abundance; Supplemental Table 6) that fit this criteria (Figure 5). These include nitrate reductase 1 (NIA1; AT1G77760) and 2 (NIA2; AT1G37130), protein kinase SnRK2.4 (AT1G10940), Rho guanyl-nucleotide exchange factor SPK1 (AT4G16340), microtubule binding protein WDL5 (AT4G32330), and winged-helix DNA-binding transcription factor family protein LARP1C (AT4G35890). NIA1 and 2 are directly related to nitrogen assimilation (Lillo, 2008; Lillo et al., 2004), while WDL5 has been implicated in mitigating ammonium toxicity through ETHYLENE INSENSITIVE 3 (EIN3) (Li et al., 2019). SnRK2.4 binds fatty acid derived lipid phosphatidic acid to associate with the plasma membrane (Julkowska et al., 2015) and responds to changes in cell osmotic status (Munnik et al., 1999), while SPK1, WDL5 and LARP1C are connected to plant hormone signaling through abscisic acid (WDL5; Yu et al., 2019), jasmonic acid (LARP1C; B. Zhang, Jia, Yang, Yan, & Han, 2012) and auxin (SPK1; Lin et al., 2012; Nakamura et al., 2018). Of these three proteins with concerted phosphorylation and abundance changes only SPK1 showed a parallel increase in abundance and phosphorylation at the same transition (Figure 5), while WDL5 and LARP1C exhibited opposing patterns of phosphorylation and abundance changes, suggesting that phosphorylation may impact their turnover. Previously, proteins involved in phytohormone signaling have been found to be regulated by both protein phosphorylation and turnover (Dai et al., 2013; Qin et al., 2014), suggesting that these three proteins may represent new examples of hormone-mediated phosphodegrons or phospho-inhibited degrons (Vu, Gevaert, & De Smet, 2018). Further examination of the ubiquitination status of these proteins and the proximity of those ubiquitin modifications to the annotated phosphorylation event are required to fully elucidate this hypothesis.

**Figure 5:**
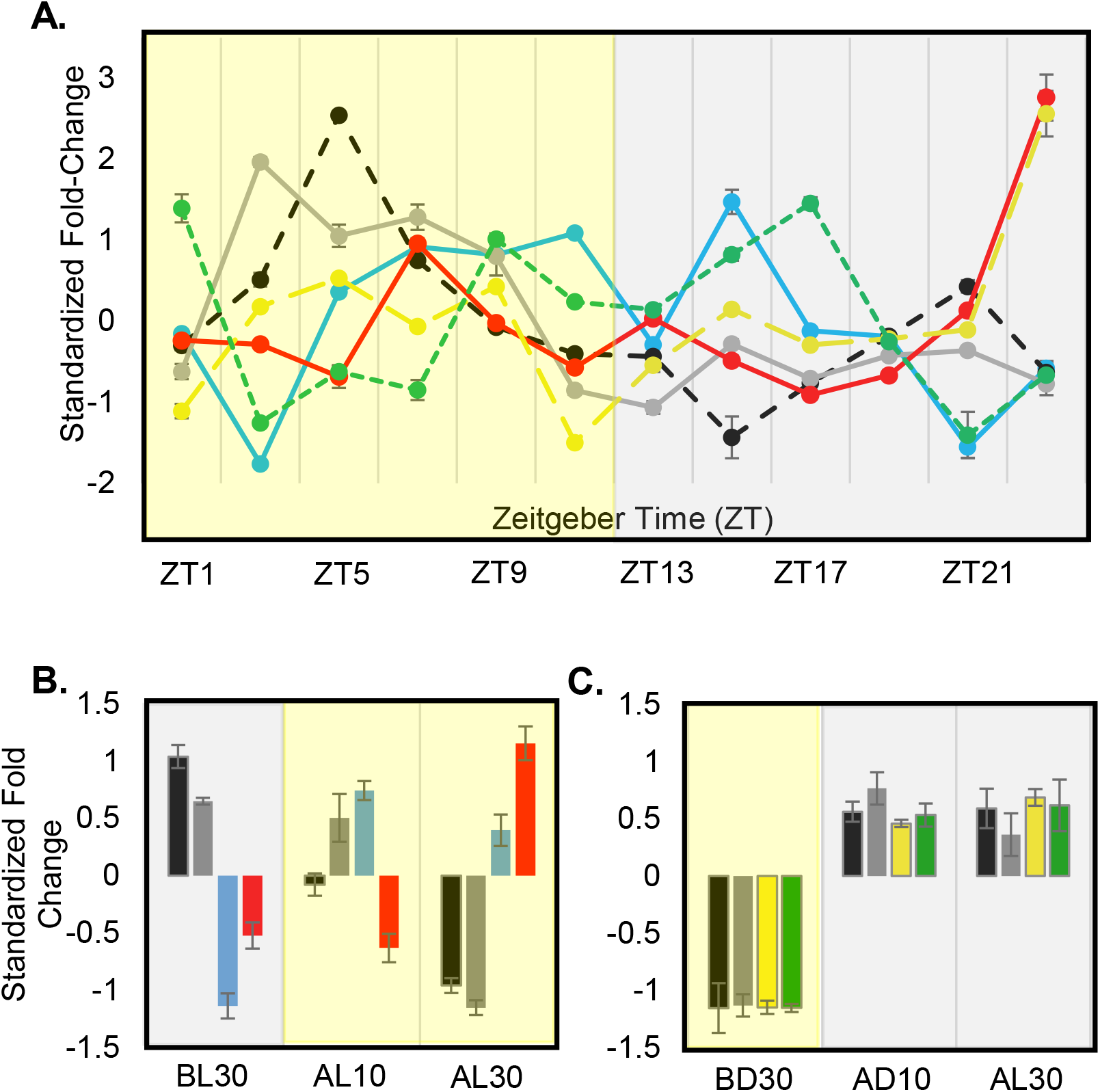
Proteins exhibiting a significant change in both diurnal protein abundance and protein phosphorylation status. Six proteins were found to significantly change in protein abundance and protein phosphorylation: AT1G10940 (SnRK2.4; blue), AT1G37130 (NIA2; black), AT1G77760 (NIA1; grey), AT4G32330 (TPX2; red), AT4G16340 (SPK1; yellow), AT4G35890 (LARP1c; green). (A) Diurnal protein abundance change profile. Standardized fold-change values are plotted relative to ZT. (B) D-L and (C) L-D phosphorylation change profiles. Standardized fold-change values are plotted relative to transition time-point either 10 or 30 minutes before light (BL), after light (AL), before dark (BD) or after dark (AD). Standard error bars are shown.

### Motif analysis reveals diurnal utilization of phosphorylation sites

We next hypothesized that we could connect our phosphoproteome data to a subset of protein kinases that may catalyze these diurnal events using a combination of motif enrichment analysis, available diurnal transcriptomic data and published literature. To understand which phosphorylation motifs are enriched in our dataset and to connect these to known protein kinases, we utilized Motif-X (motif-x.med.harvard.edu; Chou & Schwartz, 2011; Schwartz & Gygi, 2005). The significantly changing phosphorylated peptides at each transition were analyzed against all a background of all quantified phosphopeptides (P value ≤ 0.05). Motifs corresponding to serine (pS) phosphorylation sites were enriched at each transition, while enrichment of phosphorylated threonine (pT) or tyrosine (pY) motifs was absent (Supplemental Table 7). The lack of pY motif enrichment has also been reported in other studies examining phosphoproteome changes under either ED vs EN (Reiland et al., 2009; Uhrig et al., 2019) or free-running circadian cycle (Choudhary et al., 2015; Krahmer et al., 2019) experimental scenerios. Only one pT motif (pTP) has been previously associated with ED vs EN phosphoproteome changes (Uhrig et al., 2019). The lack of an enriched pTP motif here is likely due to our stringent multi-time point threshold requirement for each phosphorylation site to be considered for quantification versus the two time-point comparison previously performed between ED and EN only (Uhrig et al., 2019). Furthermore, we would expect pS motifs to be enriched over either pT or pY motifs given pS events account for 84-86% of all phosphorylation events in plants, compared to only 10-12% pT and 1-4% pY (Nakagami et al., 2010; Sugiyama et al., 2008). This makes it generally less likely to find an enrichment of pT and/or pY motifs in the phosphoproteome of plants. Of the phosphorylation sites (site probability score ≥ 0.8) we quantified, 82.8%, 16.5% and 0.7% were pS, pT and pY respectively, which aligns with previously reported distributions of phosphorylation events in *Arabidopsis thaliana* (Nakagami et al., 2010; Sugiyama et al., 2008).

At the L-D transition, we found 16 motifs of which 10 correspond to phosphorylation sites and motifs previously identified as targets of protein kinases CaMKII, PAK1, extracellular signal-regulated kinase (ERK 1/2), proto-oncogene c-RAF (RAF1), and cell division cycle 2 (CDC2) protein kinase A and B (Supplemental Table 7). Six phosphorylation sites did not correspond to known kinase motifs, and therefore likely represent currently uncharacterized and possibly plant-specific motifs considering the large expansion of protein kinases in plants relative to humans (Lehti-Shiu & Shiu, 2012; Zulawski et al., 2014). At the D-L transition, four of five identified motifs are known phosphorylation sites for checkpoint kinase 1 (CHK1), PAK2, calmodulin kinase IV (CaMKIV) and casein kinase (CKII) (Supplemental Table 7). CKII is known to phosphorylate the core circadian clock transcription factors LHY and CCA1 (Lu et al., 2011), which also peak at the D-L transition (Kusakina & Dodd, 2012).

The phosphoproteome data and motif analysis indicate that CAMKs are involved mediating L-D and D-L transition phosphosignaling and thus implicate the involvement of intracellular calcium (Ca^2+^) in circadian regulation (Marti Ruiz et al., 2018). This suggests that calcium-dependent calmodulin (CaM) protein kinase orthologs are interesting candidates for mediating circadian clock signaling. Unlike the enrichment of CKII motifs at only the D-L transition, we find enrichment of Ca^2+^ related kinases CaMKII (D-L and L-D) and CaMKIV (D-L) phosphorylation motifs at each transition (Supplemental Table 7). Previous analyses have also identified Ca^2+^ kinase motifs enriched at both ED and EN (CDPK-like motifs; Uhrig et al., 2019). Additionally,SnRK1-related motifs were identified in the phosphoproteome data from Arabidopsis CCA1-Ox plants growing in a free-running cycle (Krahmer et al., 2019). SnRK1 is a central mediator of energy signaling between different organelles and also functions to phosphorylate CDPKs (Wurzinger, Nukarinen, Nagele, Weckwerth, & Teige, 2018). Together, these studies and the results presented here suggest a broader role for Ca^2+^ in diurnal plant cell regulation during the L-D and D-L transitions.

Compared to humans, plants have more protein kinases (Lehti-Shiu & Shiu, 2012; Zulawski et al., 2014), but most of their targets remain unknown. Our phosphoproteome results, together with previously reported diurnal phosphoproteome datasets (Choudhary et al., 2015; Krahmer et al., 2019; Reiland et al., 2009; Uhrig et al., 2019) provide a compilation of phosphorylation motifs that are rapidly modified at the D-L and L-D transitions. Unfortunately, most protein kinases are outside the dynamic range of protein detection in conventional systems-level quantitative proteomic studies. However, when we integrate available transcriptional data for the diurnal expression of protein kinases (Uhrig et al., 2019) with the phosphoproteome changes uncovered here and in other studies (Choudhary et al., 2015; Krahmer et al., 2019; Reiland et al., 2009; Uhrig et al., 2019), we can begin to narrow the protein kinase sub-families and specific genes to those most likely catalyzing the observed diurnal phosphorylation events.

### Key plant processes involve independent changes in both proteome and phosphoproteome

When we queried the data for proteins that change in their abundance and/or phosphorylation status over the 24 h photoperiod, we found proteins predominantly involved in translation, cell wall biosynthesis and multiple aspects of plant metabolism. We hypothesize that these cellular processes are particularly susceptible to diurnal plant cell regulation at the protein level. The translation rates of Arabidopsis enzymes of light-induced metabolic reactions fluctuate diurnally and this correlates with their activity (Seaton et al., 2018). For example, several central metabolic enzymes are synthetized at 50 to 100% higher rates during the light phase of the photoperiod (Pal et al., 2013; Piques et al., 2009). Correspondingly, we identified 15 proteins involved in protein translation that have diurnal changes in abundance (Table 3; Supplemental Table 2). Although they belong to several clusters shown in Figure 1A, nine of the proteins are grouped in CL3, which exhibits a general protein increase at the onset of light. In addition, we found eight translation-related proteins with changes in their phosphorylation status at L-D and D-L transitions, of which 5/8 are eukaryotic initiation factor (eIF) proteins (Table 3; Supplemental Table 8). Phosphorylation is known to affect eukaryotic translation at the initiation step (Jackson, Hellen, & Pestova, 2010; Le et al., 2000; Muench et al., 2012), and numerous eIFs and ribosomal proteins show differences in phosphorylation levels between light and dark periods (Boex-Fontvieille et al., 2013; Enganti, Cho, Toperzer, Urquidi-Camancho & von Arnim, 2018; Turkina, Klang Arstrand, & Vener, 2011; Uhrig et al., 2019). Our analysis revealed additional diurnally regulated eIFs and suggests that specific translational regulation mechanisms and ribosome composition could be controlled by light changes (e.g. day versus night) and also throughout the 24h photoperiod.

**Table 3:**
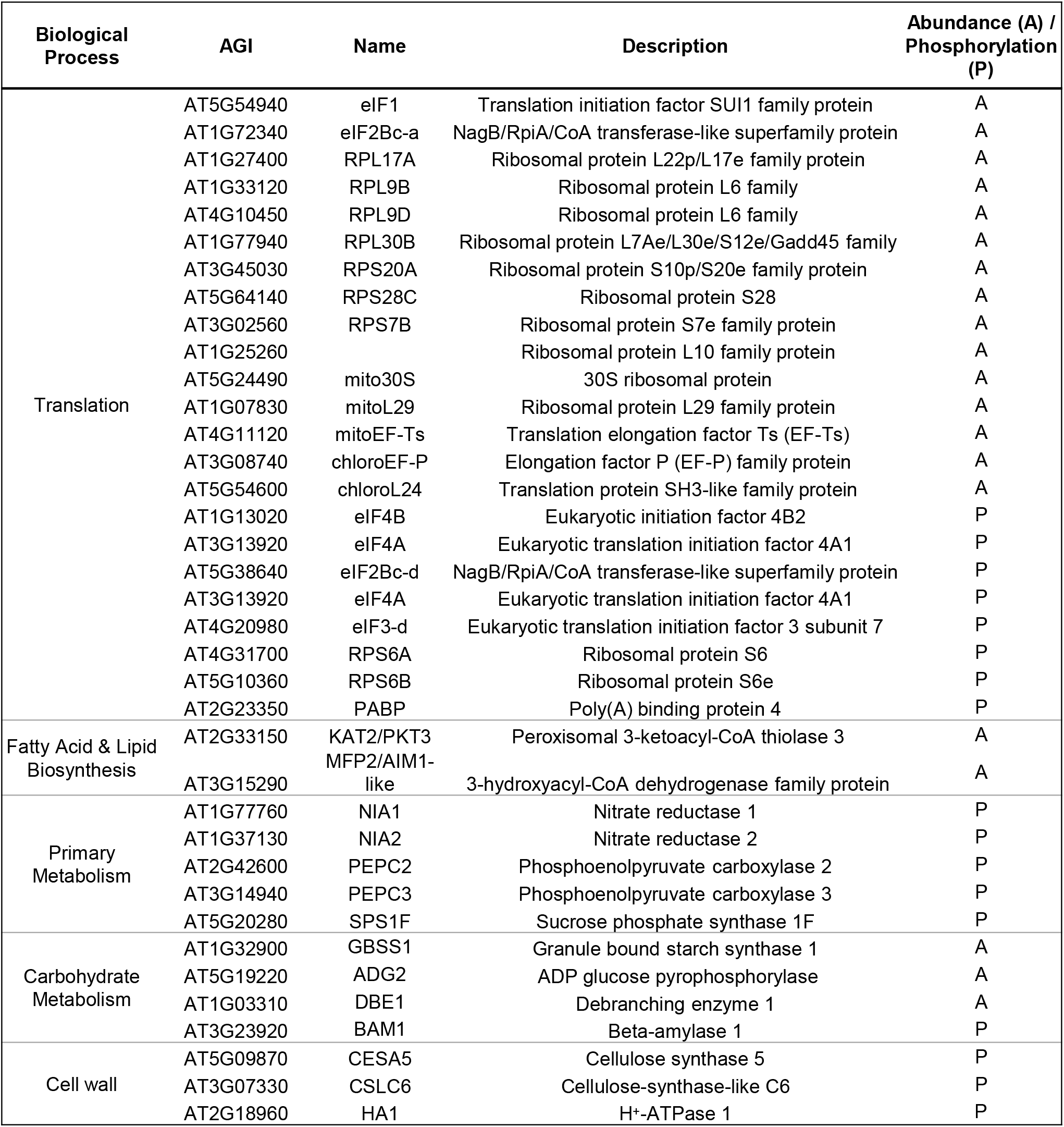
Proteins involved in plant cell processes with independent changes in abundance and/or phosphorylation.

We also find cell wall metabolic enzymes undergoing both diurnal fluctuations in protein abundance (Figure 2, Table 3; Supplemental Table 2) and changes in phosphorylation status (Figure 3, Table 3; Supplemental Table 4) at the D-L and L-D transitions. Cell wall biosynthesis is a major metabolic activity of growing plants (Barnes & Anderson, 2017; Cosgrove, 2005). We find that cellulose synthase enzymes CESA5 (AT5G09870) and CSLC6 (AT3G07330) were rapidly phosphorylated at the L-D transition. CESA5 has been shown to be phosphorylated and phosphorylation memetic-mutant enzymes increase movement of the cellulose synthase complex (CSC) in dark-grown seedlings, indicating a photoperiod-dependent regulation cell wall biosynthesis (Bischoff et al., 2011). Diurnal cellulose synthesis may also be controlled by the intracellular trafficking of CSC enzymes as a result of changes in metabolism (Ivakov et al., 2017). In dark-grown hypocotyls the ratio of CESA5 to CESA6 phosphorylation in the CSC complex is important for cellulose synthesis (Bischoff et al., 2011). Our phosphoproteome results now provide additional information on the rate of CESA5 phosphorylation at the onset of that dark period. We also find phosphorylation of the plasma membrane H^+^-ATPase HA1 (AT2G18960) at the L-D transition (Figure 3B). Phosphorylation activates H^+^-ATPases (Duby & Boutry, 2009; Sondergaard, Schulz, & Palmgren, 2004) and implicates HA1 as a primary candidate H^+^-ATPase in diurnal cell wall acidification to facilitate cell expansion during the night (Ivakov et al., 2017).

In addition to protein translation and cell wall related processes, we identified a number of enzymes involved in lipid, carbohydrate and nitrogen metabolism that change at their protein levels (Figure 2, Table 3; Supplemental Table 2) over the 24 h time-course or phosphorylation status at the D-L and L-D transitions (Table 3; Supplemental Table 9). Several of these enzymes have been previously identified as being phosphorylated (PhosPhat 4.0) (Heazlewood et al., 2008); however, our sampling of three closely spaced time-points provides new information about the rate of protein phosphorylation changes at each transition. Moreover, our results demonstrate that in Arabidopsis metabolic enzymes are subject to changes in either protein abundance or phosphorylation, or both, which likely is of regulatory relevance for metabolic pathway flux.

Our data reveals that several enzymes related to fatty acid, biotin, mitochondrial acetyl-CoA and chloroplast metabolism have diurnal changes in abundance (Figure 2; Table 3; Supplemental Table 2). Of particular interest are peroxisomal fatty acid β-oxidation enzymes 3-ketoacyl-CoA thiolase 2 (KAT2/PKT3; AT2G33150) and 3-hydroxyacyl-CoA dehydrogenase (MFP2/AIM1-like; AT3G15290). KAT2 is a central enzyme in peroxisomal fatty-acid degradation for the production of acetyl-CoA that is required for histone acetylation, which in turn affects DNA methylation (Wang et al., 2019), and ABA signaling (Jiang, Zhang, Wang, & Zhang, 2011), which is essential to daily regulation of stomatal conductance. MFP2/AIM1-like is an uncharacterized ortholog of MULTIFUNCTIONAL PROTEIN 2 (MFP2) and ENOYL-COA ISOMERASE (AIM1), which are involved in indole-3-acetic acid and jasmonic acid metabolism (Arent, Christensen, Pye, Norgaard, & Henriksen, 2010; Delker, Zolman, Miersch, & Wasternack, 2007). KAT2 loss-of-function mutants require sucrose to supplement plant acetyl-CoA production, suggesting that diurnal changes in fatty acid degradation through KAT2 and MFP2/AIM1-like are possibly tied to sucrose production and that products downstream of KAT2 and MFP2/AIM1-like (e.g. hormones) are essential to plant growth and development (Pinfield-Wells et al., 2005). Previously, fatty acid and lipid metabolism in leaves and seedlings has been suggested to be diurnally / circadian clock regulated (Gibon et al., 2006; Hsiao et al., 2014; Kim, Nusinow, Sorkin, Pruneda-Paz, & Wang, 2019; Nakamura, 2018; Nakamura et al., 2014). This includes diurnal changes in fatty acids and lipids (Gibon et al., 2006) in wild-type plants as well as diurnal changes in triacylglycerol (Hsiao et al., 2014) and phosphatidic acid (Kim et al., 2019) in the circadian clock double mutant *lhycca1.* Complementing these studies, our findings provide a new protein-level understanding of when fatty acid and lipid metabolism is diurnally impacted that differs from our current transcript / metabolite based knowledge, indicating that further protein-level investigations are required.

Furthermore, we also find diurnal changes in both protein abundance and protein phosphorylation for enzymes involved in carbohydrate metabolism (Table 3; Supplemental Table 2, 4). Starch biosynthesis and degradation is diurnally regulated to manage the primary carbon stores in plants (Kotting, Kossmann, Zeeman, & Lloyd, 2010). For example, granule bound starch synthase 1 (GBSS1; AT1G32900) levels increase preceding the D-L transition, likely in anticipation of starch granule formation (Szydlowski et al., 2011). Debranching enzyme 1 (DBE1, AT1G03310) increases in abundance at the end of the light period to facilitate effective starch degradation in the dark (Delatte, Trevisan, Parker, & Zeeman, 2005). Other enzymes such as beta-amylase 1 (BAM1; AT3G23920) were phosphorylated immediately after the onset of light. Although the function of BAM1 phosphorylation is currently unknown, our results provide information to understand its regulation in stomatal starch degradation and sensitivity to osmotic changes in rosettes (Zanella et al., 2016).

Lastly, we identified enzymes in nitrogen metabolism that changed their phosphorylation status at the D-L and L-D transitions (Table 3; Supplemental Figure 4; Supplemental Table 9). This predominantly involved NITRATE REDUCTASE 1 (NIA1; AT1G77760) and 2 (NIA2; AT1G37130) proteins. NIA1 and NIA2 are regulated both transcriptionally and post-translationally by phosphorylation (Lillo, 2008; Lillo et al., 2004, Wang, Du, & Song, 2011). Our results further the understanding of NIA regulation by newly defining a rate of change in the phosphorylation of these related isozymes at the L-D and D-L transitions, while also defining when peak NIA1 and NIA2 protein levels precisely occur relative to peak transcript levels (Table 3; Supplemental Figure 4; Supplemental Table 9). *NIA1* and *NIA2* maintain tissue-specific gene expression profiles, with *NIA1* expression generally complementing that of *NIA2* in the same organ. NIA1 was predominantly found in leaves, while NIA2 was predominantly found in meristematic tissue (Olas & Wahl, 2019). We analyzed whole Arabidopsis rosettes before bolting, of which developing leaves and apical meristematic tissue comprises only a small amount of total tissue. We therefore propose that the observed difference in NIA1 and NIA2 phosphorylation rates at these known regulatory phosphorylation sites reflect a higher sensitivity of NIA2 to changes in nitrate levels in meristematic and developing tissues (Olas et al., 2019).

Overall, what our analysis of the phosphoproteome at three D-L and L-D time-points shows is the dynamics of phosphorylation events at both transitions. Plant genomes often encode multiple forms of enzymes (isozymes) in metabolic pathways, therefore knowing the temporal rate at which related co-expressed protein orthologs are modified by PTMs such as protein phosphorylation provides more detailed information about their cellular regulation. This information is particularly useful when deciding which protein isoform may be best for engineering increased pathway flux if two are present simultaneously. NIA1 and NIA2 represent good examples of how resolving differences in PTM rates helps us better understand the role of PTMs play in temporal protein regulation. Lastly, we hypothesize that the rate at which different phosphorylation events on a protein are temporally fluctuating can be combined with enzyme kinetics to better define how metabolic flux through multiple enzyme reactions are fine-tuned by PTMs versus changes in protein abundance. Collectively, insights such as these will not only help us better understand precise regulatory differences between related orthologs (e.g. NIA1 and NIA2), but will also be broadly applicable to the other enzymes found in our dataset for future research and more targeted experimentation with these enzymes.

## CONCLUSION

To date, detailed analyses of plant functions during a 24 h diurnal cycle have predominantly focused on genome-wide changes in gene expression. Transcript-level changes can serve as a proxy for protein-level changes, but in plants transcript levels often do not correlate with protein abundance. While proteomes have a narrower dynamic range than transcriptomes, they nevertheless complement transcriptome studies because they provide direct insights into protein-level changes. Our quantitative combined analysis of the proteome over a 12 h light : 12 h dark 24 h photoperiod and the phosphoproteome at the L-D and D-L transitions during the diurnal cycle in a single experimental workflow has generated new information on diurnal abundance fluctuations and/or phosphorylation changes for Arabidopsis proteins involved in different cellular and biological processes (Figure 6). The identified proteins and phosphoproteins provide a useful basis for further experimental studies. In particular, understanding the specific functions of diurnally fluctuating ribosomal proteins involved in translation considering that hundreds of ribosomal protein isoforms are encoded by plant genomes with little information available to decipher their combinatorial assembly. Furthermore, the regulation of protein translation in plants at the protein complex level remains poorly understood, but specific time-of-day abundance peaks for these proteins suggests that temporal differences in the ribosome complex exists which likely correlated with the specific time-of-day requirements of the plant cell. Further elucidation of ribosome and protein translation regulation will be instrumental in filling the current knowledge gap between the transcriptome and proteome. Lastly, our phosphoproteome analysis during the transitions from D-L and L-D provides new information about candidate protein kinases catalyzing diurnal phosphorylation events at each transition, providing new opportunities for future systems-level and targeted studies.

**Figure 6:**
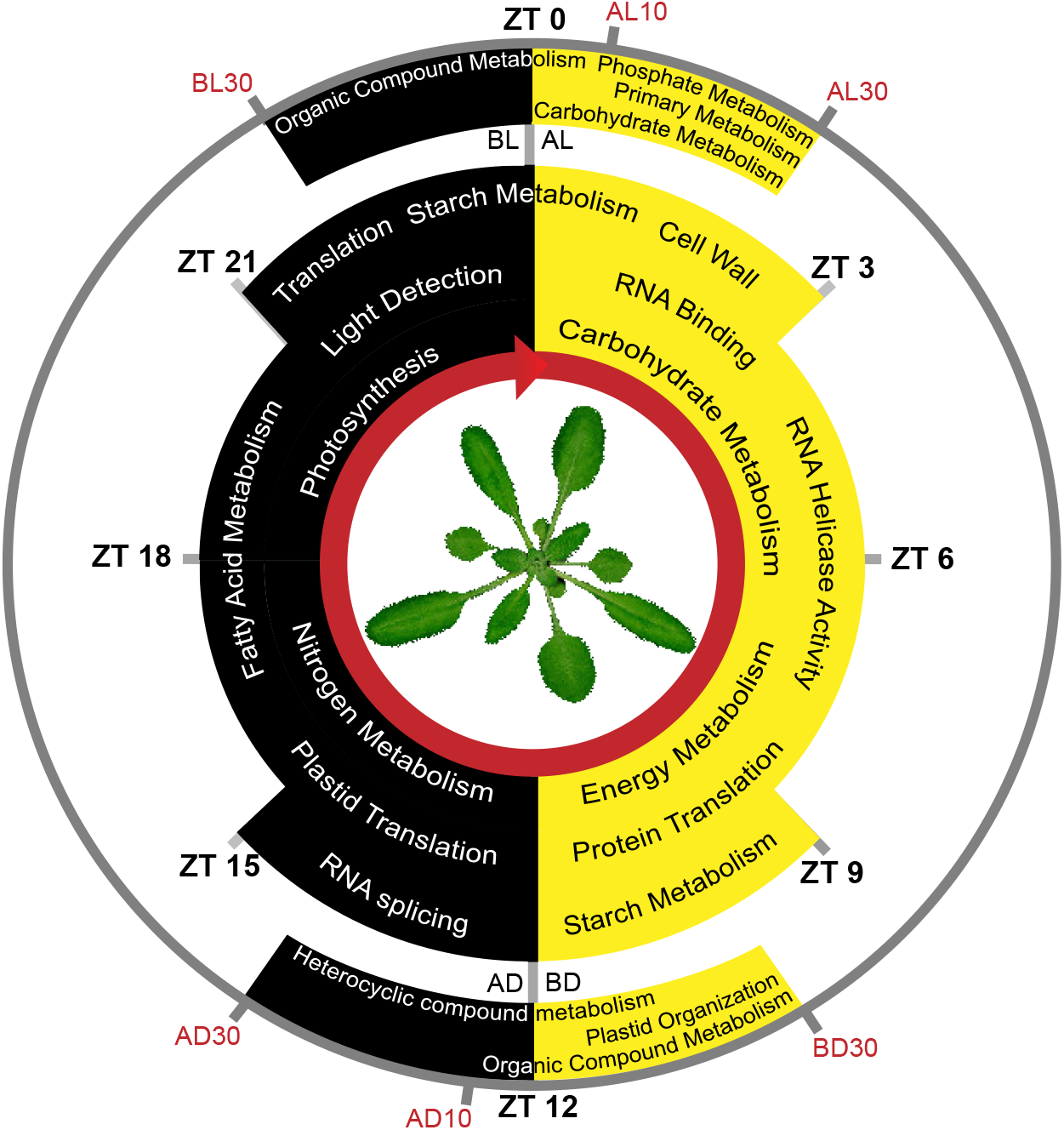
Schematic representation of Arabidopsis cellular and biological processes with diurnal fluctuations in protein abundance or protein phosphorylation. The inner three circles show terms of processes involving proteins with a maximal change in abundance during the day (yellow) or night (black). The outer circle show terms of processes involving proteins with changes in protein phosphorylation at the dark-to-light (D-L) transition (top) or light-to-dark (L-D) transition (bottom). The segments of each inner circle relative to ZT0 (day) or ZT12 (night) represent the approximate time interval in which proteins (ZT) and phosphoproteins (30 min before light or dark, 10 and 30 min after light or dark) involved in each process have their maximal change. The cellular and biological terms shown here were obtained by GO term enrichment of each protein and phosphoprotein cluster as outlined in Materials and Methods.

## Supporting information

SuppFig1

SuppFig2

SuppFig3

SuppFig4

SuppTable1

SuppTable2

SuppTable3

SuppTable4

SuppTable5

SuppTable6

SuppTable7

SuppTable8

SuppTable9

SuppData1

SuppData2

SuppData3

SuppData4

SuppData5

SuppData6

SuppData7

## ACKNOWLEDGEMENTS

This work was funded by TiMet - Linking the Clock to Metabolism (Grant Agreement 245143) supported by the European Commission (FP7-KBBE-2009-3). Experiments conducted at Forschungszentrum Jülich were partly funded by the Helmholtz Association.

## SUPPORTING INFORMATION

### Supplemental Figures

**Supplemental Figure 1: Schematic depiction of the experimental workflow.**

The total proteome and phosphoproteome experimental workflow is shown in black and blue, respectively. Light and dark boxes represent the 12 h light : 12 h dark photoperiod. The numbers on top of the boxes represent the tissue harvest times for the total proteome analysis (Zeitgeber time; ZT). The numbers below the boxes represent the tissue harvest times for the phosphoproteome analysis (minutes before or after a transition from L-D and D-L).

**Supplemental Figure 2: Hierarchical heatmap of significantly changing diurnal phosphopeptides at the D-L transition.**

The hierarchical heatmap was generated using the R package Pheatmap and Euclidean distance. Standardized relative log2 FC in phosphopeptide abundance is shown along with the corresponding AGI and phosphopeptide with phosphorylation site probabilities. GO terms of proteins in the heatmap clusters are shown on the right together with their predicted subcellular localization (SUBAcon). The segments of the circles represent the nucleus (red), cytosol (orange), plastid (green), mitochondria (blue), plasma membrane (purple) and other (black) localizations. The numbers below each pie chart represent the unique protein identifications. The time points of sampling for phosphoprotein analysis were 30 min before light (BL30), 10 min after light (AL10) and 30 min after light (AL30).

**Supplemental Figure 3: Hierarchical heat map of significantly changing diurnal phosphopeptides at the L-D transition.**

The hierarchical heat map was generated using the R package Pheatmap and Euclidean distance. Standardized relative log2 FC in phosphopeptide abundance is shown along with the corresponding AGI and phosphopeptide with phosphorylation site probabilities. GO terms of proteins in the heatmap clusters are shown on the right together with their predicted subcellular localization (SUBAcon). The segments of the circles represent the nucleus (red), cytosol (orange), plastid (green), mitochondria (blue), plasma membrane (purple) and other (black) localizations. The numbers below each pie chart represent the number of unique protein identifications. The time points of sampling for phosphoprotein analysis were 30 min before dark (BD30), 10 min after dark (AD10) and 30 min after dark (AD30).

**Supplemental Figure 4: Diurnal phosphorylation of nitrate reductase 1 (NIA1) and 2 (NIA2).** (A-B) Diurnal fluctuations of NIA1 and 2 mRNA and protein levels, and phosphorylation status. Relative changes in mRNA and protein levels were assessed over 24 h. Transcript data was extracted from Diurnal DB (http://diurnal.mocklerlab.org/). Relative changes in protein phosphorylation were measured at the D-L and L-D transitions only (see Materials and Methods). (C) Model of NIA2 protein structure including molybdenum cofactor (MoCo), dimerization (Dimer), cytochrome b5 (Cyt B), FAD and NADH binding domains in addition to hinge regions 1 and 2. Phosphorylation of the three annotated phosphorylation sites in NIA2 shown as circles is light-dependent (yellow), dark-dependent (blue) and nitric oxide-induced (white; Wang et al; 2011).

### Supplemental Tables

Supplemental Table 1: All identified and quantified proteins.

Supplemental Table 2: Significantly changing diurnal proteins.

Supplemental Table 3: All identified and quantified phosphoproteins.

Supplemental Table 4: Significantly changing diurnal phosphoproteins.

Supplemental Table 5: Benchmark phosphoproteins.

Supplemental Table 6: Comparative proteome and phosphoproteome analysis.

Supplemental Table 7: MotifX data for D-L and L-D transitions.

Supplemental Table 8: Standardized D-L and L-D changes in the phosphorylation of protein translation.

Supplemental Table 9: Standardized D-L and L-D transition phosphopeptide rates-of-change.

### Supplemental Data

Supplemental Data 1-6: The matched transcript and protein expression profiles for genes in clusters 1 – 6 respectively in Figure 1.

Supplemental Data 7: Comparison of changing diurnal proteome and a circadian proteome reported by Kramer et al. (2019).

