## Supplementary material for "Diurnal Dynamics of the Arabidopsis Rosette Proteome and Phosphoproteome": SuppFig1

Supplemental Figure 1

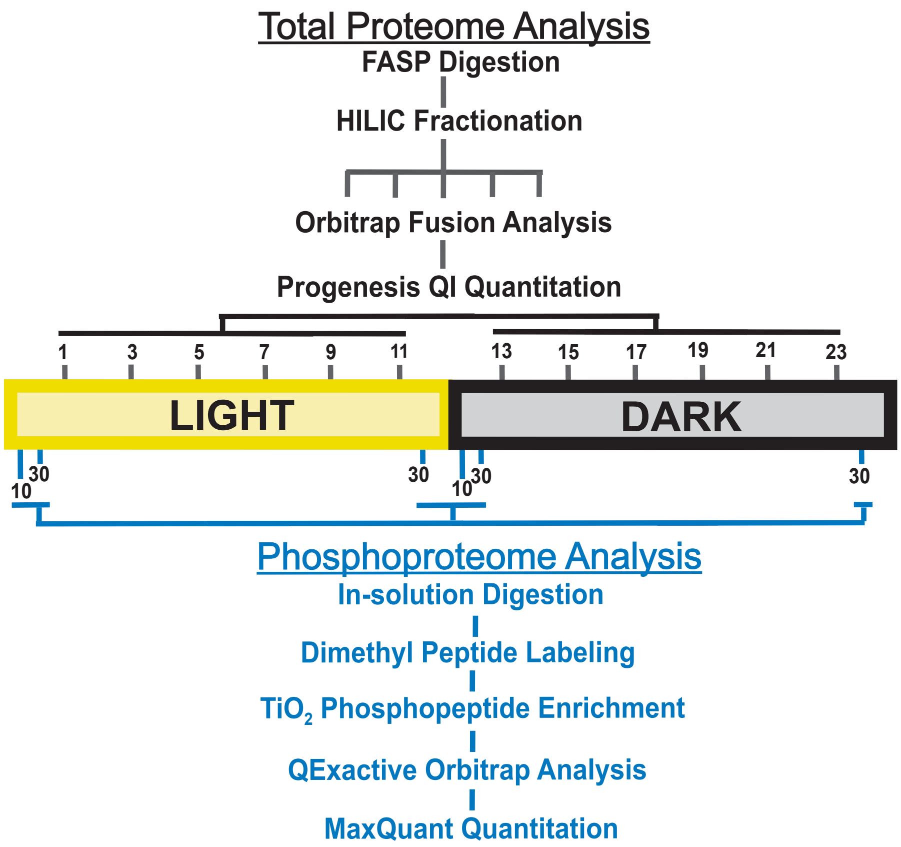


**Supplemental Figure 1: Schematic depiction of experimental workflow.** The total proteome and phosphoproteome experimental workflow is shown in black and blue, respectively. Light and dark boxes represent the 12 h light : 12 h dark photoperiod. The numbers on top of the boxes represent the tissue harvest times for the total proteome analysis (Zeitgeber time; ZT). The numbers below the boxes represent the tissue harvest times for the phosphoproteome analysis (minutes before or after a transition from L-D and D-L).
