## Supplementary material for "Diurnal Dynamics of the Arabidopsis Rosette Proteome and Phosphoproteome": SuppFig2

Supplemental Figure 2

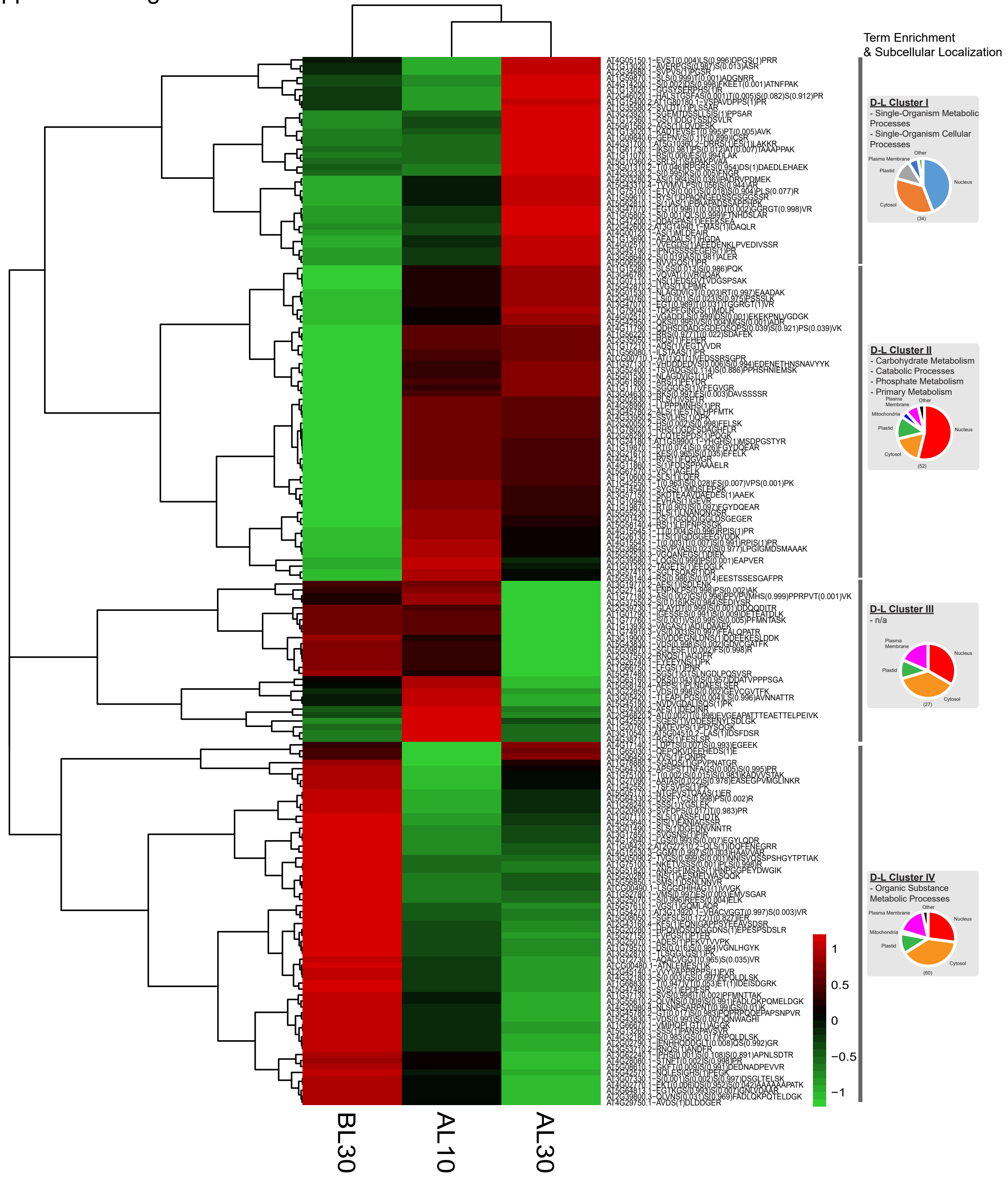


**Supplemental Figure 2: Hierarchical heatmap of significantly changing diurnal phosphopeptides at the D-L transition.** Hierarchical heatmap was generated using the R package Pheatmap using euclidean distance. Standardized relative log2 FC in phosphopeptide abundance is shown along with the corresponding AGI and phosphopeptide with phosphorylation site probabilities. GO terms of proteins in the clusters are depicted along with SUBAcon determined subcellular localization of each protein corresponding to each phosphopeptide. Nucleus (blue), cytosol (orange), plastid (grey), mitochondria (yellow), plasma membrane (purple) and other (green) localizations are shown. The numbers below each chart represents the number of unique protein identifications. BL30, AL10 and AL30 describe time-points respectively; 30 min before light (BL30), 10 min after light (AL10) and 30 min after light (AL30).
