## Supplementary material for "Diurnal Dynamics of the Arabidopsis Rosette Proteome and Phosphoproteome": SuppFig3

Supplemental Figure 3

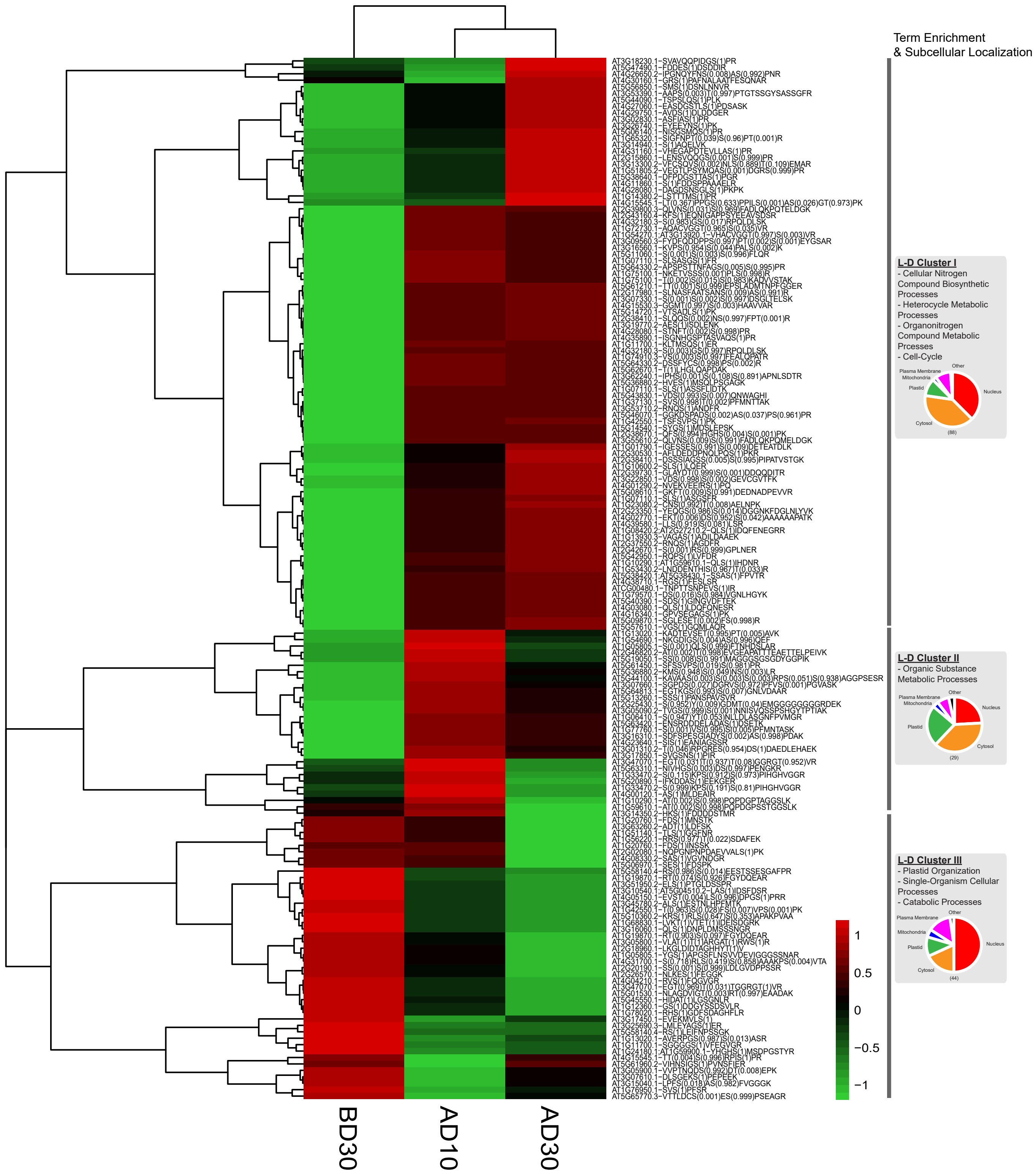


**Supplemental Figure 3: Hierarchical heat map of significantly changing diurnal phosphopeptides at the L-D transition.** The hierarchical heat map was generated using the R package Pheatmap and Euclidean distance. Standardized relative log2 FC in phosphopeptide abundance is shown along with the corresponding AGI and phosphopeptide with phosphorylation site probabilities. GO terms of proteins in the heatmap clusters are shown on the right together with their predicted subcellular localization (SUBAcon). The segments of the circles represent the nucleus (red), cytosol (orange), plastid (green), mitochondria (blue), plasma membrane (purple) and other (black) localizations. The numbers below each pie chart represent the number of unique protein identifications. The time points of sampling for phosphoprotein analysis were 30 min before dark (BD30), 10 min after dark (AD10) and 30 min after dark (AD30).
