## Supplementary material for "Diurnal Dynamics of the Arabidopsis Rosette Proteome and Phosphoproteome": SuppFig4

Supplemental Figure 4

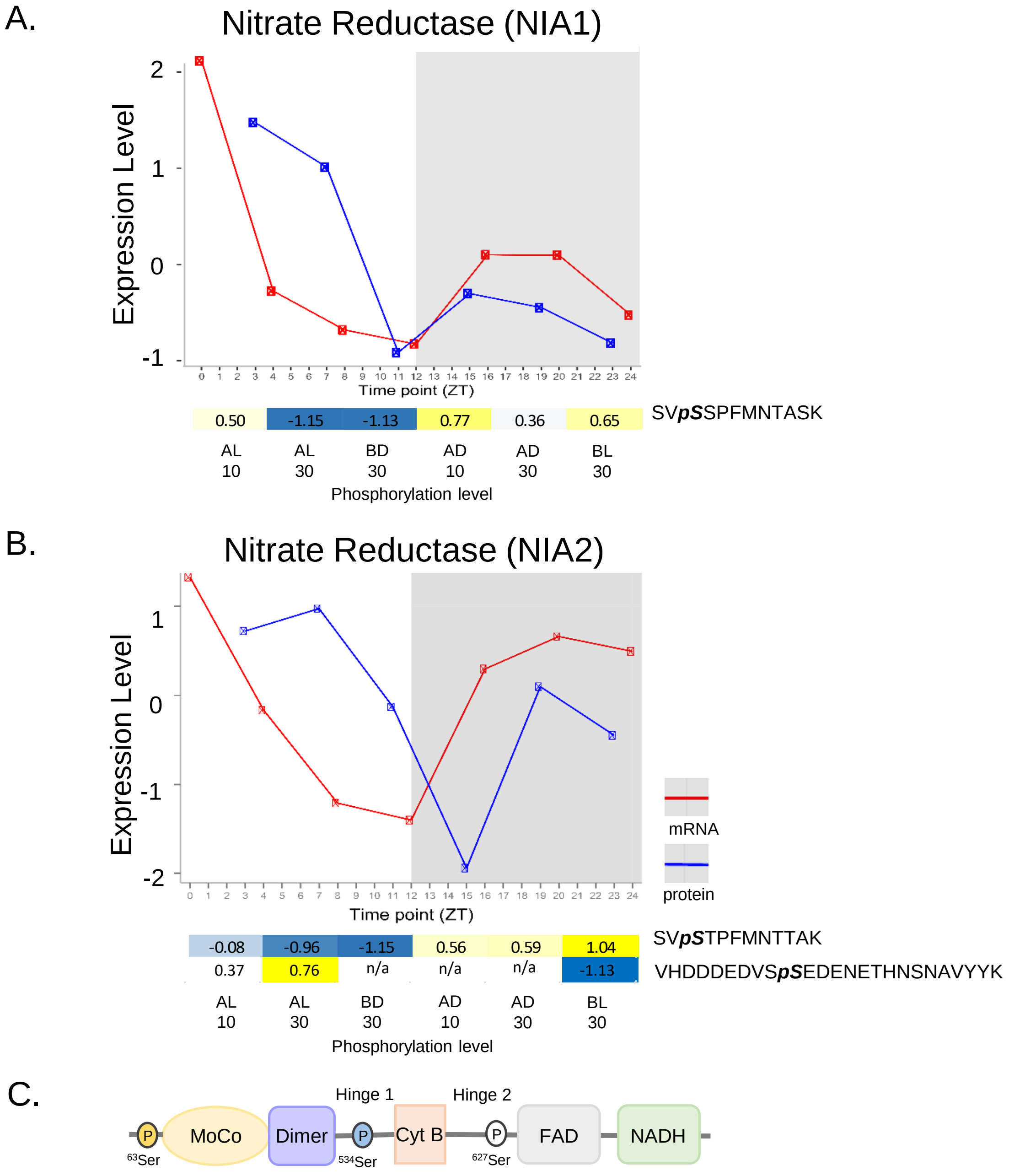


**Supplemental Figure 4 : Diurnal phosphorylation of nitrate reductase 1 (NIA1) and 2 (NIA2).** (A-B) Diurnal fluctuations in NIA1 and 2 mRNA, proteome and phosphorylation status. Relative changes in mRNA and protein levels were assessed over 24h. Transcript data was extracted from Diurnal DB (DB). Relative changes in protein phosphorylation were measure at the D-L and L-D transitions only (see materials and methods). CC represents the correlation coefficient. (C) model of NIA2 protein structure including: molybdenum cofactor (MoCo), dimerization (Dimer), cytochrome b5 (Cyt B), FAD and NADH-bind domains in addition to hinge regions 1 and 2. Three annotated phosphorylation sites of NIA2. Yellow, blue and white phosphorylation sites represent light, dark and light-independent induced phosphorylation events. The white phosphorylation event has been shown to be nitric oxide induced (Wang et al 2011) .
