## Supplementary material for "Diurnal Dynamics of the Arabidopsis Rosette Proteome and Phosphoproteome": SuppTable6

### Supplemental Table 6

**Supplemental Table 6 : Comparative Proteome and Phosphoproteome Analysis.** Quantified proteins and significantly (Sig.) changing phosphoproteins were compared to (1) assess how many phosphoproteins were also quantified at the proteome level and by extension, (2) what percentage of phosphorylation changes were due to a change in protein abundance.

| Sig. Changing Phosphoproteins (226 proteins total) | Total: Proteome Analysis | Overlap |
|---|---|---|
| vs. Total Quantified Proteins | 4762 | 60% |
| vs. Total Changing Proteins (*p-val* ≤ 0.05) | 288 | 2.1% |
| vs. Total Non-Changing Proteins L-D (*p-val* ≤ 0.05) | 1624 | 25% |
| vs. Total Non-Changing Proteins D-L (*p-val* ≤ 0.05) | 514 | 7.1% |
