## Supplementary material for "Diurnal Dynamics of the Arabidopsis Rosette Proteome and Phosphoproteome": SuppData1

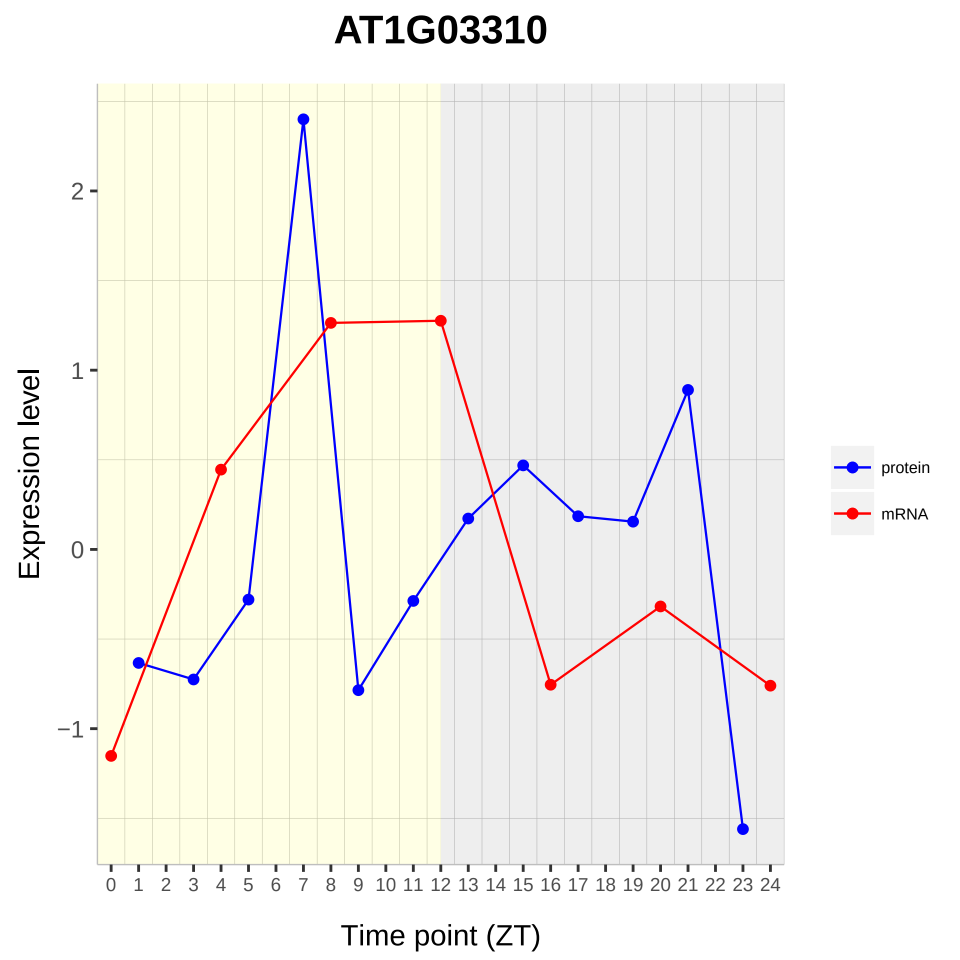

AT1G03310
Expression level
Time point (ZT)
2
1
0
−1
0 1 2 3 4 5 6 7 8 9 10 11 12 13 14 15 16 17 18 19 20 21 22 23 24
protein
mRNA

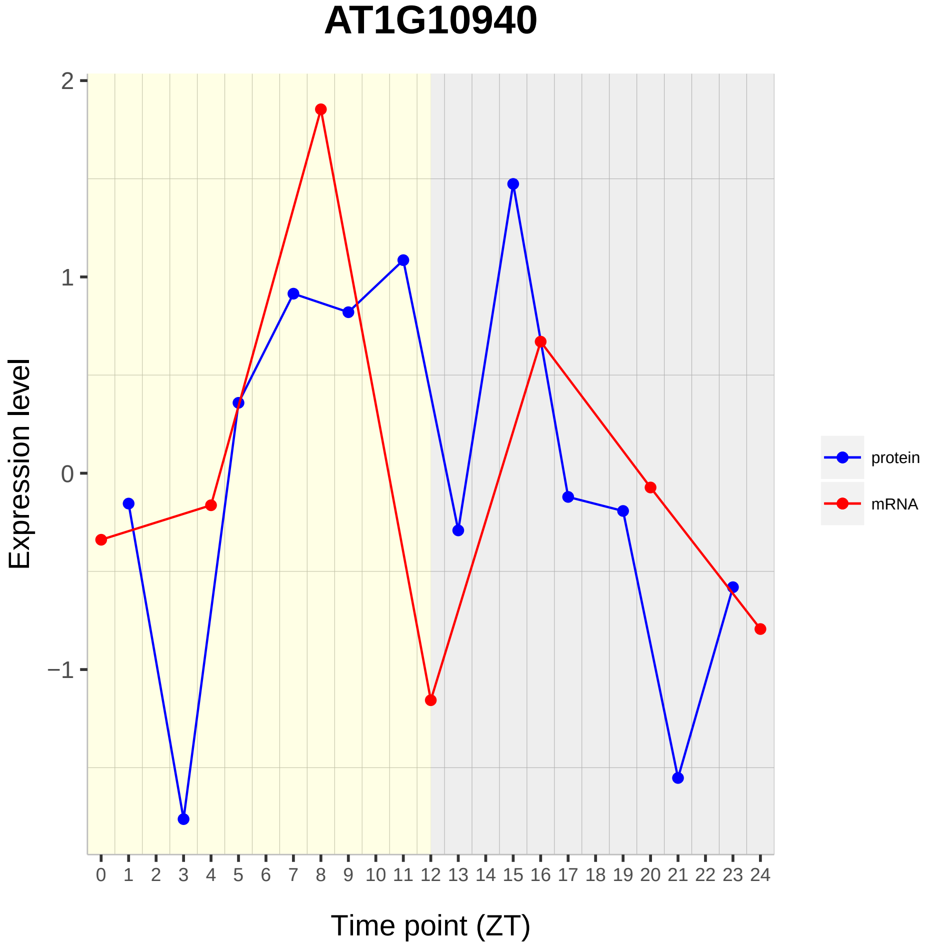

AT1G10940
Expression level
Time point (ZT)
2
1
0
−1
protein
mRNA

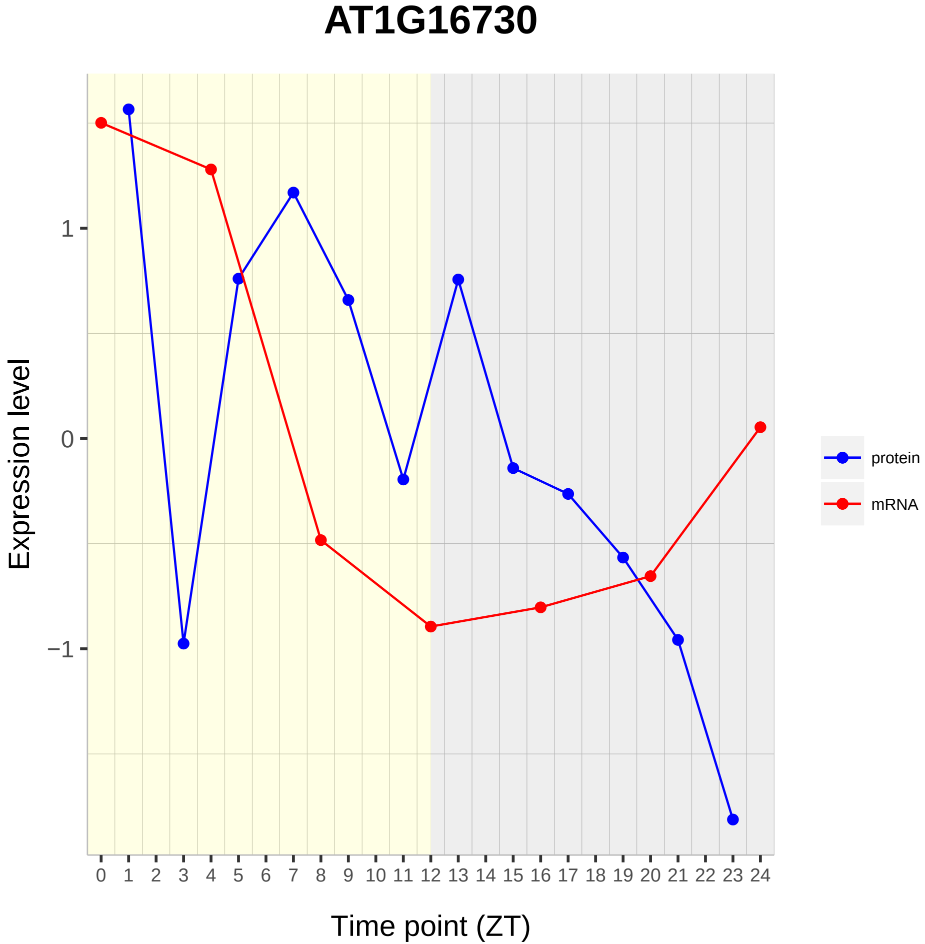

AT1G16730
Expression level
1
0
−1
0 1 2 3 4 5 6 7 8 9 10 11 12 13 14 15 16 17 18 19 20 21 22 23 24
Time point (ZT)
protein
mRNA

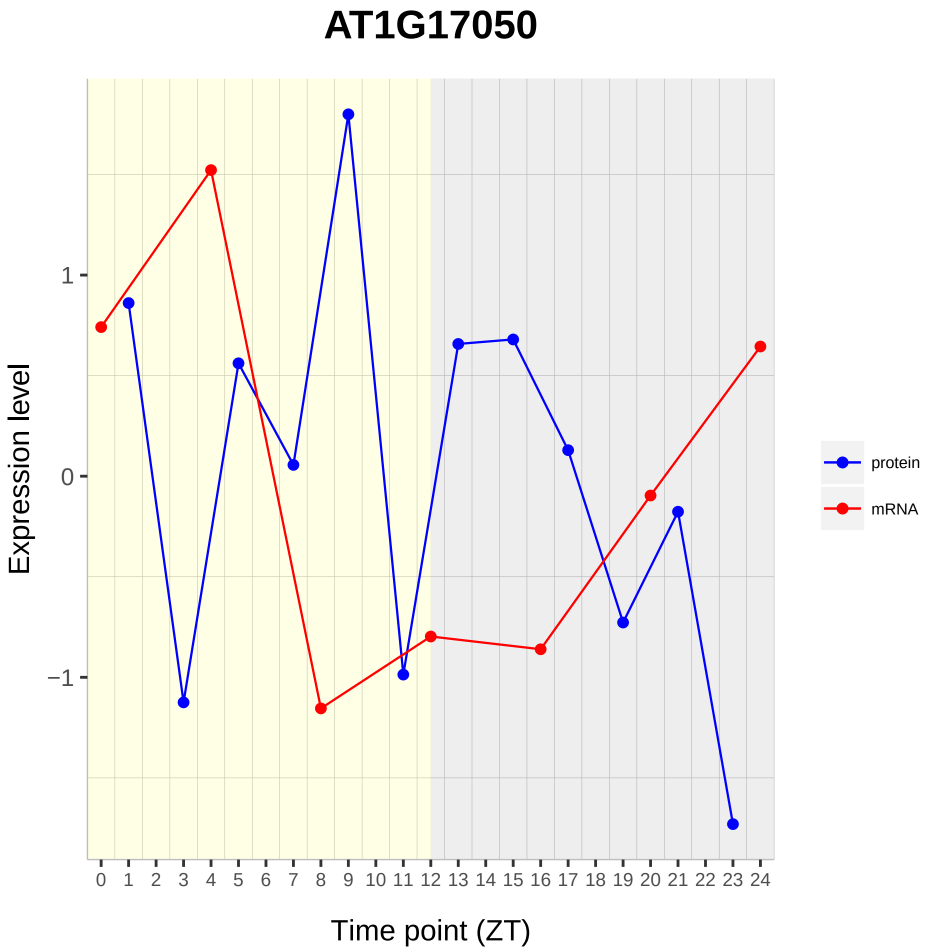
AT1G17050
Expression level
1
0
−1
Time point (ZT)
0 1 2 3 4 5 6 7 8 9 10 11 12 13 14 15 16 17 18 19 20 21 22 23 24
protein
mRNA

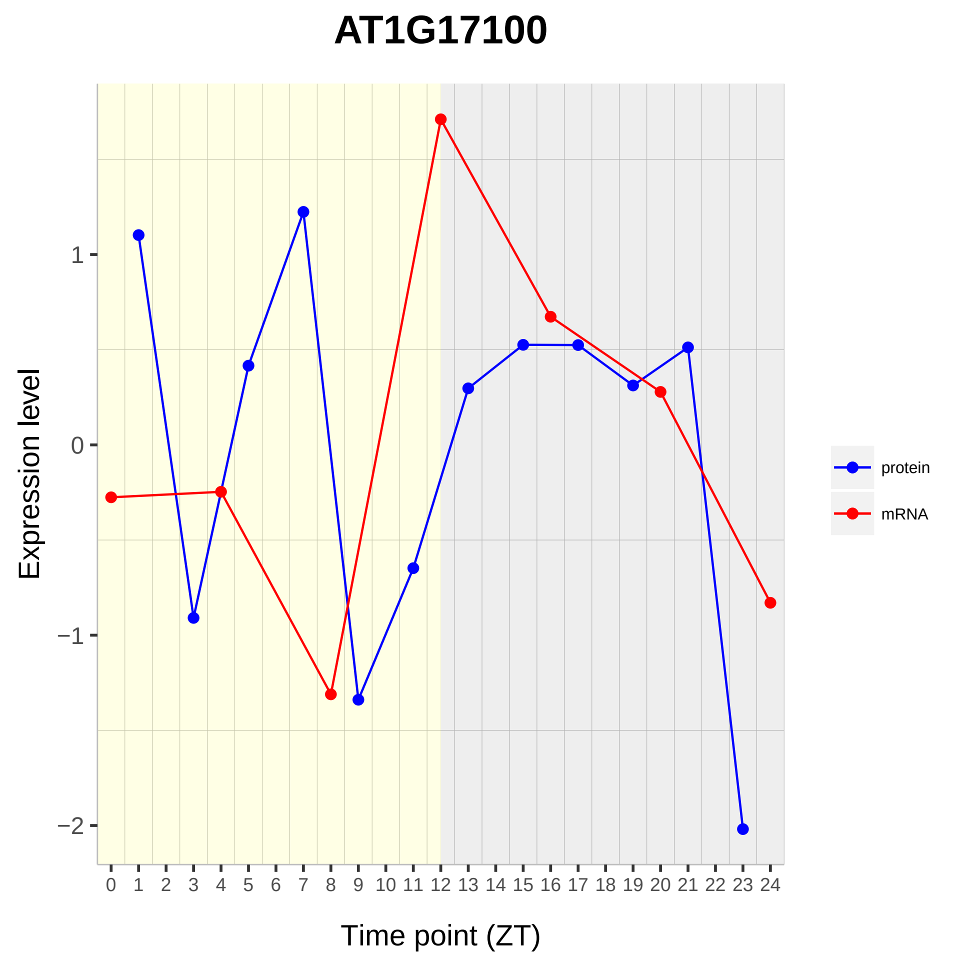

AT1G17100
Expression level
1
0
−1
−2
0 1 2 3 4 5 6 7 8 9 10 11 12 13 14 15 16 17 18 19 20 21 22 23 24
Time point (ZT)
protein
mRNA

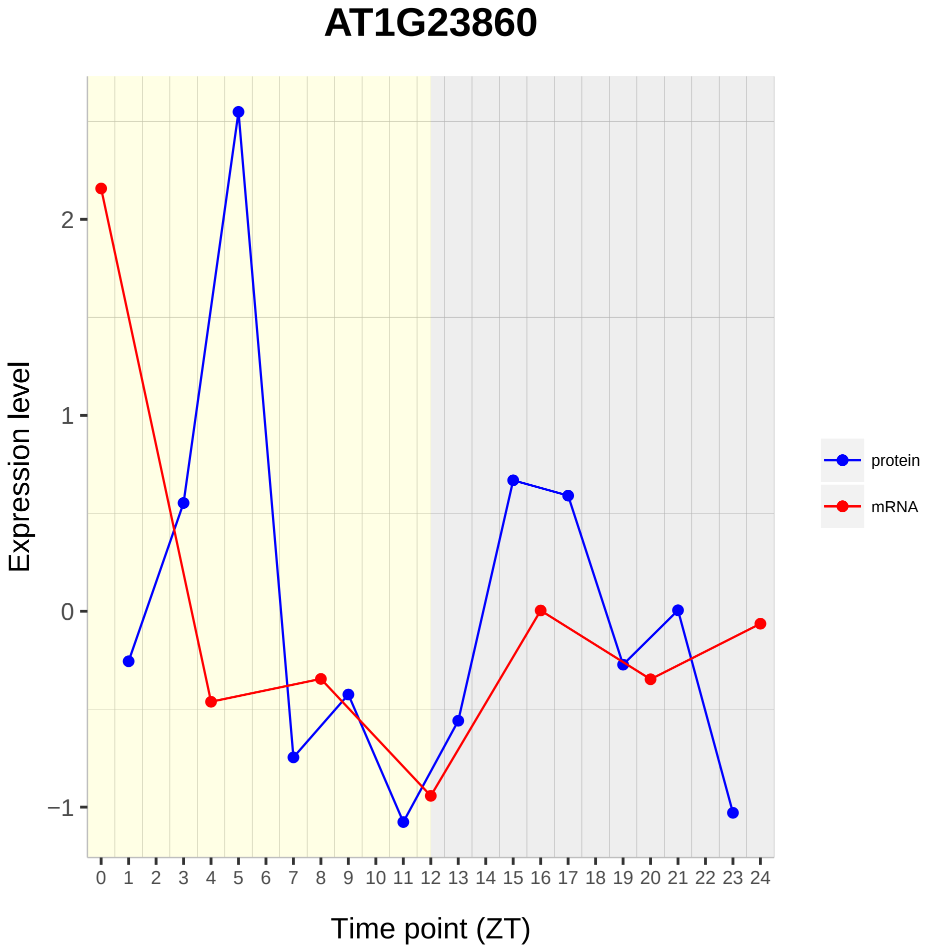

AT1G23860
Expression level
2
1
0
−1
0 1 2 3 4 5 6 7 8 9 10 11 12 13 14 15 16 17 18 19 20 21 22 23 24
Time point (ZT)
protein
mRNA

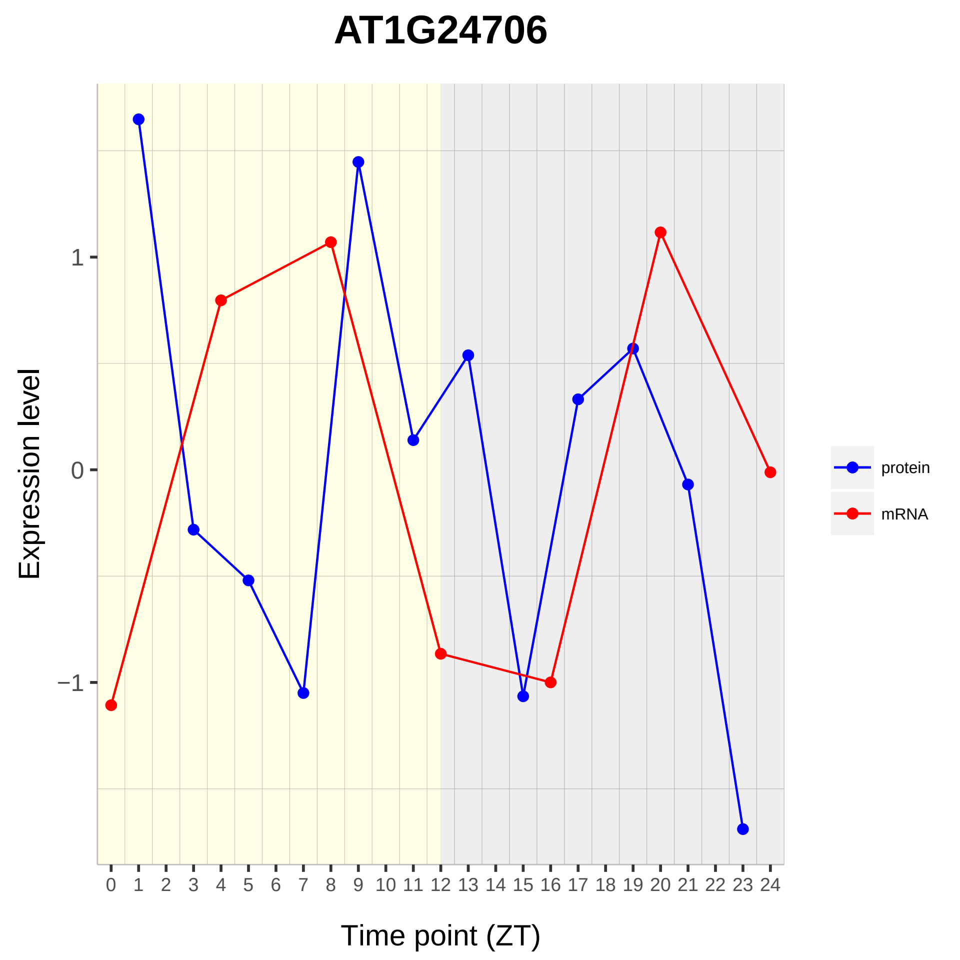

AT1G24706
Expression level
1
0
−1
0 1 2 3 4 5 6 7 8 9 10 11 12 13 14 15 16 17 18 19 20 21 22 23 24
Time point (ZT)
protein
mRNA

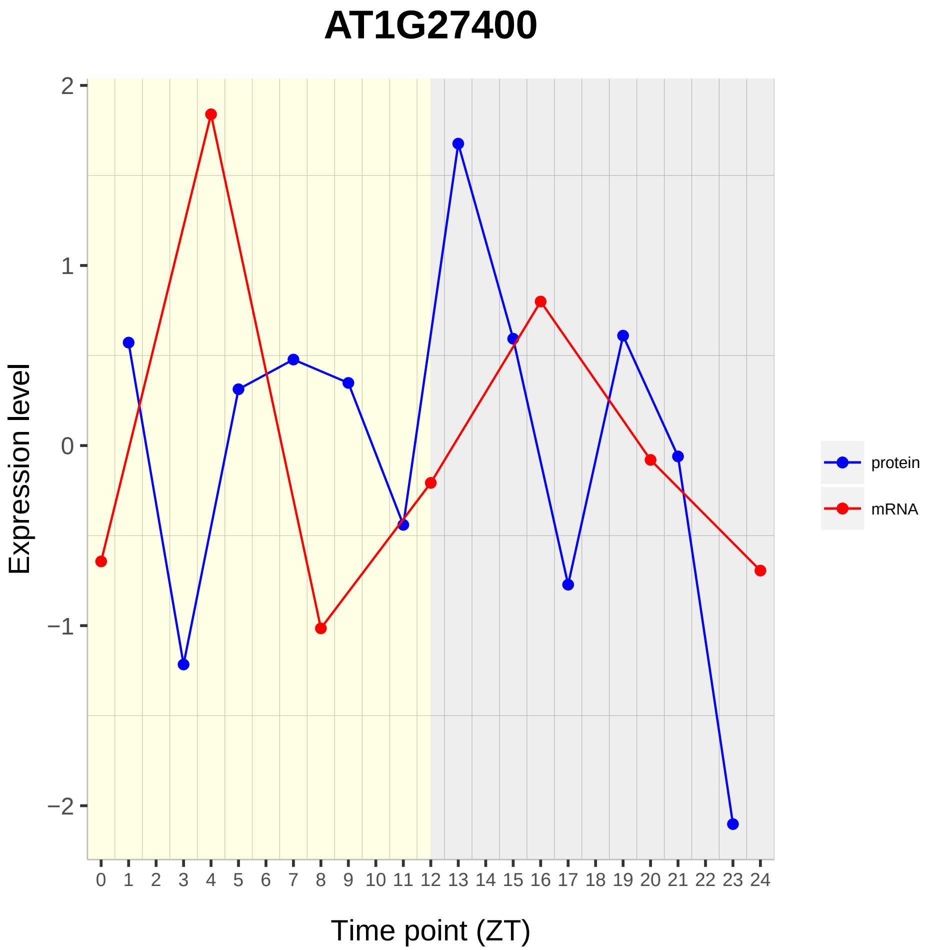
AT1G27400
Expression level
2
1
0
−1
−2
0 1 2 3 4 5 6 7 8 9 10 11 12 13 14 15 16 17 18 19 20 21 22 23 24
Time point (ZT)
protein
mRNA

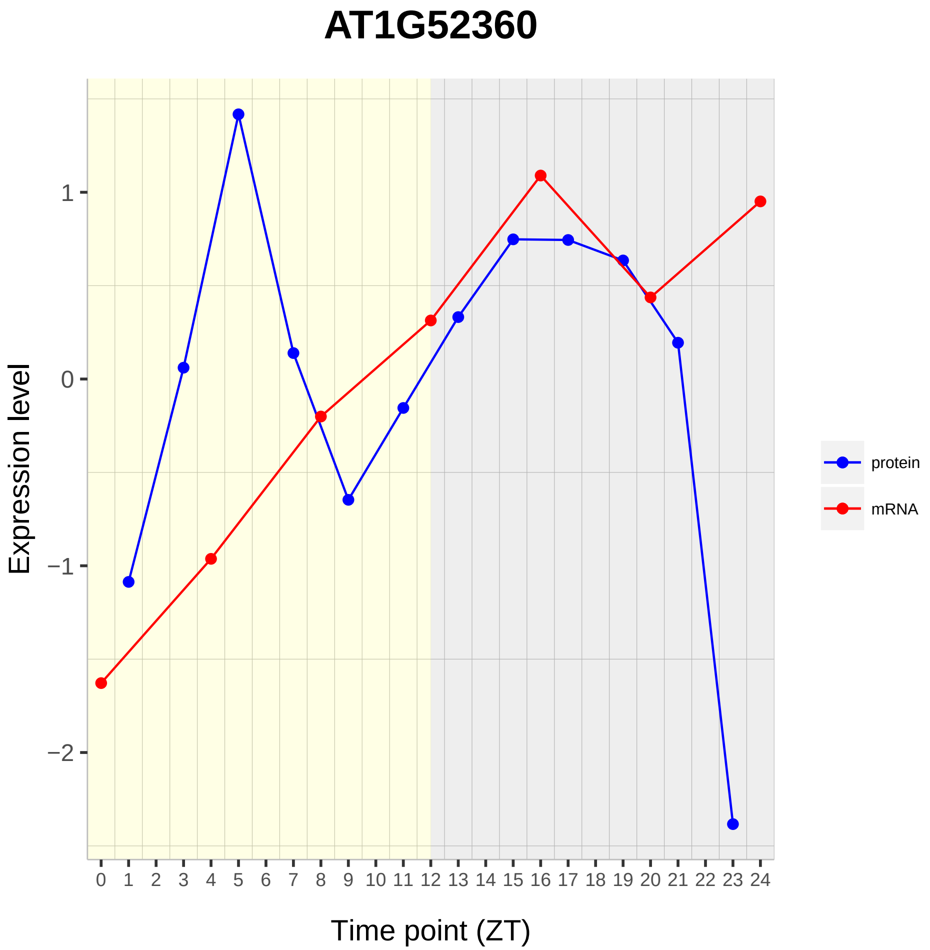
AT1G52360
Expression level
1
0
−1
−2
Time point (ZT)
protein
mRNA

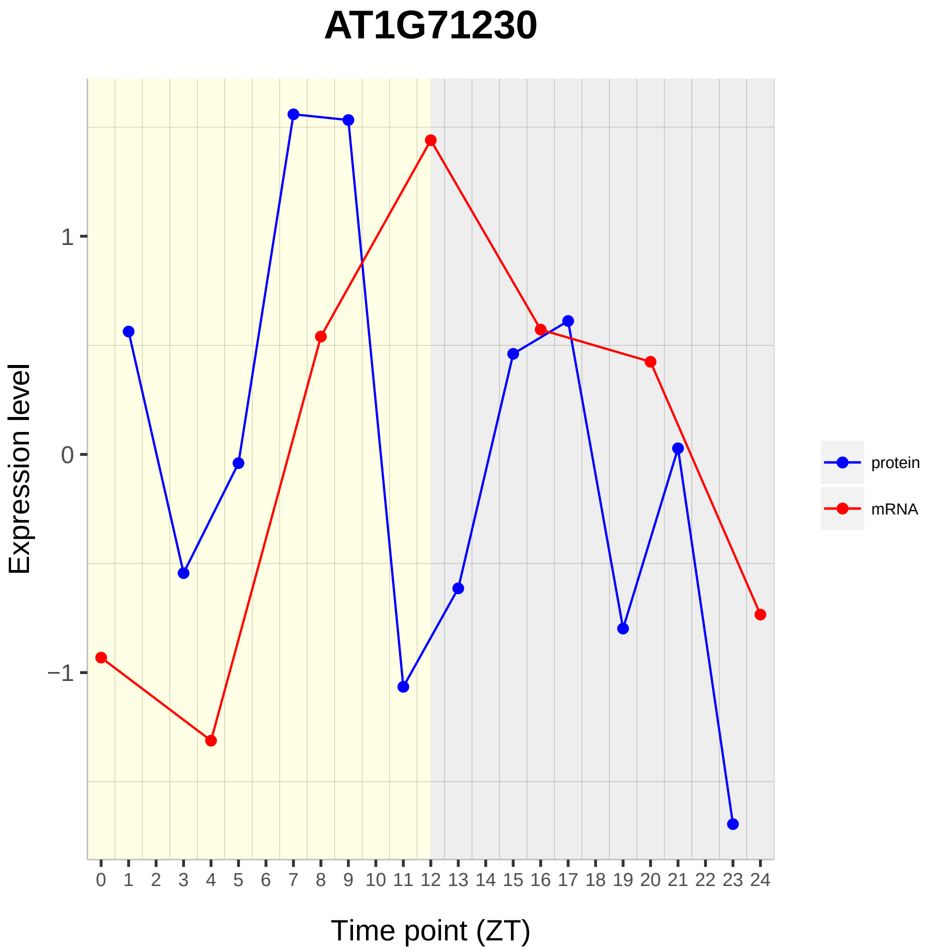
AT1G71230
Expression level
1
0
−1
Time point (ZT)
protein
mRNA

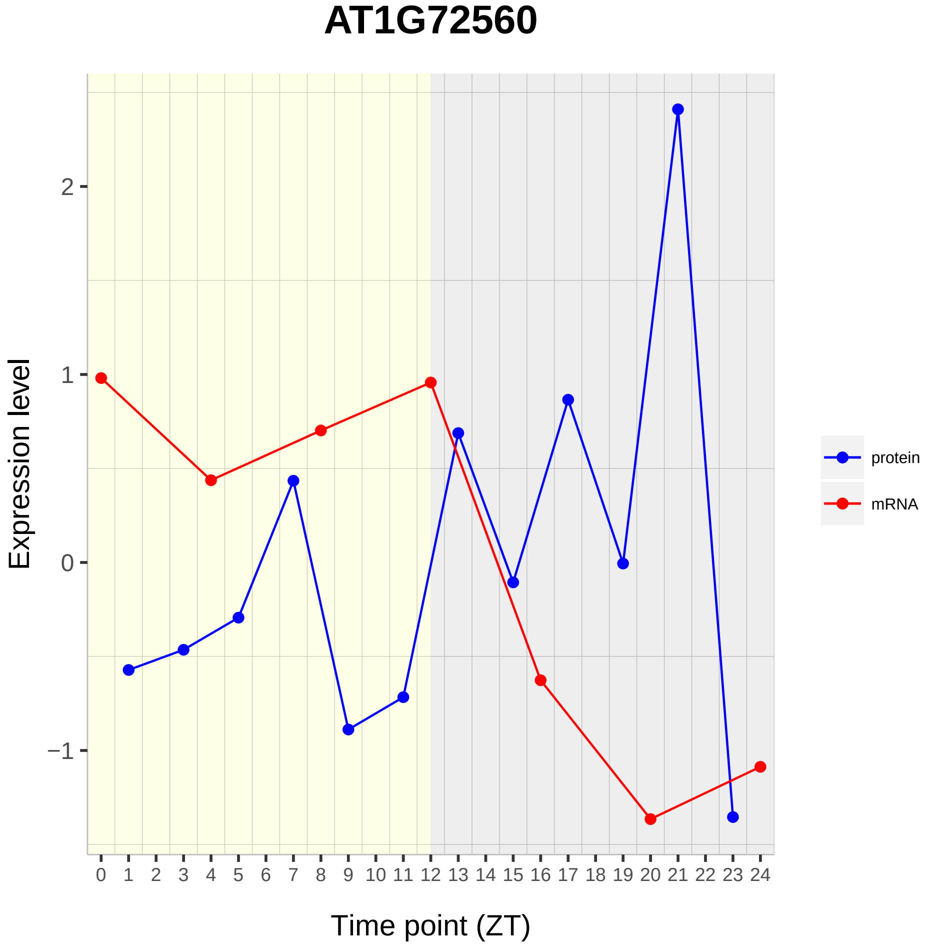

AT1G72560
Expression level
2
1
0
−1
Time point (ZT)
protein
mRNA

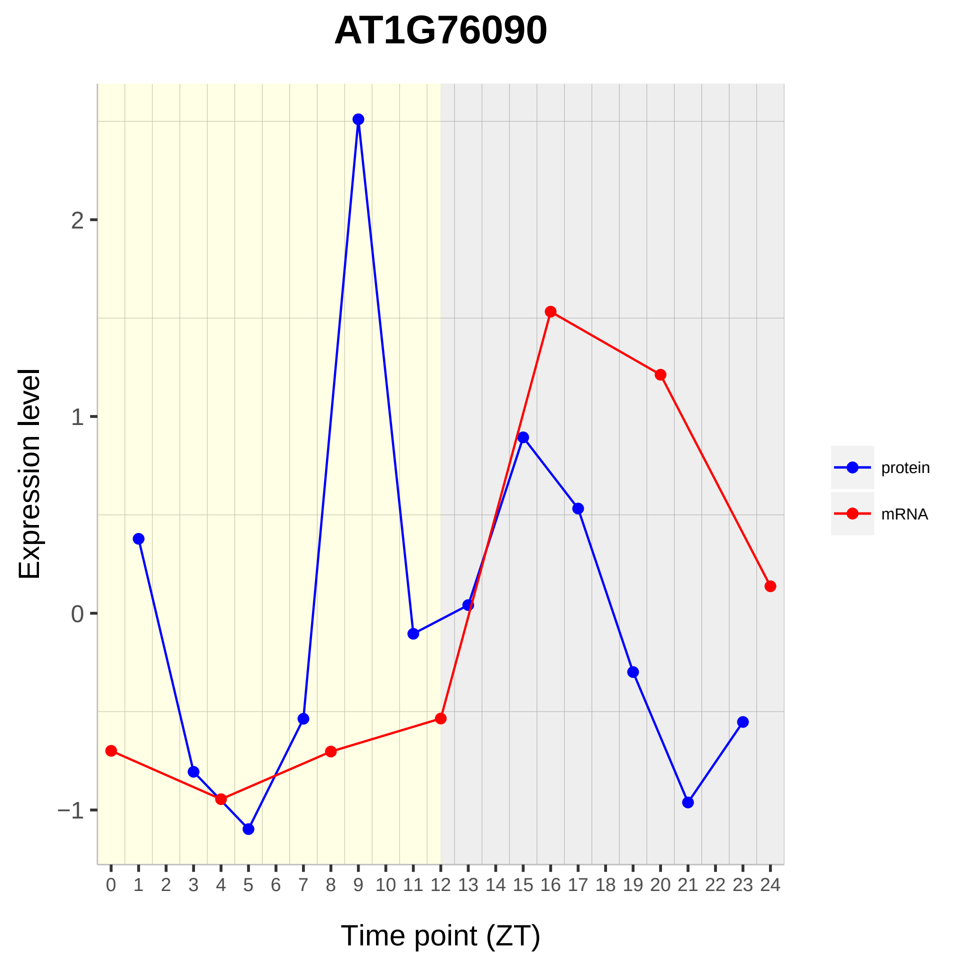

AT1G76090
Expression level
2
1
0
−1
0 1 2 3 4 5 6 7 8 9 10 11 12 13 14 15 16 17 18 19 20 21 22 23 24
Time point (ZT)
protein
mRNA

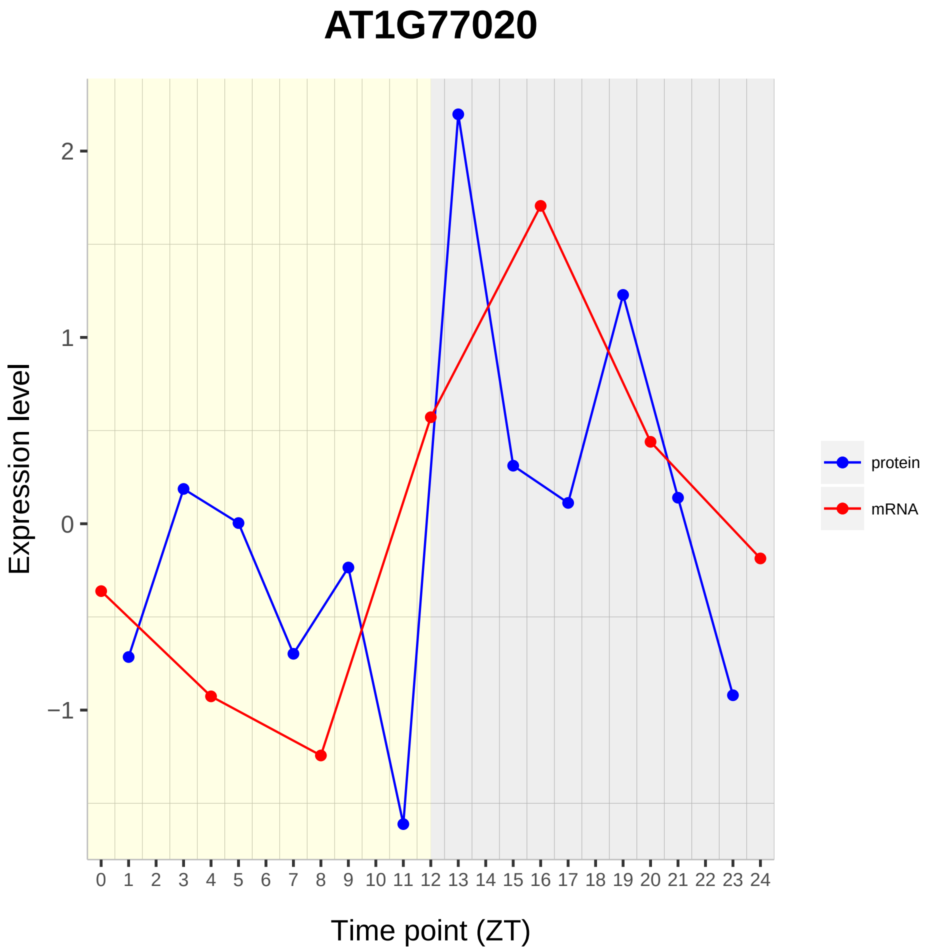

AT1G77020
Expression level
2
1
0
−1
0 1 2 3 4 5 6 7 8 9 10 11 12 13 14 15 16 17 18 19 20 21 22 23 24
Time point (ZT)
protein
mRNA

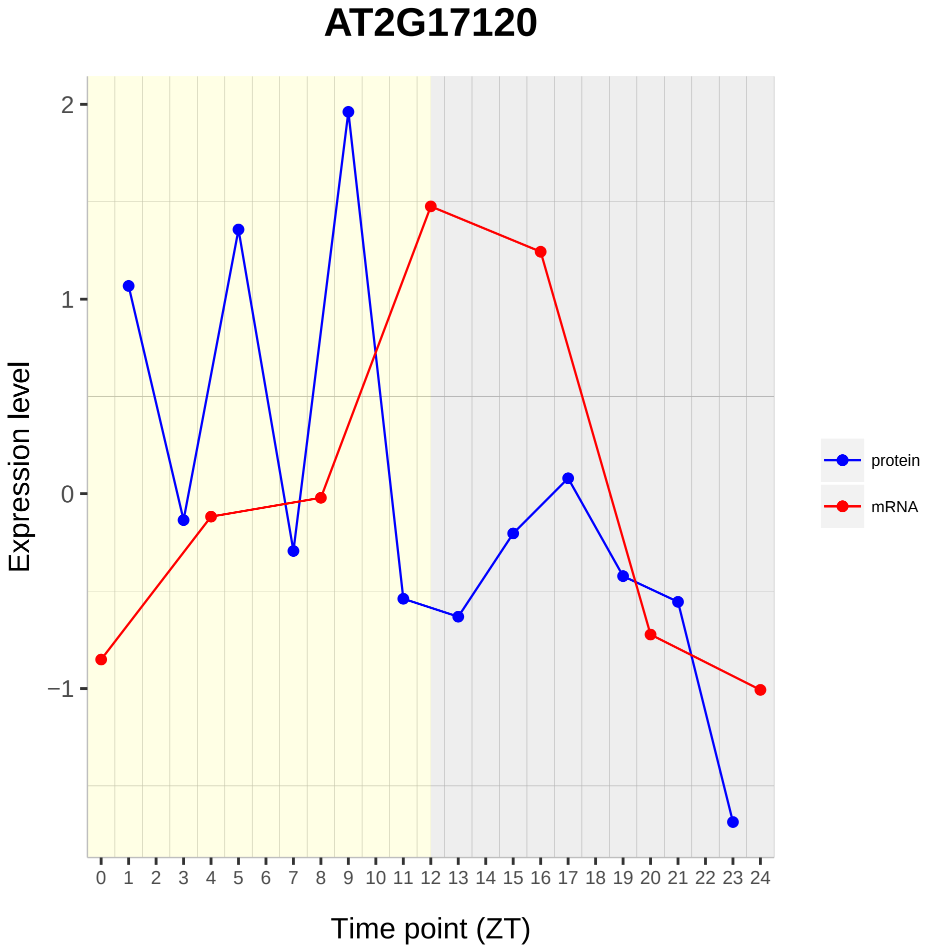

AT2G17120
Expression level
Time point (ZT)
2
1
0
−1
0 1 2 3 4 5 6 7 8 9 10 11 12 13 14 15 16 17 18 19 20 21 22 23 24
protein
mRNA

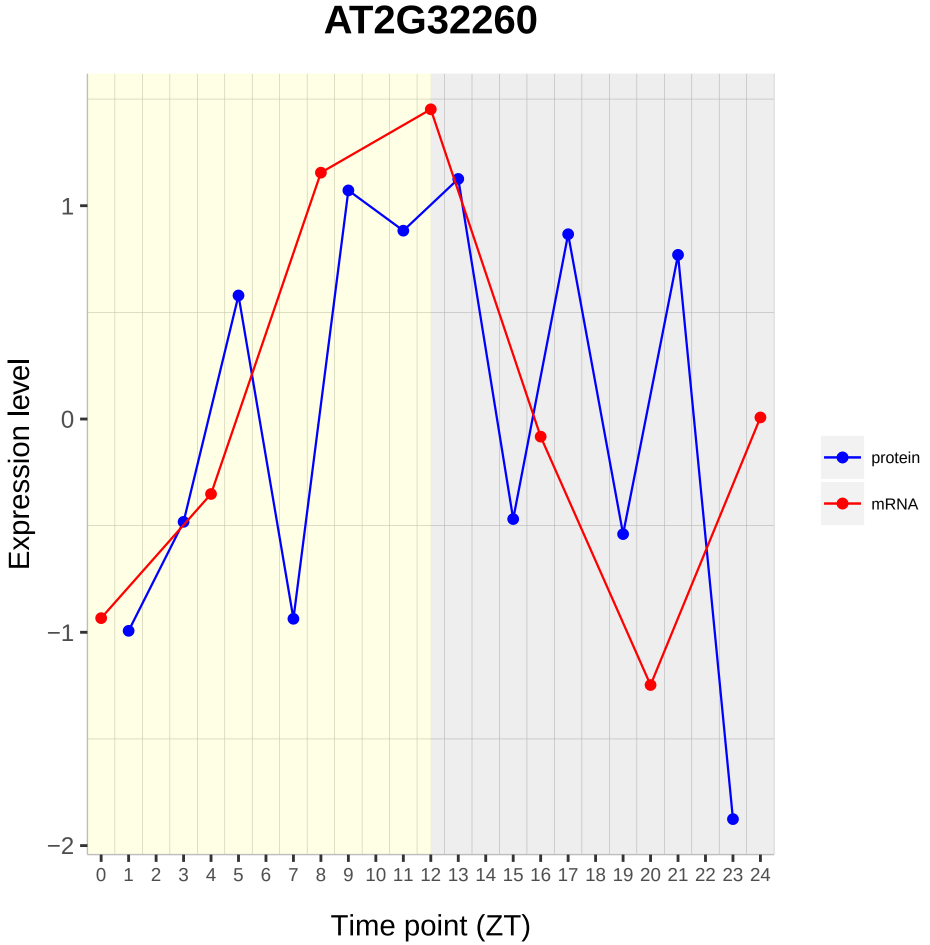

AT2G32260
Expression level
1
0
−1
−2
0 1 2 3 4 5 6 7 8 9 10 11 12 13 14 15 16 17 18 19 20 21 22 23 24
Time point (ZT)
protein
mRNA

# AT2G41090

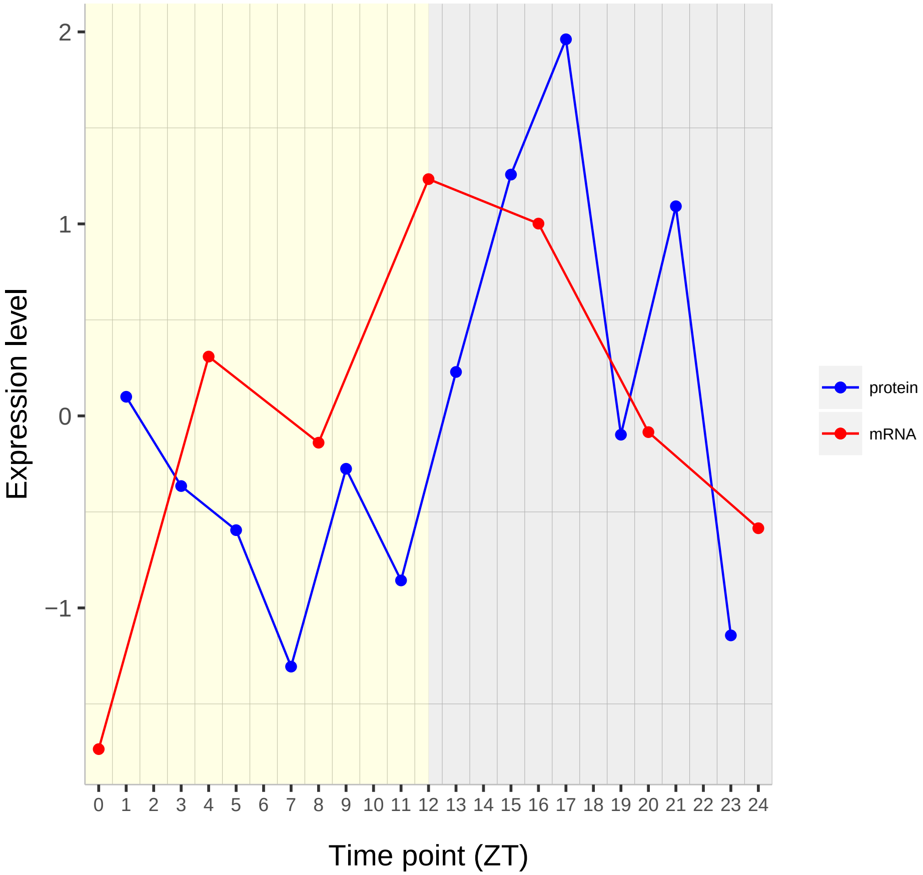

Expression level
2
1
0
−1
0 1 2 3 4 5 6 7 8 9 10 11 12 13 14 15 16 17 18 19 20 21 22 23 24
Time point (ZT)
protein
mRNA

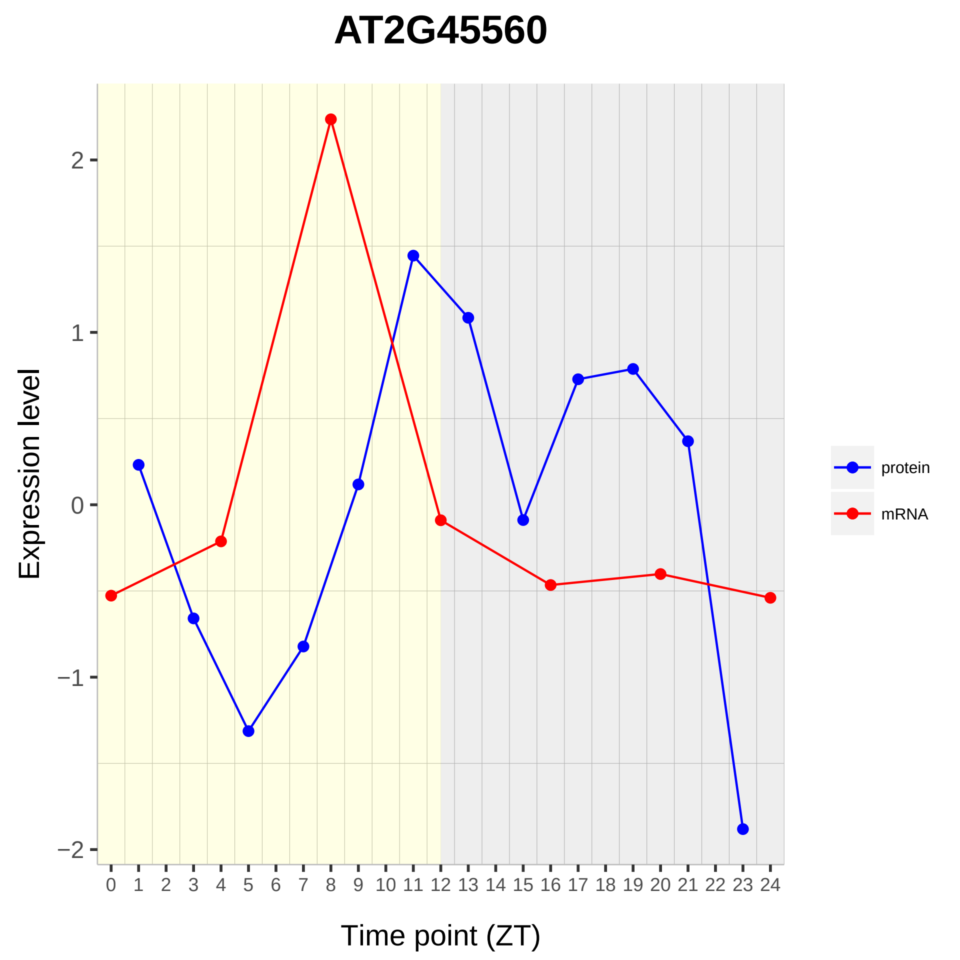

AT2G45560
Expression level
Time point (ZT)
2
1
0
−1
−2
0 1 2 3 4 5 6 7 8 9 10 11 12 13 14 15 16 17 18 19 20 21 22 23 24
protein
mRNA

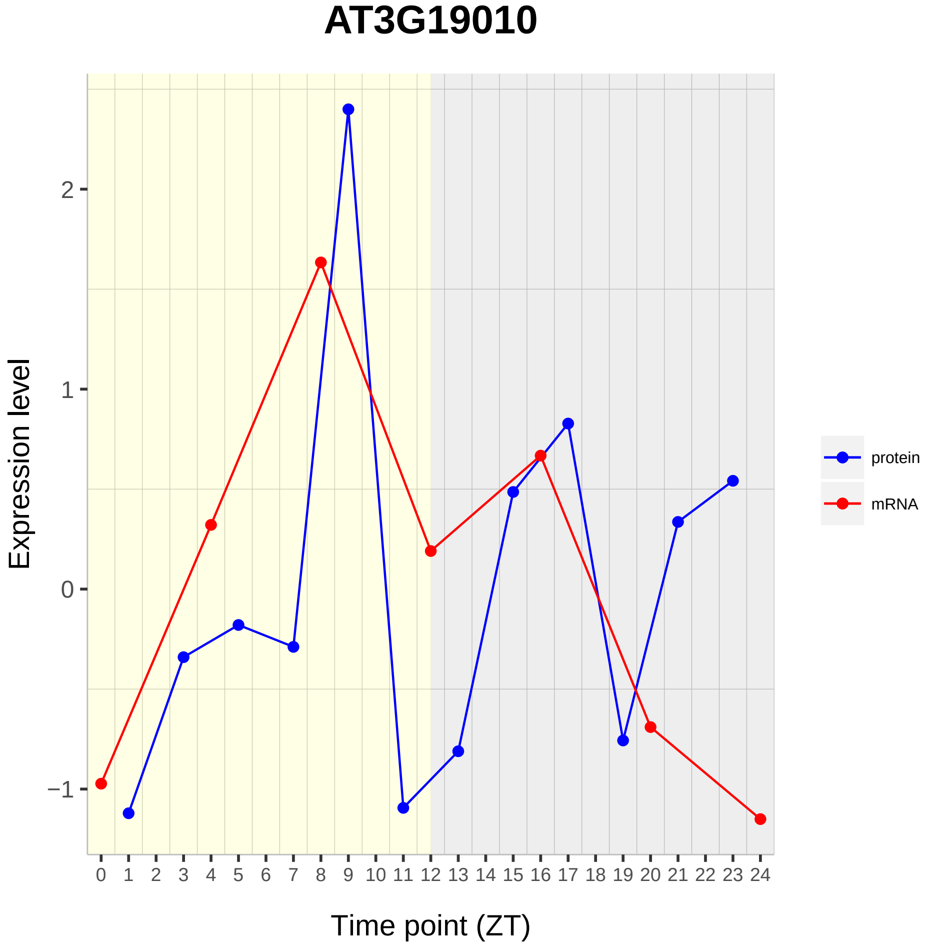

AT3G19010
Expression level
Time point (ZT)
2
1
0
−1
protein
mRNA

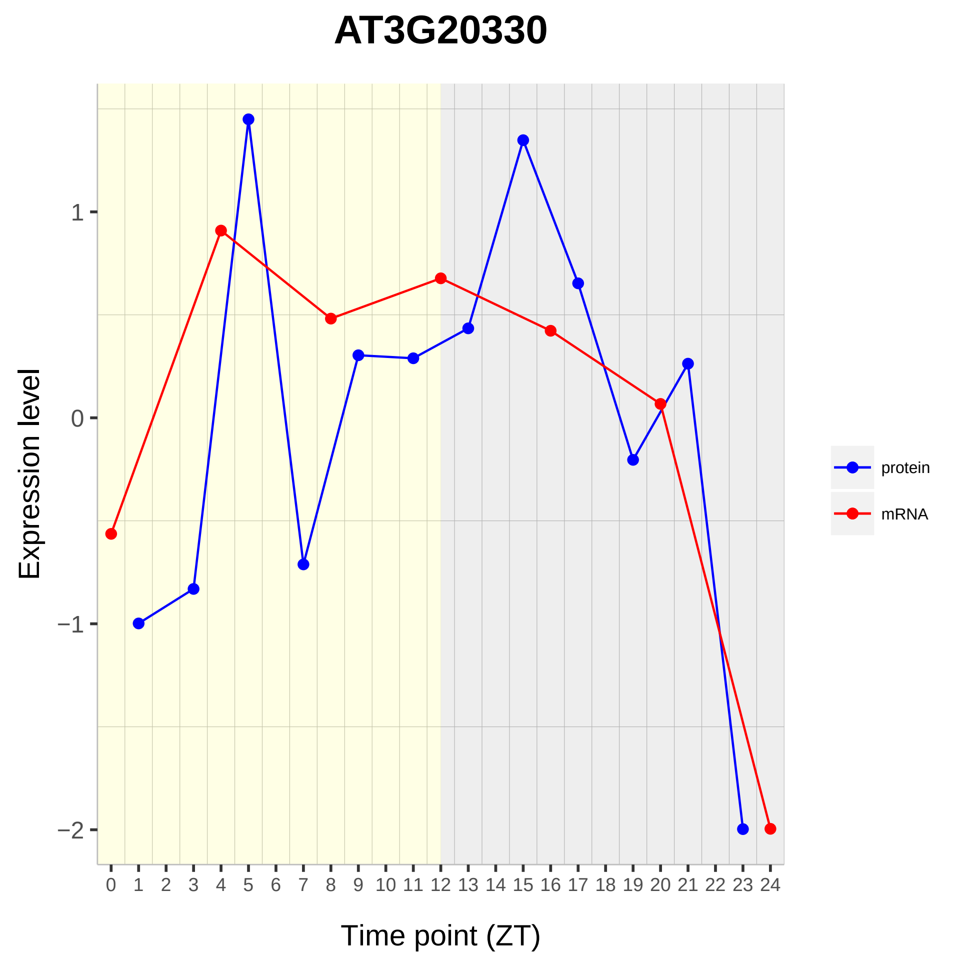

AT3G20330
Expression level
1
0
−1
−2
Time point (ZT)
protein
mRNA

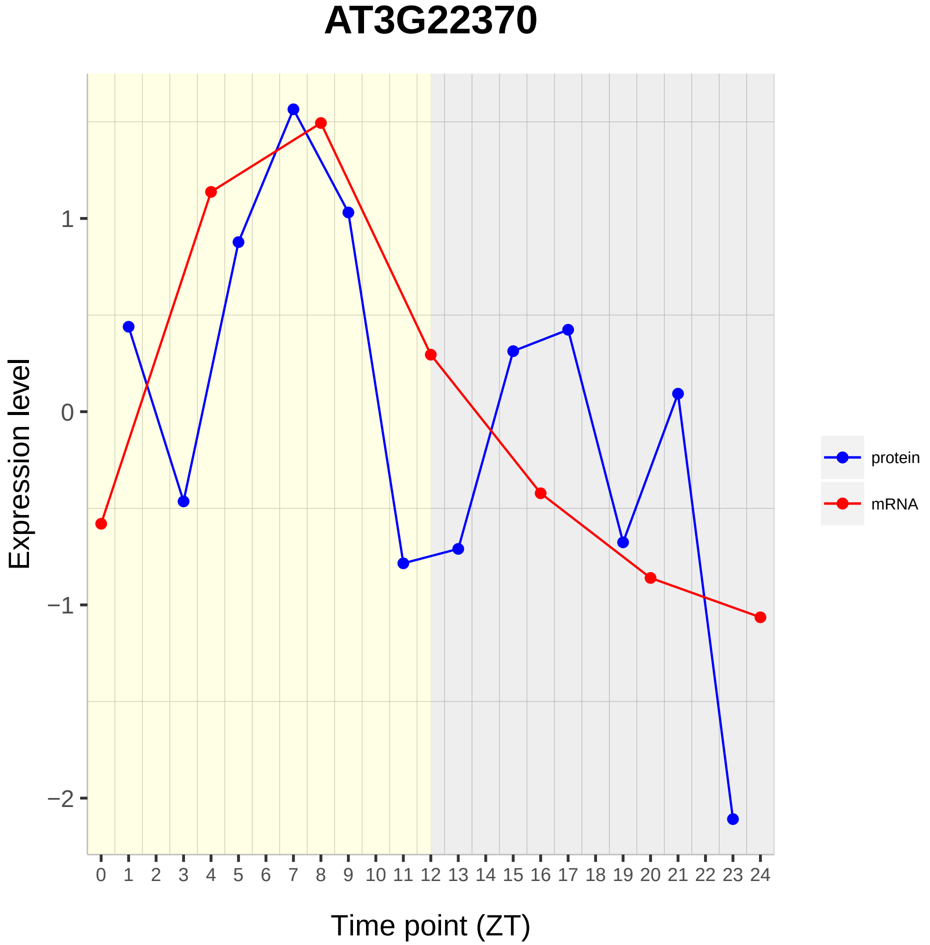
AT3G22370
Expression level
1
0
−1
−2
0 1 2 3 4 5 6 7 8 9 10 11 12 13 14 15 16 17 18 19 20 21 22 23 24
Time point (ZT)
protein
mRNA

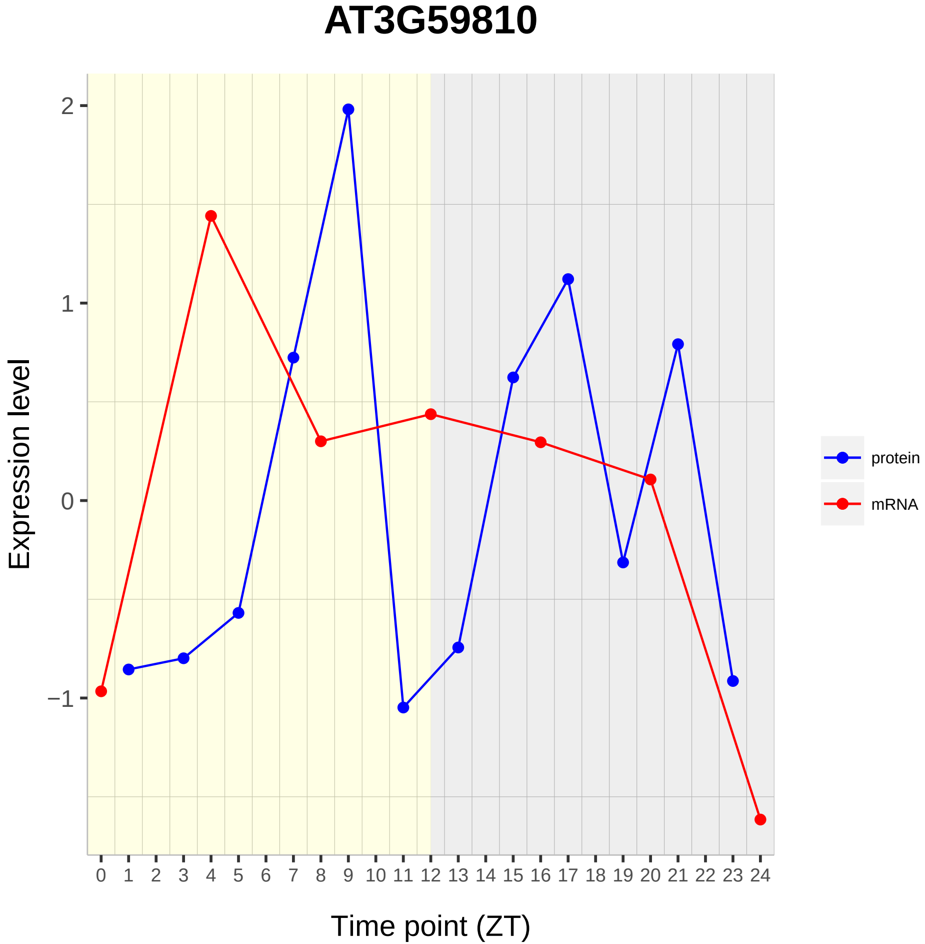
AT3G59810
Expression level
Time point (ZT)
2
1
0
−1
0 1 2 3 4 5 6 7 8 9 10 11 12 13 14 15 16 17 18 19 20 21 22 23 24
protein
mRNA

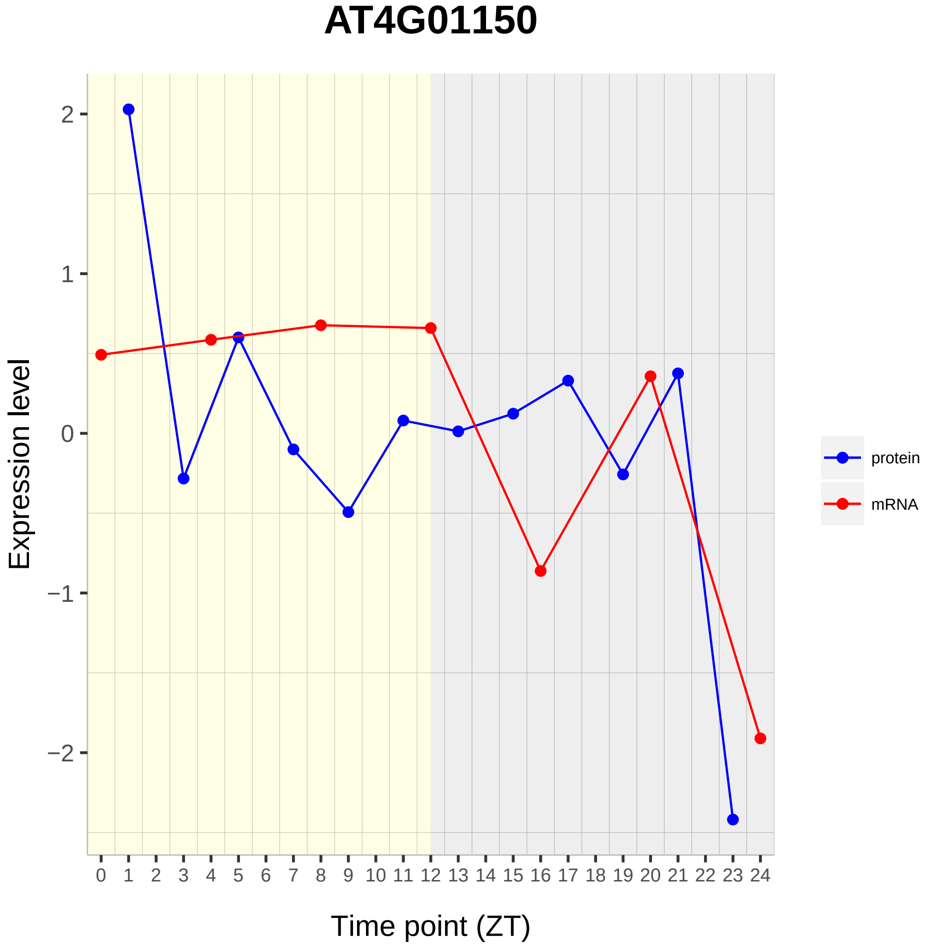

AT4G01150
Expression level
2
1
0
−1
−2
0 1 2 3 4 5 6 7 8 9 10 11 12 13 14 15 16 17 18 19 20 21 22 23 24
Time point (ZT)
protein
mRNA

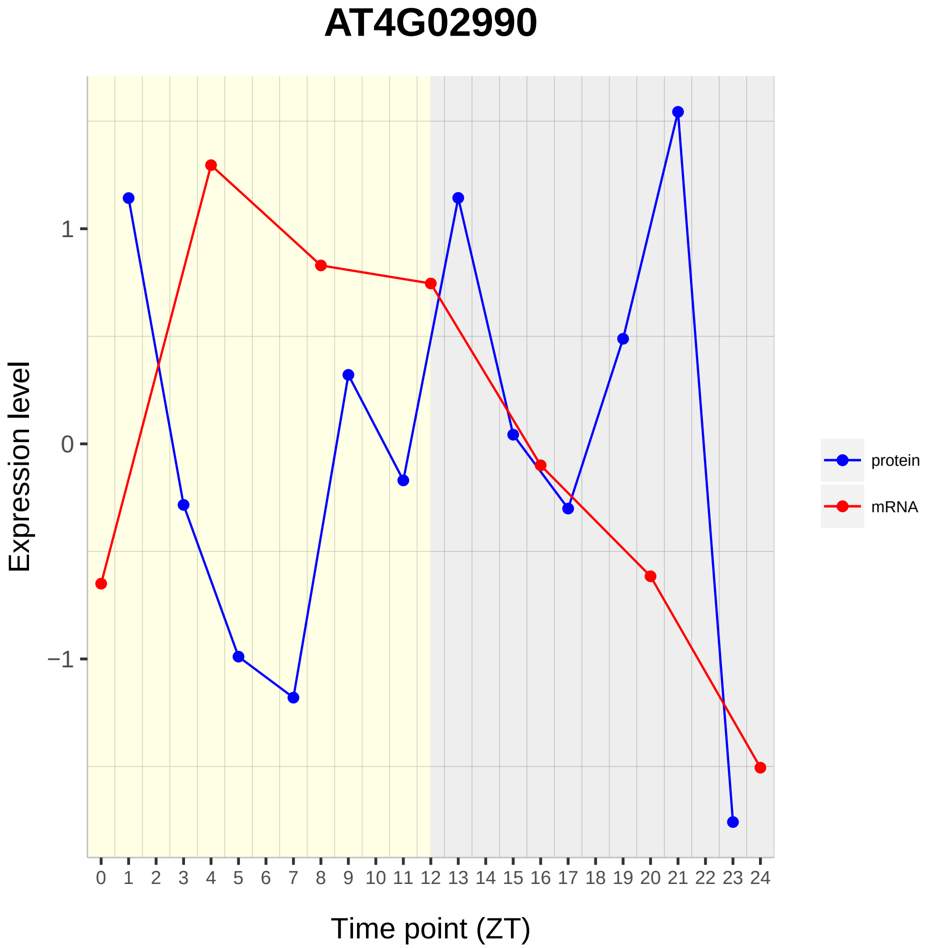
AT4G02990
Expression level
1
0
−1
0 1 2 3 4 5 6 7 8 9 10 11 12 13 14 15 16 17 18 19 20 21 22 23 24
Time point (ZT)
protein
mRNA

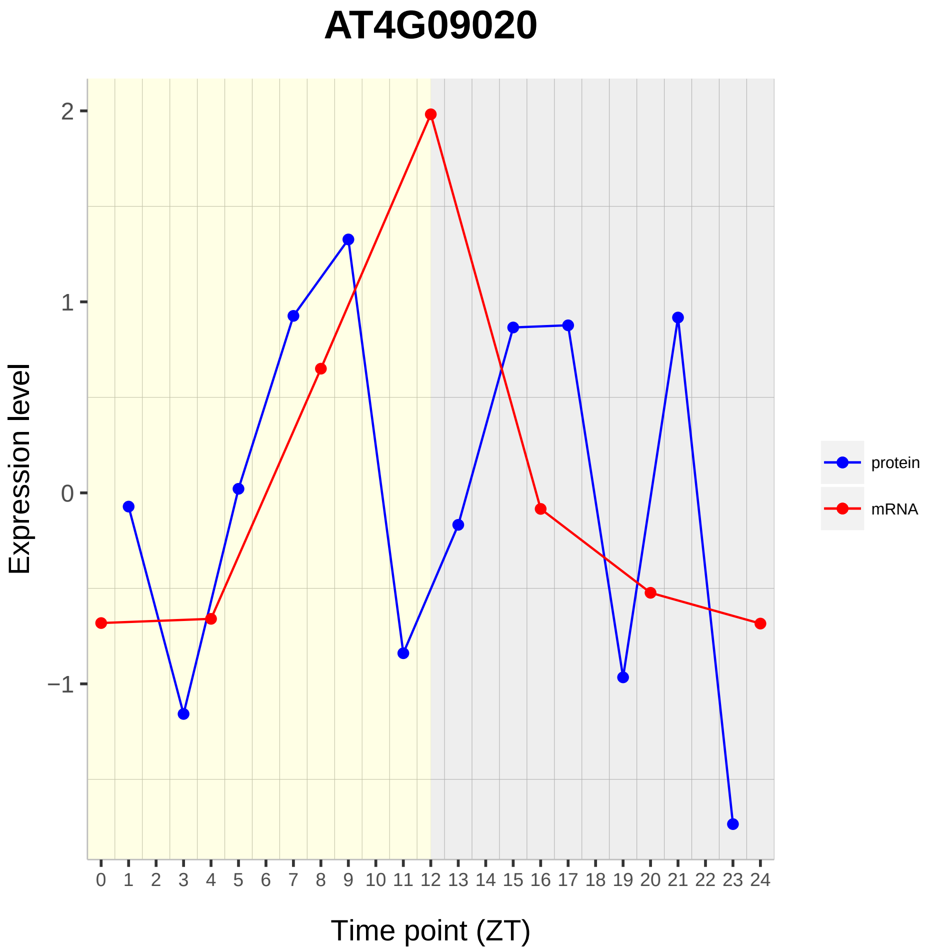
AT4G09020
Expression level
Time point (ZT)
protein
mRNA
2
1
0
−1
0 1 2 3 4 5 6 7 8 9 10 11 12 13 14 15 16 17 18 19 20 21 22 23 24

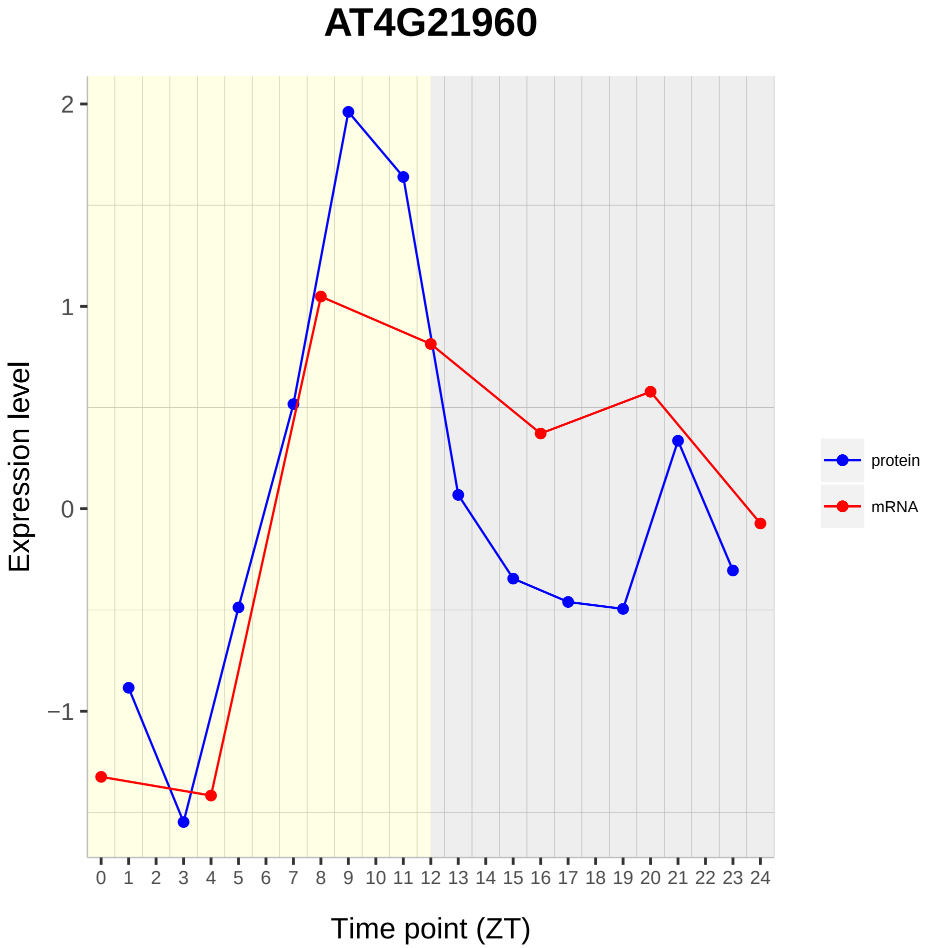
AT4G21960
Expression level
Time point (ZT)
2
1
0
−1
protein
mRNA

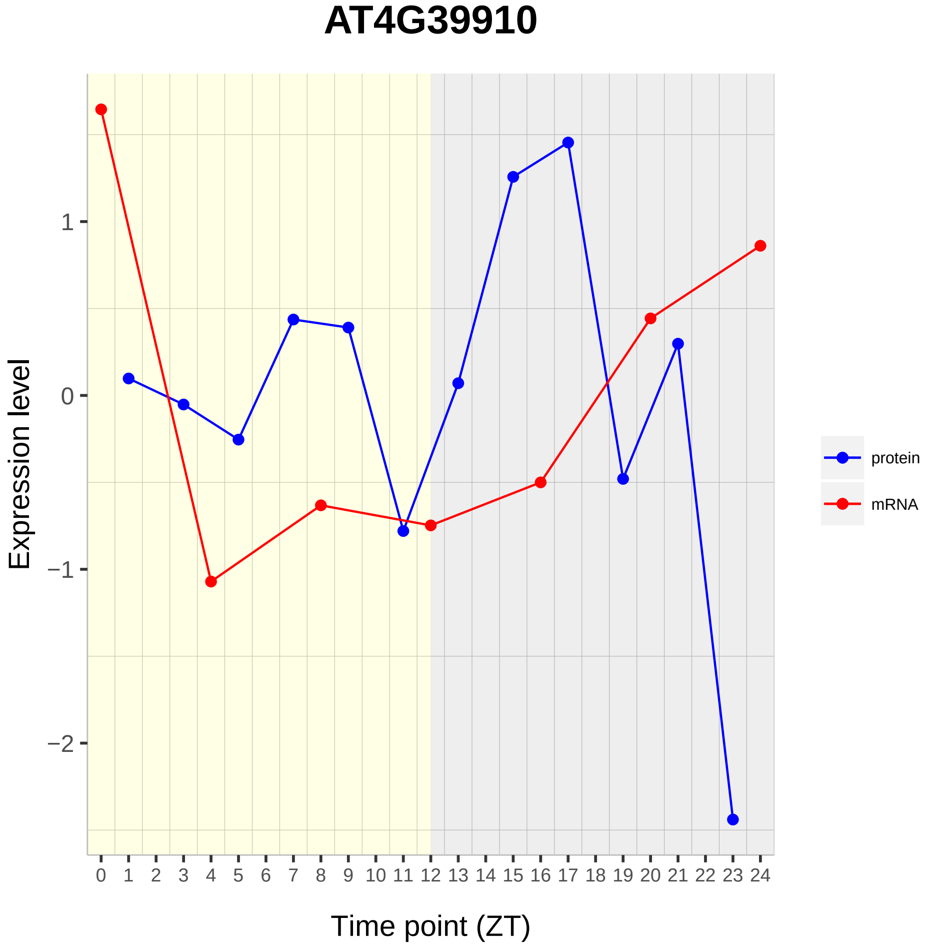

AT4G39910
Expression level
1
0
−1
−2
Time point (ZT)
protein
mRNA

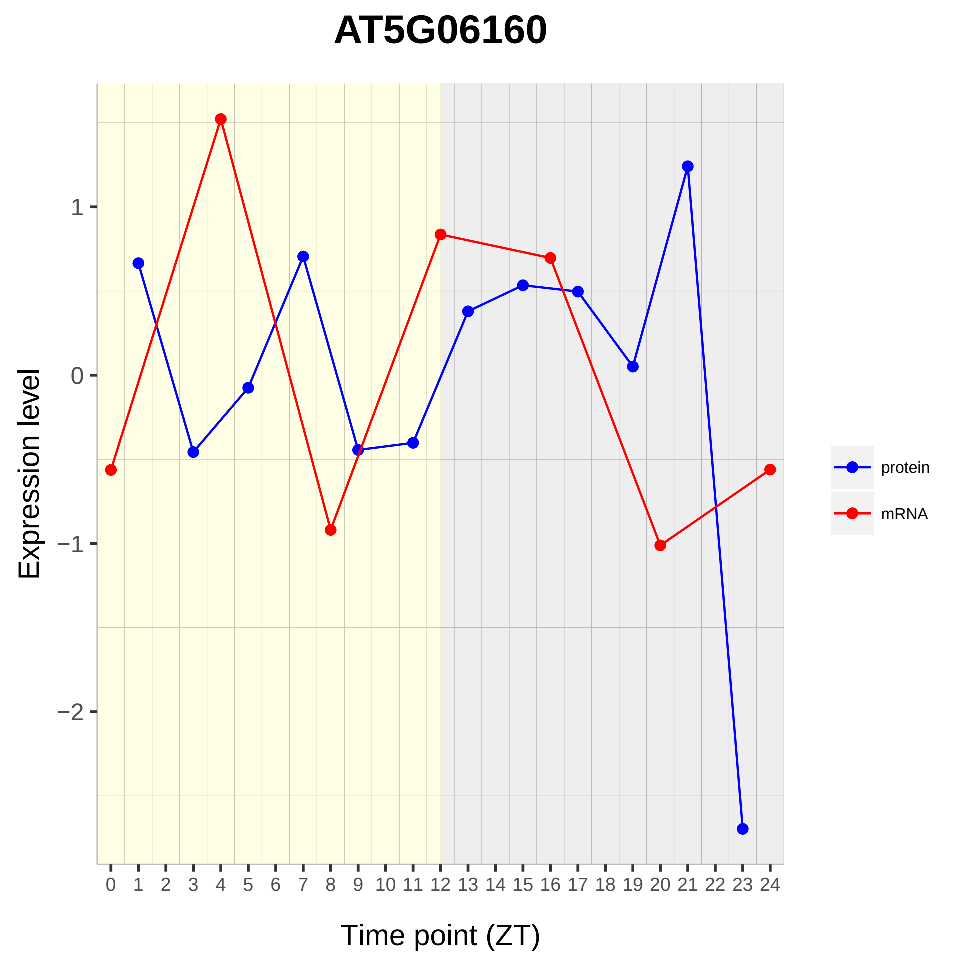

AT5G06160
Expression level
Time point (ZT)
1
0
−1
−2
0 1 2 3 4 5 6 7 8 9 10 11 12 13 14 15 16 17 18 19 20 21 22 23 24
protein
mRNA

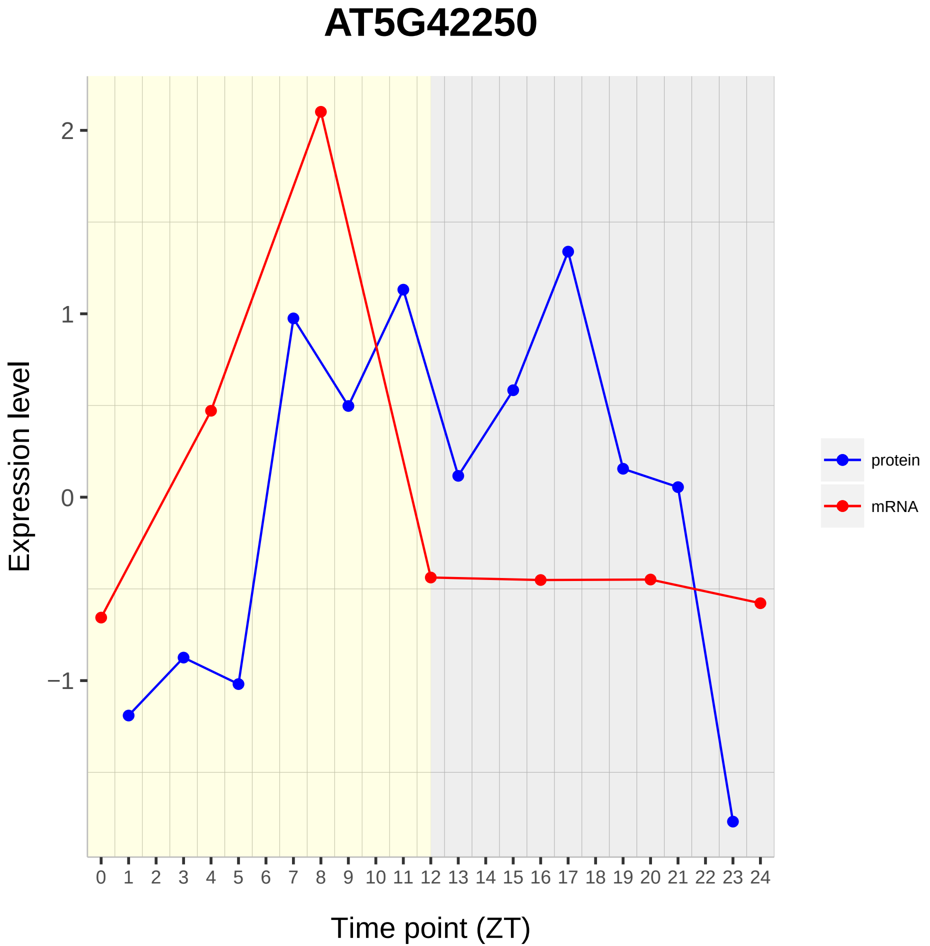
AT5G42250
Expression level
2
1
0
−1
0 1 2 3 4 5 6 7 8 9 10 11 12 13 14 15 16 17 18 19 20 21 22 23 24
Time point (ZT)
protein
mRNA

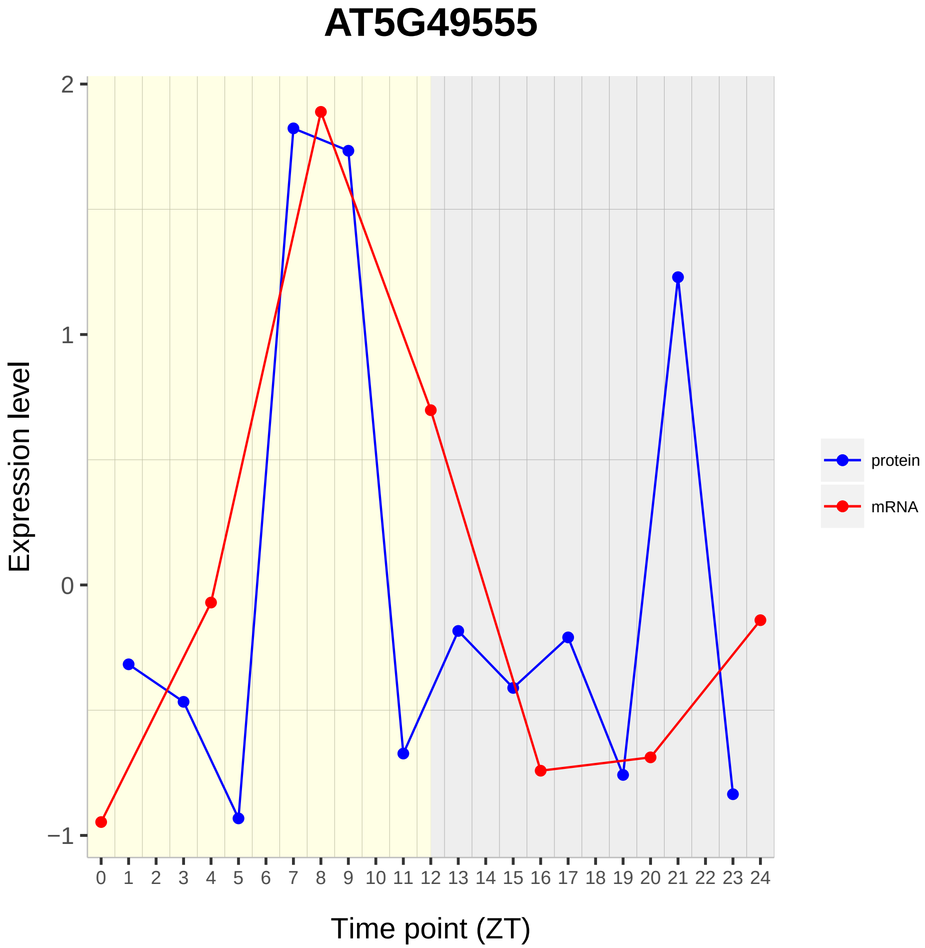
AT5G49555
Expression level
2
1
0
−1
0 1 2 3 4 5 6 7 8 9 10 11 12 13 14 15 16 17 18 19 20 21 22 23 24
Time point (ZT)
protein
mRNA

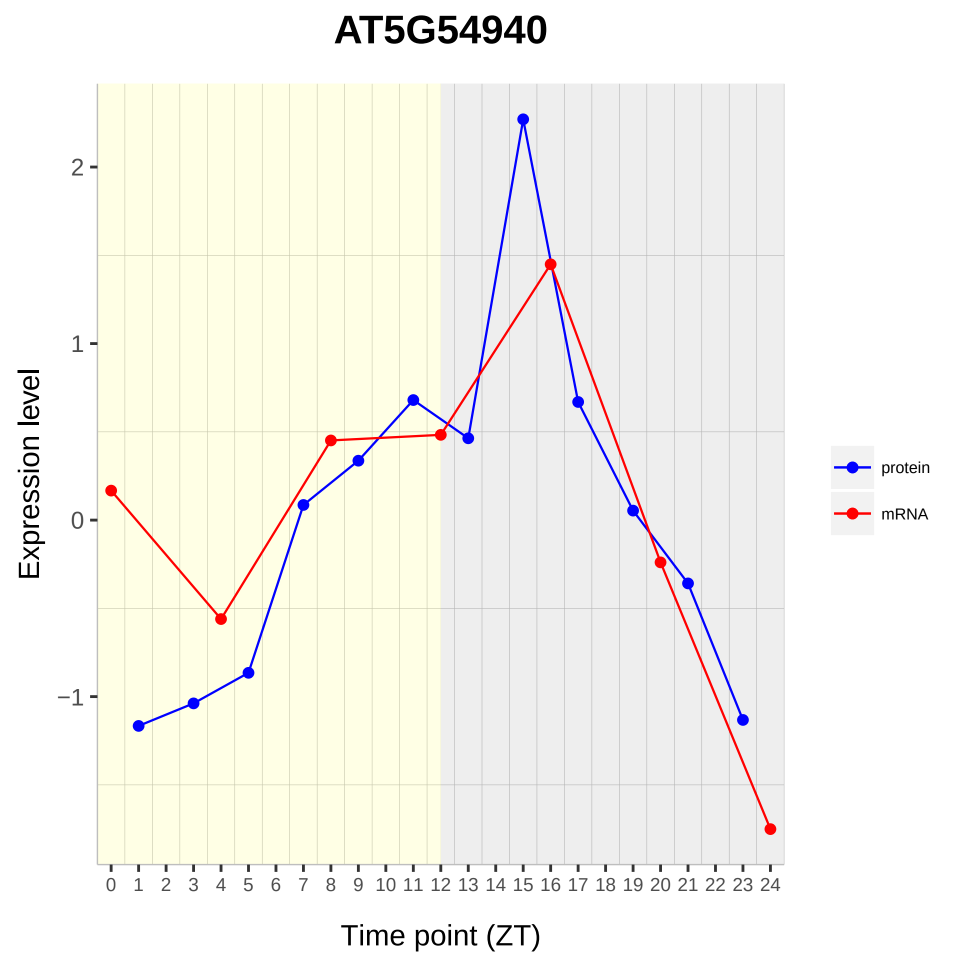

AT5G54940
Expression level
2
1
0
−1
Time point (ZT)
protein
mRNA

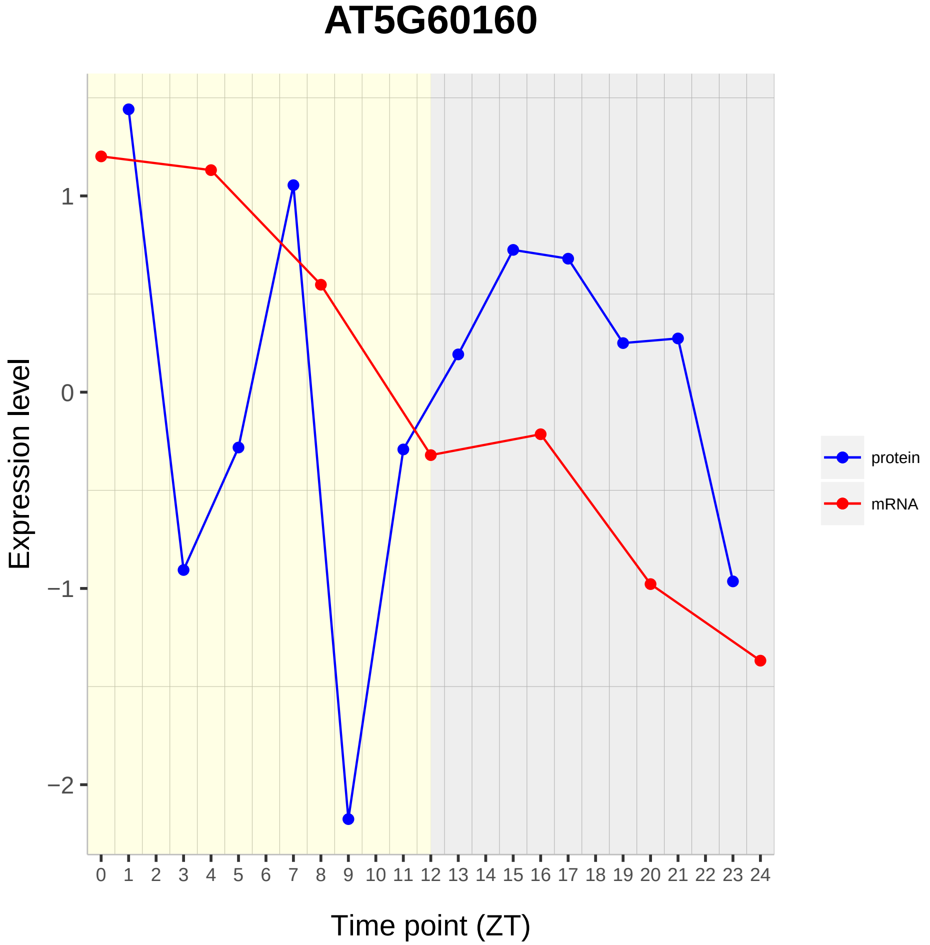

AT5G60160
Expression level
1
0
−1
−2
0 1 2 3 4 5 6 7 8 9 10 11 12 13 14 15 16 17 18 19 20 21 22 23 24
Time point (ZT)
protein
mRNA

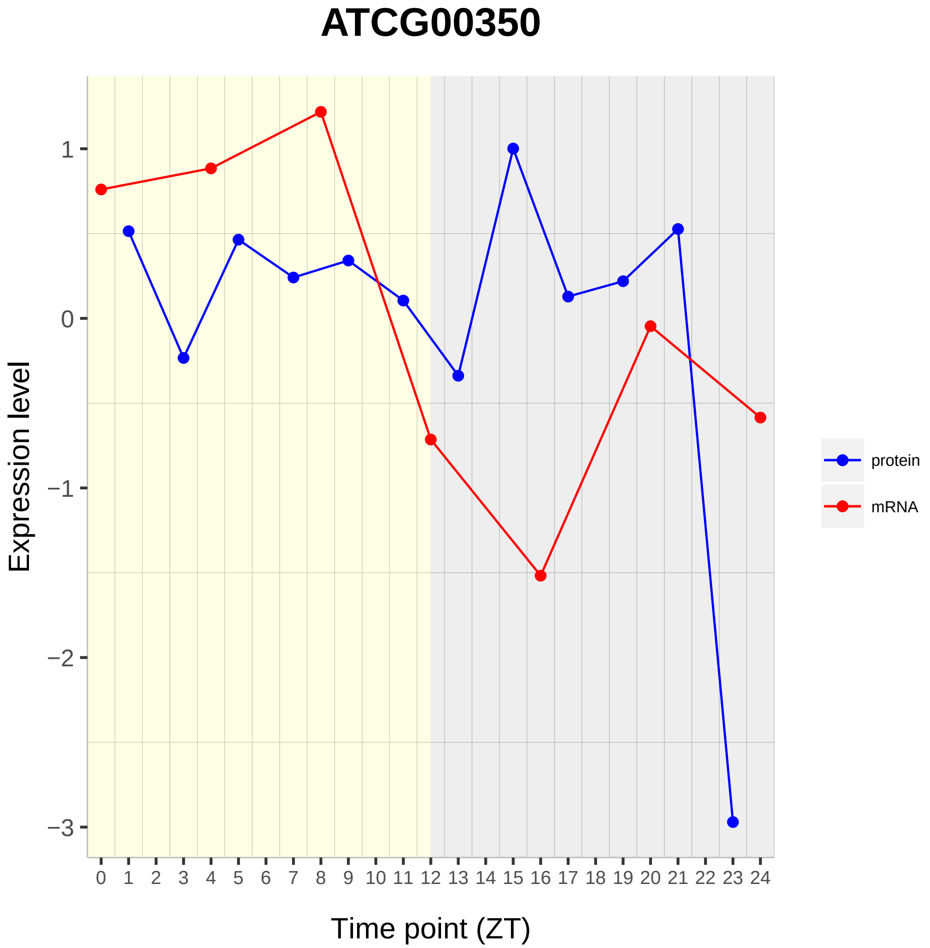

ATCG00350
Expression level
1
0
−1
−2
−3
0 1 2 3 4 5 6 7 8 9 10 11 12 13 14 15 16 17 18 19 20 21 22 23 24
Time point (ZT)
protein
mRNA
