## Supplementary material for "Diurnal Dynamics of the Arabidopsis Rosette Proteome and Phosphoproteome": SuppData2

# AT1G02100

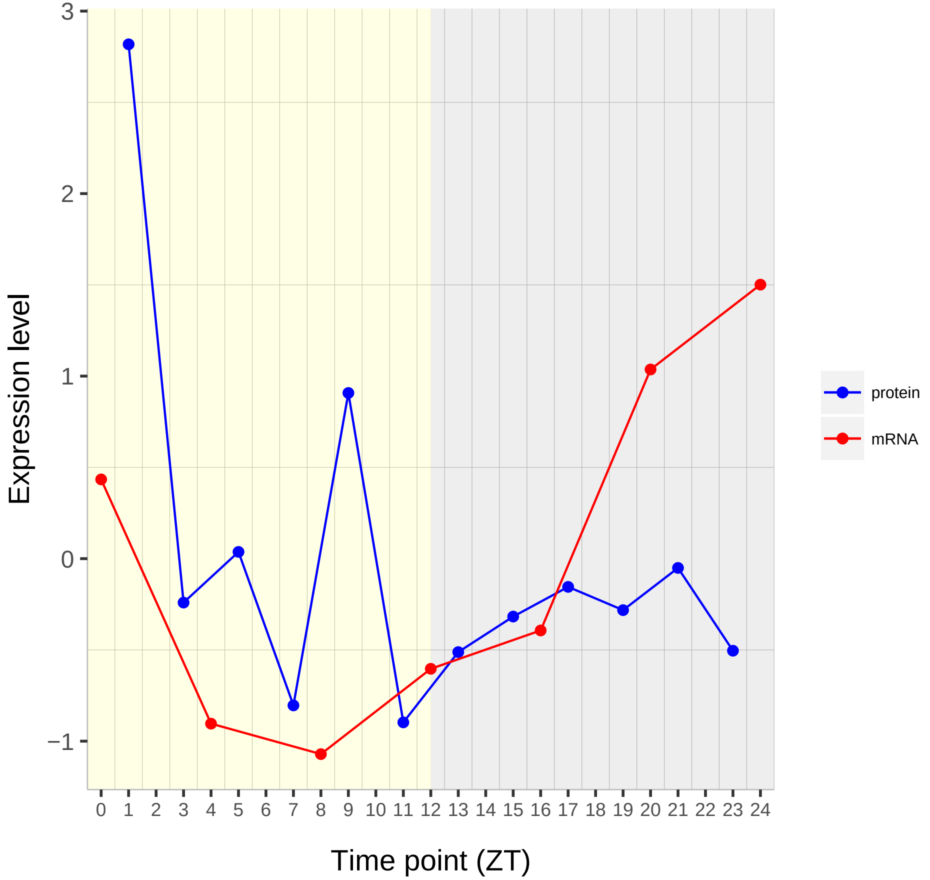

Expression level
3
2
1
0
−1
0 1 2 3 4 5 6 7 8 9 10 11 12 13 14 15 16 17 18 19 20 21 22 23 24
Time point (ZT)
protein
mRNA

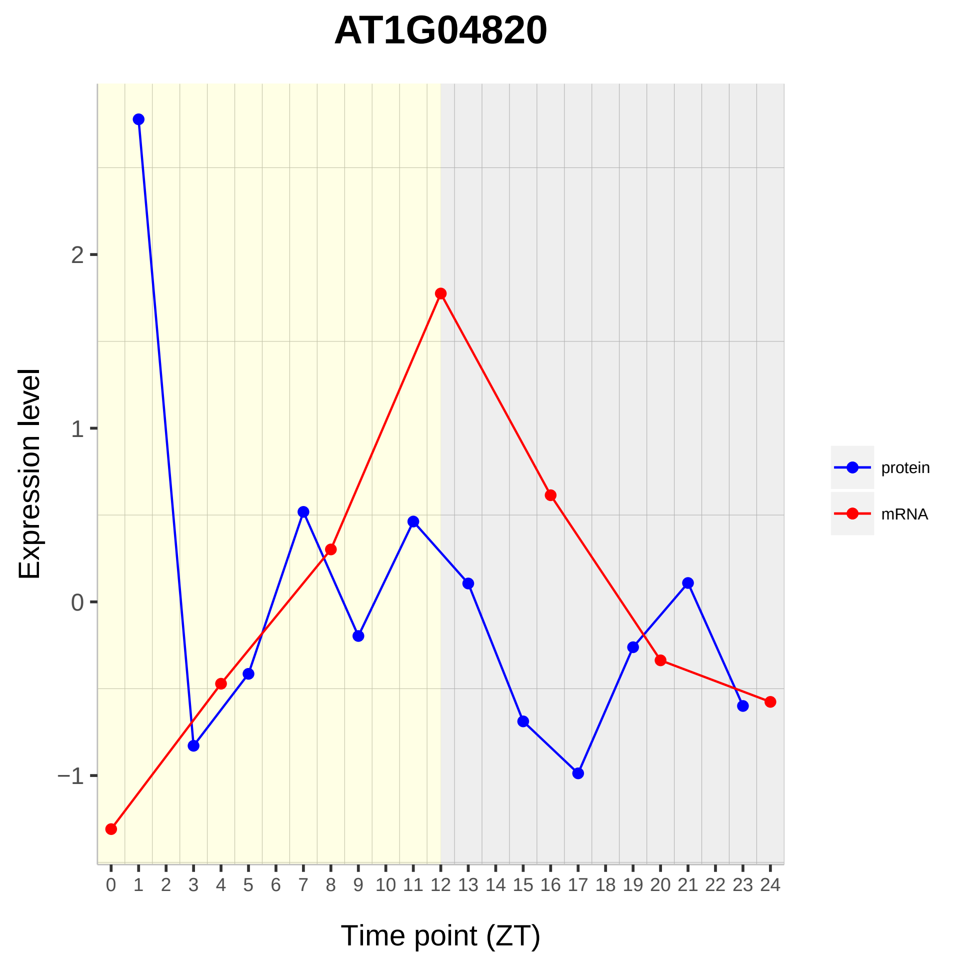

AT1G04820
Expression level
2
1
0
−1
Time point (ZT)
protein
mRNA

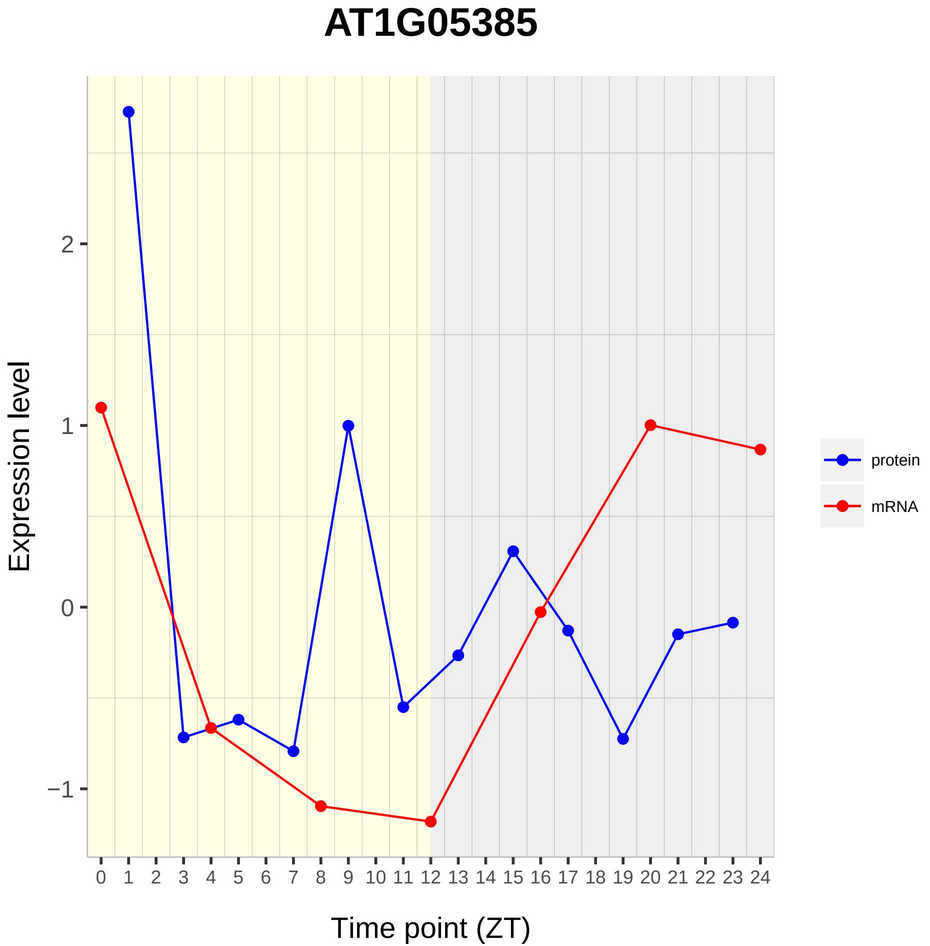

AT1G05385
Expression level
2
1
0
−1
Time point (ZT)
protein
mRNA

# AT1G05520

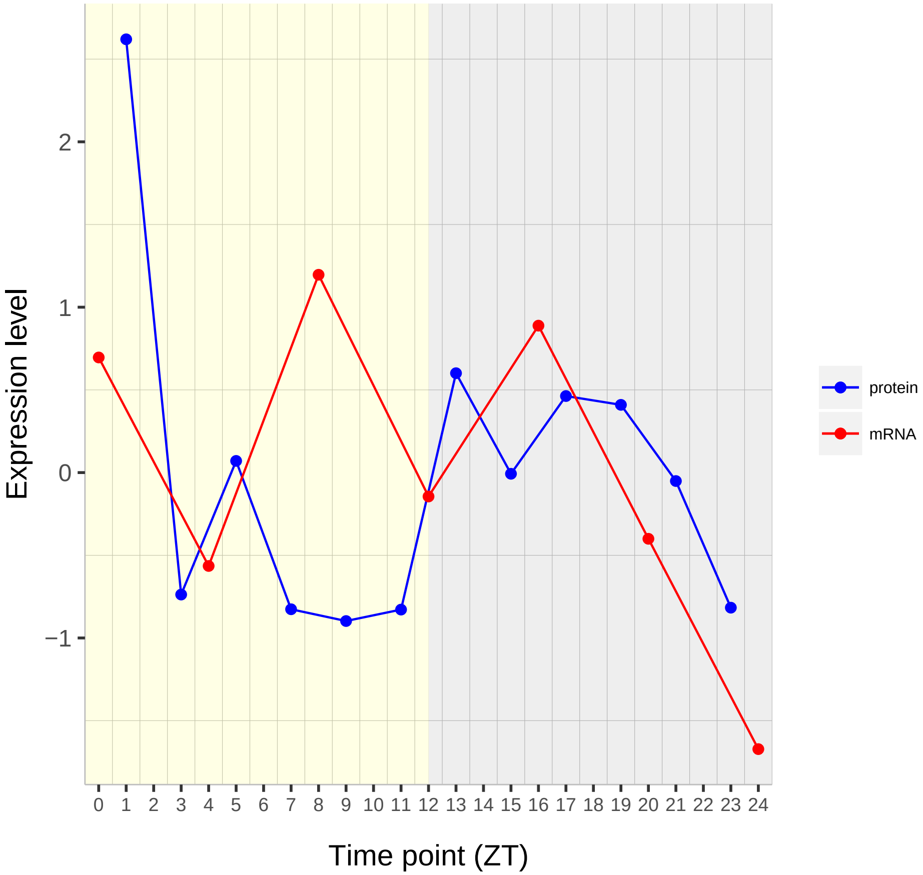

Expression level
2
1
0
−1
Time point (ZT)
protein
mRNA

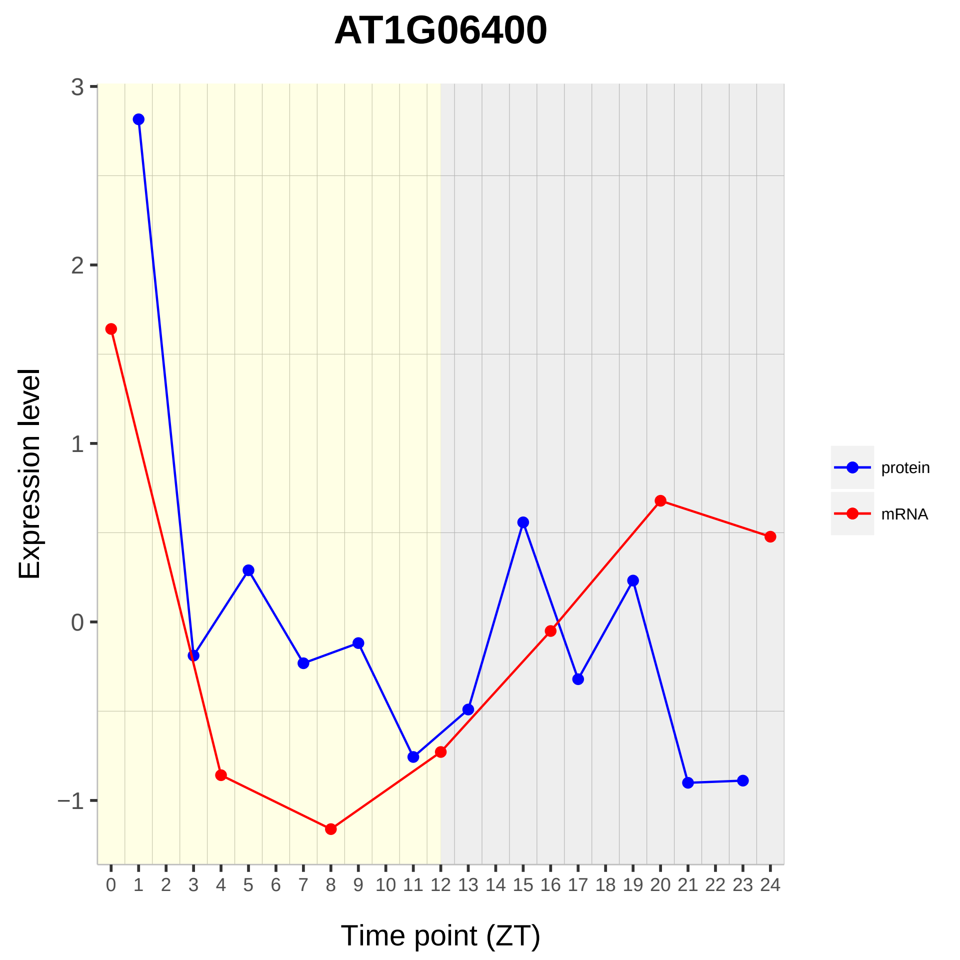

AT1G06400
Expression level
Time point (ZT)
3
2
1
0
−1
0 1 2 3 4 5 6 7 8 9 10 11 12 13 14 15 16 17 18 19 20 21 22 23 24
protein
mRNA

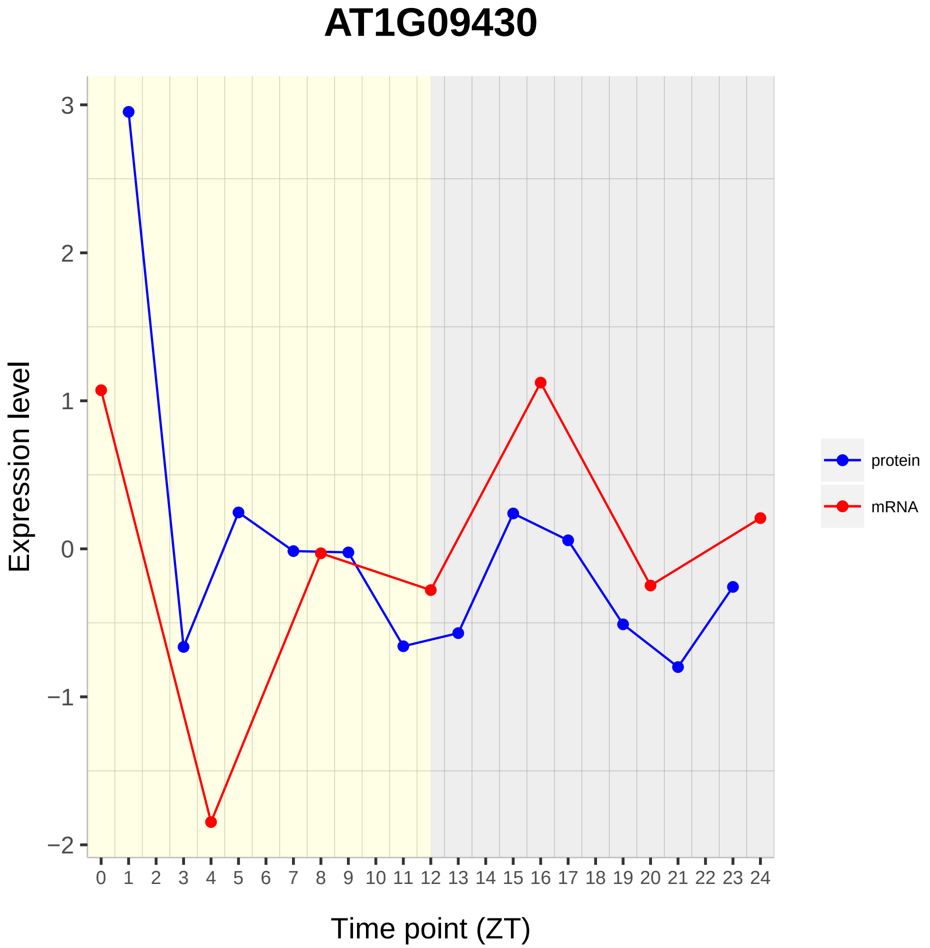

AT1G09430
Expression level
3
2
1
0
−1
−2
0 1 2 3 4 5 6 7 8 9 10 11 12 13 14 15 16 17 18 19 20 21 22 23 24
Time point (ZT)
protein
mRNA

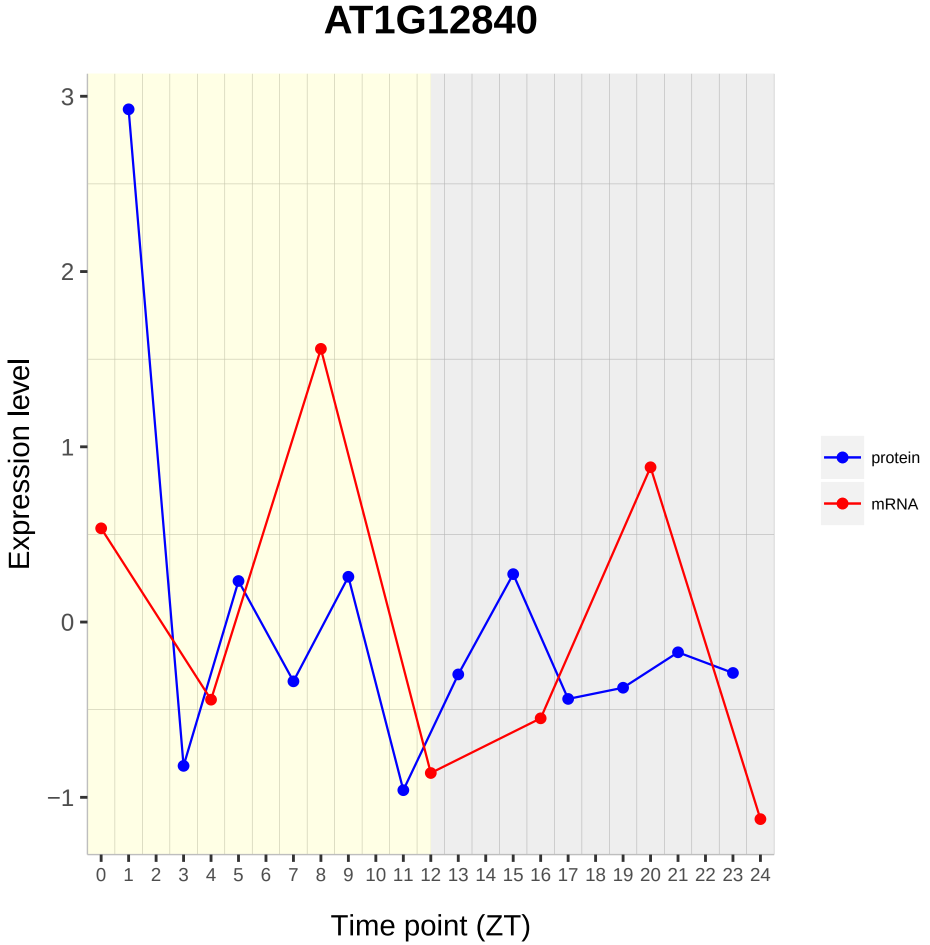
AT1G12840
Expression level
Time point (ZT)
protein
mRNA
3
2
1
0
−1
0 1 2 3 4 5 6 7 8 9 10 11 12 13 14 15 16 17 18 19 20 21 22 23 24

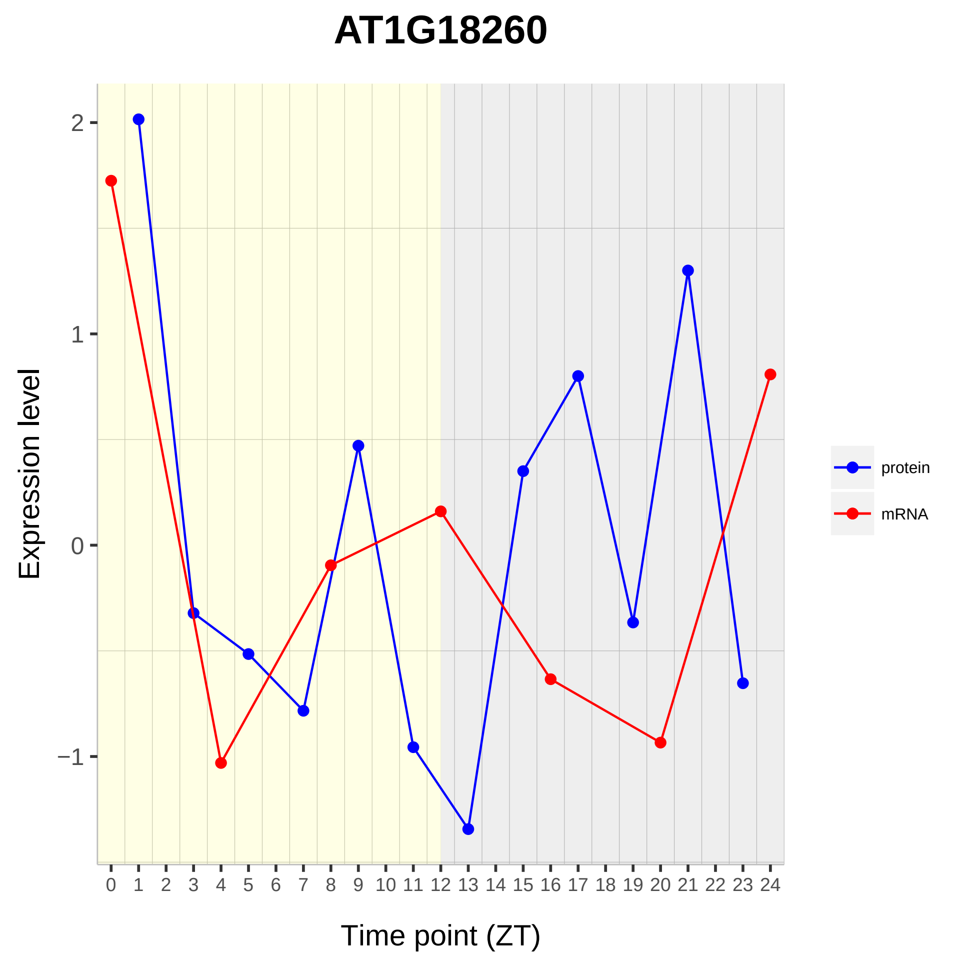

AT1G18260
Expression level
2
1
0
−1
0 1 2 3 4 5 6 7 8 9 10 11 12 13 14 15 16 17 18 19 20 21 22 23 24
Time point (ZT)
protein
mRNA

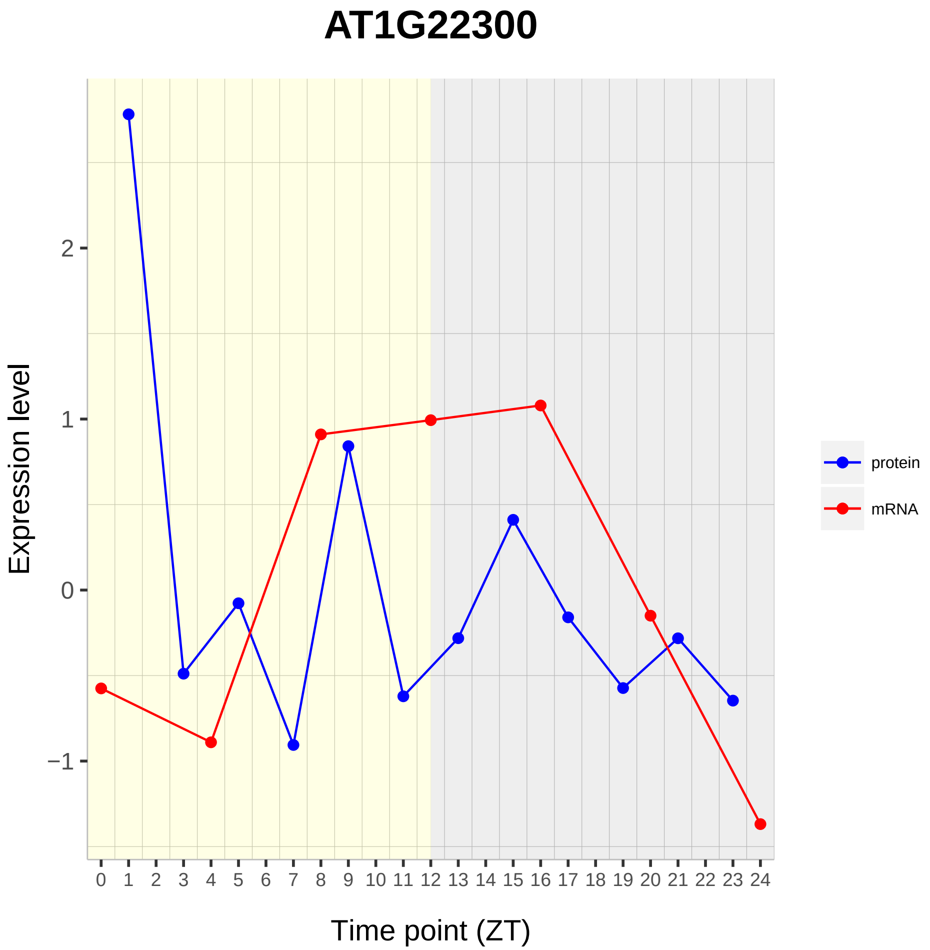
AT1G22300
Expression level
2
1
0
−1
0 1 2 3 4 5 6 7 8 9 10 11 12 13 14 15 16 17 18 19 20 21 22 23 24
Time point (ZT)
protein
mRNA

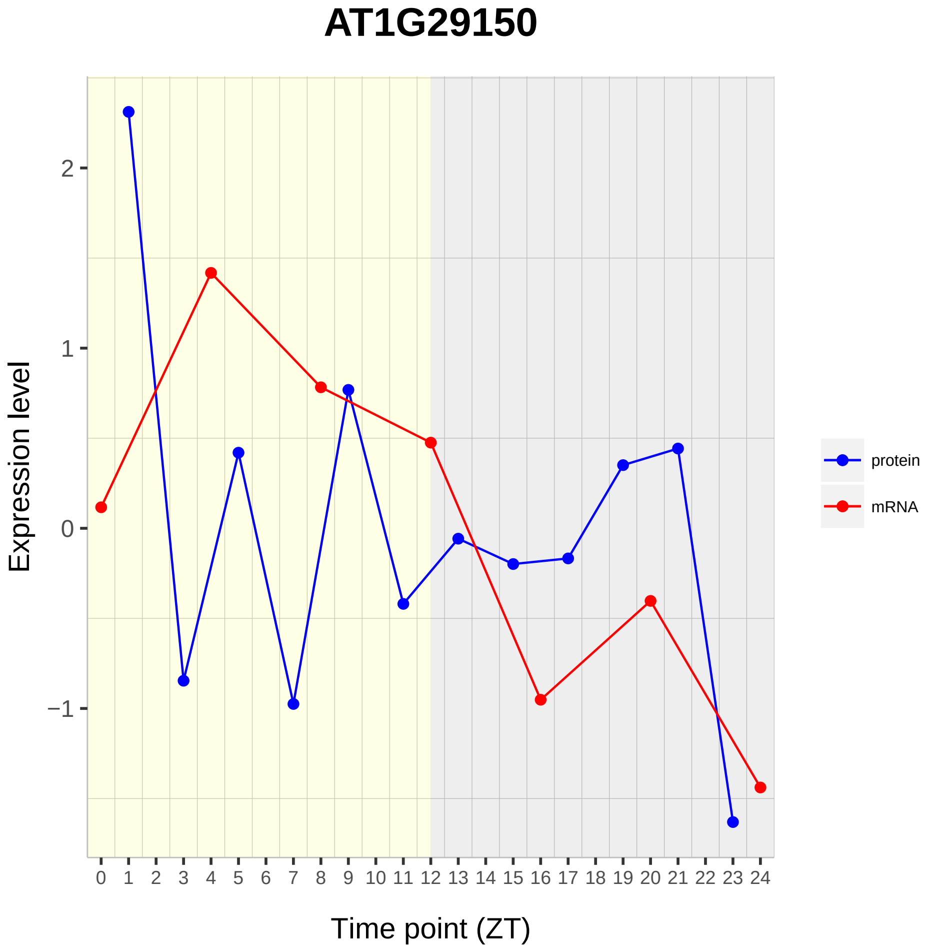

AT1G29150
Expression level
2
1
0
−1
0 1 2 3 4 5 6 7 8 9 10 11 12 13 14 15 16 17 18 19 20 21 22 23 24
Time point (ZT)
protein
mRNA

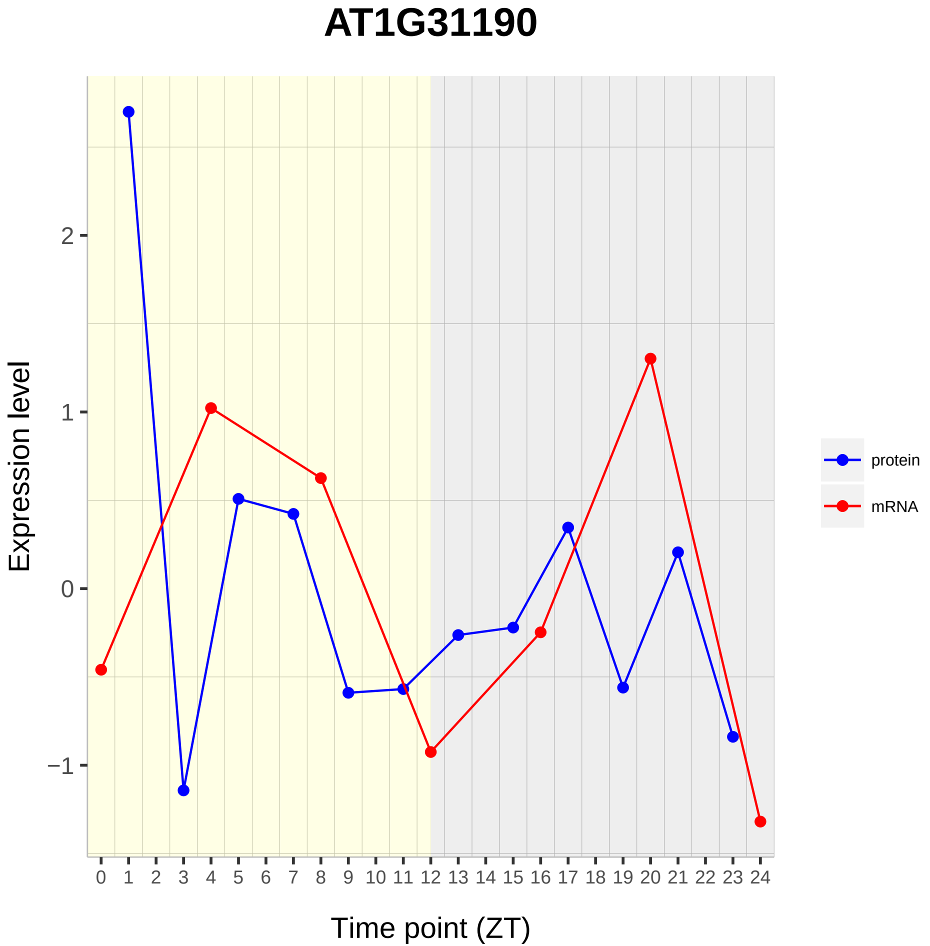
AT1G31190
Expression level
2
1
0
−1
0 1 2 3 4 5 6 7 8 9 10 11 12 13 14 15 16 17 18 19 20 21 22 23 24
Time point (ZT)
protein
mRNA

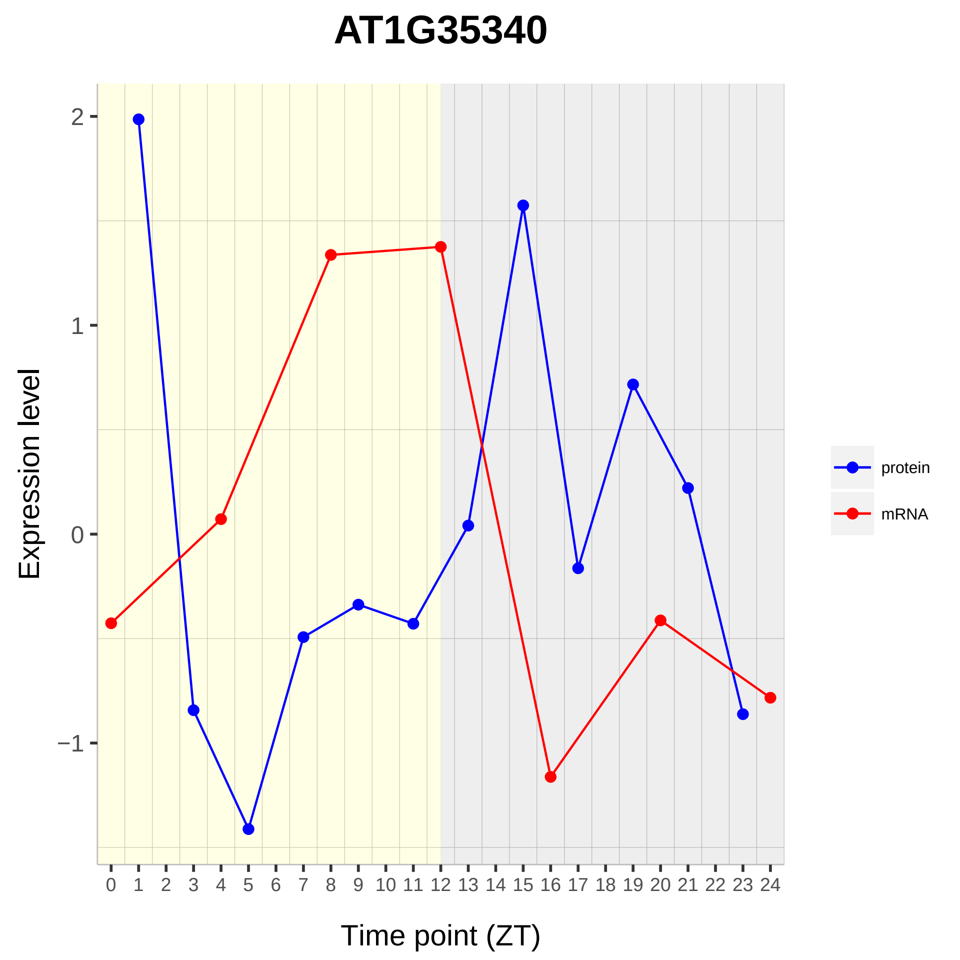

AT1G35340
Expression level
2
1
0
−1
0 1 2 3 4 5 6 7 8 9 10 11 12 13 14 15 16 17 18 19 20 21 22 23 24
Time point (ZT)
protein
mRNA

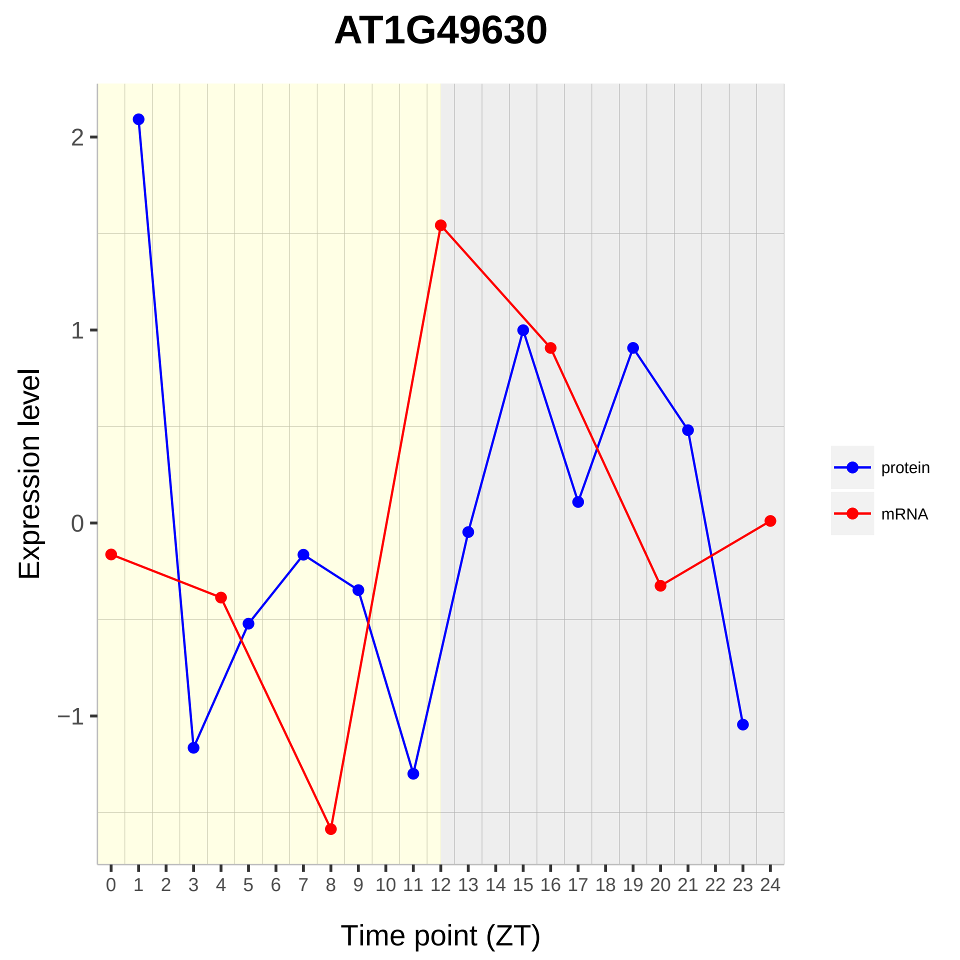

AT1G49630
Expression level
2
1
0
−1
Time point (ZT)
protein
mRNA

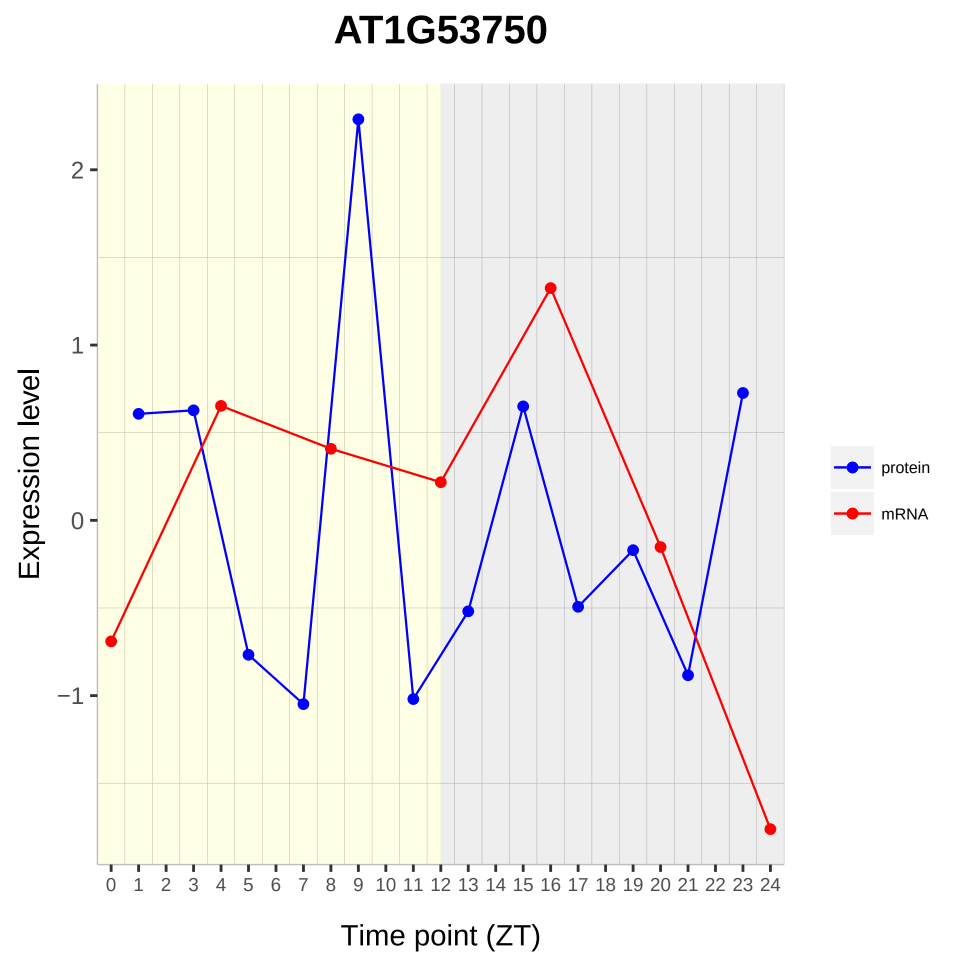
AT1G53750
Expression level
Time point (ZT)
2
1
0
−1
0 1 2 3 4 5 6 7 8 9 10 11 12 13 14 15 16 17 18 19 20 21 22 23 24
protein
mRNA

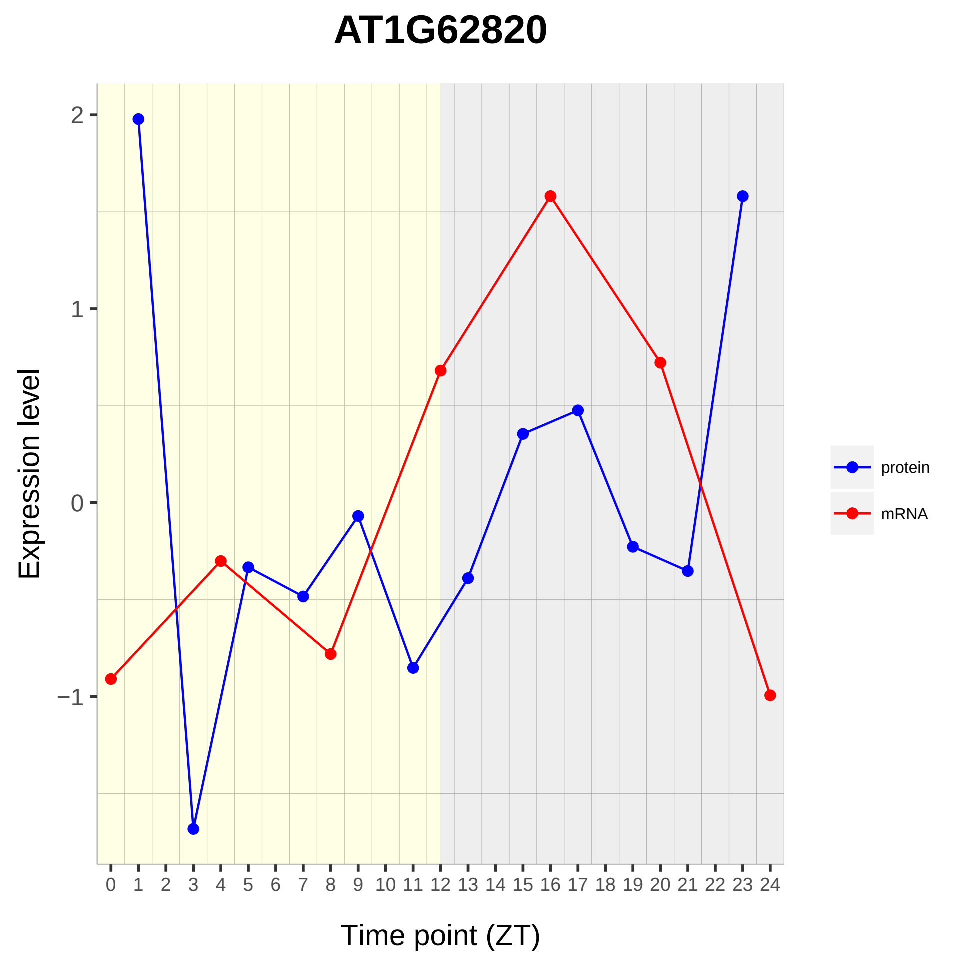

AT1G62820
Expression level
Time point (ZT)
2
1
0
−1
0 1 2 3 4 5 6 7 8 9 10 11 12 13 14 15 16 17 18 19 20 21 22 23 24
protein
mRNA

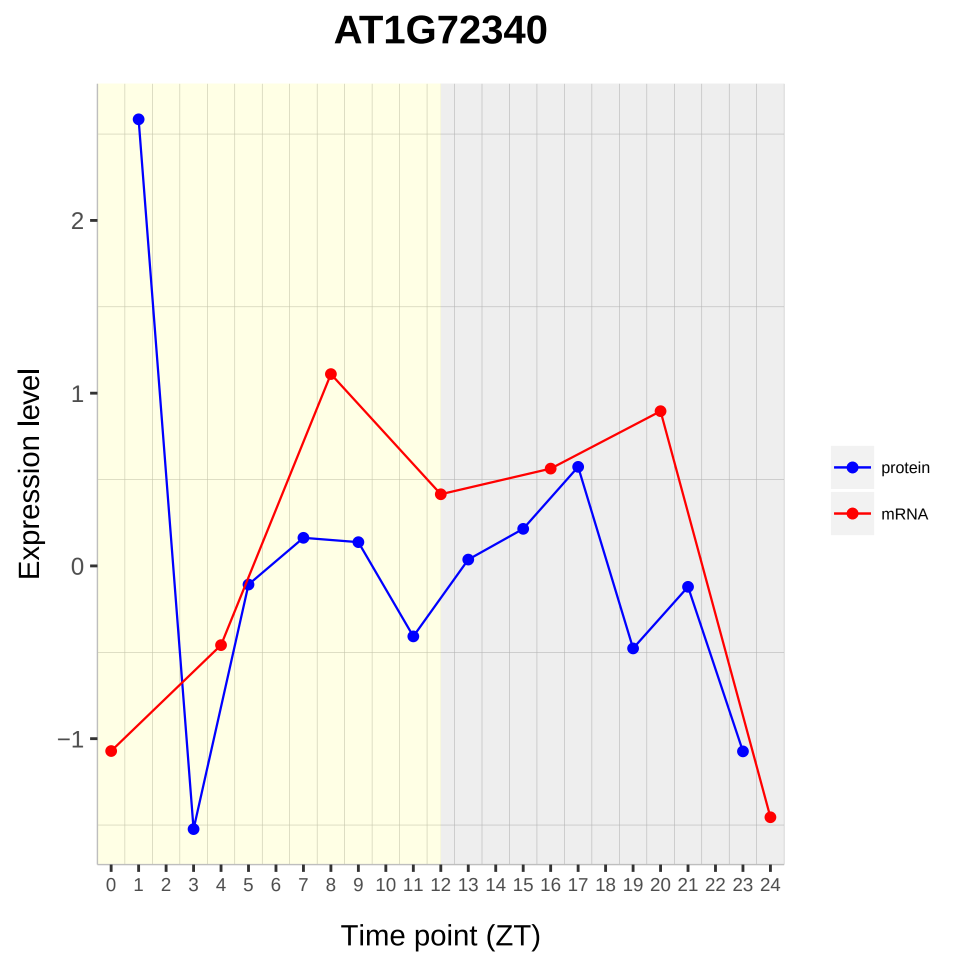

AT1G72340
Expression level
Time point (ZT)
protein
mRNA
2
1
0
−1
0 1 2 3 4 5 6 7 8 9 10 11 12 13 14 15 16 17 18 19 20 21 22 23 24

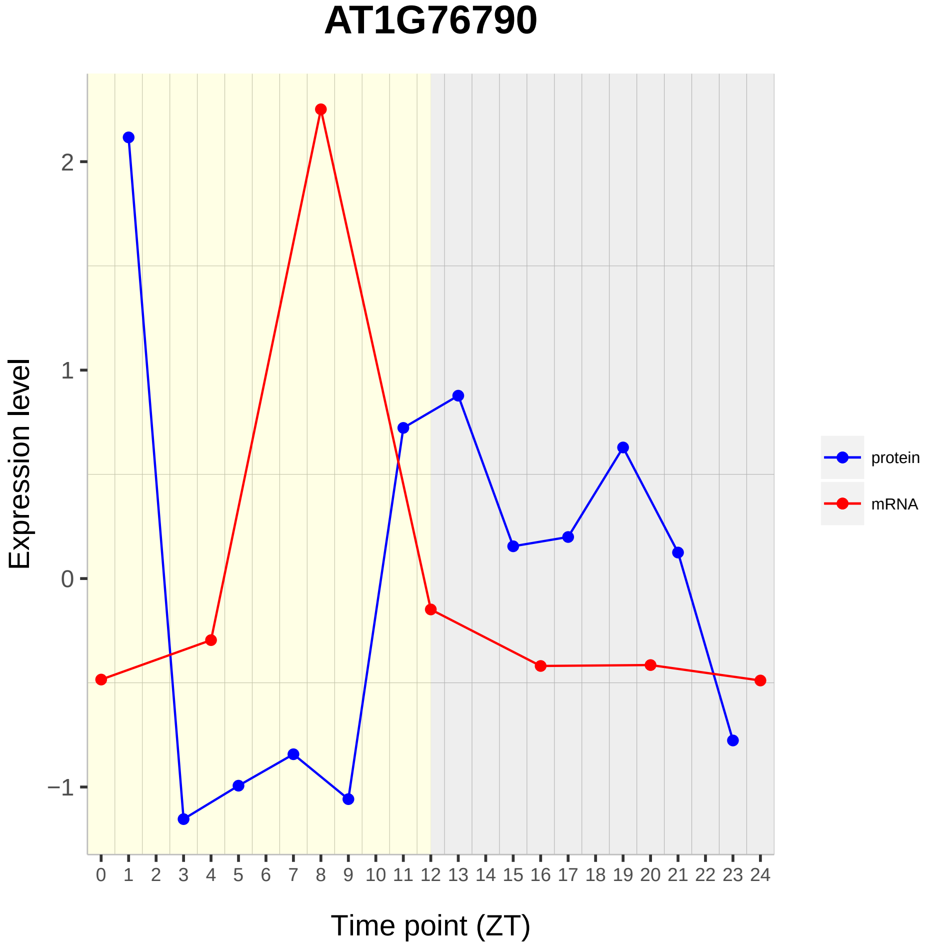

AT1G76790
Expression level
2
1
0
−1
Time point (ZT)
protein
mRNA

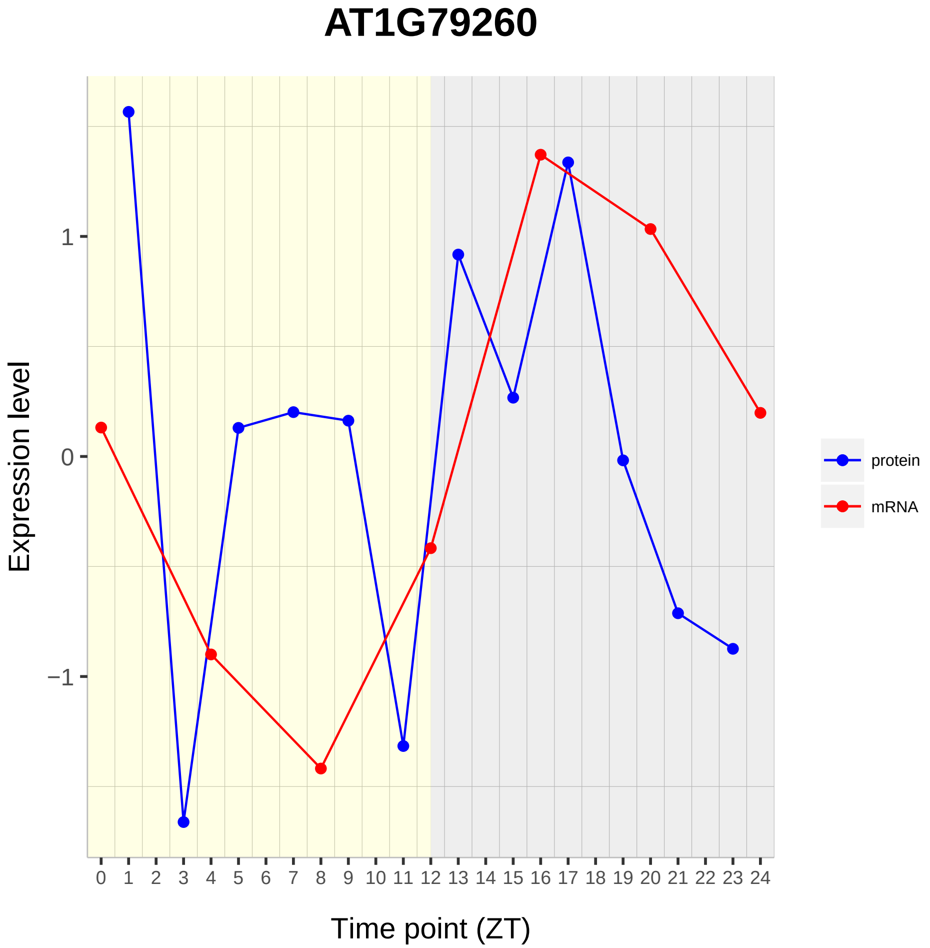
AT1G79260
Expression level
1
0
−1
0 1 2 3 4 5 6 7 8 9 10 11 12 13 14 15 16 17 18 19 20 21 22 23 24
Time point (ZT)
protein
mRNA

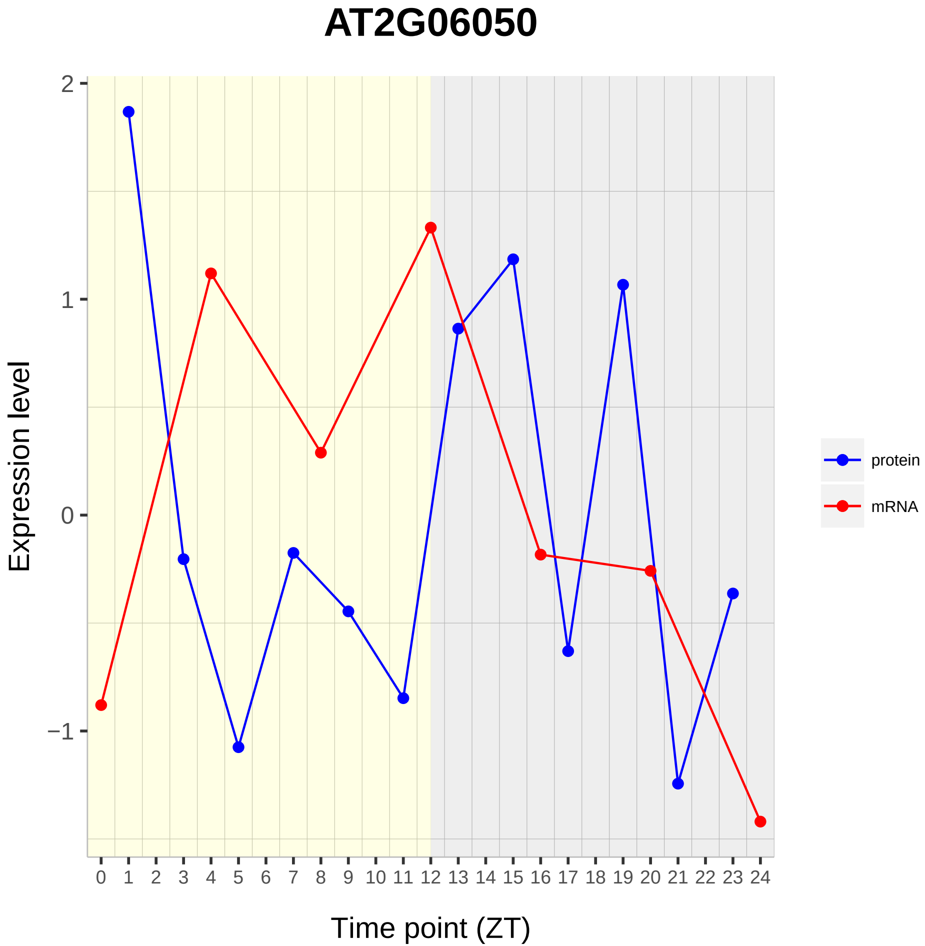
AT2G06050
2
1
0
−1
Expression level
0 1 2 3 4 5 6 7 8 9 10 11 12 13 14 15 16 17 18 19 20 21 22 23 24
Time point (ZT)
protein
mRNA

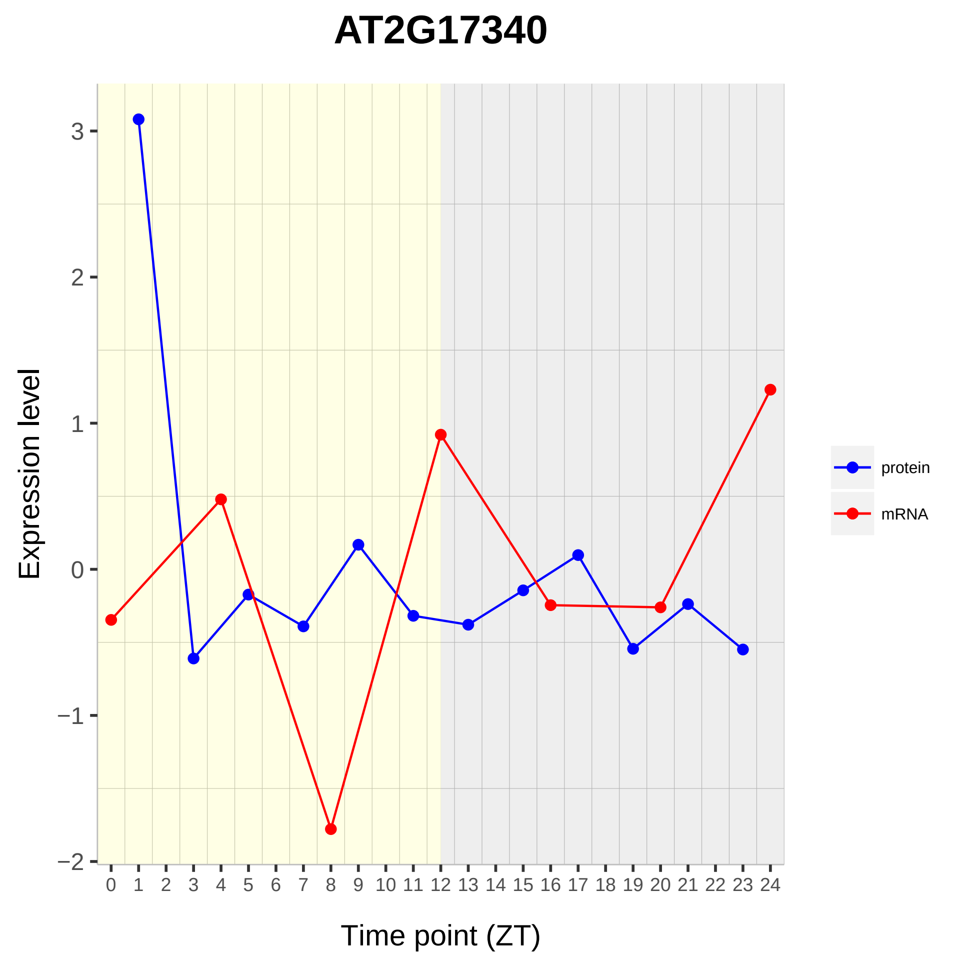

AT2G17340
Expression level
3
2
1
0
−1
−2
Time point (ZT)
protein
mRNA

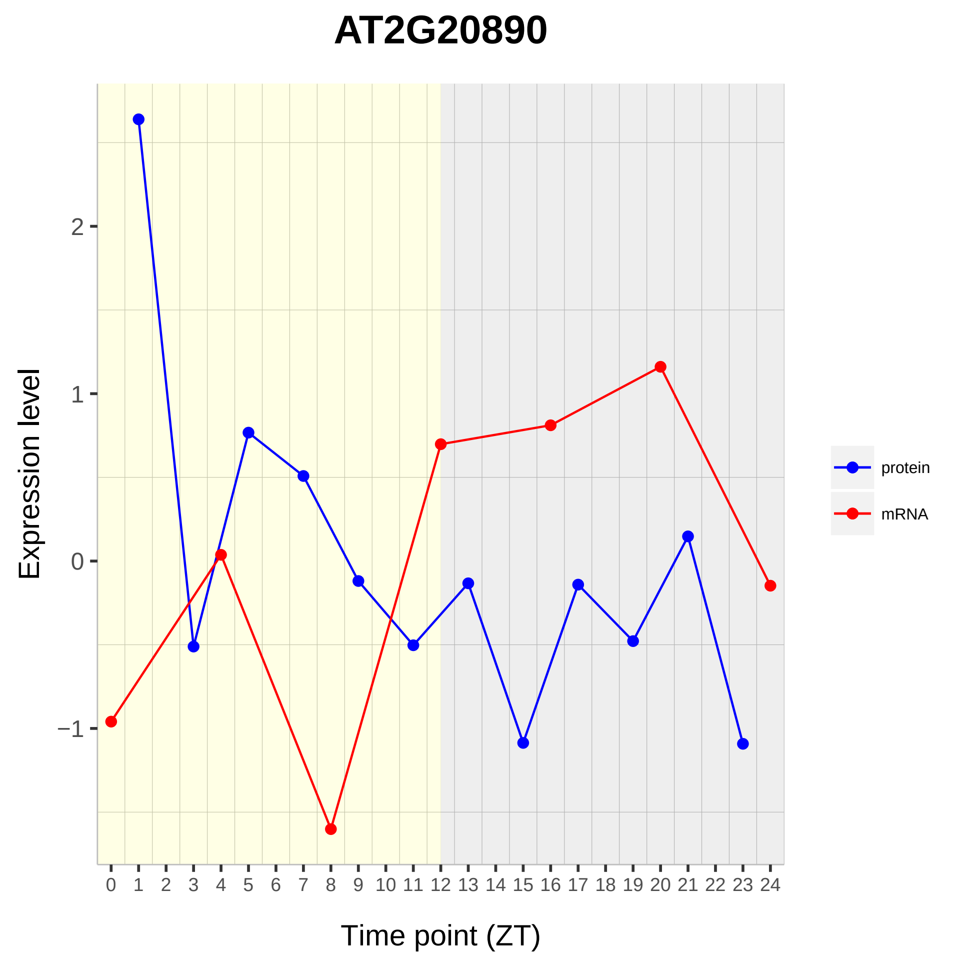

AT2G20890
Expression level
2
1
0
−1
0 1 2 3 4 5 6 7 8 9 10 11 12 13 14 15 16 17 18 19 20 21 22 23 24
Time point (ZT)
protein
mRNA

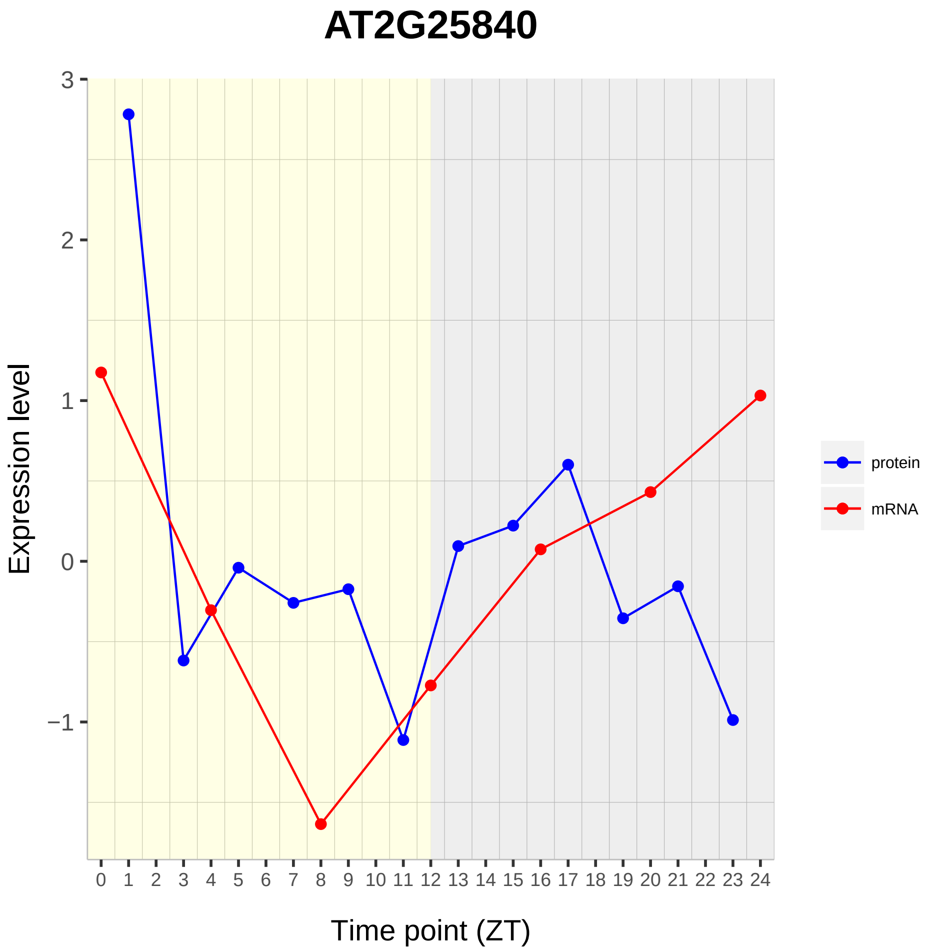
AT2G25840
Expression level
3
2
1
0
−1
0 1 2 3 4 5 6 7 8 9 10 11 12 13 14 15 16 17 18 19 20 21 22 23 24
Time point (ZT)
protein
mRNA

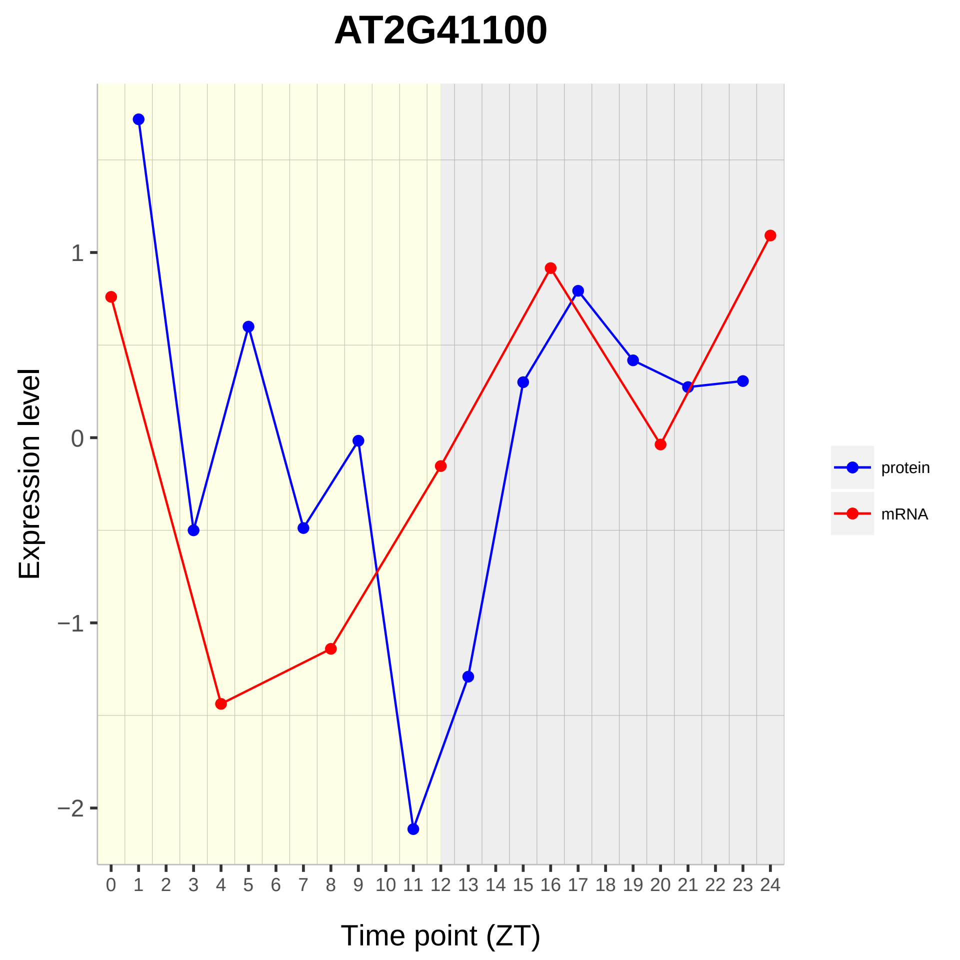
AT2G41100
Expression level
Time point (ZT)
1
0
−1
−2
0 1 2 3 4 5 6 7 8 9 10 11 12 13 14 15 16 17 18 19 20 21 22 23 24
protein
mRNA

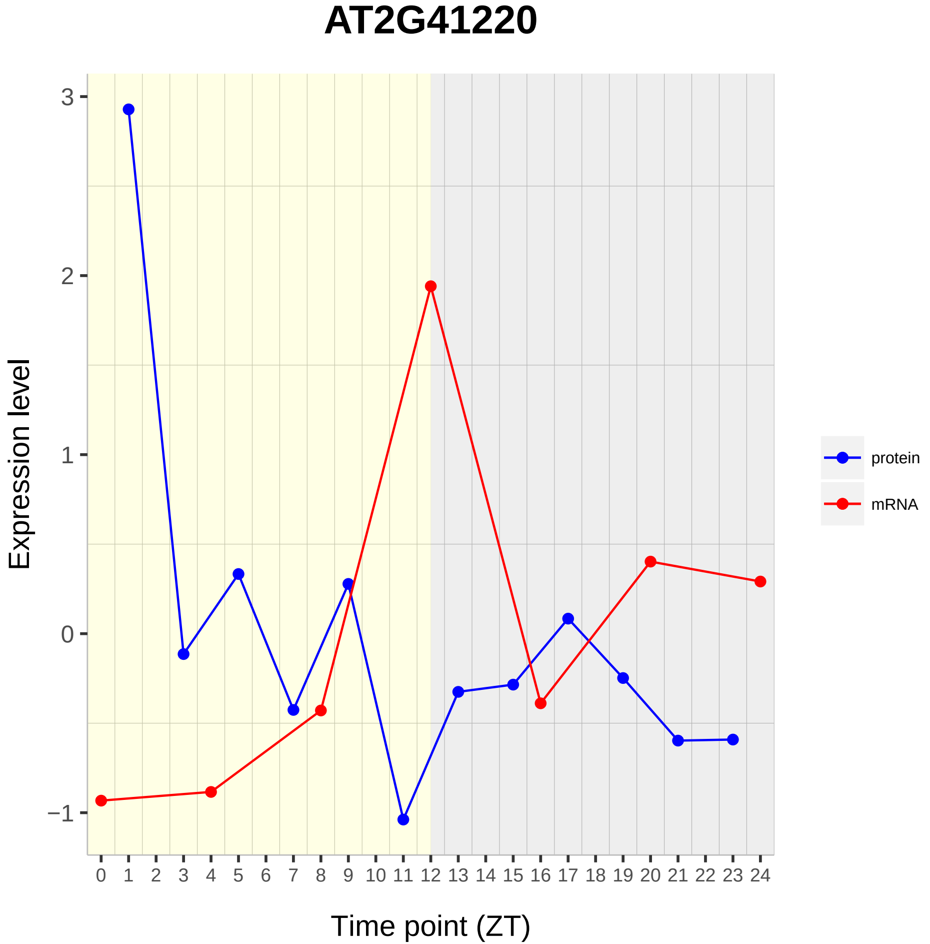
AT2G41220
Expression level
Time point (ZT)
protein
mRNA
3
2
1
0
−1
0 1 2 3 4 5 6 7 8 9 10 11 12 13 14 15 16 17 18 19 20 21 22 23 24

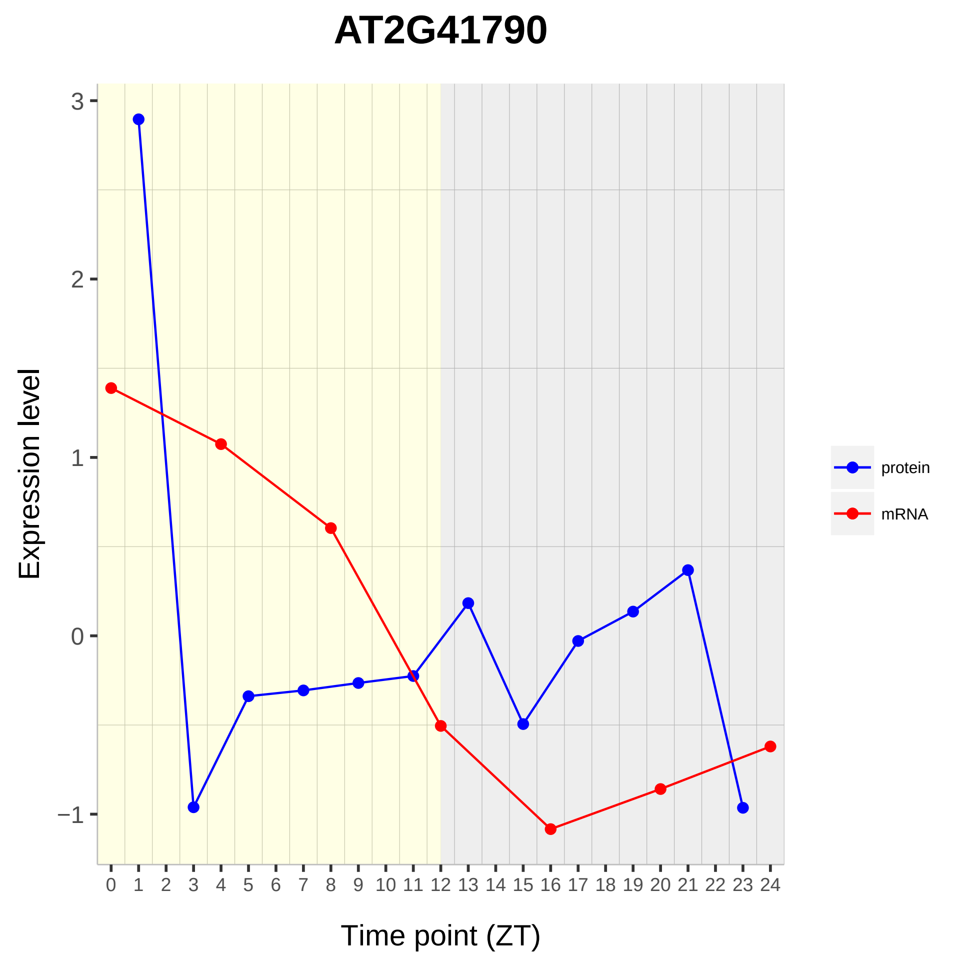

AT2G41790
Expression level
Time point (ZT)
3
2
1
0
−1
protein
mRNA

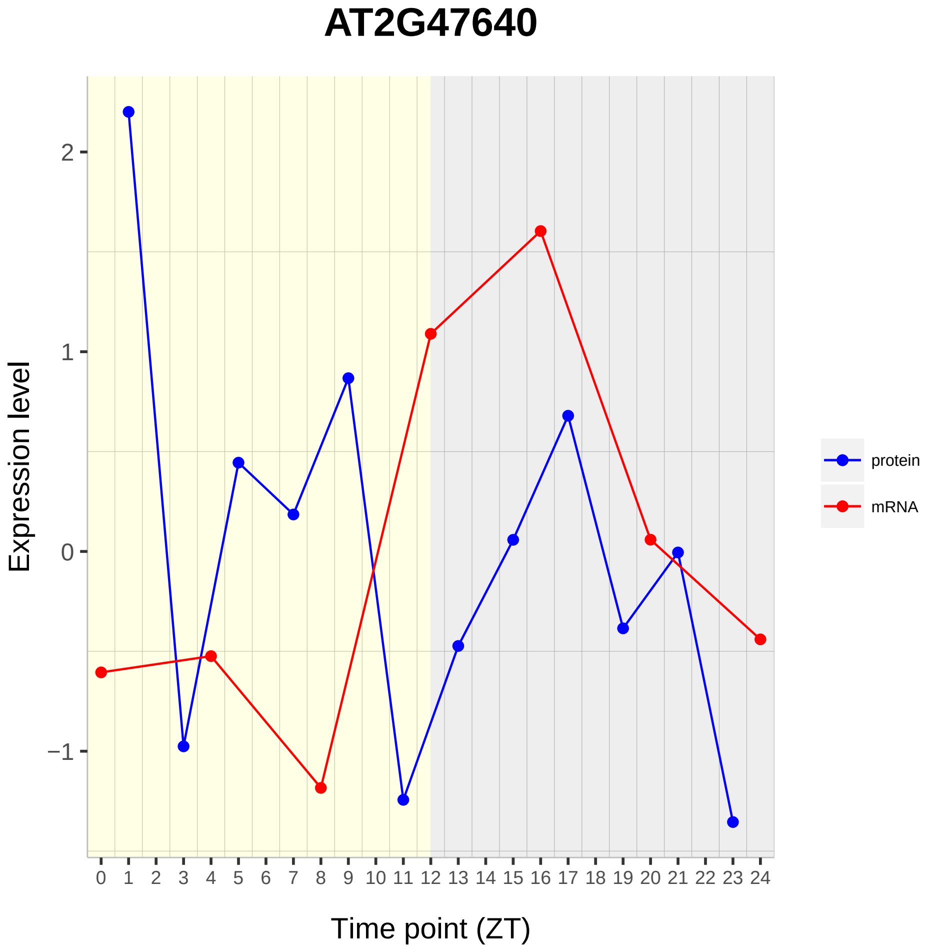

AT2G47640
Expression level
2
1
0
−1
Time point (ZT)
protein
mRNA

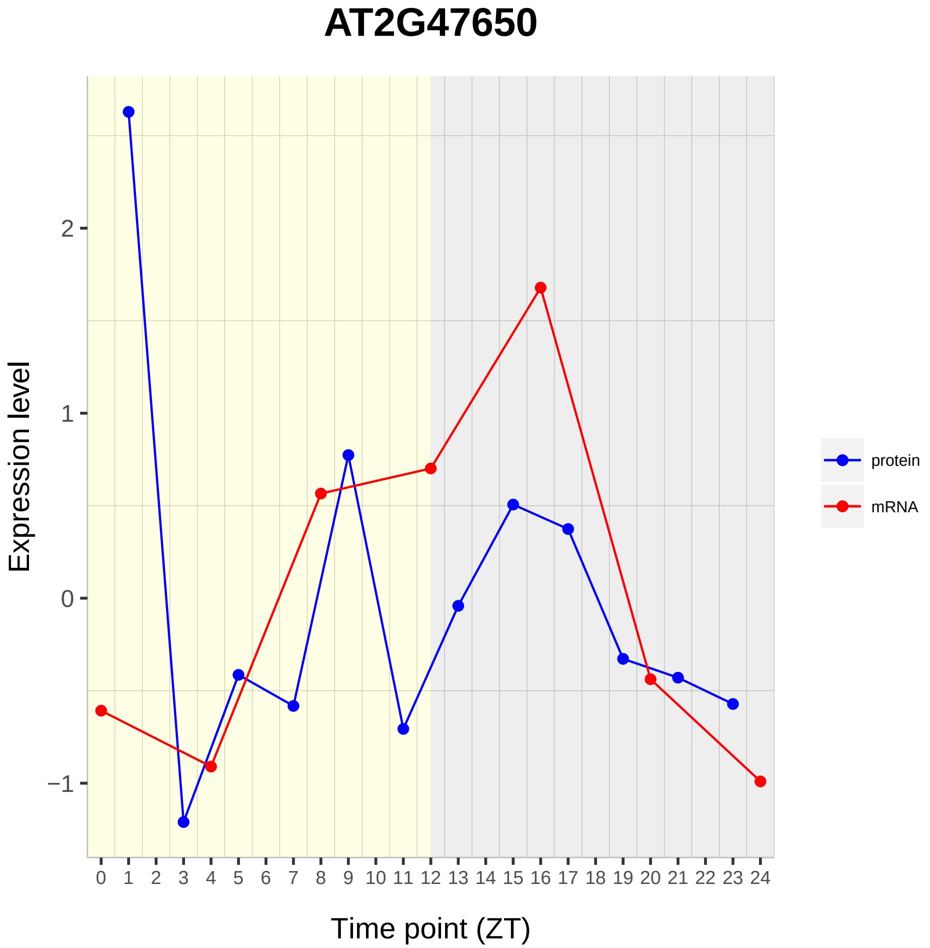
AT2G47650
Expression level
2
1
0
−1
0 1 2 3 4 5 6 7 8 9 10 11 12 13 14 15 16 17 18 19 20 21 22 23 24
Time point (ZT)
protein
mRNA

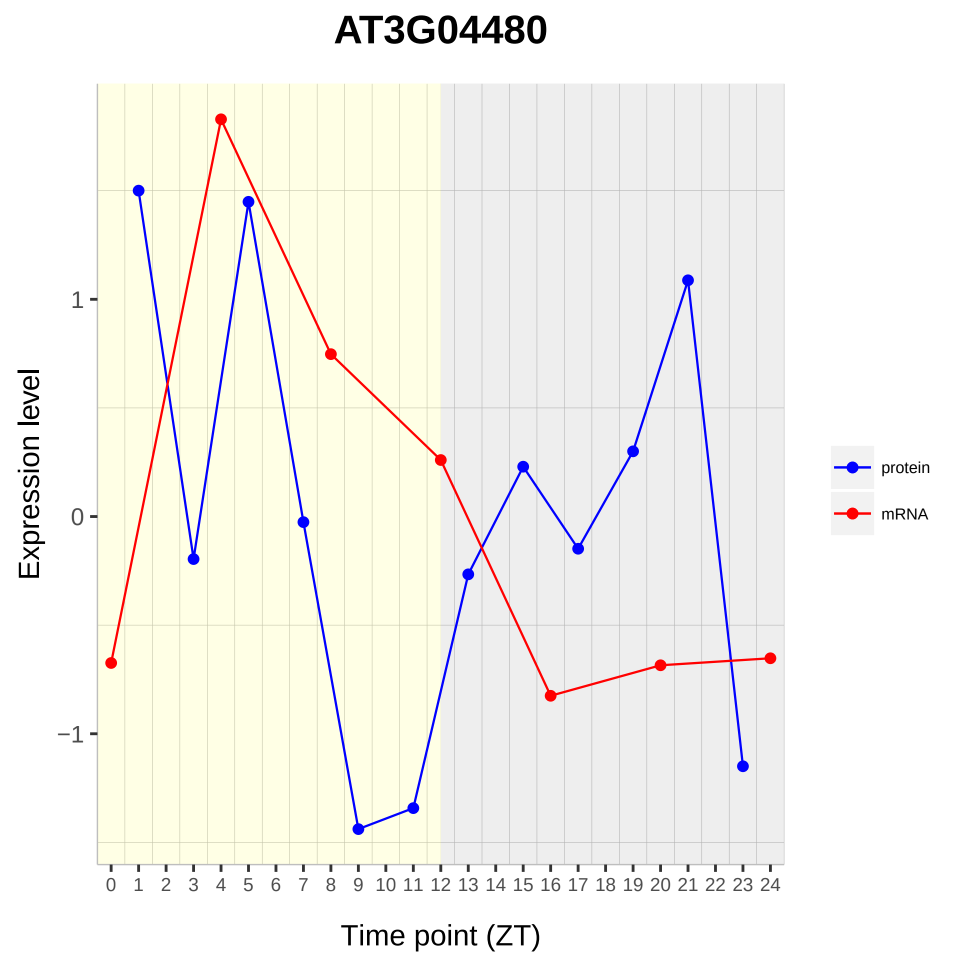

AT3G04480
Expression level
1
0
−1
Time point (ZT)
protein
mRNA

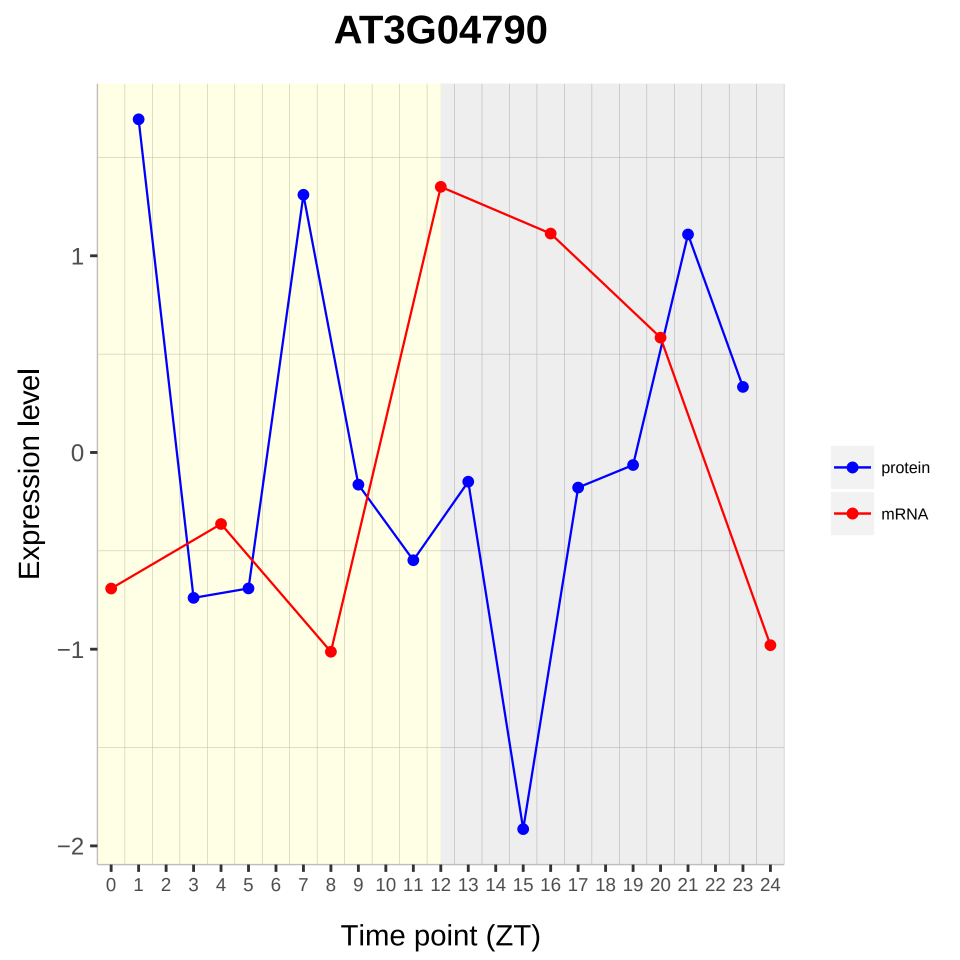

AT3G04790
Expression level
Time point (ZT)
1
0
−1
−2
0 1 2 3 4 5 6 7 8 9 10 11 12 13 14 15 16 17 18 19 20 21 22 23 24
protein
mRNA

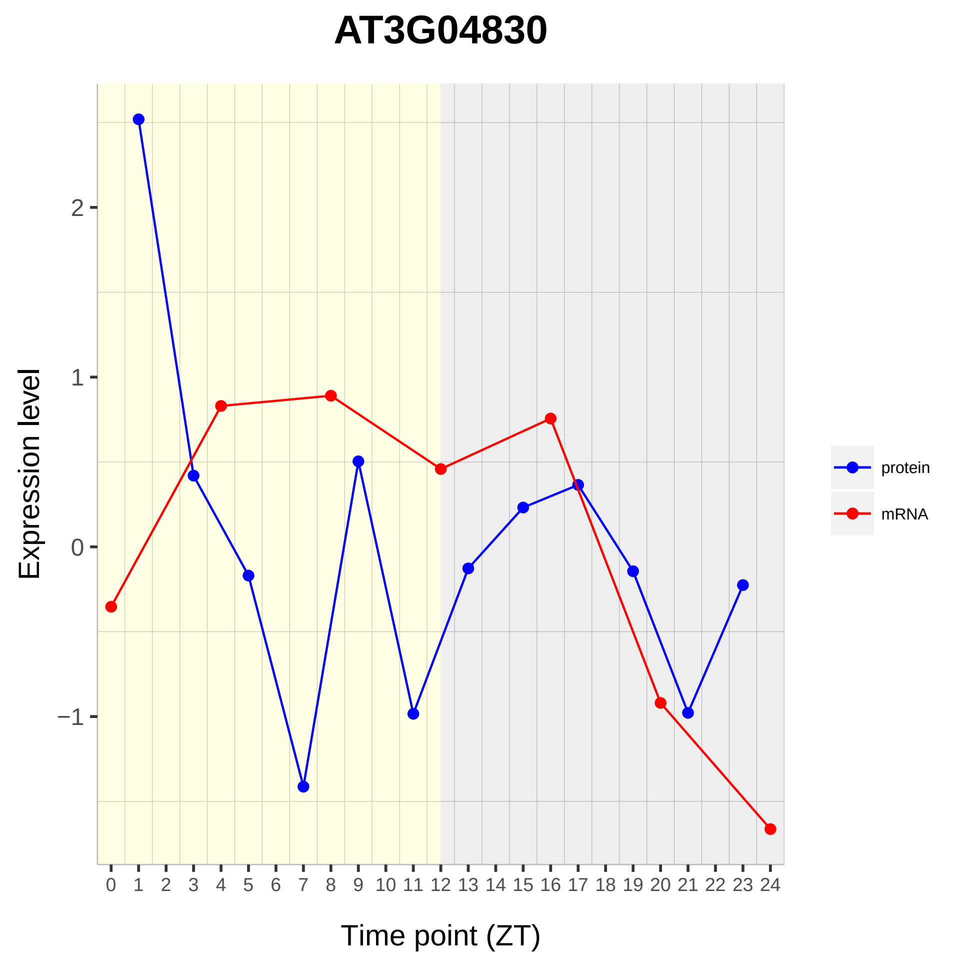

AT3G04830
Expression level
2
1
0
−1
Time point (ZT)
protein
mRNA

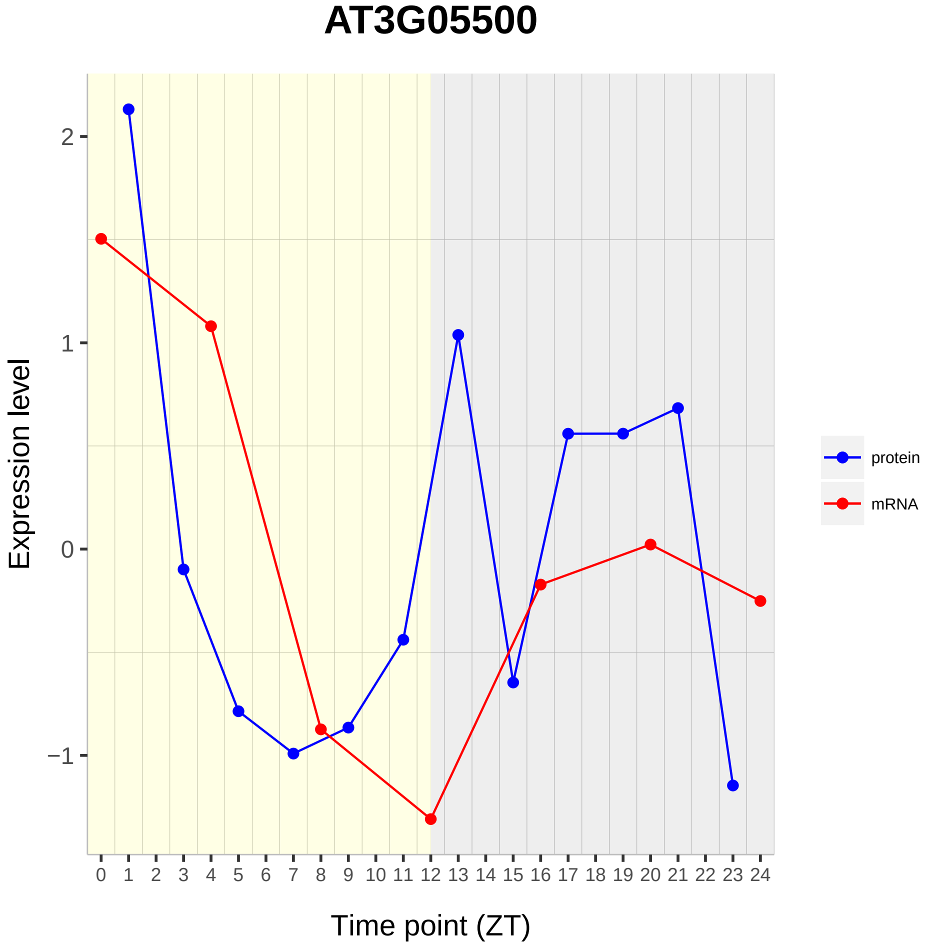

AT3G05500
Expression level
Time point (ZT)
2
1
0
−1
protein
mRNA

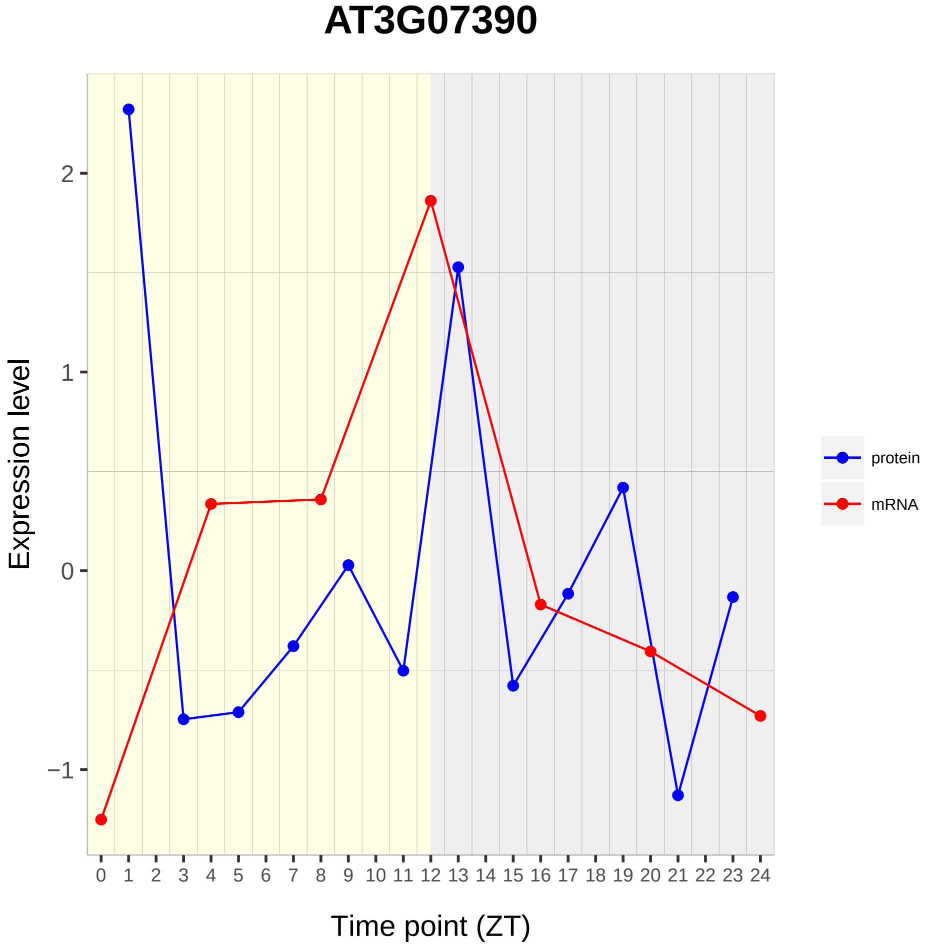
AT3G07390
Expression level
2
1
0
−1
0 1 2 3 4 5 6 7 8 9 10 11 12 13 14 15 16 17 18 19 20 21 22 23 24
Time point (ZT)
protein
mRNA

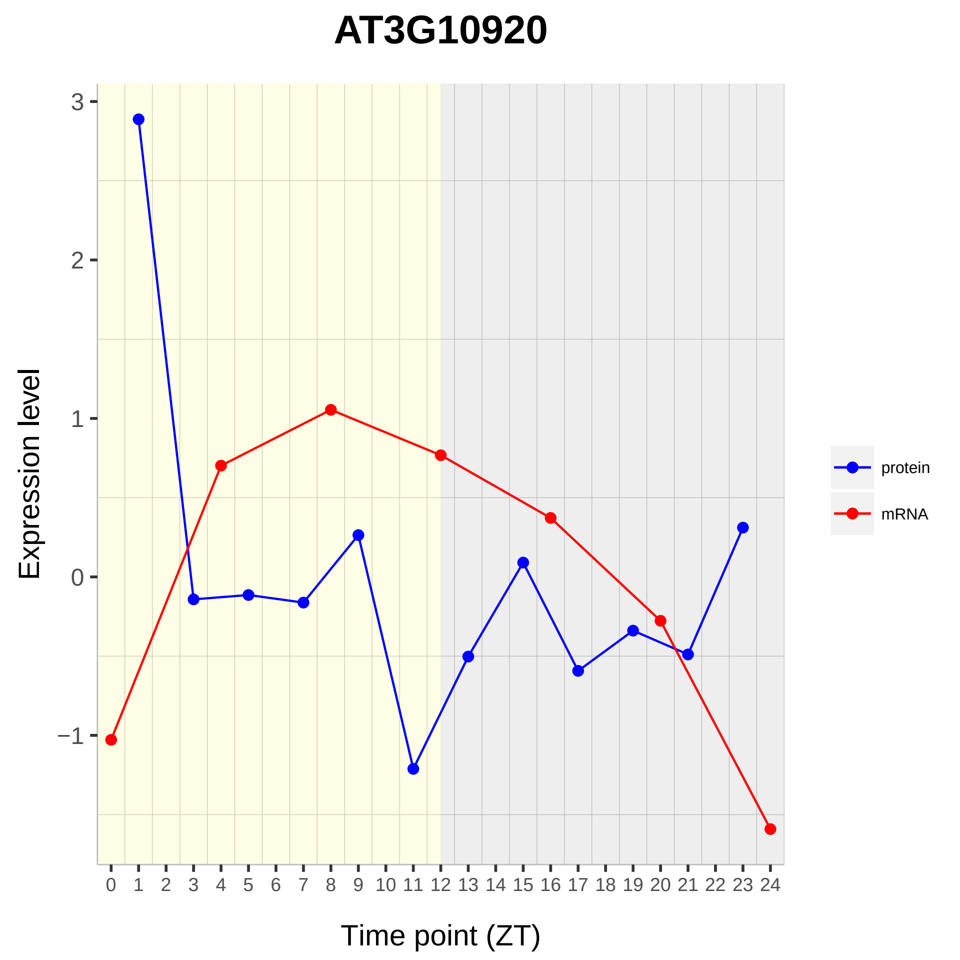
AT3G10920
Expression level
Time point (ZT)
3
2
1
0
−1
0 1 2 3 4 5 6 7 8 9 10 11 12 13 14 15 16 17 18 19 20 21 22 23 24
protein
mRNA

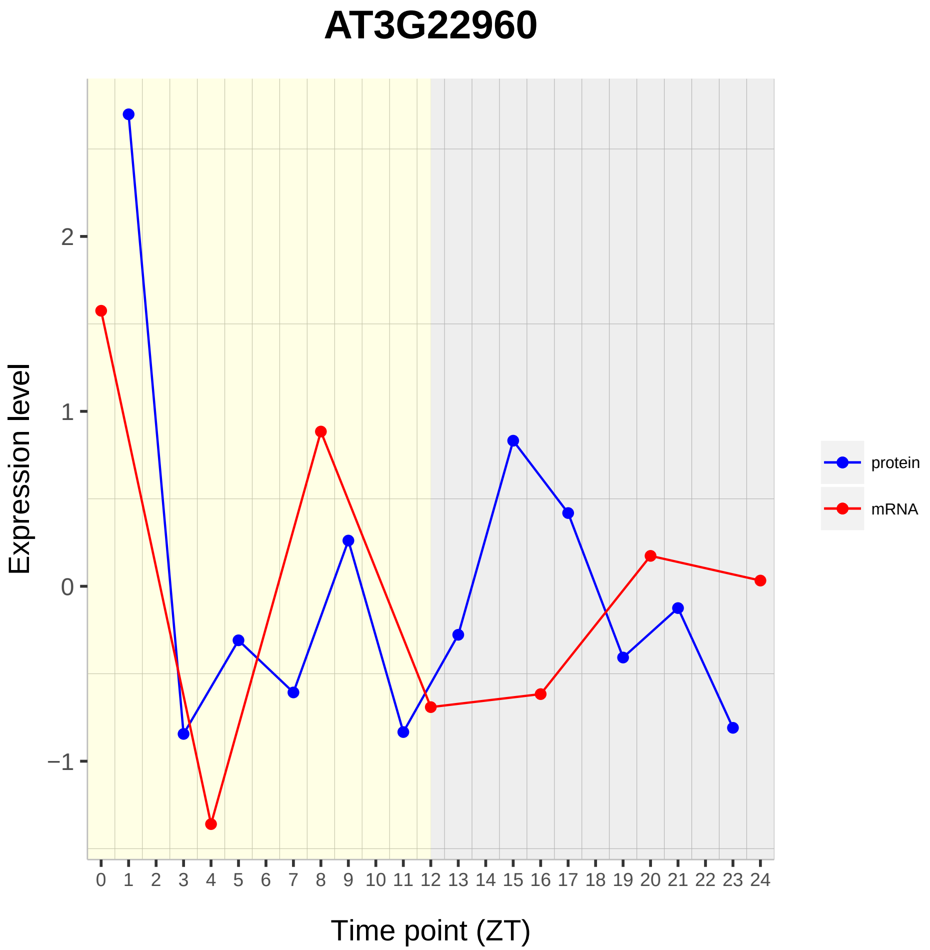

AT3G22960
Expression level
2
1
0
−1
Time point (ZT)
protein
mRNA

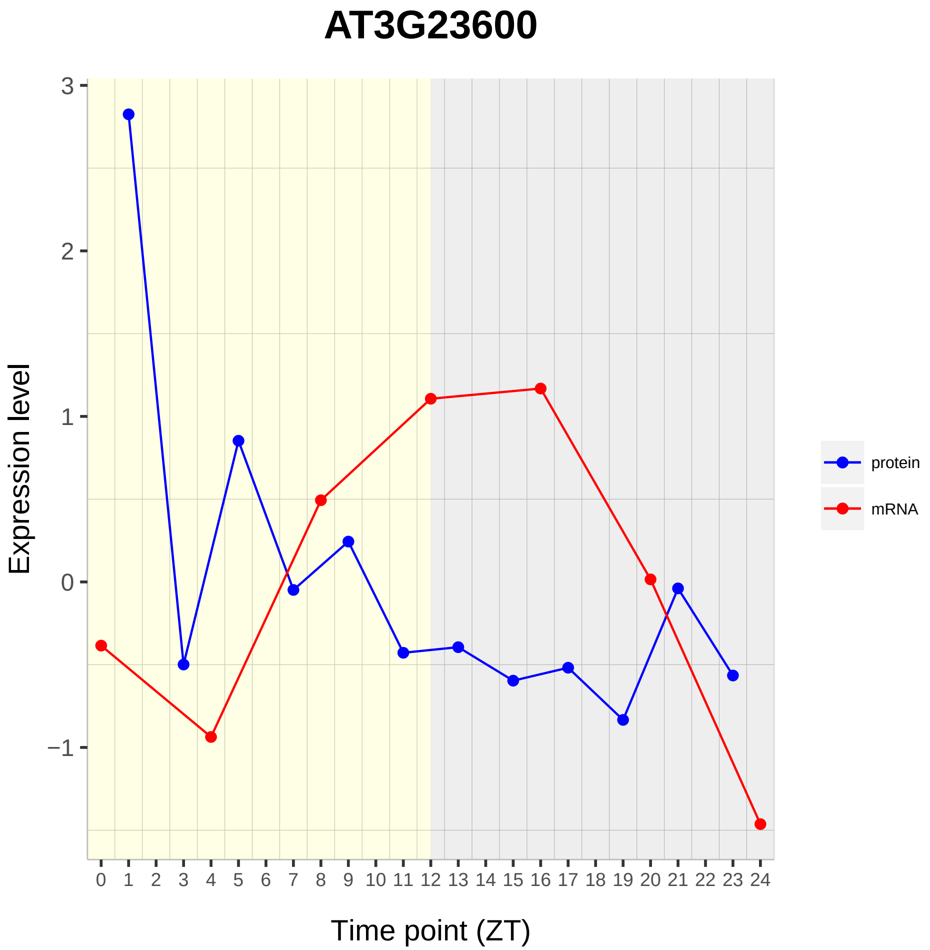

AT3G23600
Expression level
Time point (ZT)
3
2
1
0
−1
0 1 2 3 4 5 6 7 8 9 10 11 12 13 14 15 16 17 18 19 20 21 22 23 24
protein
mRNA

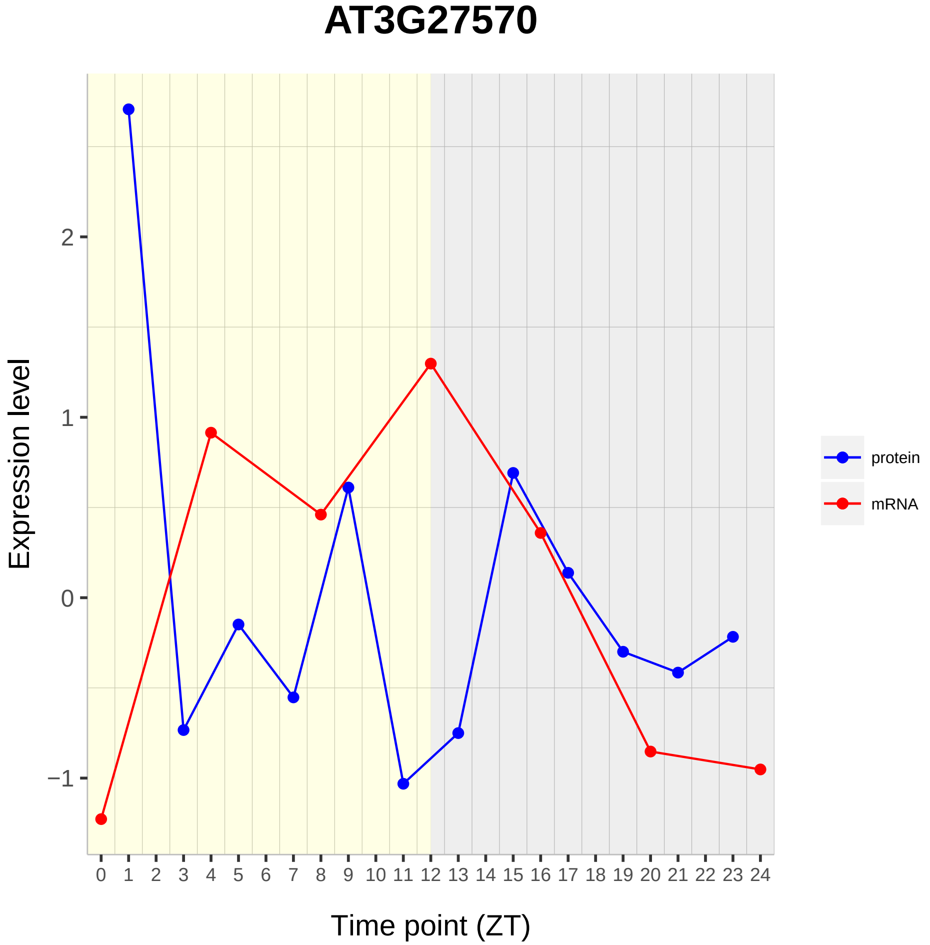

AT3G27570
Expression level
Time point (ZT)
2
1
0
−1
0 1 2 3 4 5 6 7 8 9 10 11 12 13 14 15 16 17 18 19 20 21 22 23 24
protein
mRNA

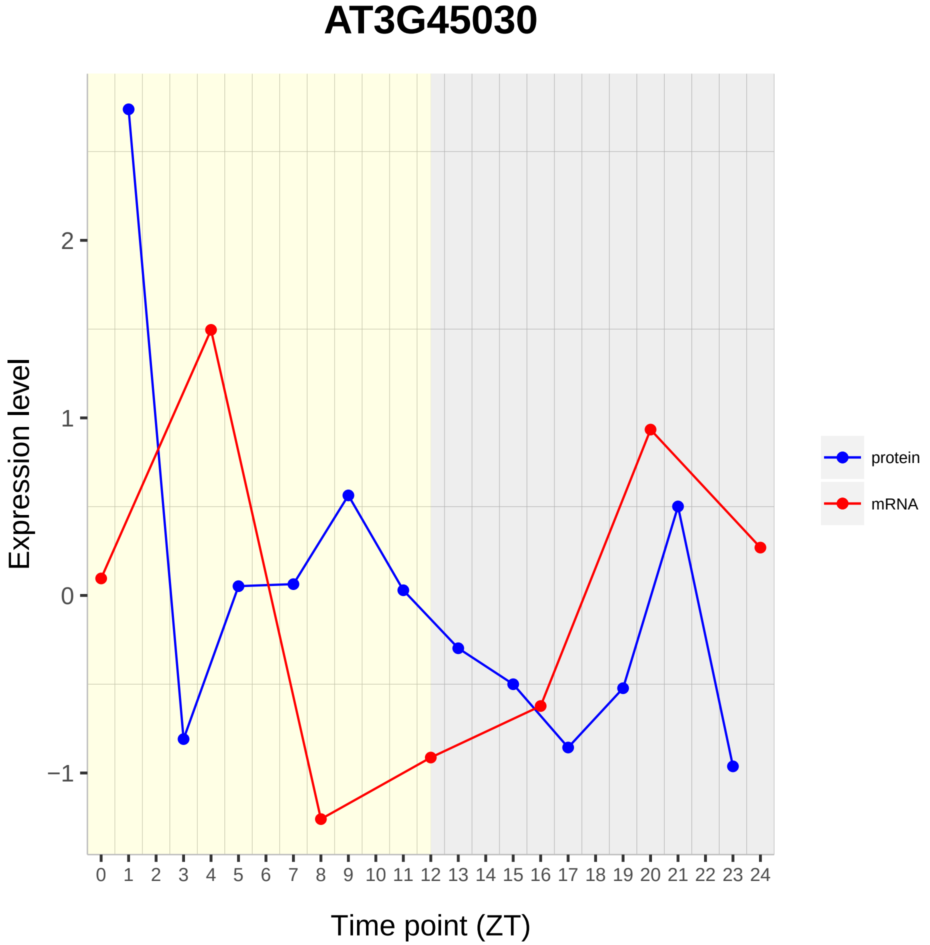

AT3G45030
Expression level
2
1
0
−1
Time point (ZT)
protein
mRNA

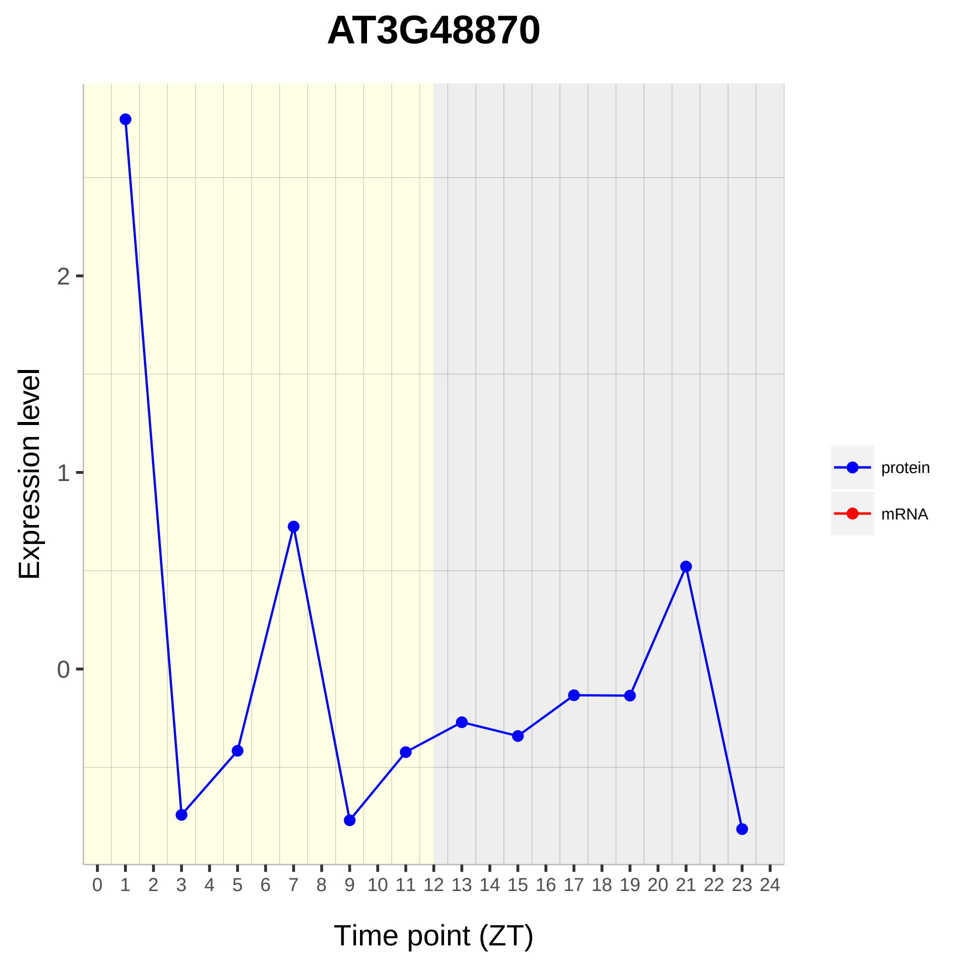
AT3G48870
Expression level
2
1
0
0 1 2 3 4 5 6 7 8 9 10 11 12 13 14 15 16 17 18 19 20 21 22 23 24
Time point (ZT)
protein
mRNA

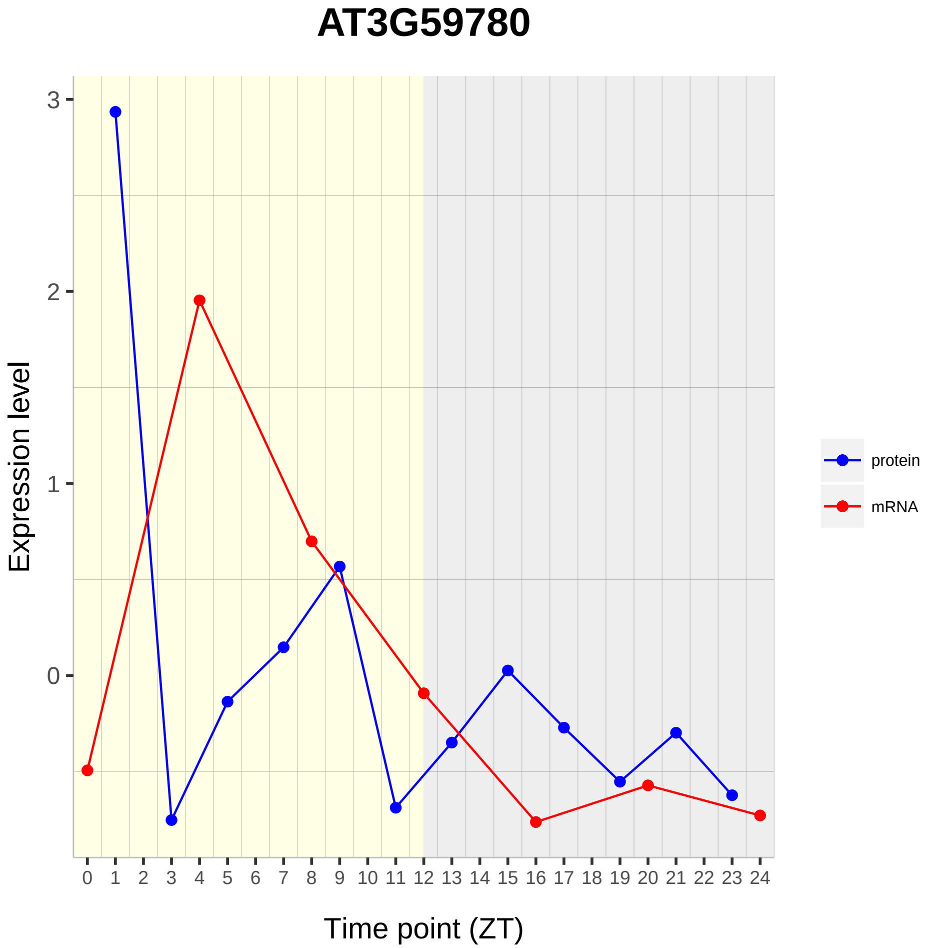

AT3G59780
Expression level
3
2
1
0
Time point (ZT)
protein
mRNA

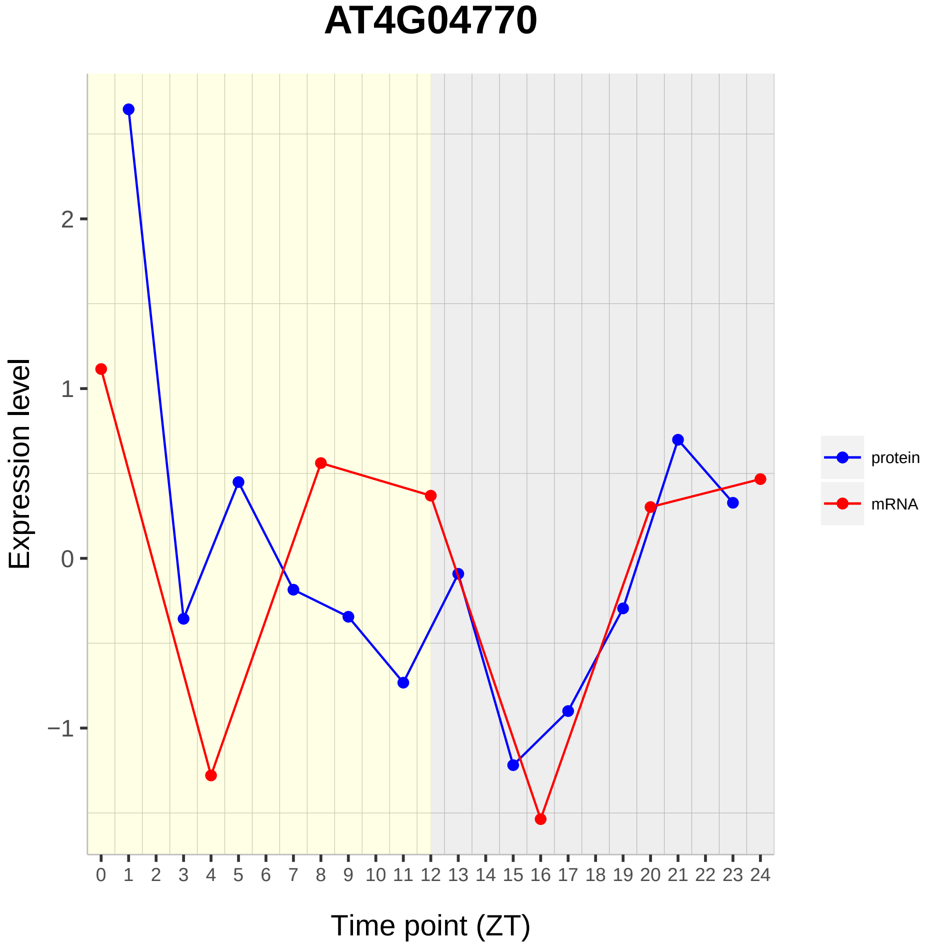

AT4G04770
Expression level
2
1
0
−1
0 1 2 3 4 5 6 7 8 9 10 11 12 13 14 15 16 17 18 19 20 21 22 23 24
Time point (ZT)
protein
mRNA

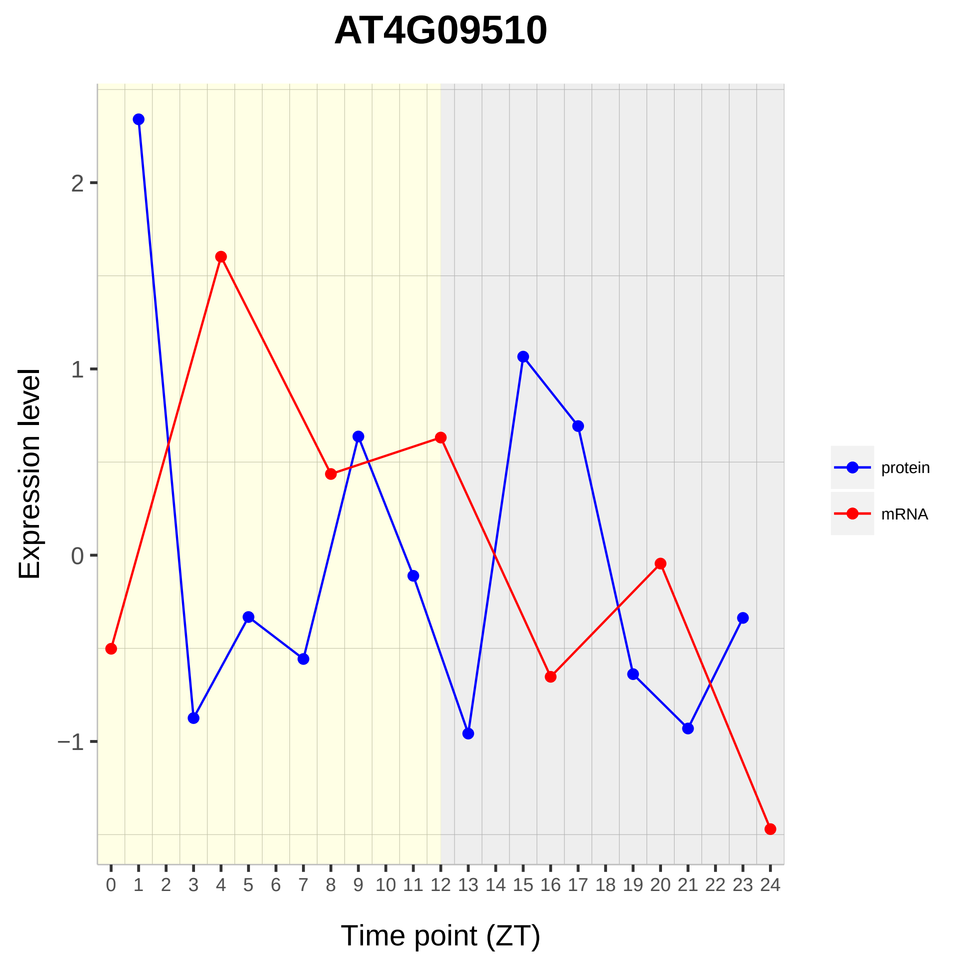
AT4G09510
Expression level
Time point (ZT)
2
1
0
−1
protein
mRNA

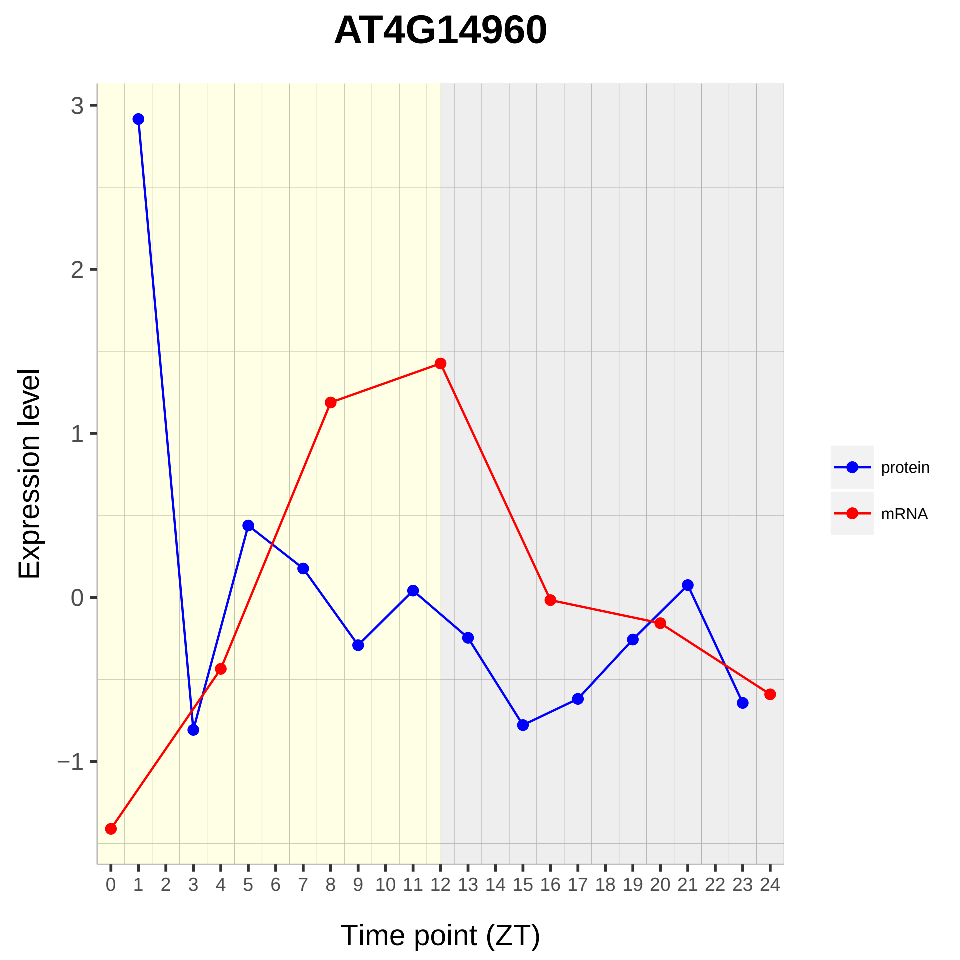

AT4G14960
Expression level
3
2
1
0
−1
Time point (ZT)
0 1 2 3 4 5 6 7 8 9 10 11 12 13 14 15 16 17 18 19 20 21 22 23 24
protein
mRNA

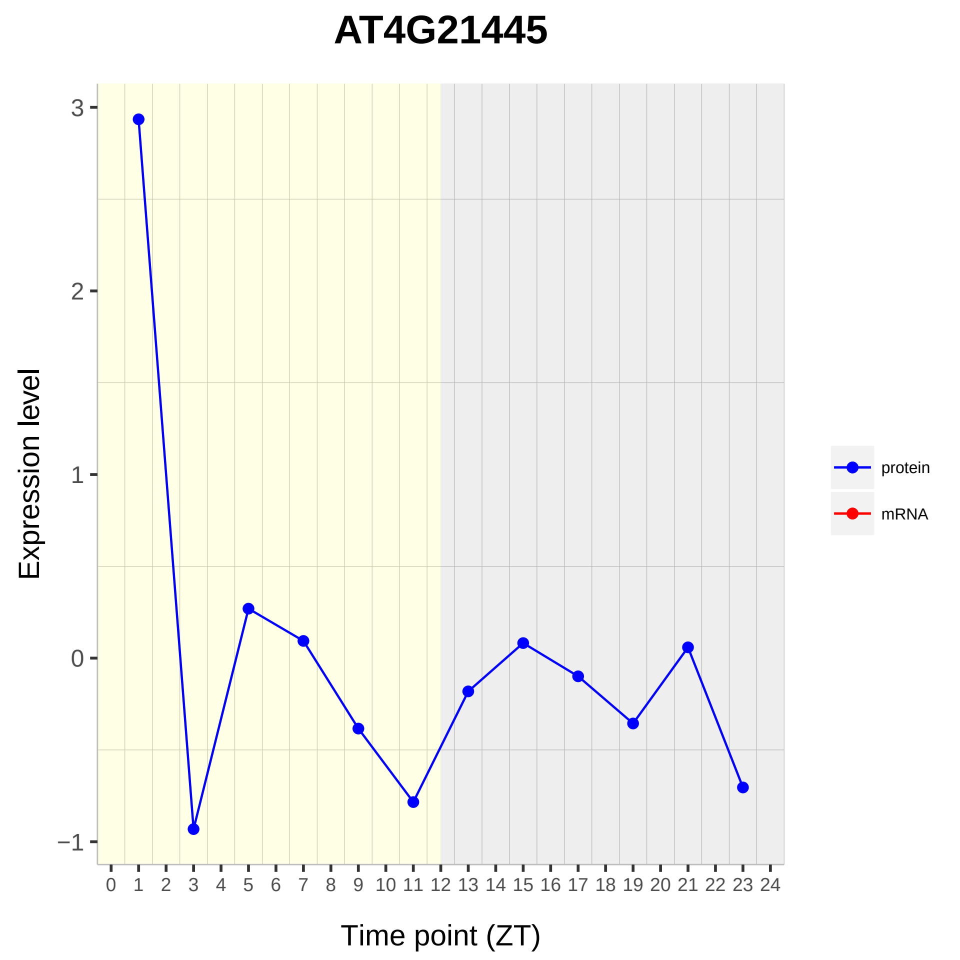
AT4G21445
Expression level
3
2
1
0
−1
0 1 2 3 4 5 6 7 8 9 10 11 12 13 14 15 16 17 18 19 20 21 22 23 24
Time point (ZT)
protein
mRNA

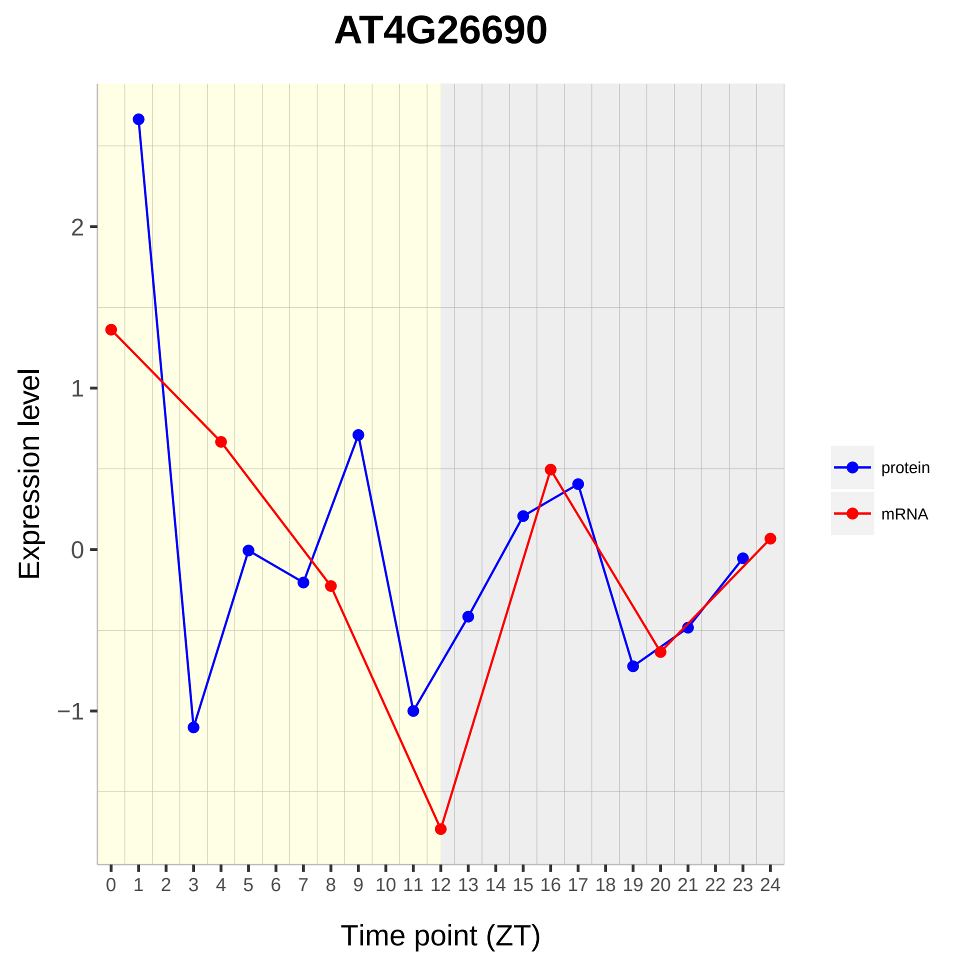

AT4G26690
Expression level
Time point (ZT)
2
1
0
−1
0 1 2 3 4 5 6 7 8 9 10 11 12 13 14 15 16 17 18 19 20 21 22 23 24
protein
mRNA

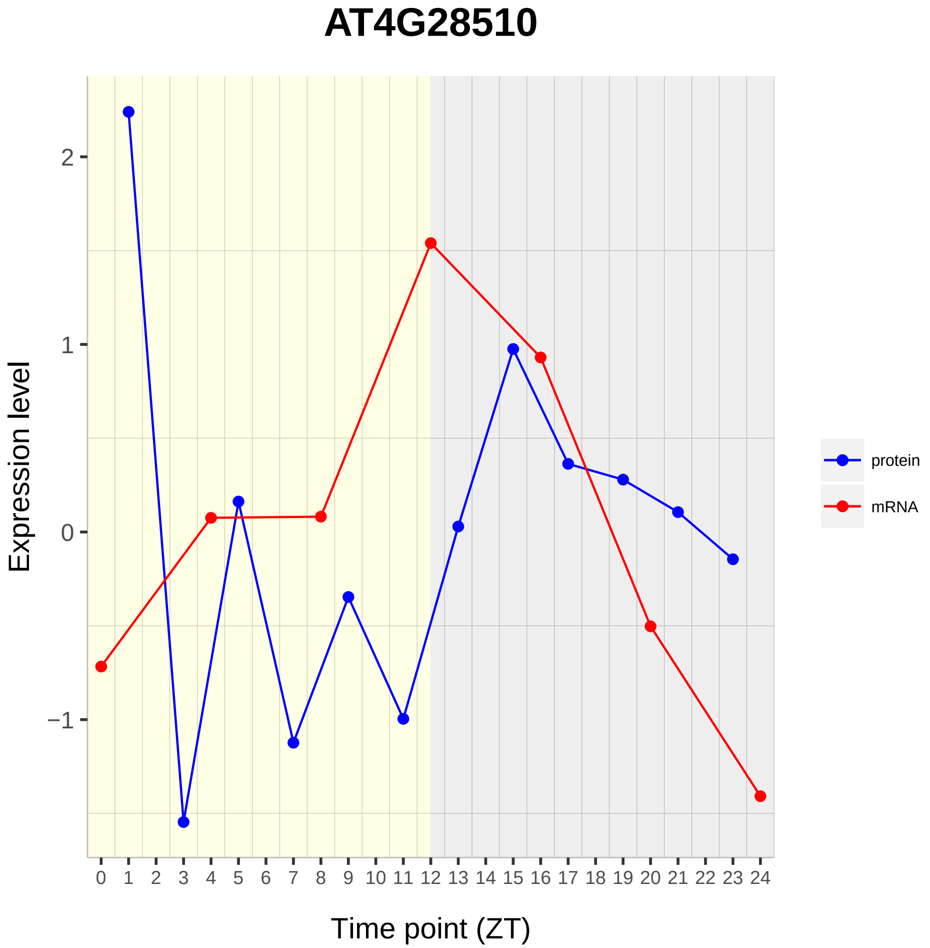

AT4G28510
Expression level
Time point (ZT)
2
1
0
−1
protein
mRNA

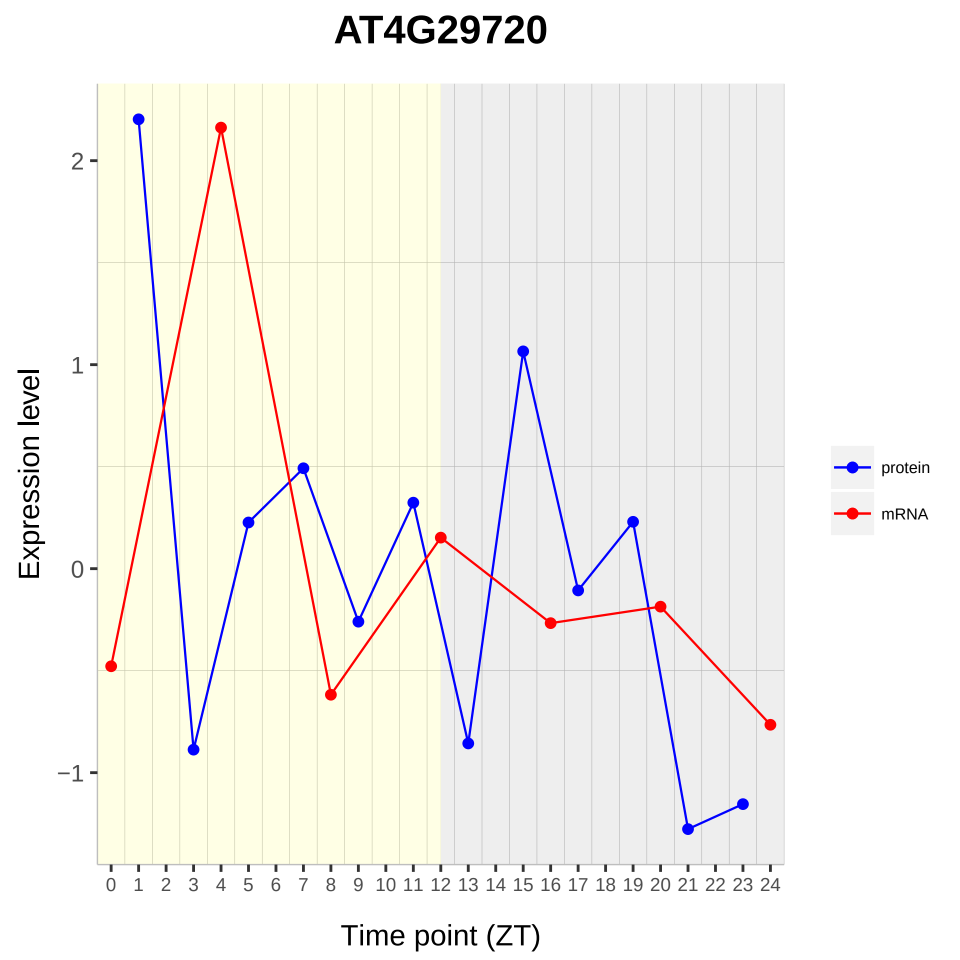

AT4G29720
Expression level
2
1
0
−1
Time point (ZT)
protein
mRNA

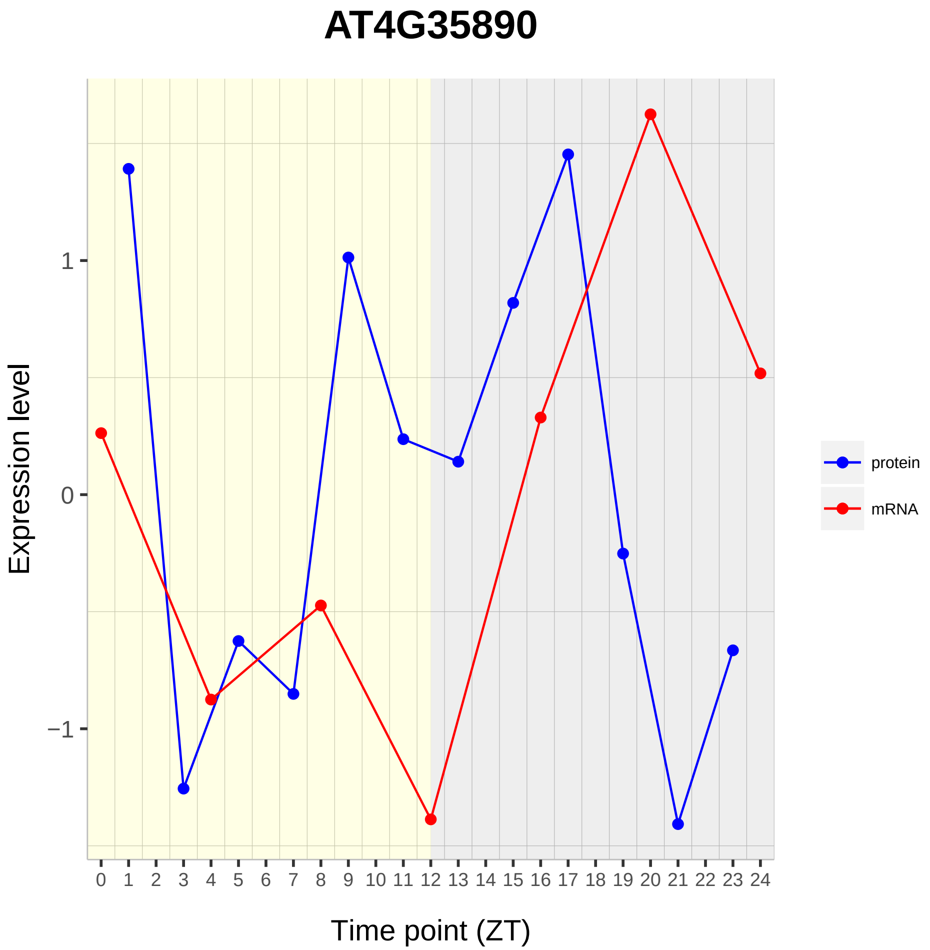
AT4G35890
Expression level
1
0
−1
0 1 2 3 4 5 6 7 8 9 10 11 12 13 14 15 16 17 18 19 20 21 22 23 24
Time point (ZT)
protein
mRNA

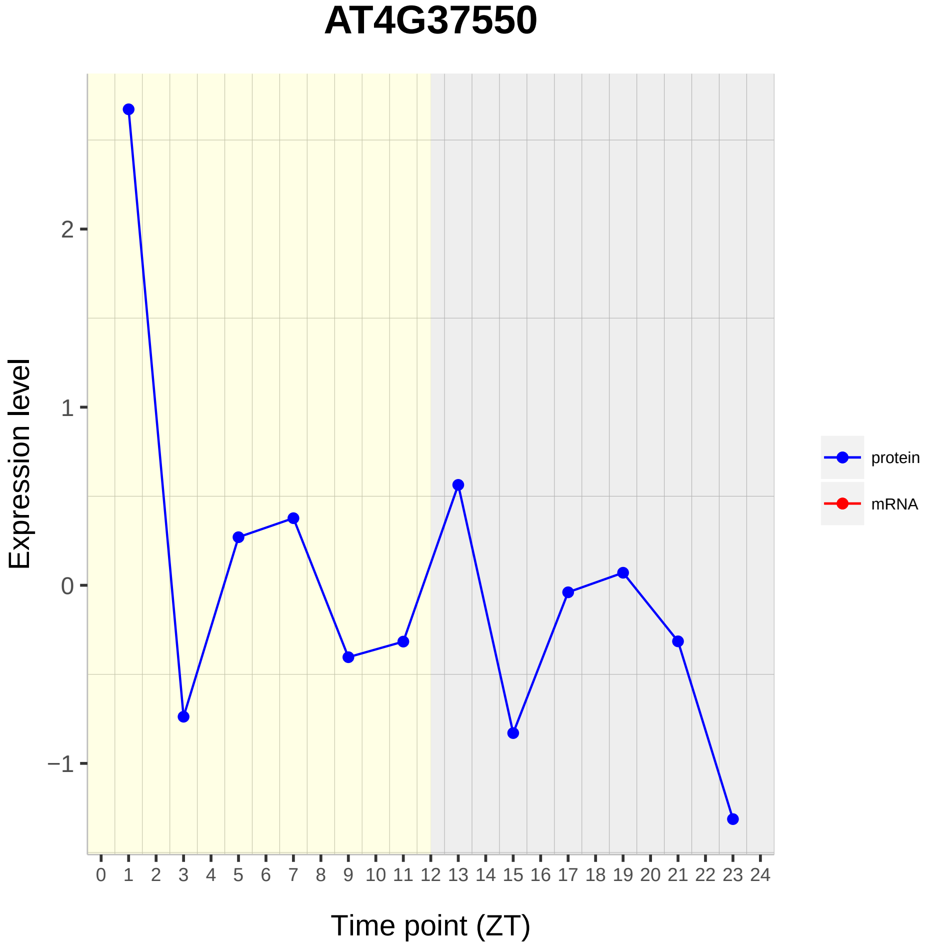

AT4G37550
Expression level
Time point (ZT)
2
1
0
−1
0 1 2 3 4 5 6 7 8 9 10 11 12 13 14 15 16 17 18 19 20 21 22 23 24
protein
mRNA

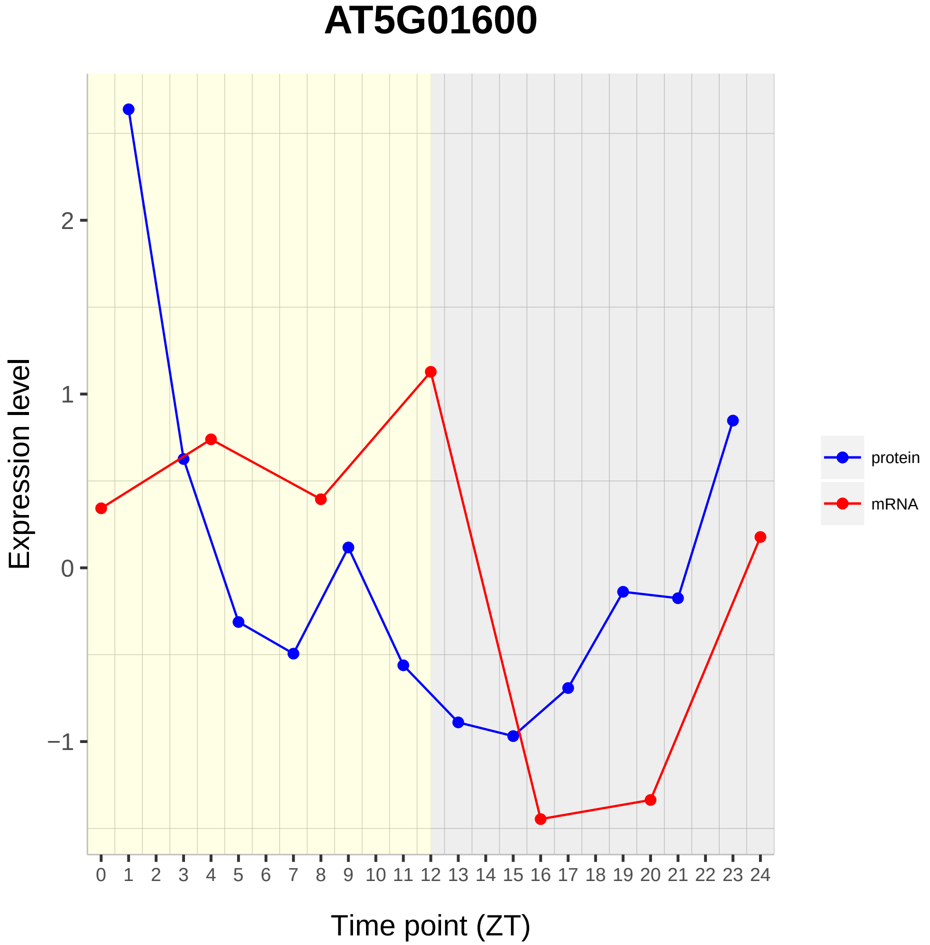
AT5G01600
Expression level
2
1
0
−1
0 1 2 3 4 5 6 7 8 9 10 11 12 13 14 15 16 17 18 19 20 21 22 23 24
Time point (ZT)
protein
mRNA

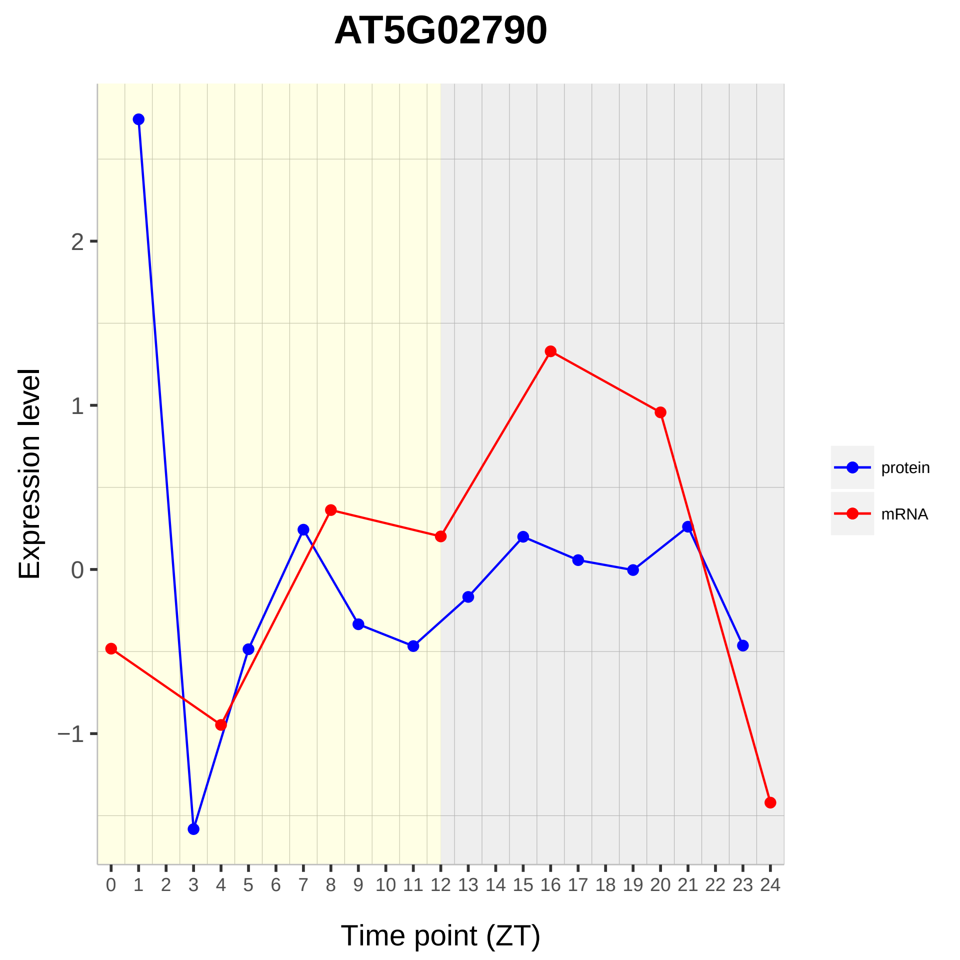

AT5G02790
Expression level
Time point (ZT)
2
1
0
−1
0 1 2 3 4 5 6 7 8 9 10 11 12 13 14 15 16 17 18 19 20 21 22 23 24
protein
mRNA

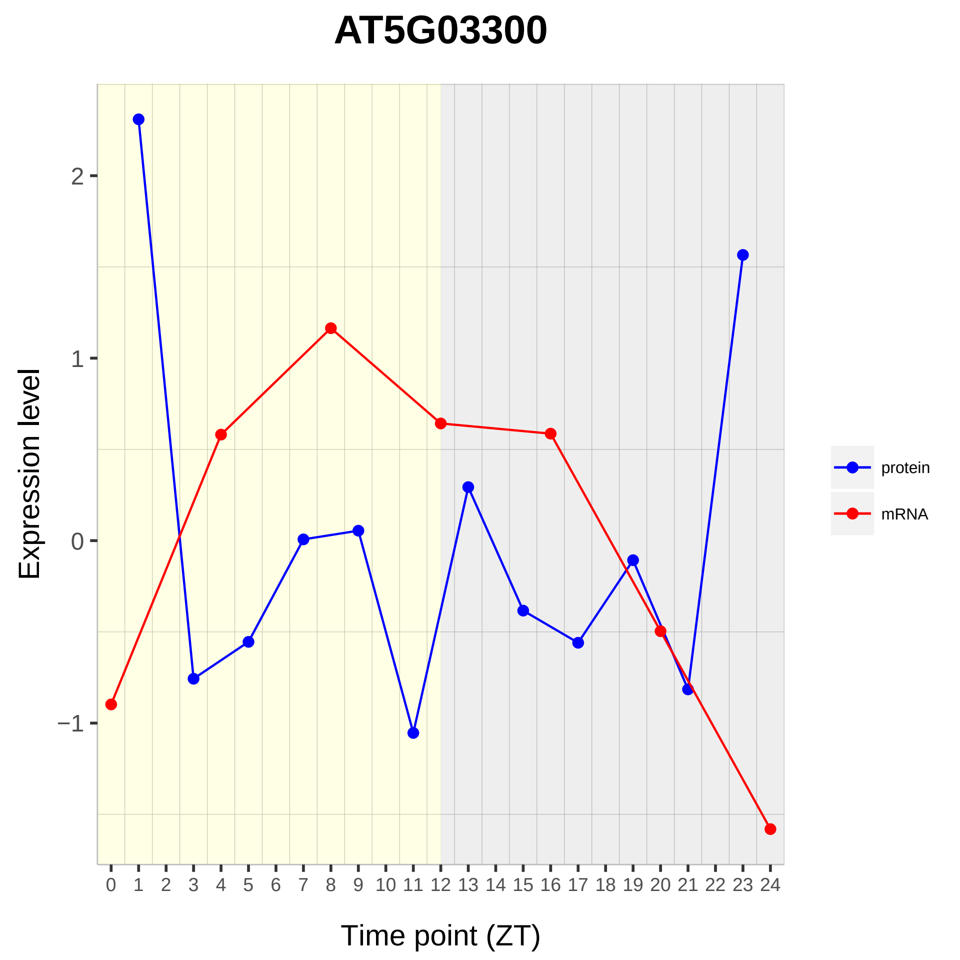

AT5G03300
Expression level
2
1
0
−1
0 1 2 3 4 5 6 7 8 9 10 11 12 13 14 15 16 17 18 19 20 21 22 23 24
Time point (ZT)
protein
mRNA

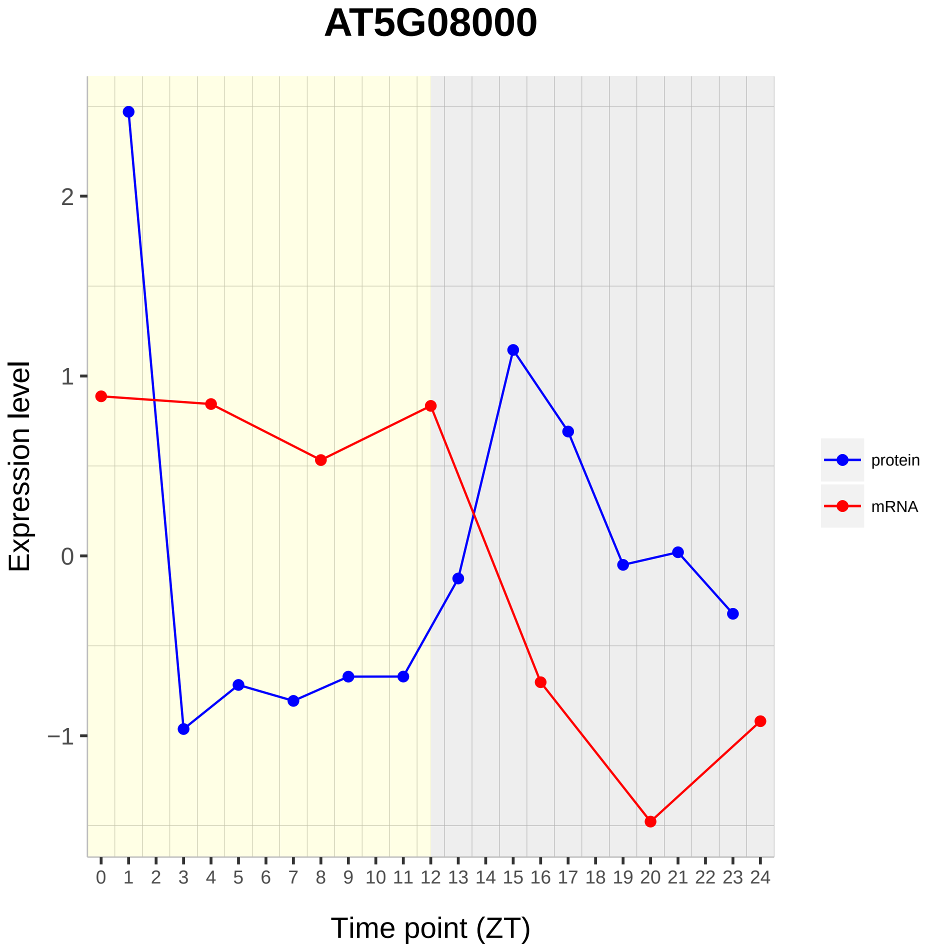
AT5G08000
Expression level
Time point (ZT)
protein
mRNA
2
1
0
−1
0 1 2 3 4 5 6 7 8 9 10 11 12 13 14 15 16 17 18 19 20 21 22 23 24

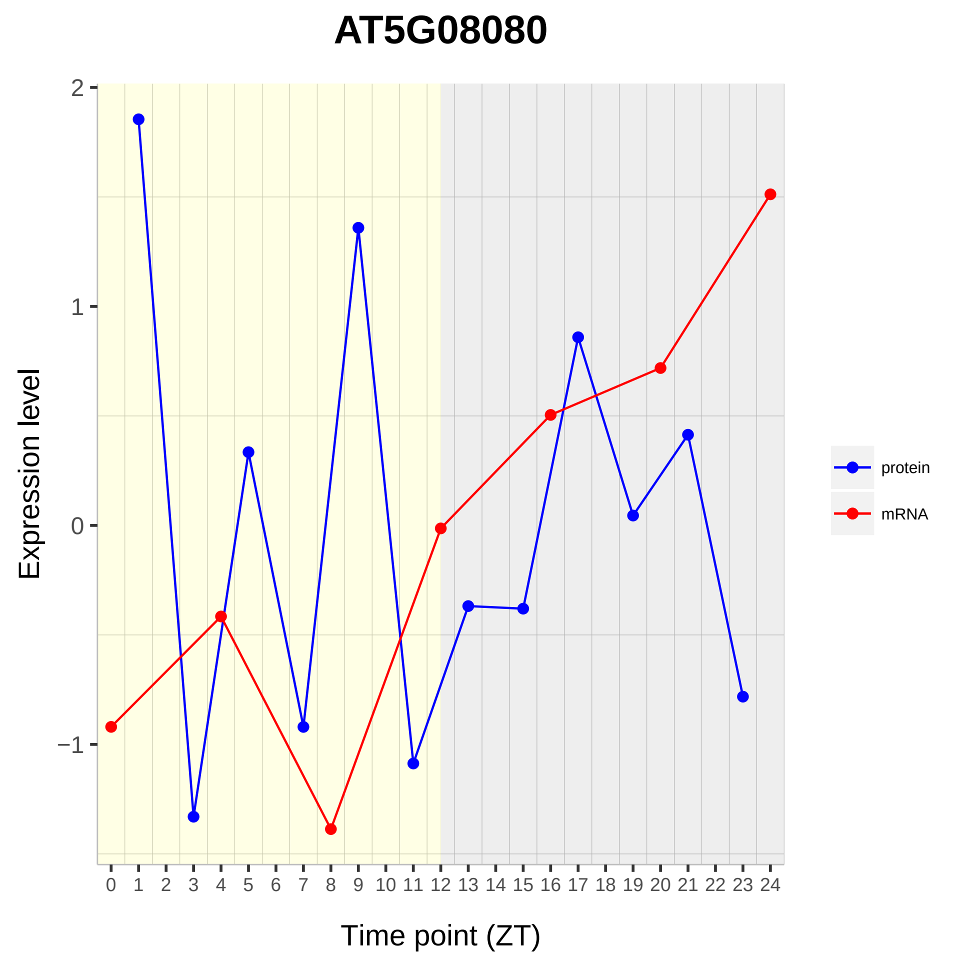

AT5G08080
Expression level
2
1
0
−1
Time point (ZT)
protein
mRNA

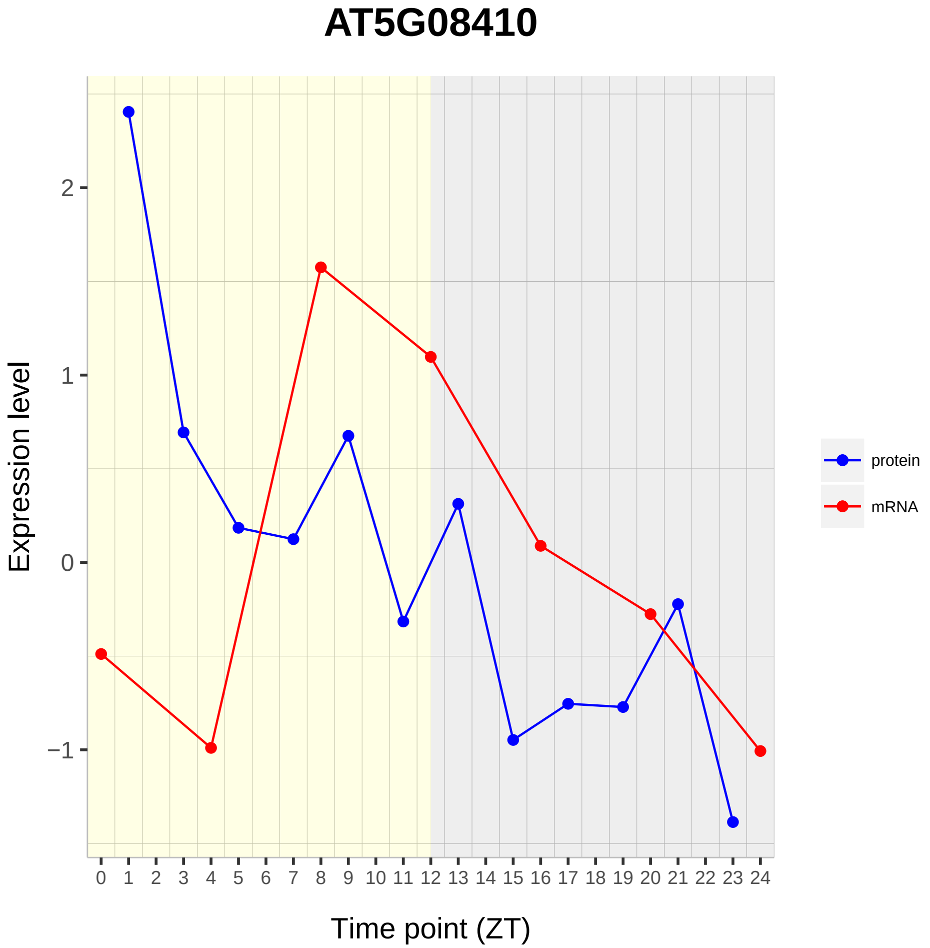
AT5G08410
Expression level
Time point (ZT)
2
1
0
−1
0 1 2 3 4 5 6 7 8 9 10 11 12 13 14 15 16 17 18 19 20 21 22 23 24
protein
mRNA

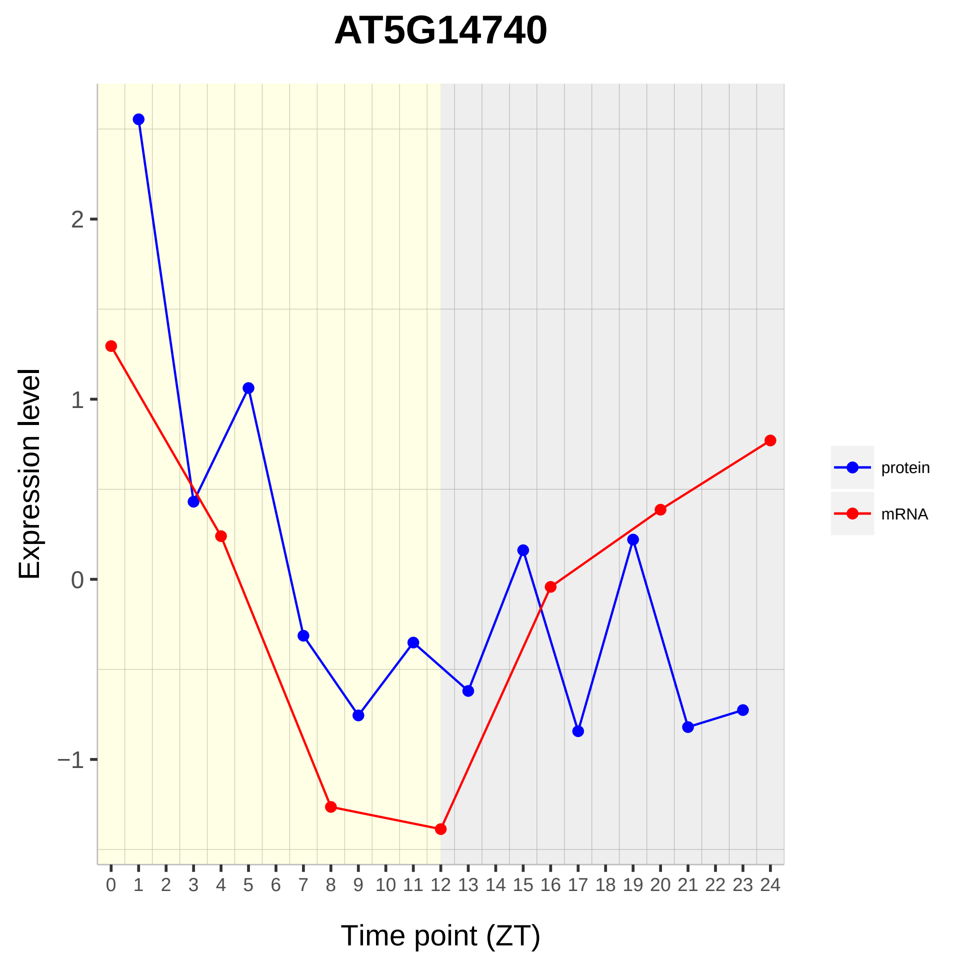

AT5G14740
Expression level
2
1
0
−1
0 1 2 3 4 5 6 7 8 9 10 11 12 13 14 15 16 17 18 19 20 21 22 23 24
Time point (ZT)
protein
mRNA

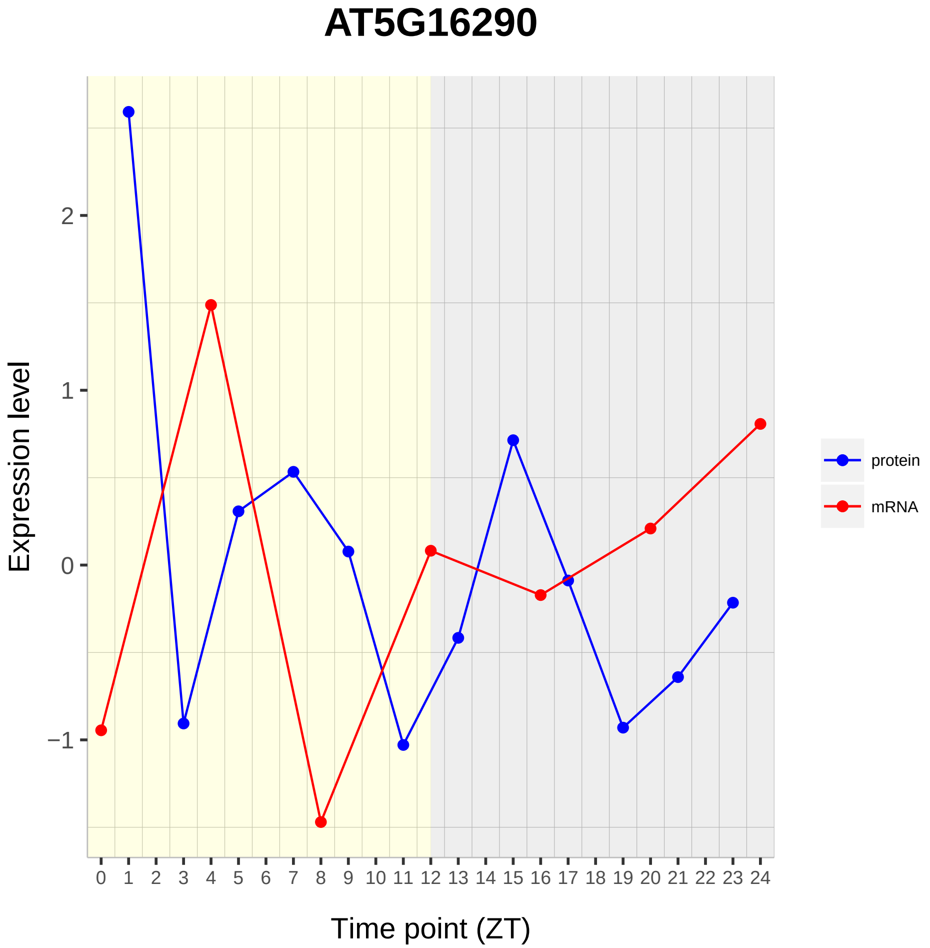
AT5G16290
Expression level
2
1
0
−1
0 1 2 3 4 5 6 7 8 9 10 11 12 13 14 15 16 17 18 19 20 21 22 23 24
Time point (ZT)
protein
mRNA

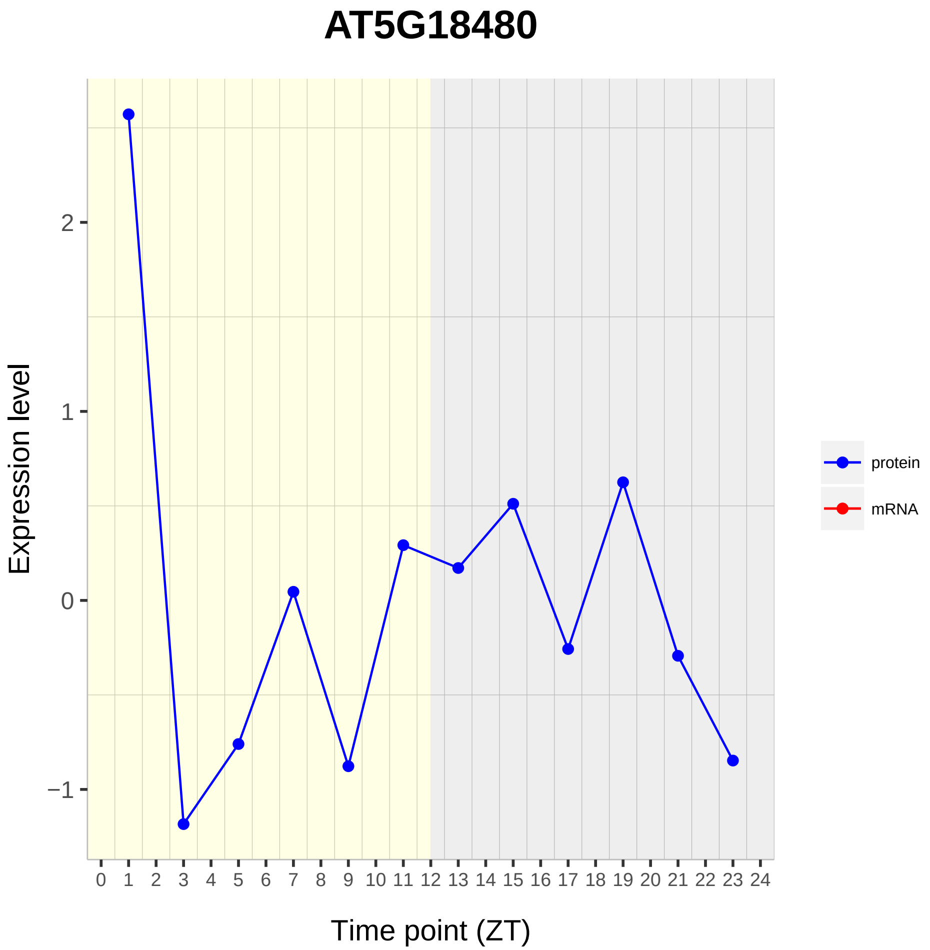
AT5G18480
Expression level
2
1
0
−1
0 1 2 3 4 5 6 7 8 9 10 11 12 13 14 15 16 17 18 19 20 21 22 23 24
Time point (ZT)
protein
mRNA

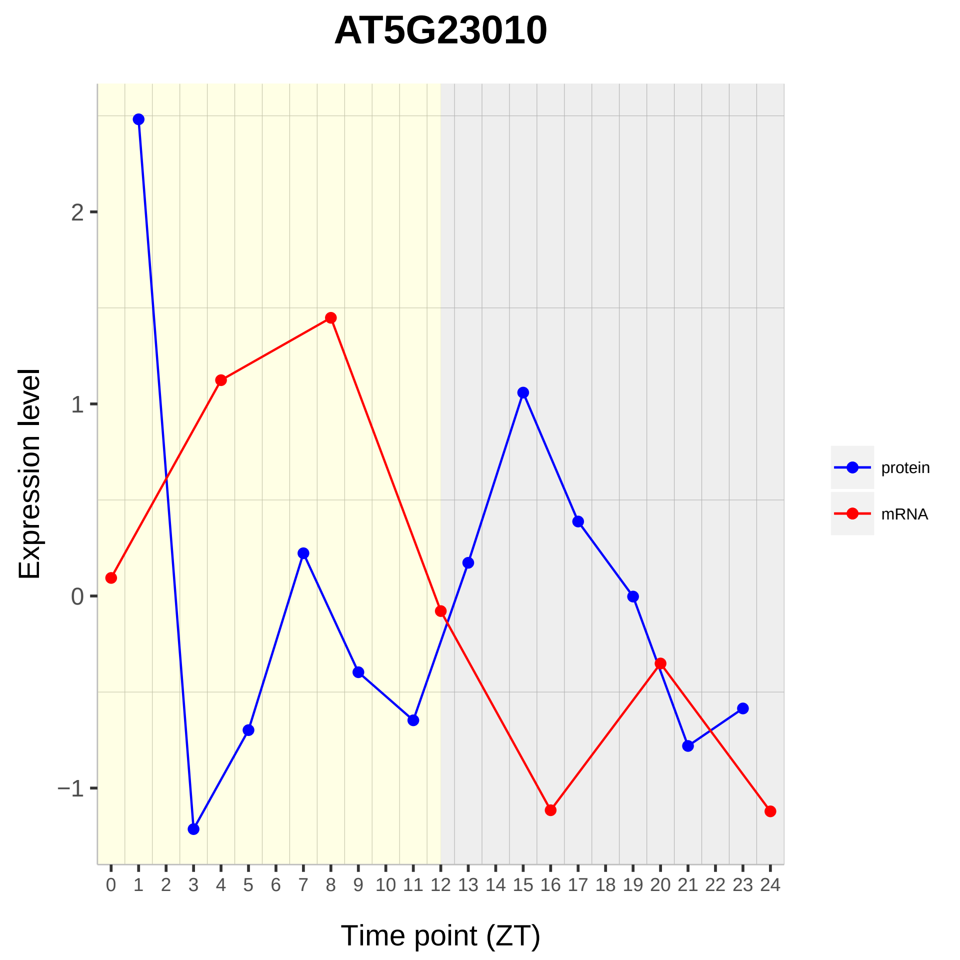
AT5G23010
Expression level
2
1
0
−1
0 1 2 3 4 5 6 7 8 9 10 11 12 13 14 15 16 17 18 19 20 21 22 23 24
Time point (ZT)
protein
mRNA

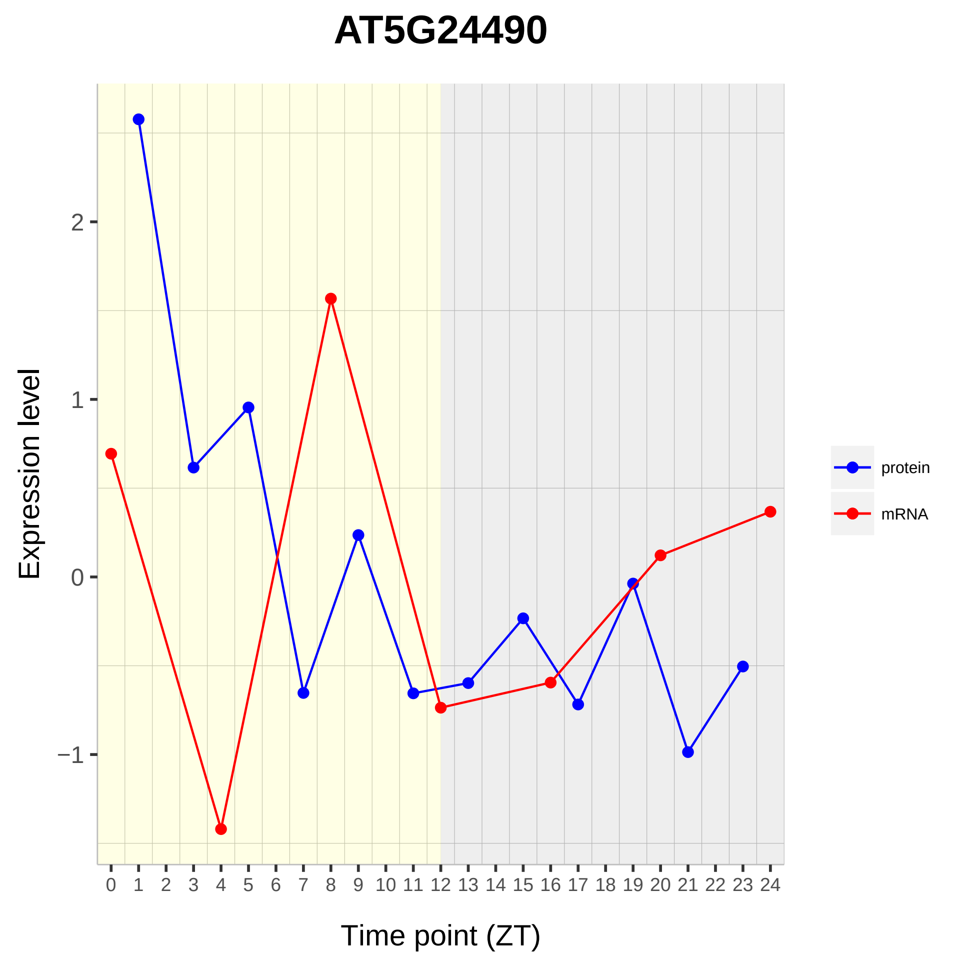
AT5G24490
Expression level
2
1
0
−1
0 1 2 3 4 5 6 7 8 9 10 11 12 13 14 15 16 17 18 19 20 21 22 23 24
Time point (ZT)
protein
mRNA

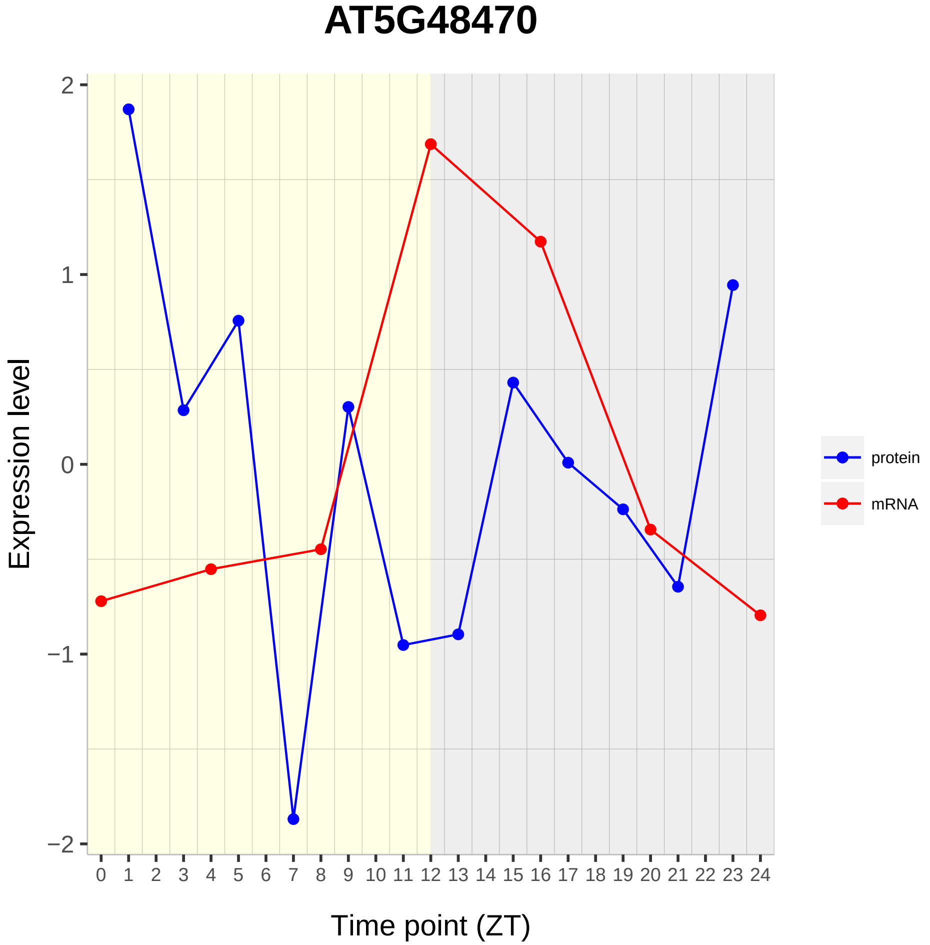
AT5G48470
Expression level
Time point (ZT)
2
1
0
−1
−2
protein
mRNA

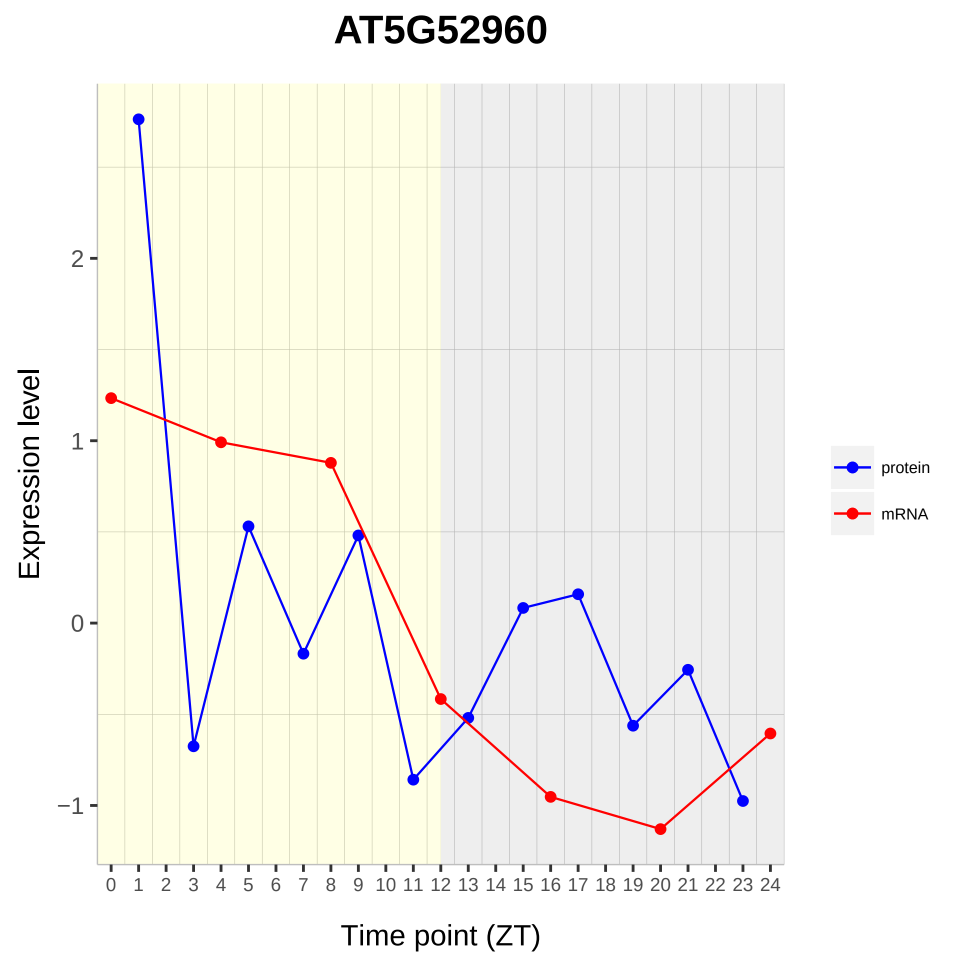
AT5G52960
Expression level
2
1
0
−1
0 1 2 3 4 5 6 7 8 9 10 11 12 13 14 15 16 17 18 19 20 21 22 23 24
Time point (ZT)
protein
mRNA

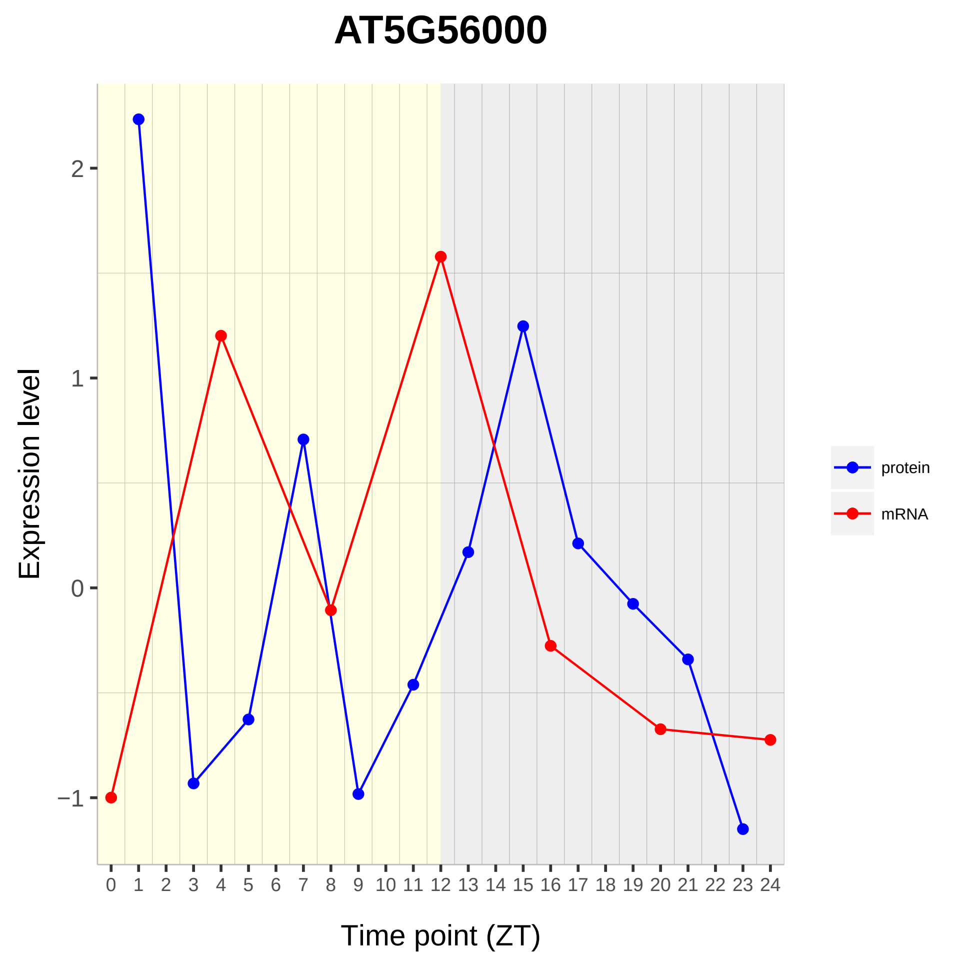

AT5G56000
Expression level
Time point (ZT)
2
1
0
−1
0 1 2 3 4 5 6 7 8 9 10 11 12 13 14 15 16 17 18 19 20 21 22 23 24
protein
mRNA

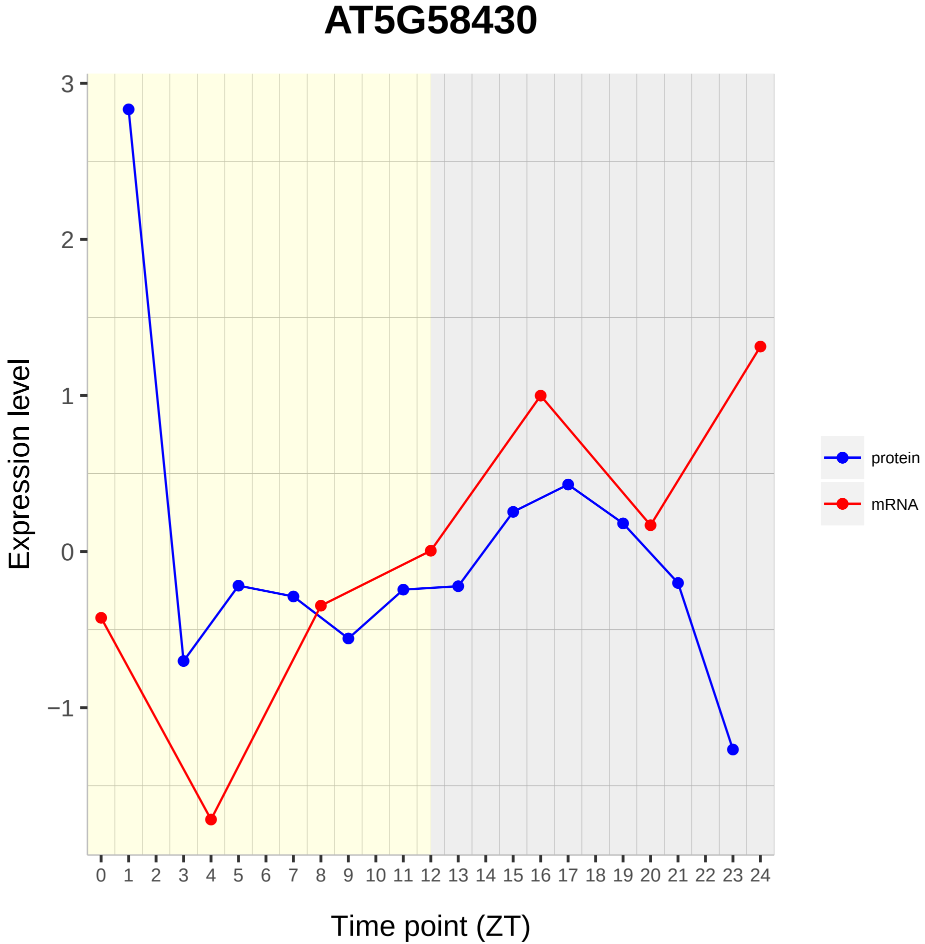

AT5G58430
Expression level
Time point (ZT)
3
2
1
0
−1
protein
mRNA

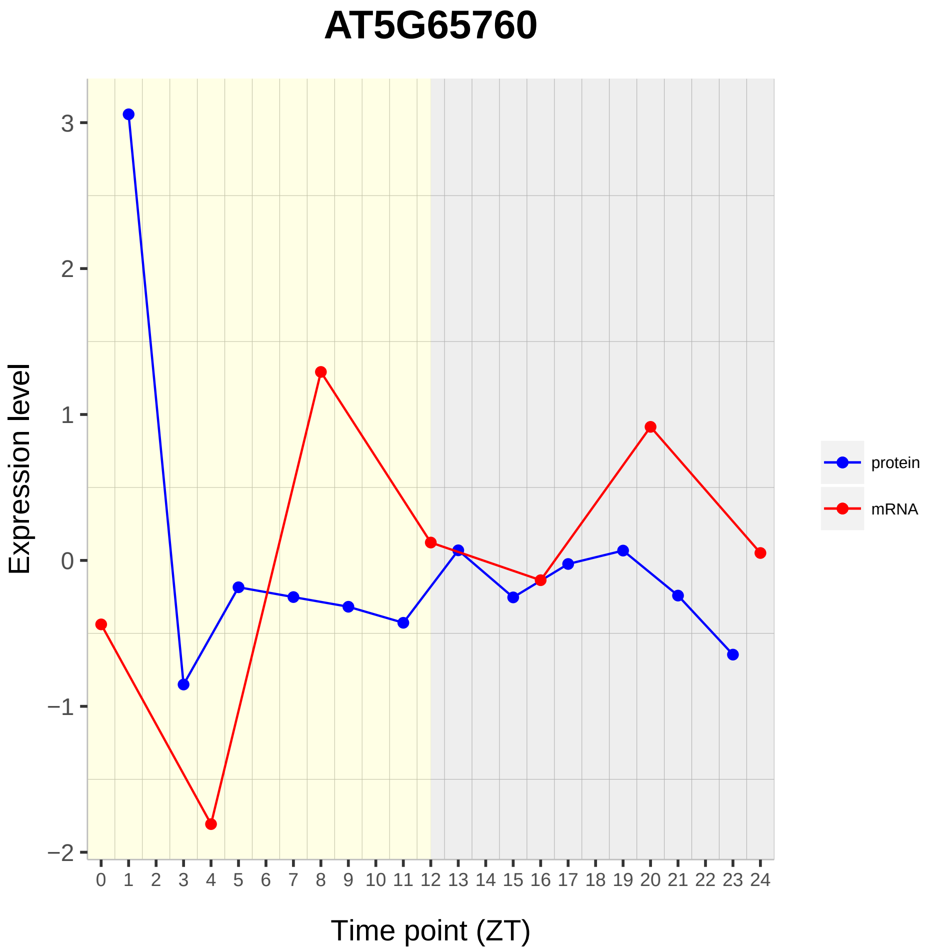

AT5G65760
Expression level
Time point (ZT)
3
2
1
0
−1
−2
0 1 2 3 4 5 6 7 8 9 10 11 12 13 14 15 16 17 18 19 20 21 22 23 24
protein
mRNA

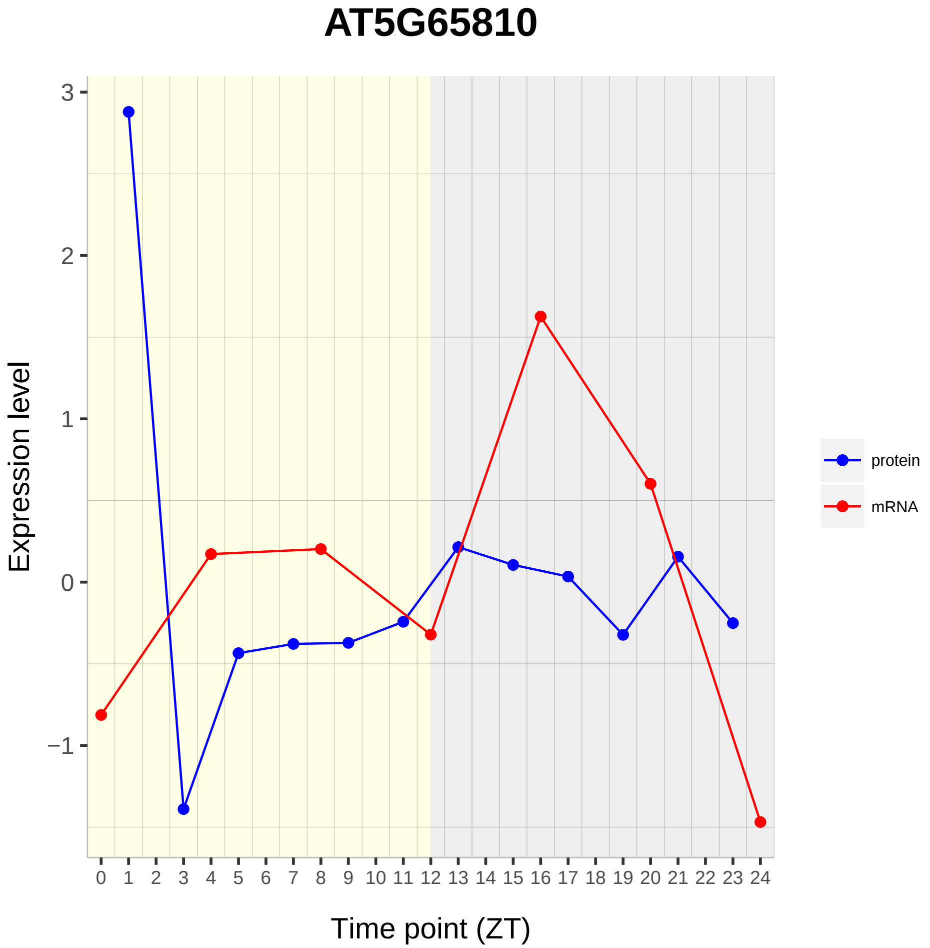

AT5G65810
Expression level
3
2
1
0
−1
0 1 2 3 4 5 6 7 8 9 10 11 12 13 14 15 16 17 18 19 20 21 22 23 24
Time point (ZT)
protein
mRNA
