## Supplementary material for "Diurnal Dynamics of the Arabidopsis Rosette Proteome and Phosphoproteome": SuppData3

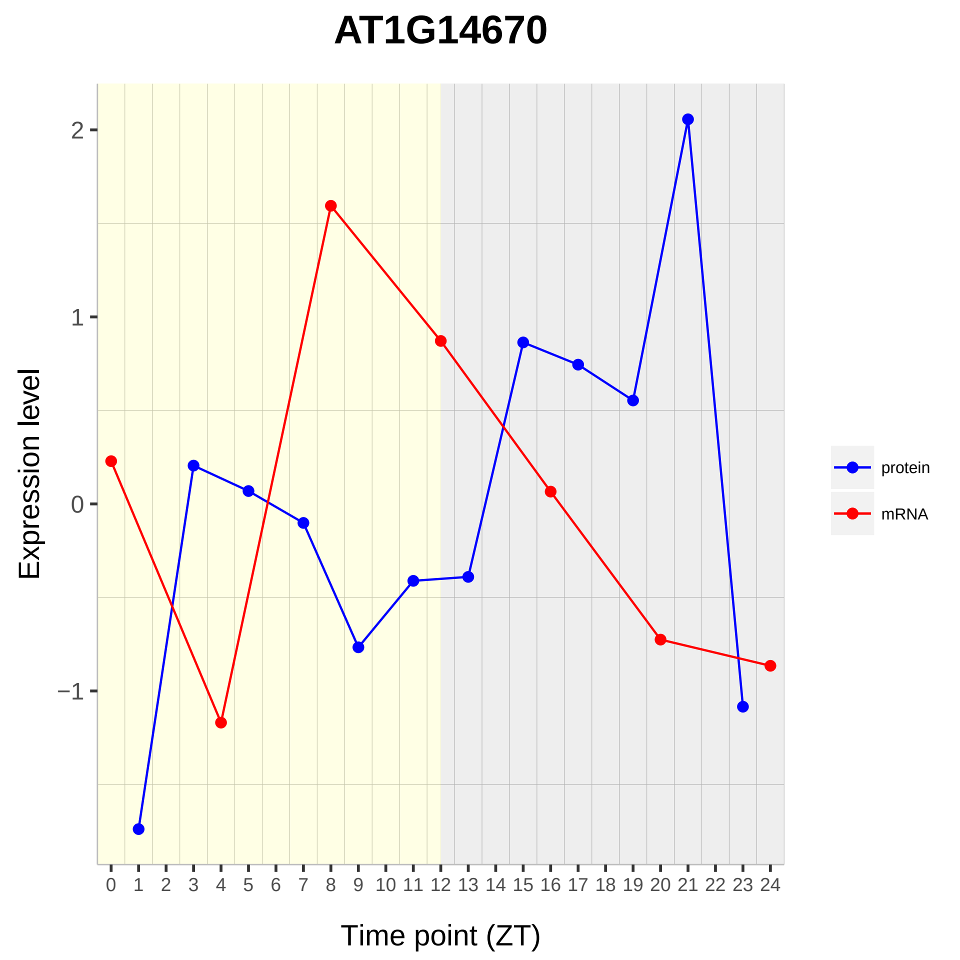

AT1G14670
Expression level
Time point (ZT)
2
1
0
−1
0 1 2 3 4 5 6 7 8 9 10 11 12 13 14 15 16 17 18 19 20 21 22 23 24
protein
mRNA

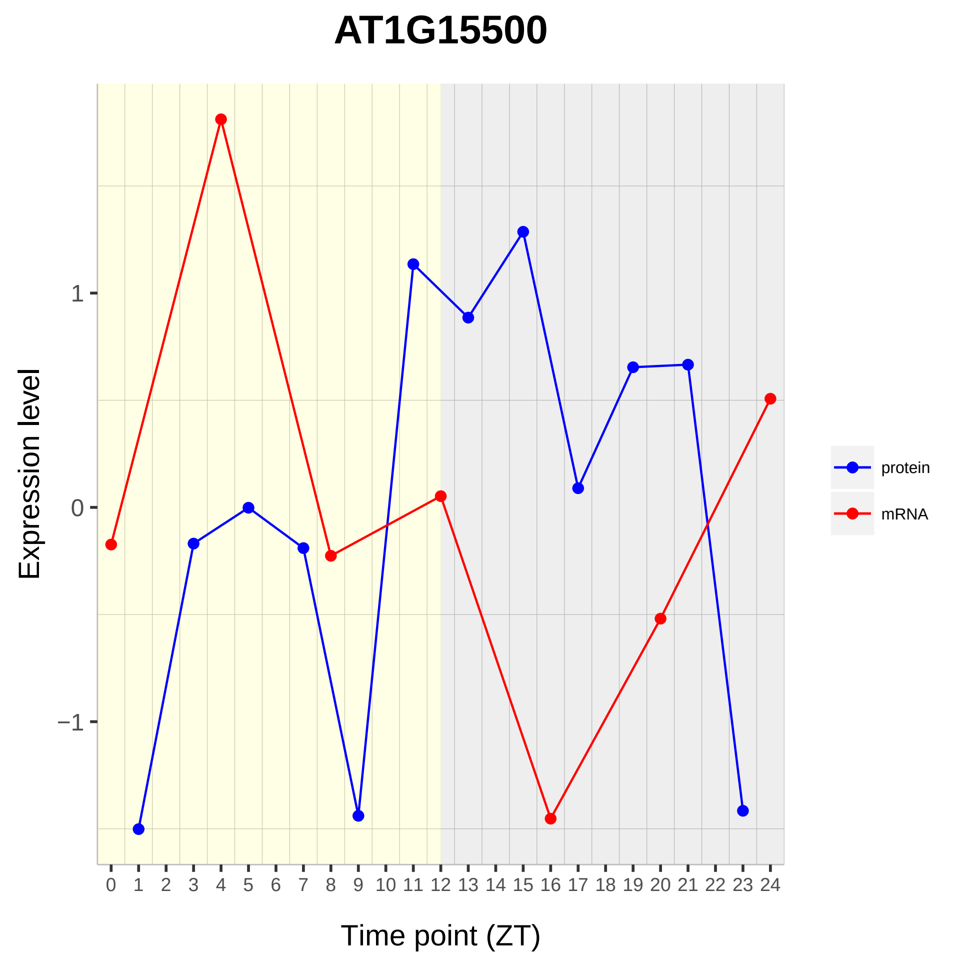
AT1G15500
Expression level
1
0
−1
Time point (ZT)
protein
mRNA

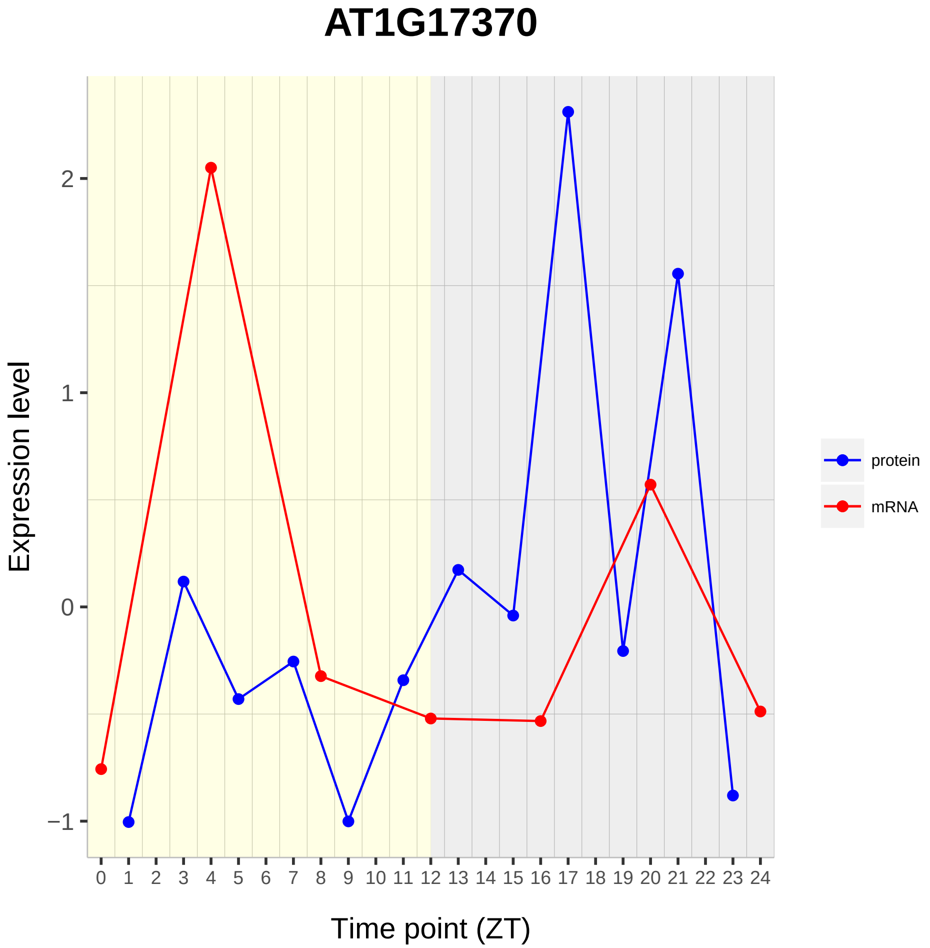

AT1G17370
Expression level
2
1
0
−1
Time point (ZT)
protein
mRNA

# AT1G25260

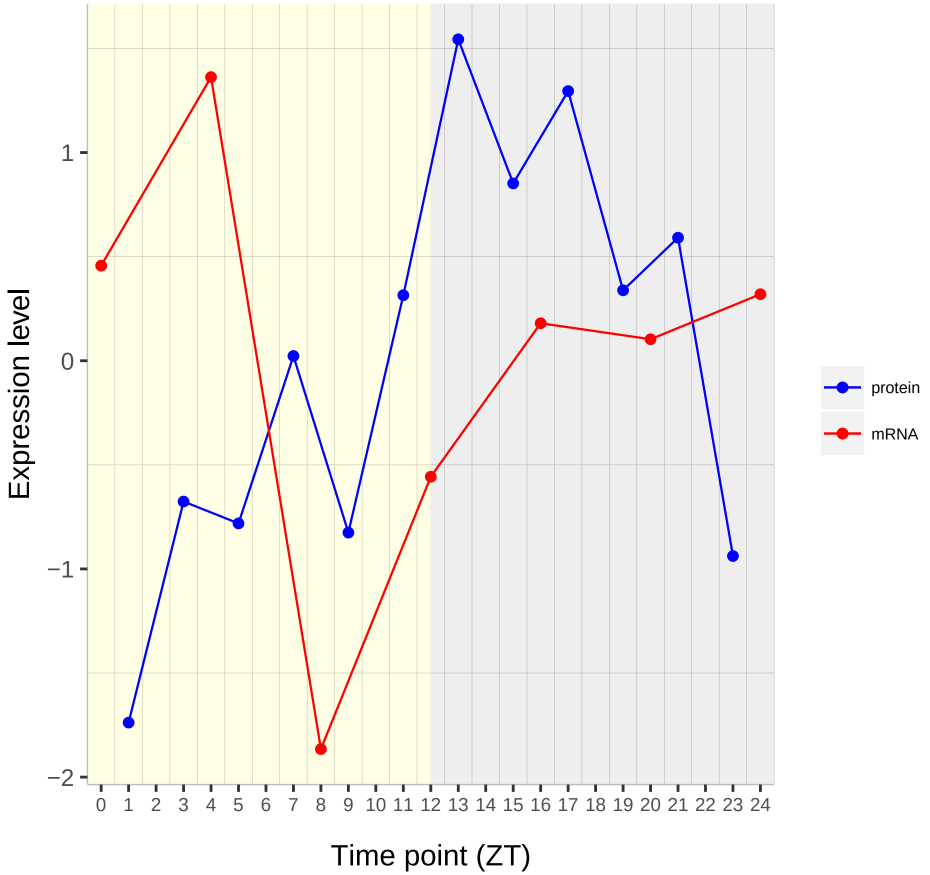

Expression level
1
0
−1
−2
0 1 2 3 4 5 6 7 8 9 10 11 12 13 14 15 16 17 18 19 20 21 22 23 24
Time point (ZT)
protein
mRNA

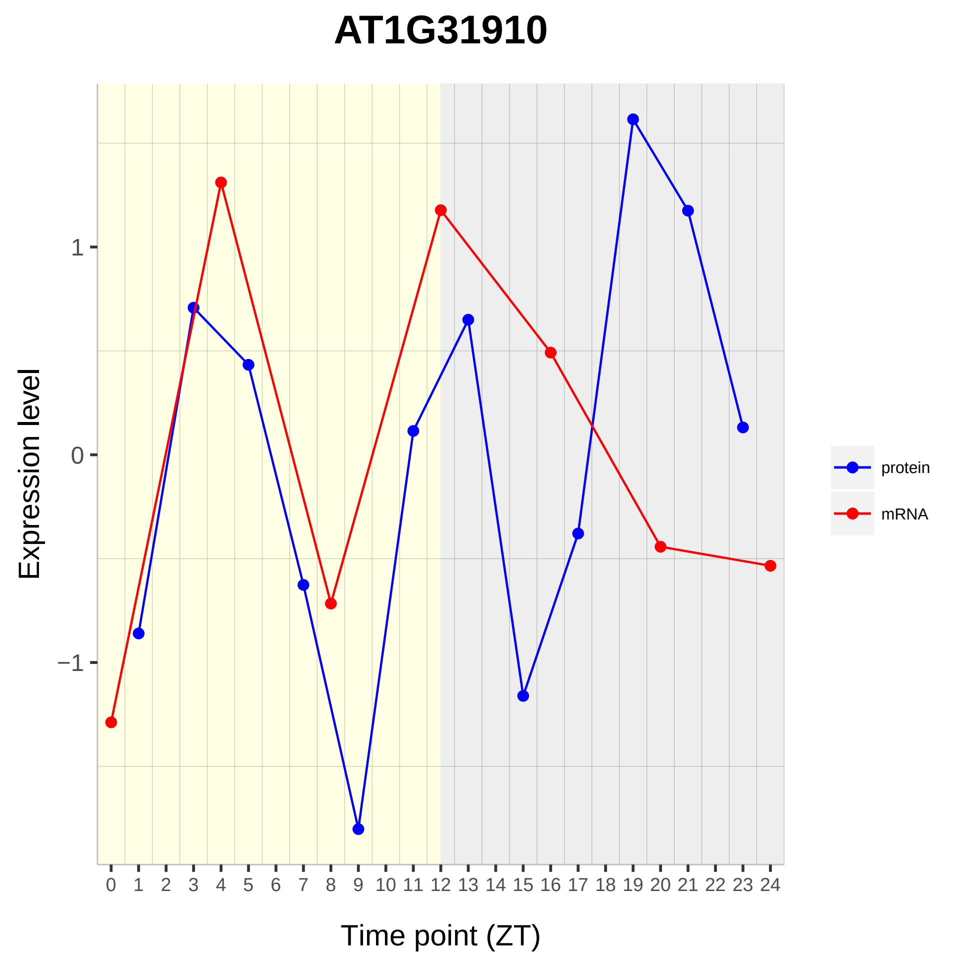

AT1G31910
Expression level
1
0
−1
0 1 2 3 4 5 6 7 8 9 10 11 12 13 14 15 16 17 18 19 20 21 22 23 24
Time point (ZT)
protein
mRNA

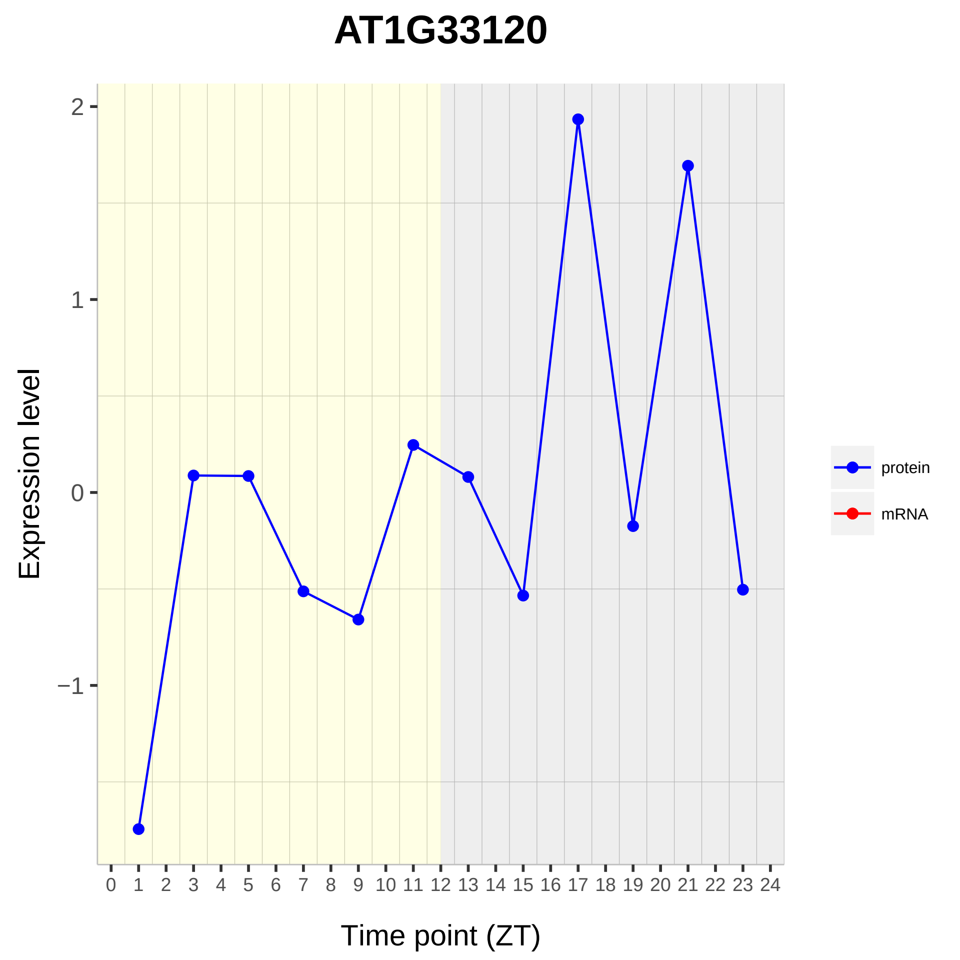

AT1G33120
Expression level
Time point (ZT)
protein
mRNA
2
1
0
−1
0 1 2 3 4 5 6 7 8 9 10 11 12 13 14 15 16 17 18 19 20 21 22 23 24

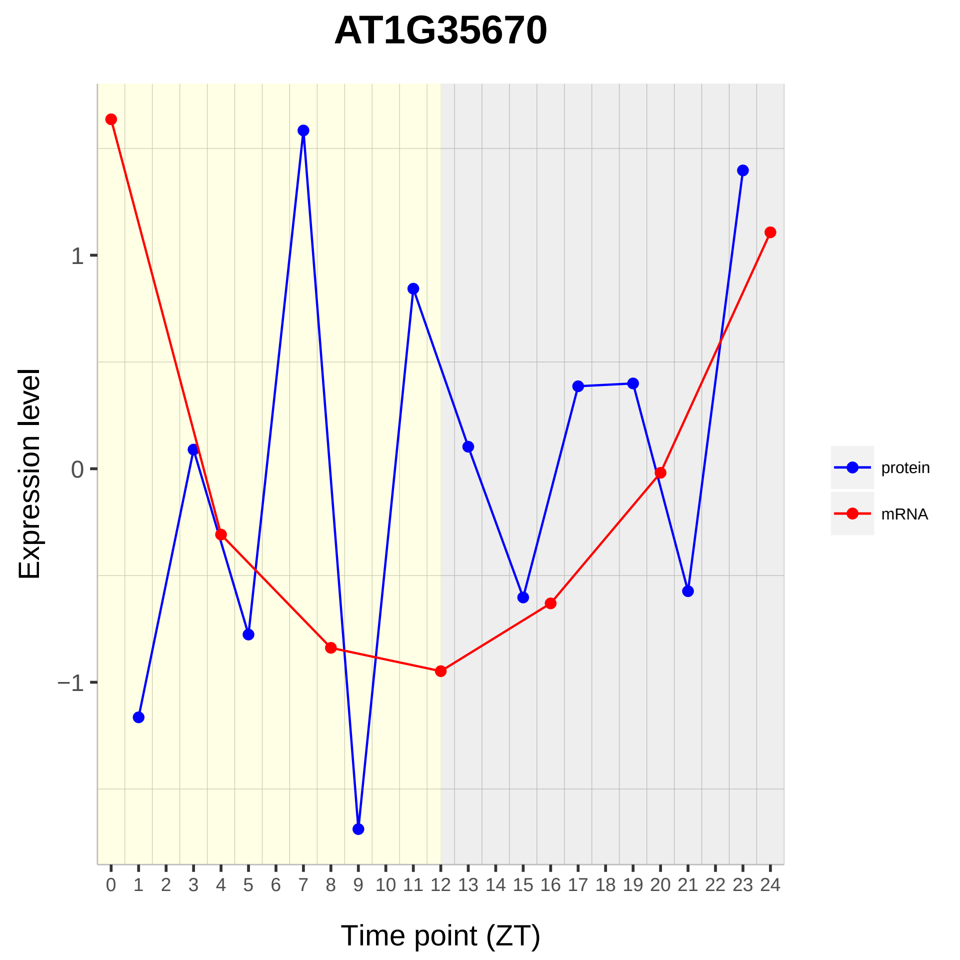
AT1G35670
Expression level
1
0
−1
Time point (ZT)
protein
mRNA

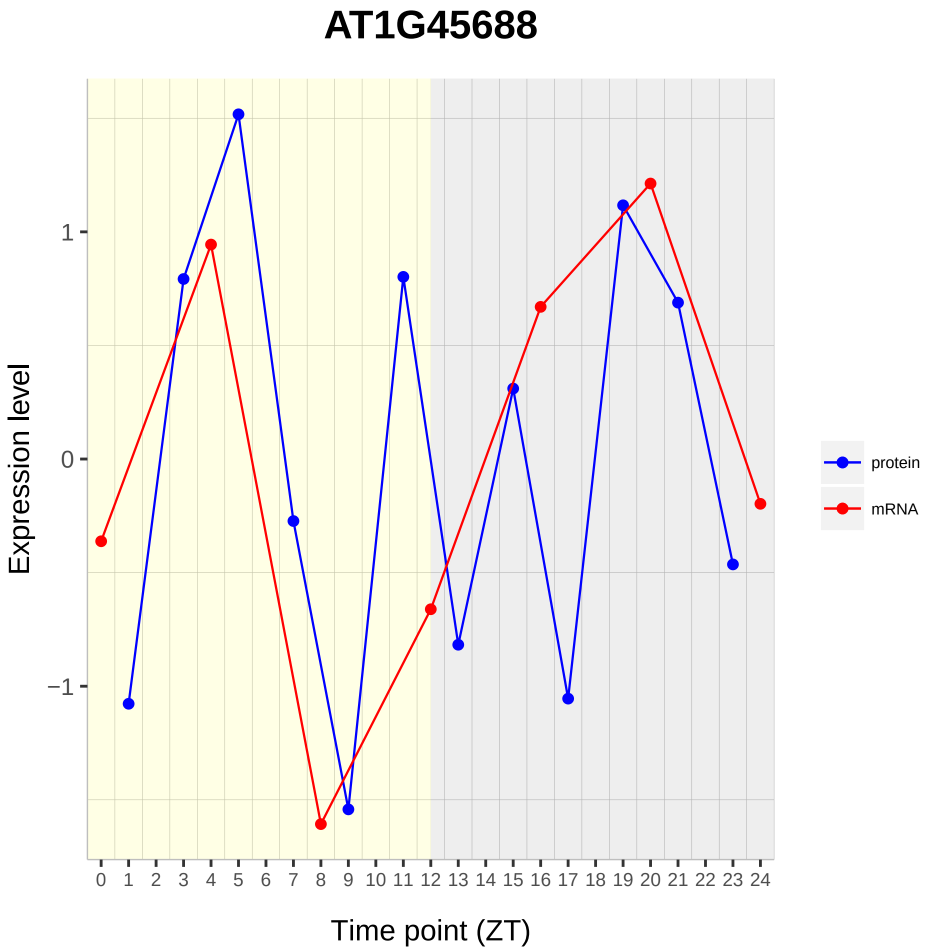
AT1G45688
Expression level
1
0
−1
Time point (ZT)
protein
mRNA

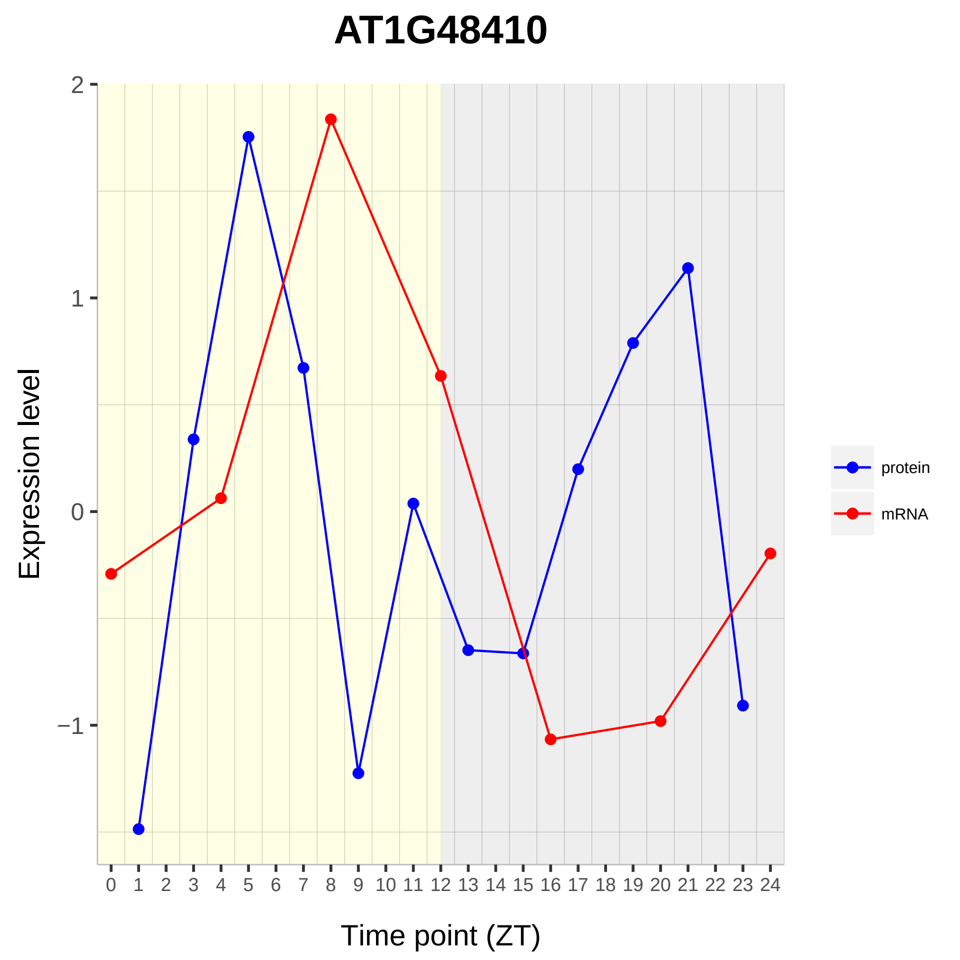

AT1G48410
Expression level
2
1
0
−1
0 1 2 3 4 5 6 7 8 9 10 11 12 13 14 15 16 17 18 19 20 21 22 23 24
Time point (ZT)
protein
mRNA

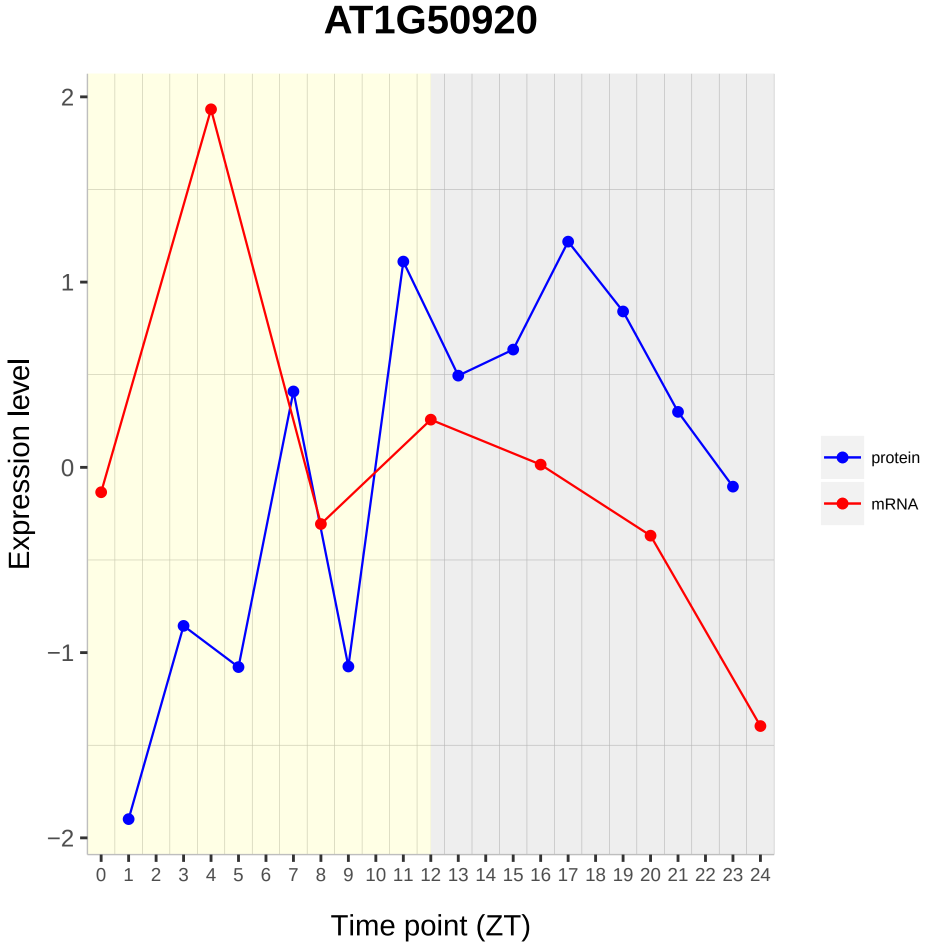

AT1G50920
Expression level
Time point (ZT)
protein
mRNA
2
1
0
−1
−2
0 1 2 3 4 5 6 7 8 9 10 11 12 13 14 15 16 17 18 19 20 21 22 23 24

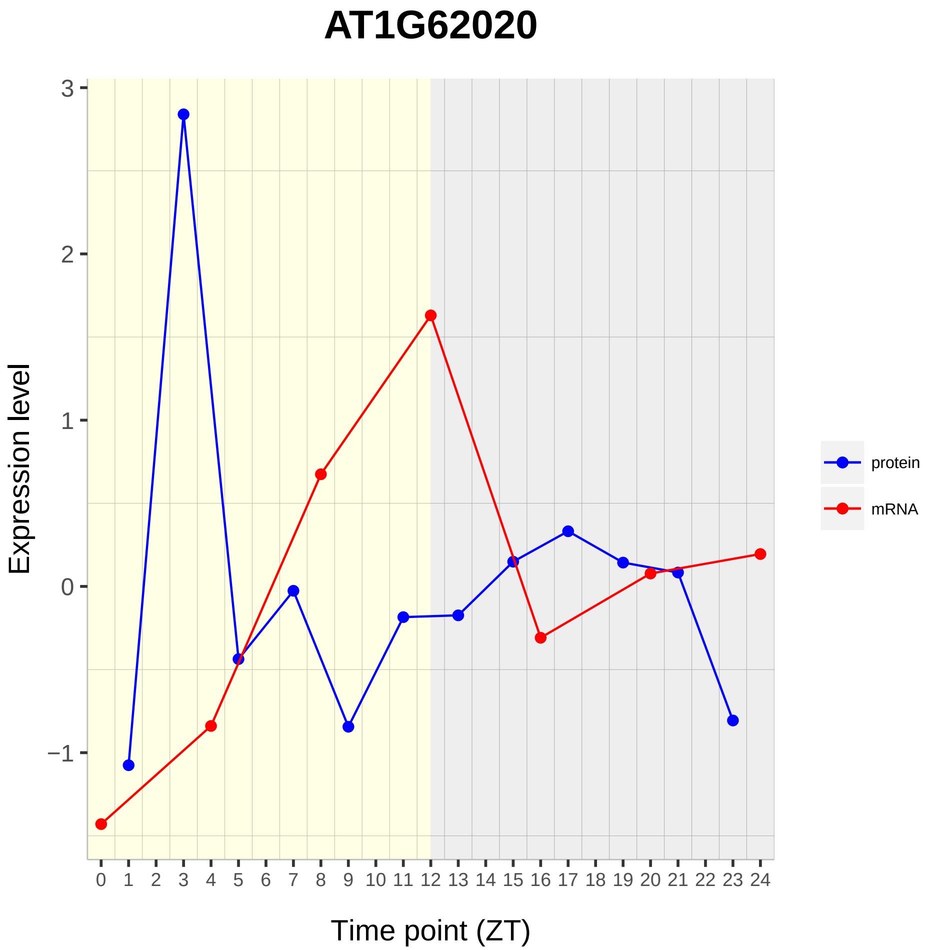

AT1G62020
Expression level
Time point (ZT)
3
2
1
0
−1
0 1 2 3 4 5 6 7 8 9 10 11 12 13 14 15 16 17 18 19 20 21 22 23 24
protein
mRNA

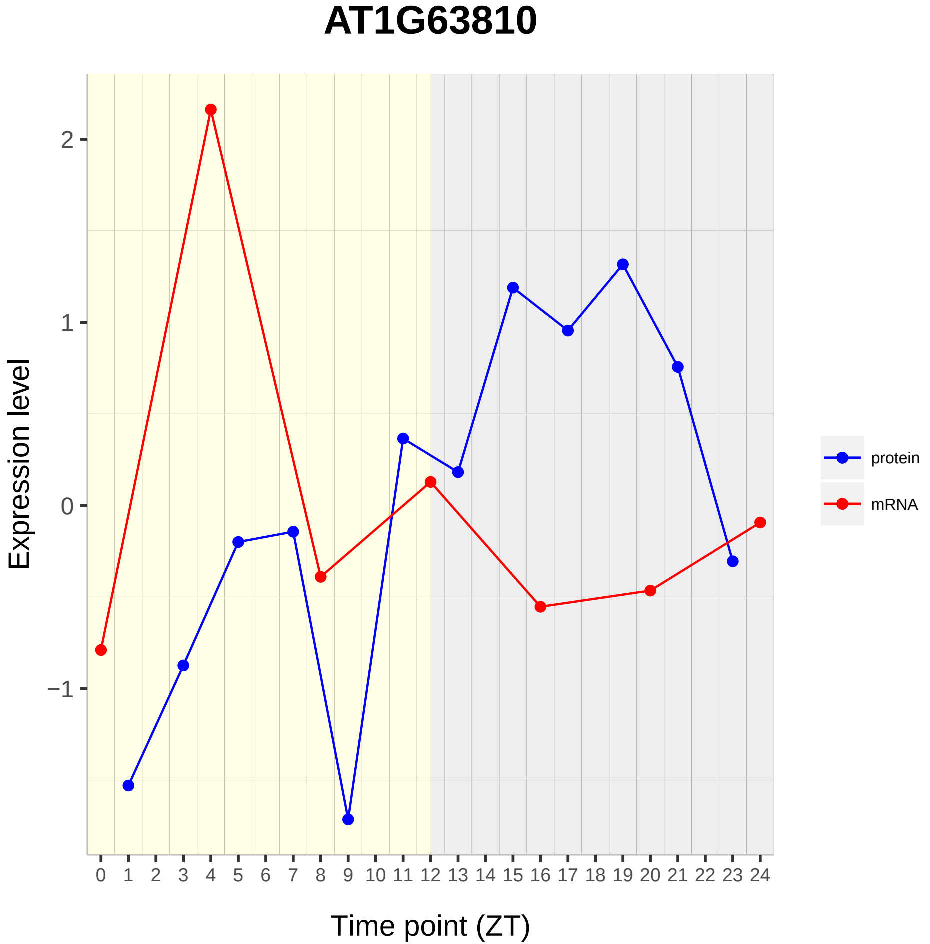

AT1G63810
Expression level
Time point (ZT)
2
1
0
−1
protein
mRNA

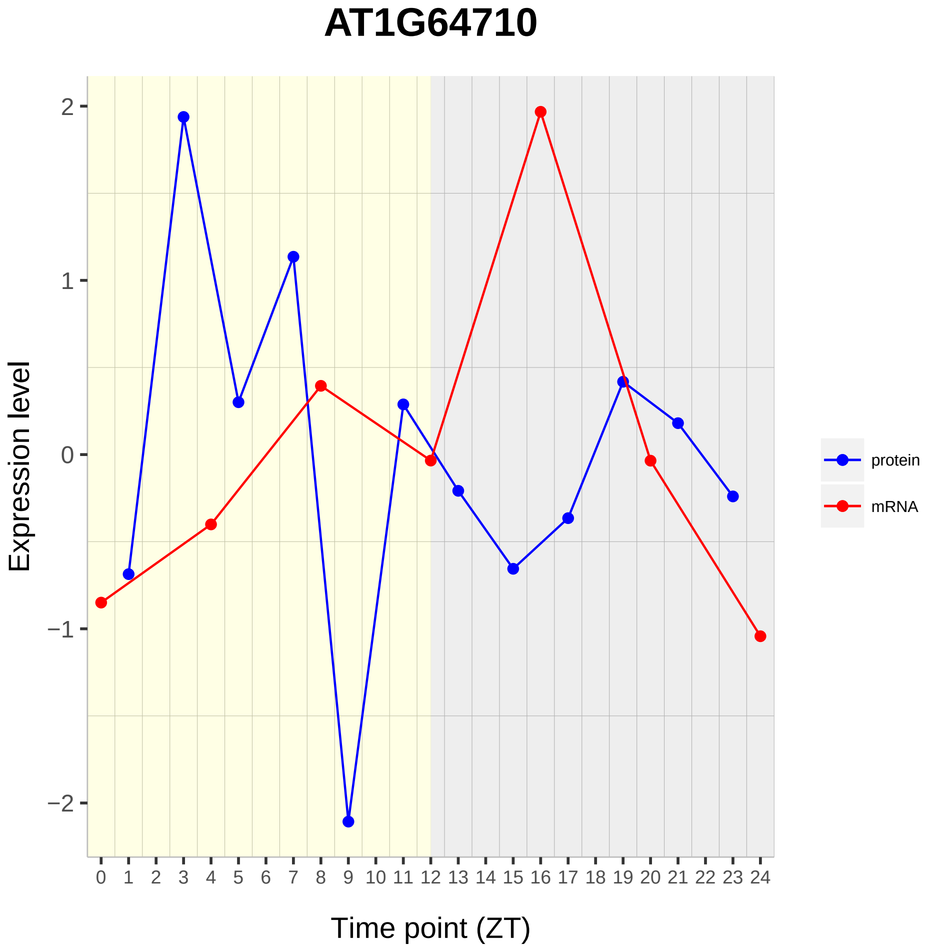

AT1G64710
Expression level
2
1
0
−1
−2
Time point (ZT)
protein
mRNA

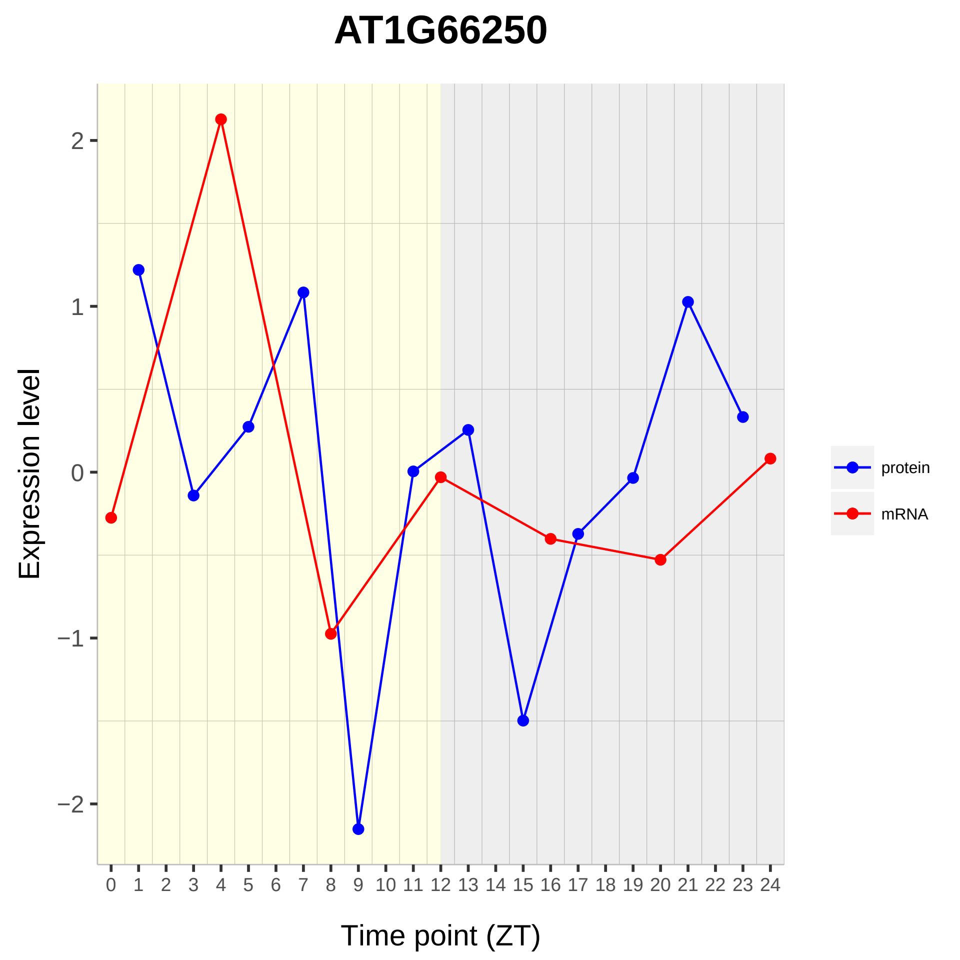
AT1G66250
Expression level
2
1
0
−1
−2
0 1 2 3 4 5 6 7 8 9 10 11 12 13 14 15 16 17 18 19 20 21 22 23 24
Time point (ZT)
protein
mRNA

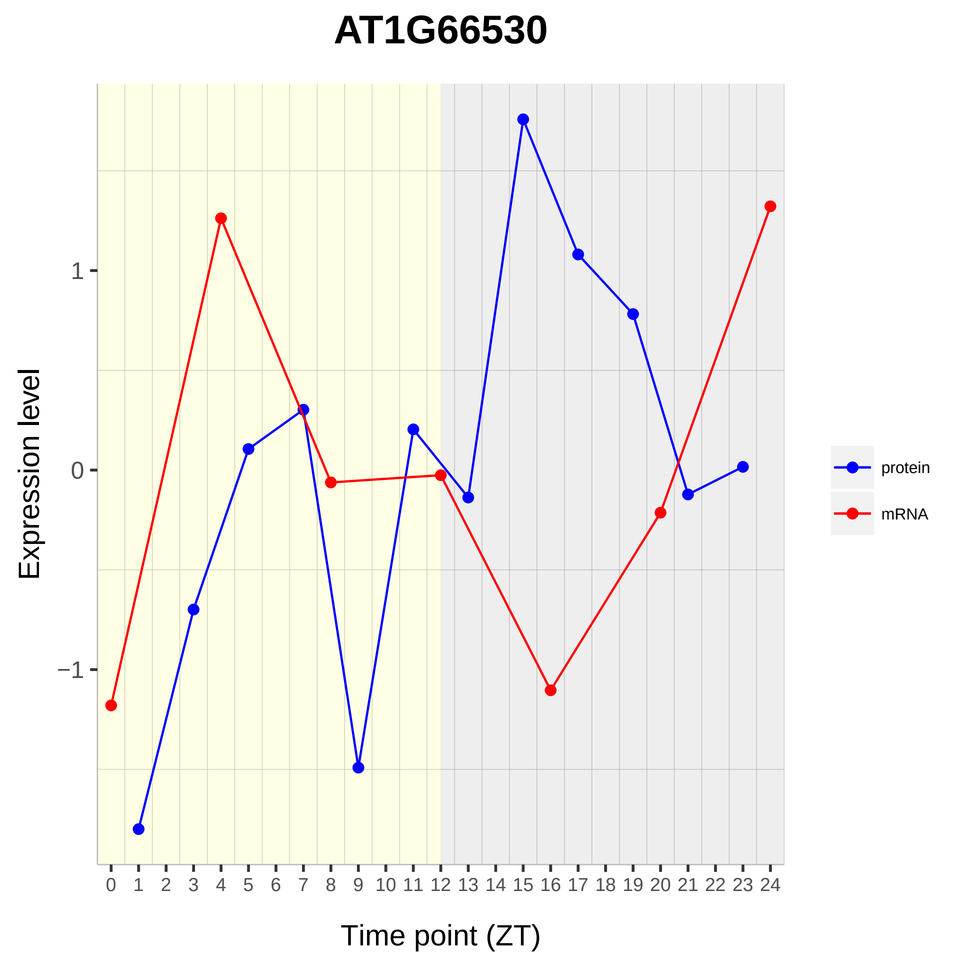

AT1G66530
Expression level
1
0
−1
0 1 2 3 4 5 6 7 8 9 10 11 12 13 14 15 16 17 18 19 20 21 22 23 24
Time point (ZT)
protein
mRNA

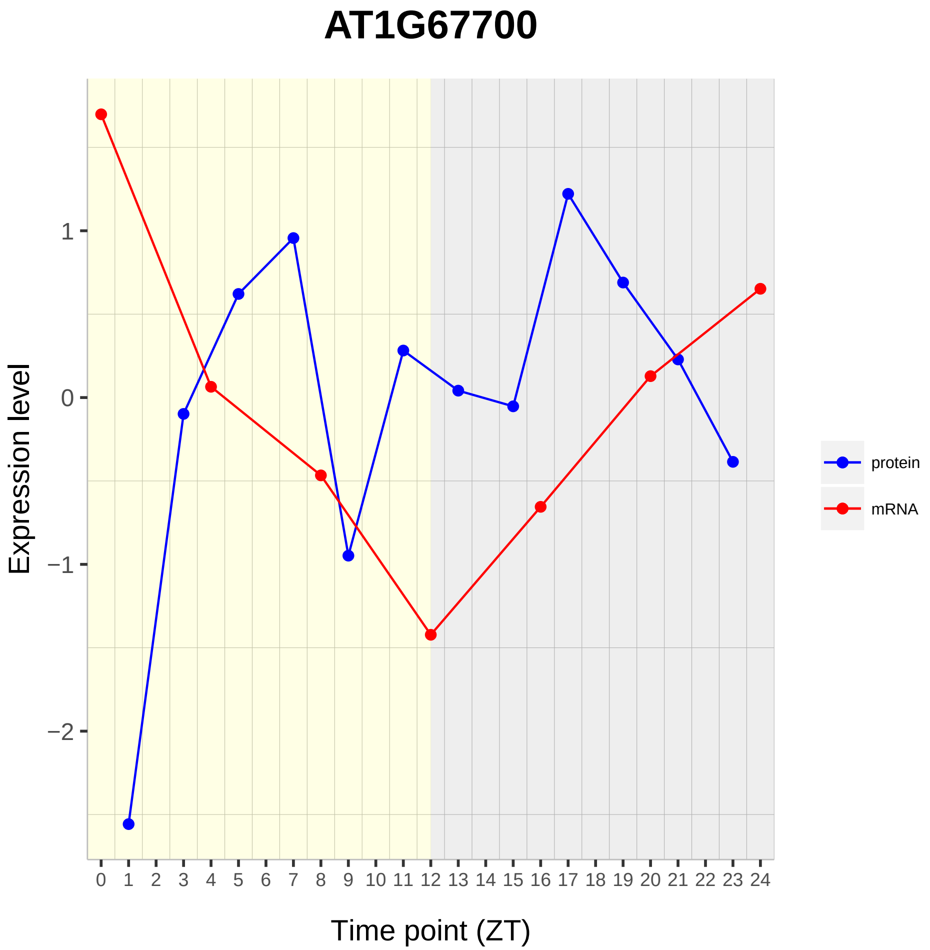
AT1G67700
Expression level
Time point (ZT)
1
0
−1
−2
0 1 2 3 4 5 6 7 8 9 10 11 12 13 14 15 16 17 18 19 20 21 22 23 24
protein
mRNA

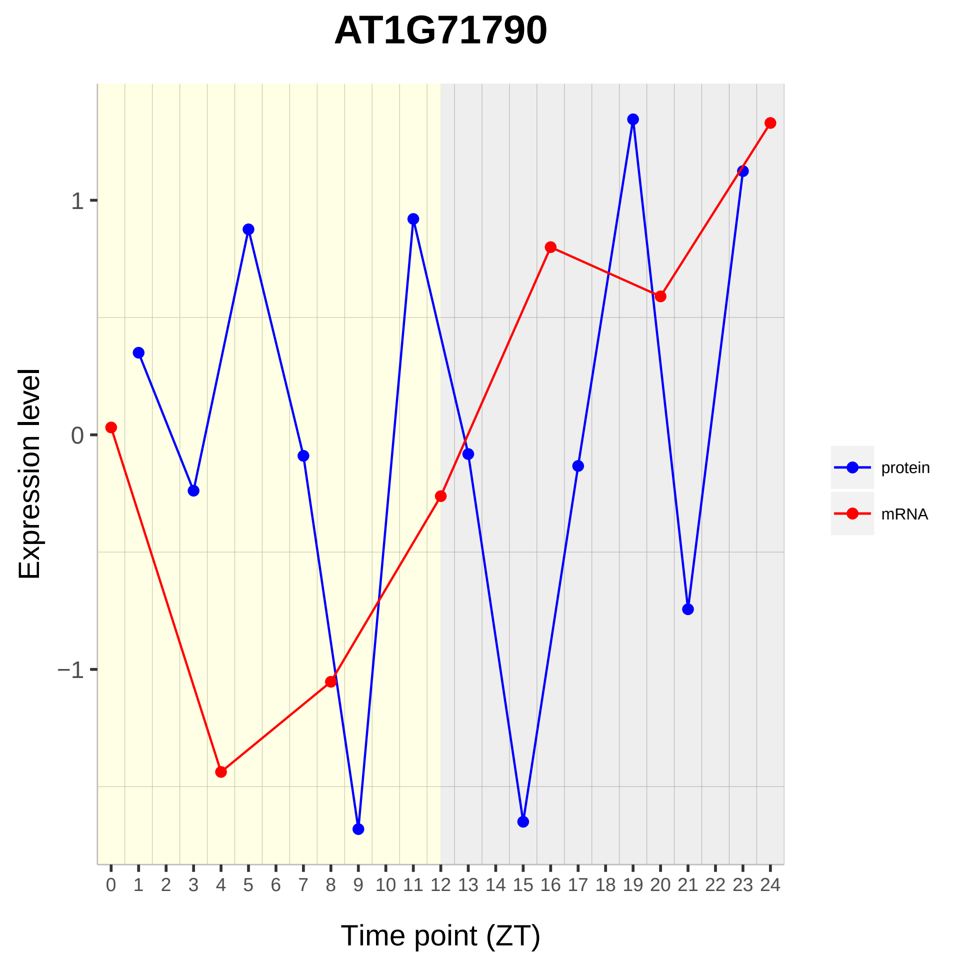

AT1G71790
Expression level
1
0
−1
Time point (ZT)
protein
mRNA

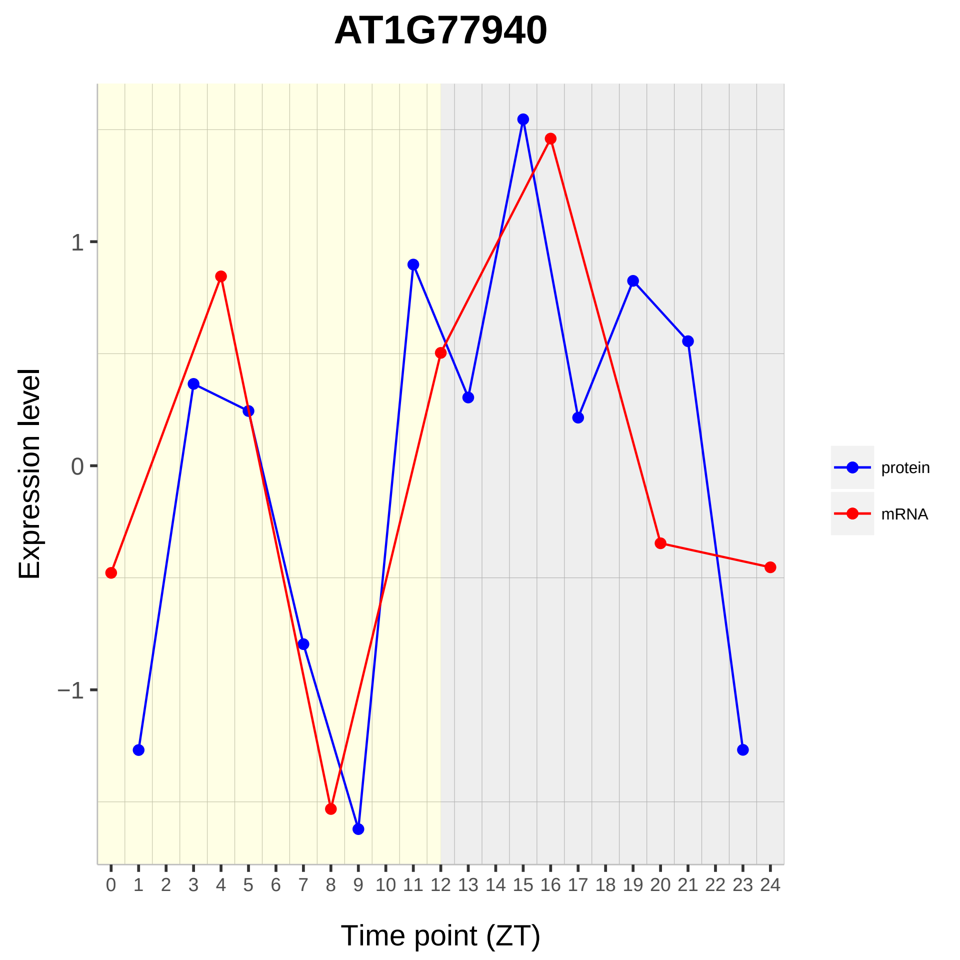
AT1G77940
Expression level
1
0
−1
0 1 2 3 4 5 6 7 8 9 10 11 12 13 14 15 16 17 18 19 20 21 22 23 24
Time point (ZT)
protein
mRNA

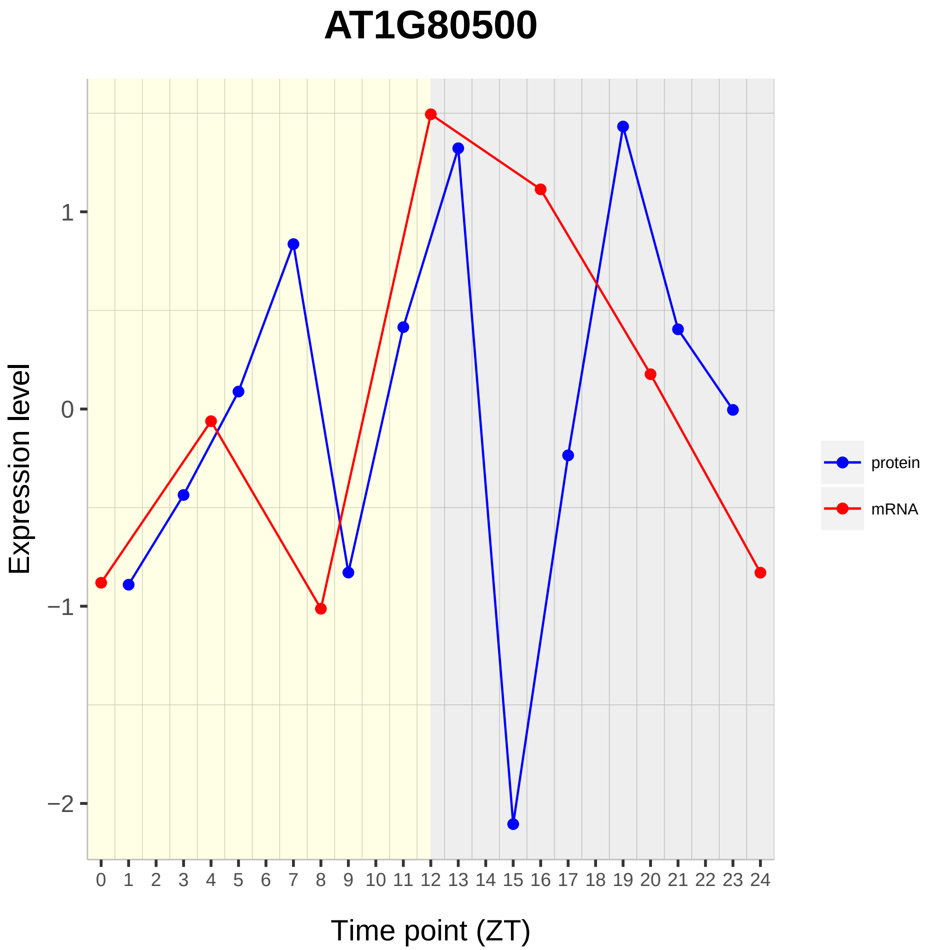
AT1G80500
Expression level
Time point (ZT)
1
0
−1
−2
0 1 2 3 4 5 6 7 8 9 10 11 12 13 14 15 16 17 18 19 20 21 22 23 24
protein
mRNA

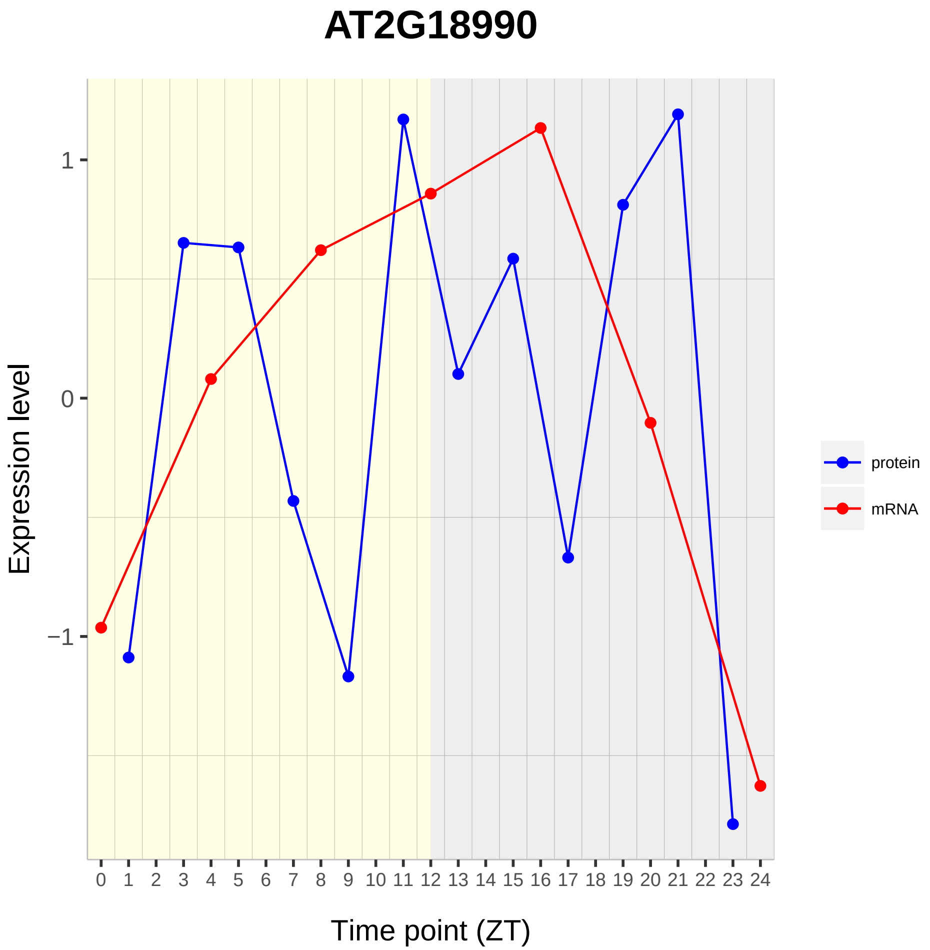

AT2G18990
Expression level
Time point (ZT)
protein
mRNA
1
0
−1

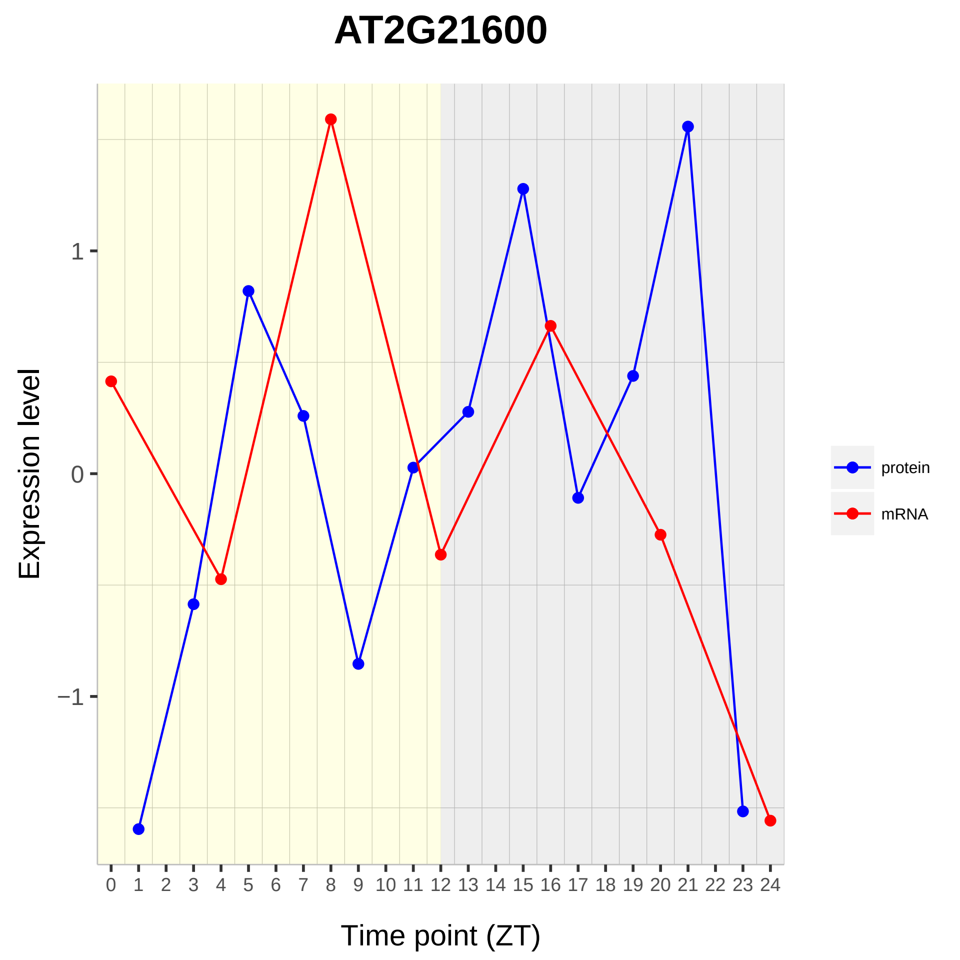
AT2G21600
Expression level
1
0
−1
Time point (ZT)
protein
mRNA

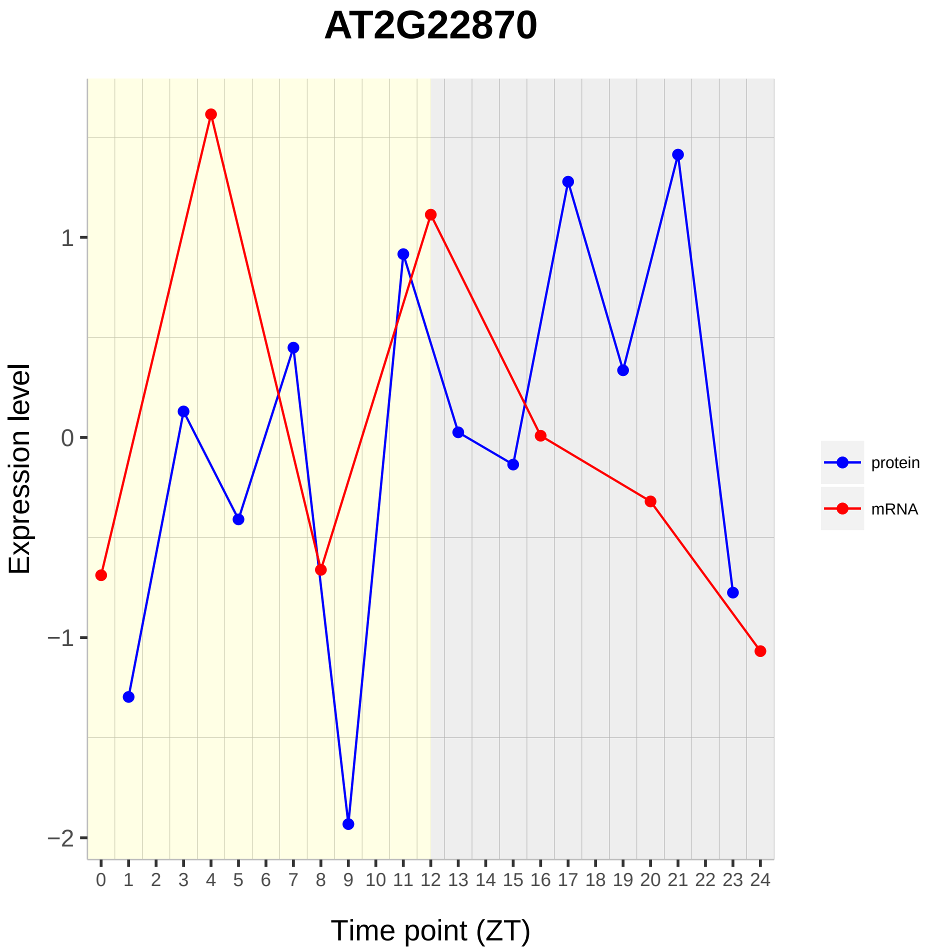

AT2G22870
Expression level
1
0
−1
−2
0 1 2 3 4 5 6 7 8 9 10 11 12 13 14 15 16 17 18 19 20 21 22 23 24
Time point (ZT)
protein
mRNA

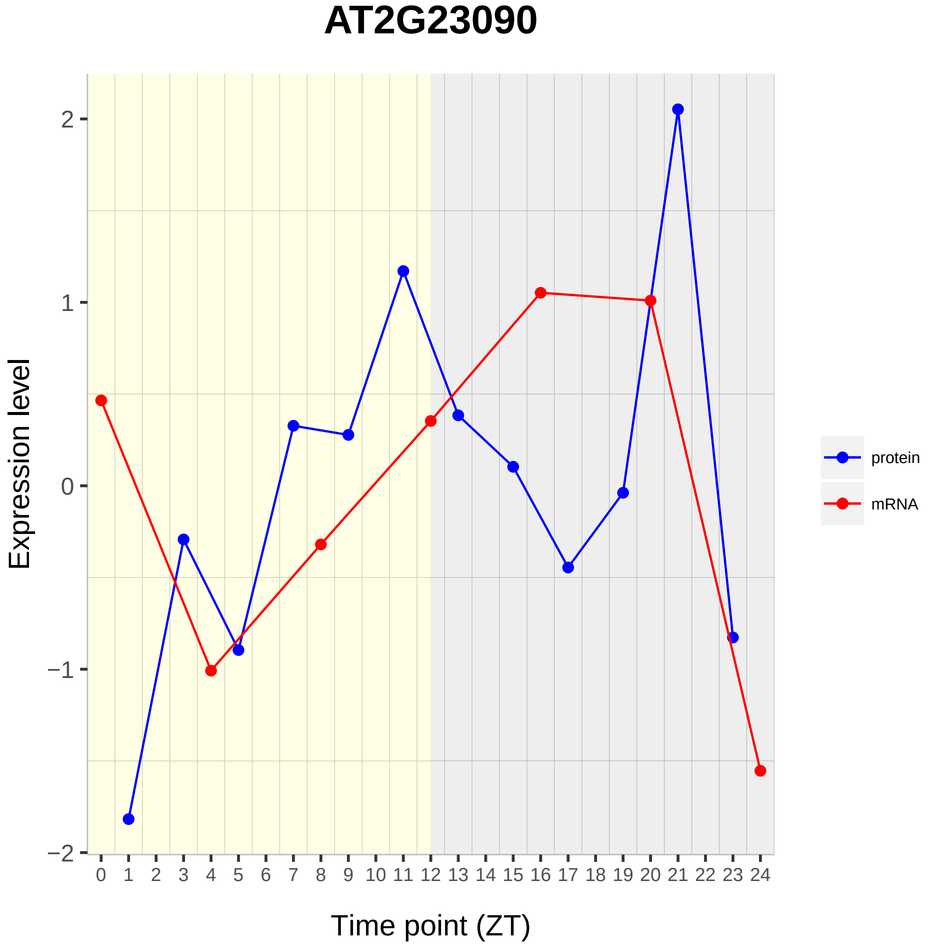

AT2G23090
Expression level
Time point (ZT)
2
1
0
−1
−2
0 1 2 3 4 5 6 7 8 9 10 11 12 13 14 15 16 17 18 19 20 21 22 23 24
protein
mRNA

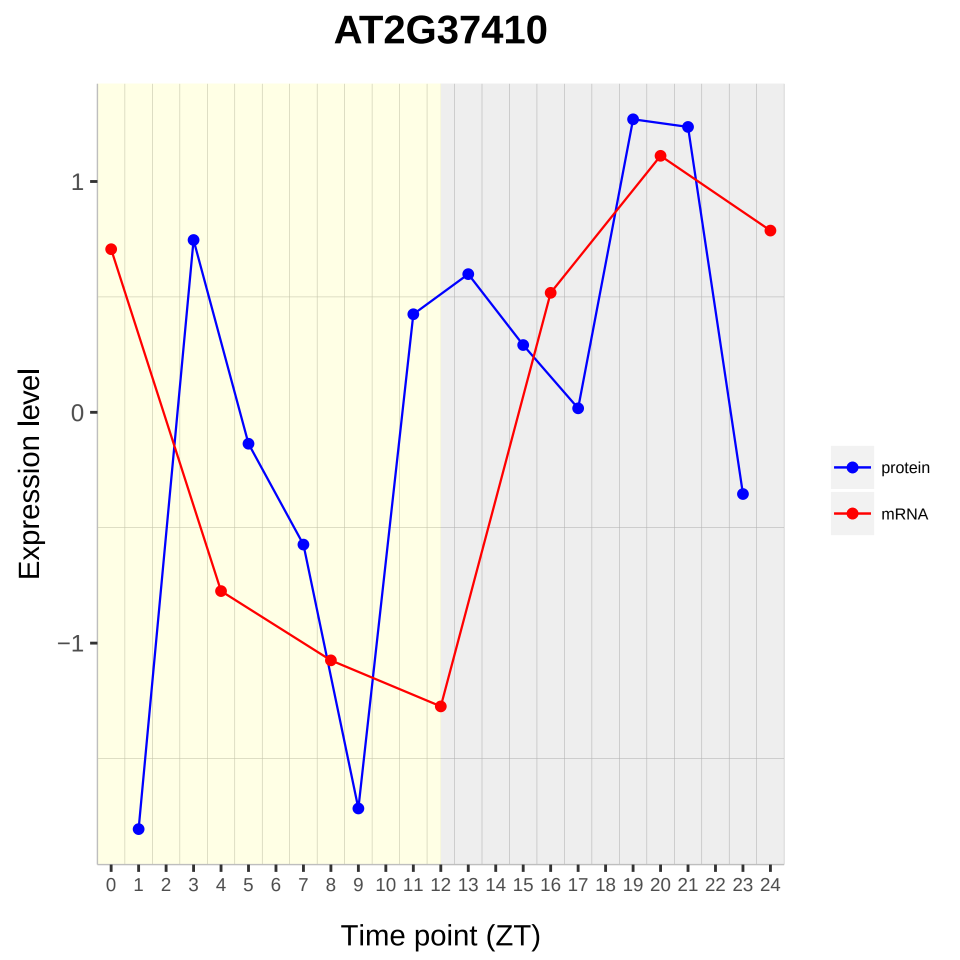

AT2G37410
Expression level
1
0
−1
Time point (ZT)
protein
mRNA

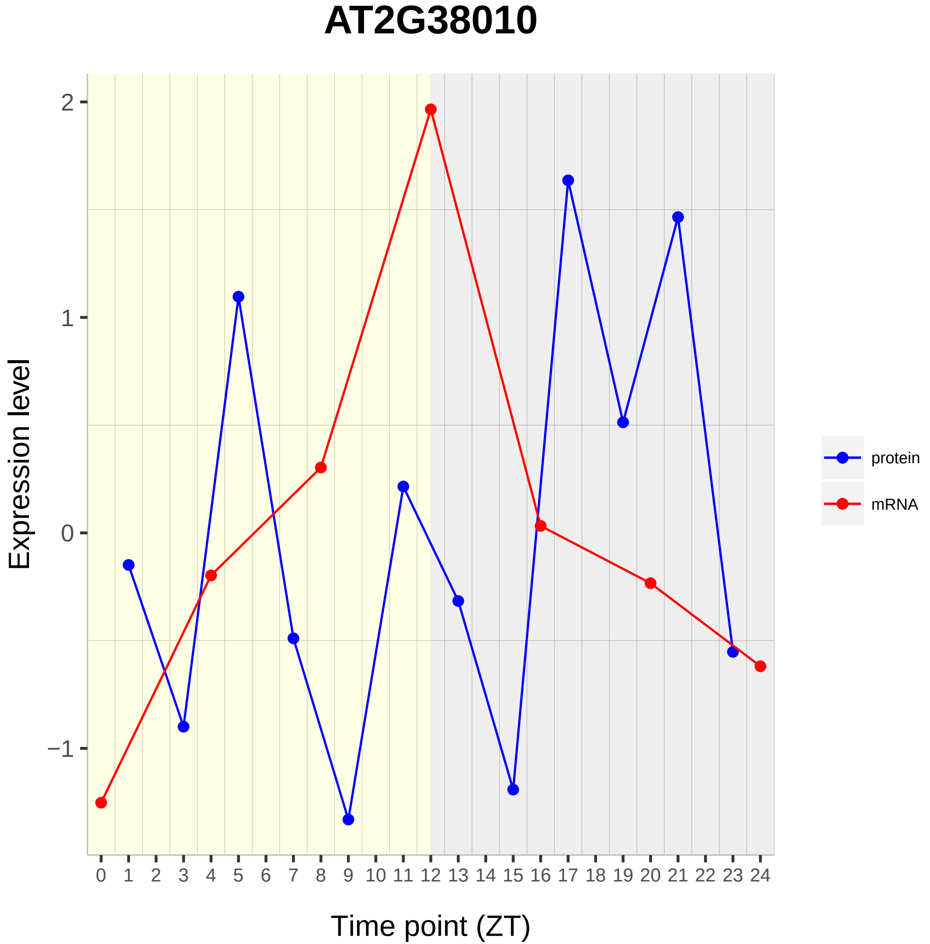

AT2G38010
Expression level
2
1
0
−1
0 1 2 3 4 5 6 7 8 9 10 11 12 13 14 15 16 17 18 19 20 21 22 23 24
Time point (ZT)
protein
mRNA

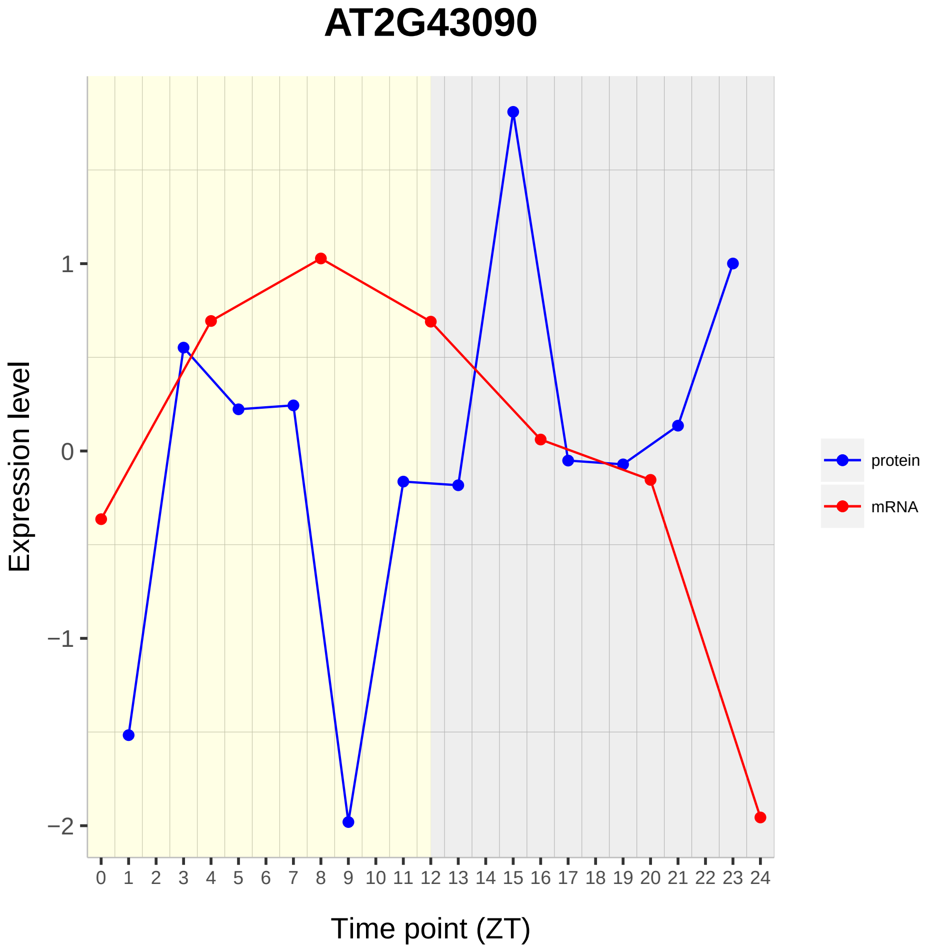
AT2G43090
Expression level
Time point (ZT)
1
0
−1
−2
0 1 2 3 4 5 6 7 8 9 10 11 12 13 14 15 16 17 18 19 20 21 22 23 24
protein
mRNA

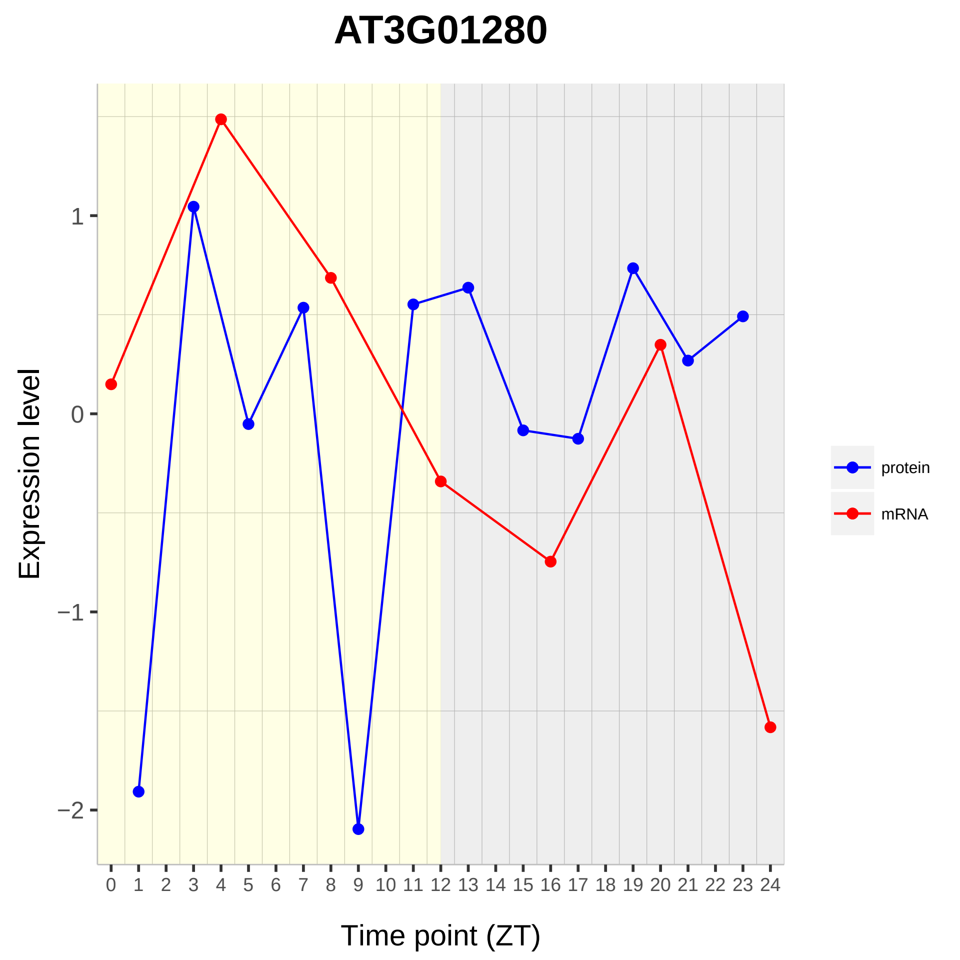

AT3G01280
Expression level
1
0
−1
−2
0 1 2 3 4 5 6 7 8 9 10 11 12 13 14 15 16 17 18 19 20 21 22 23 24
Time point (ZT)
protein
mRNA

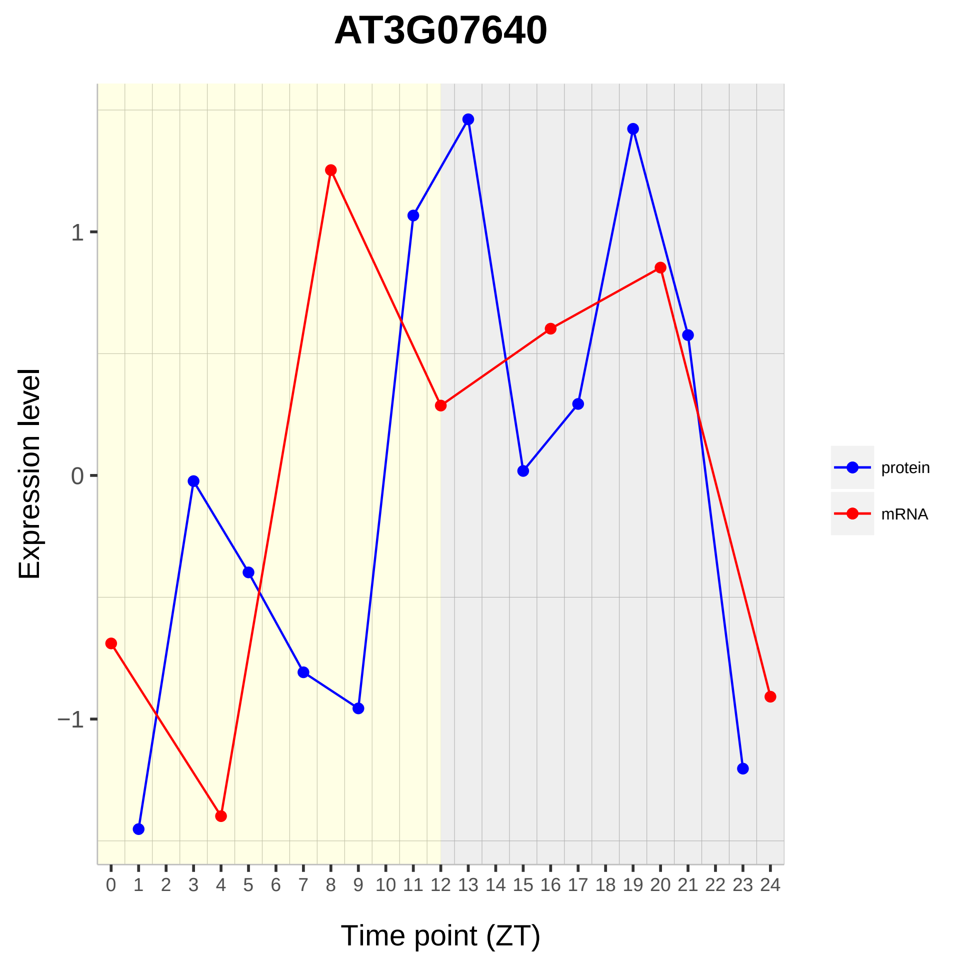
AT3G07640
Expression level
1
0
−1
Time point (ZT)
protein
mRNA

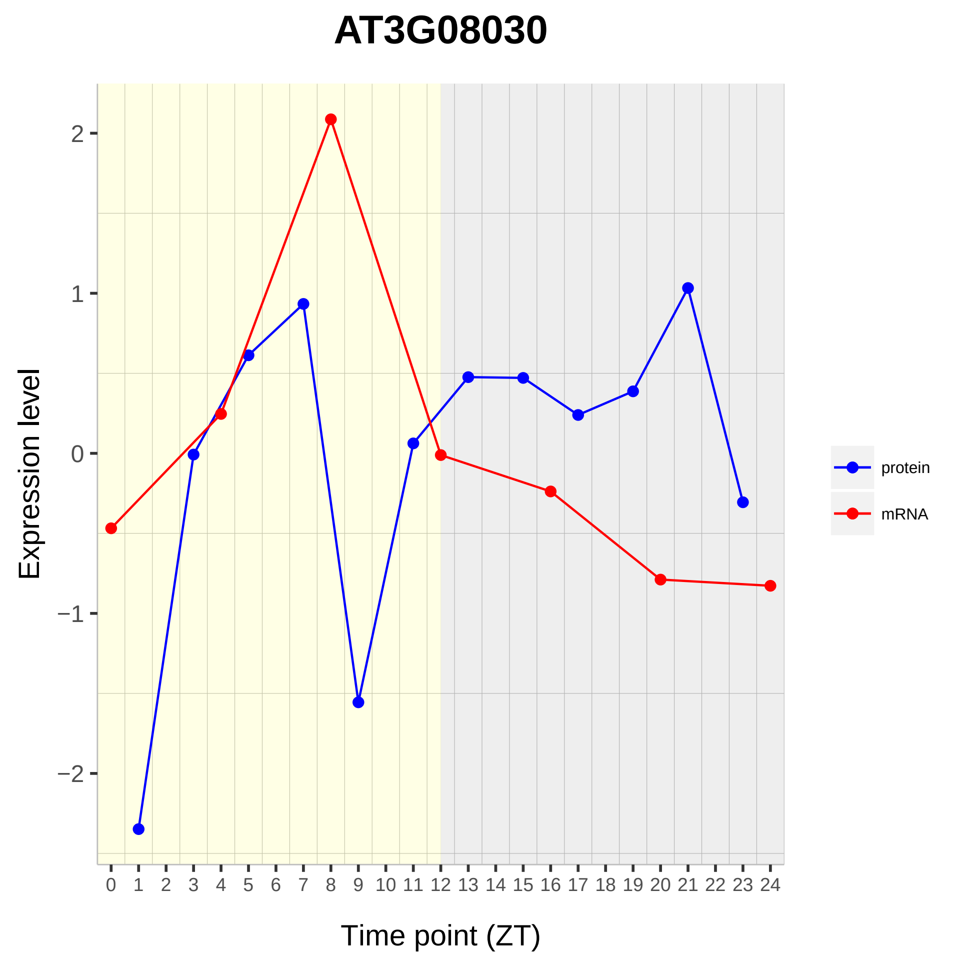

AT3G08030
Expression level
Time point (ZT)
2
1
0
−1
−2
protein
mRNA

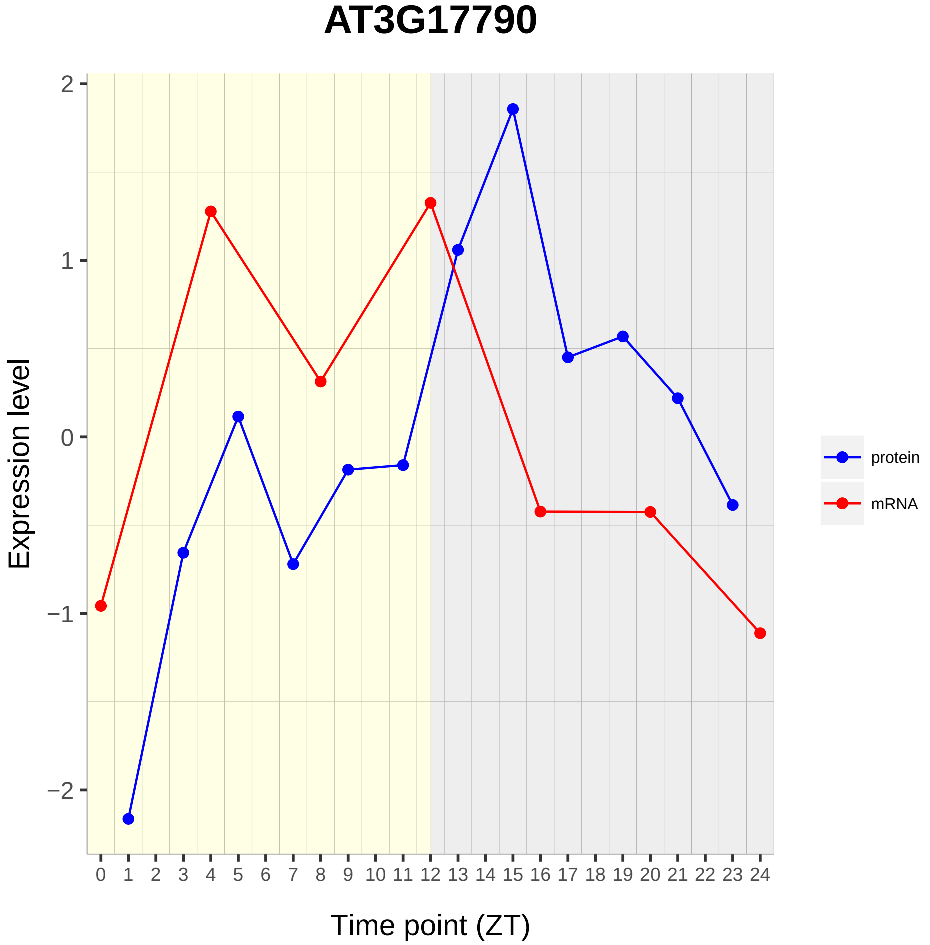
AT3G17790
Expression level
Time point (ZT)
protein
mRNA
2
1
0
−1
−2
0 1 2 3 4 5 6 7 8 9 10 11 12 13 14 15 16 17 18 19 20 21 22 23 24

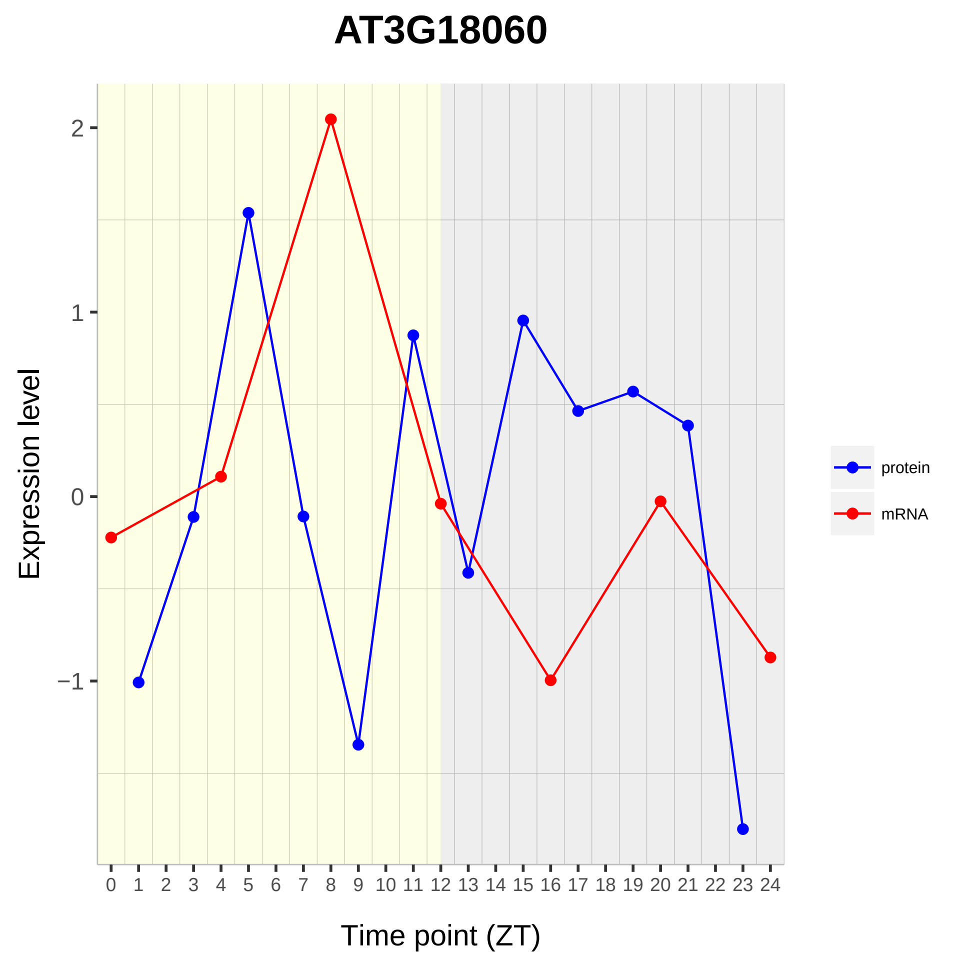
AT3G18060
Expression level
Time point (ZT)
2
1
0
−1
0 1 2 3 4 5 6 7 8 9 10 11 12 13 14 15 16 17 18 19 20 21 22 23 24
protein
mRNA

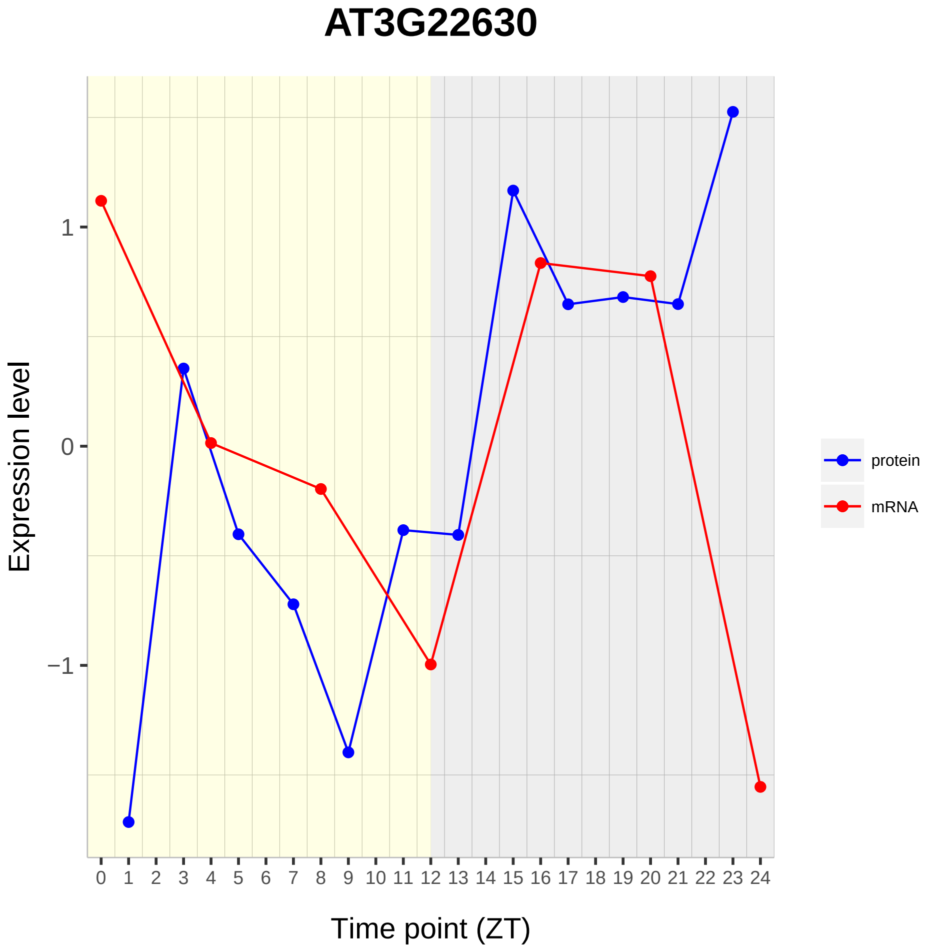

AT3G22630
Expression level
1
0
−1
Time point (ZT)
protein
mRNA

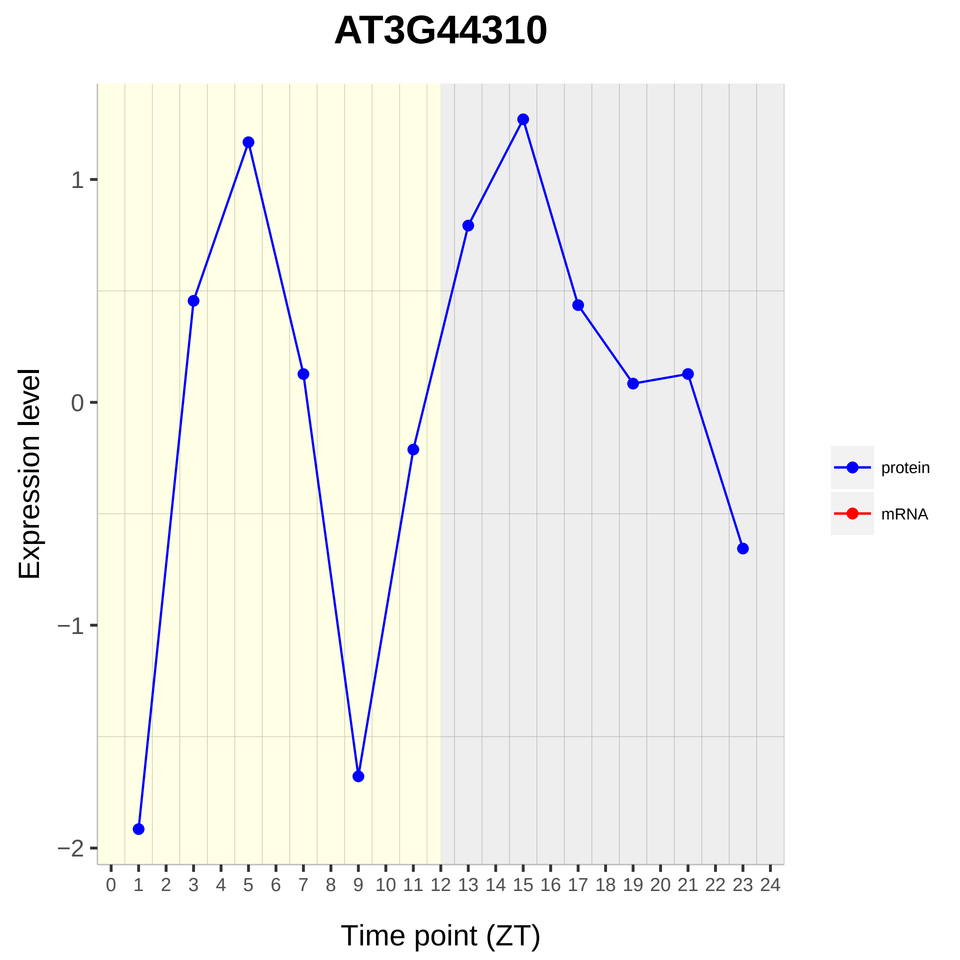

AT3G44310
Expression level
1
0
−1
−2
0 1 2 3 4 5 6 7 8 9 10 11 12 13 14 15 16 17 18 19 20 21 22 23 24
Time point (ZT)
protein
mRNA

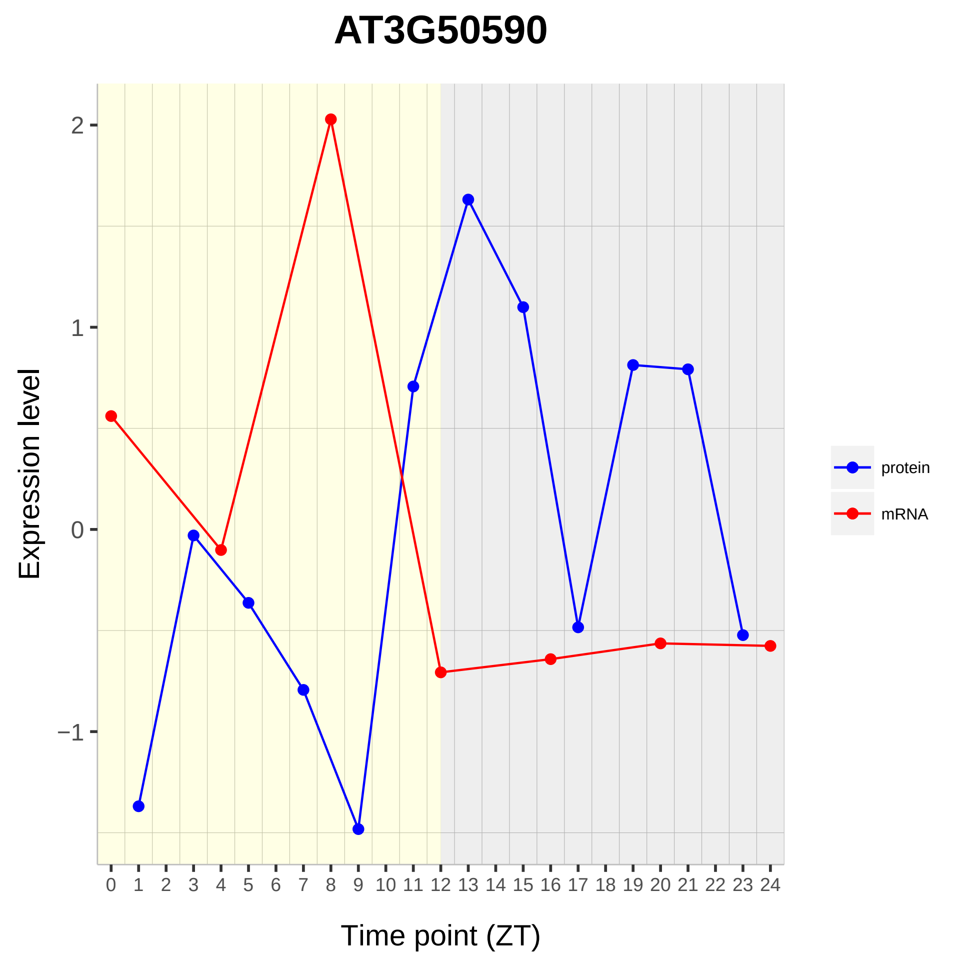
AT3G50590
Expression level
Time point (ZT)
2
1
0
−1
0 1 2 3 4 5 6 7 8 9 10 11 12 13 14 15 16 17 18 19 20 21 22 23 24
protein
mRNA

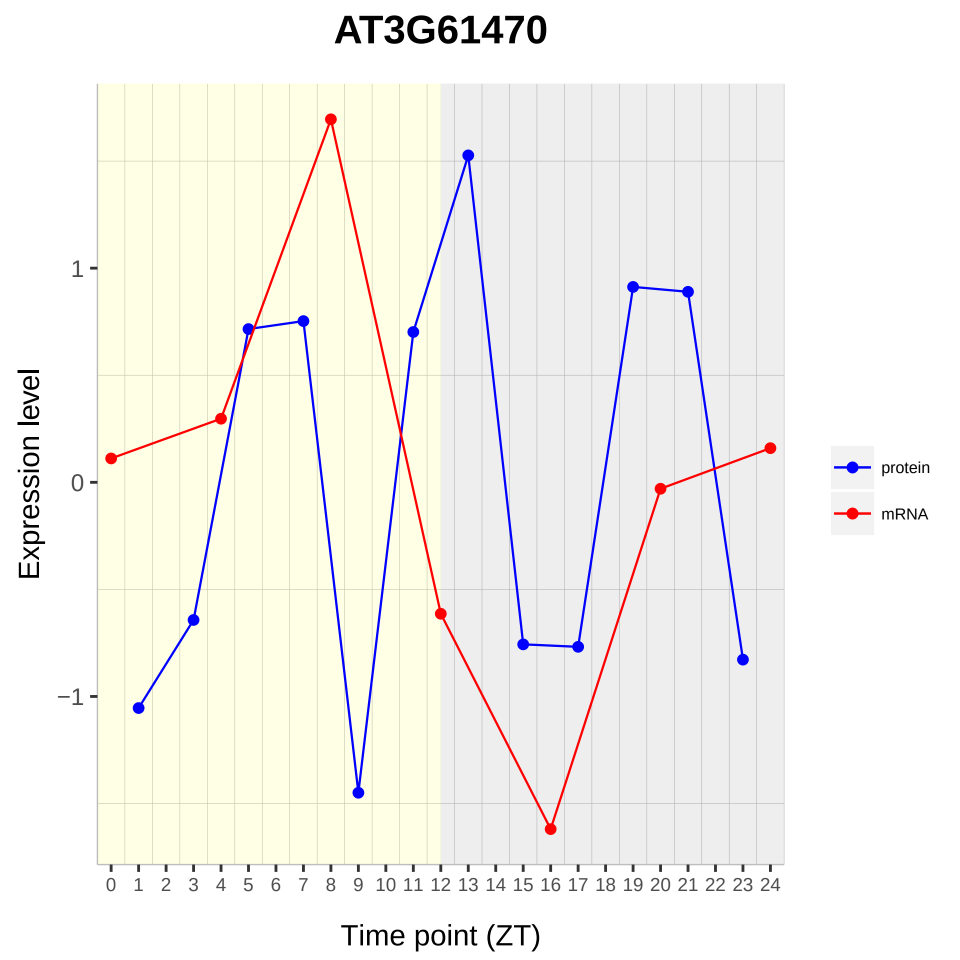

AT3G61470
Expression level
1
0
−1
Time point (ZT)
protein
mRNA

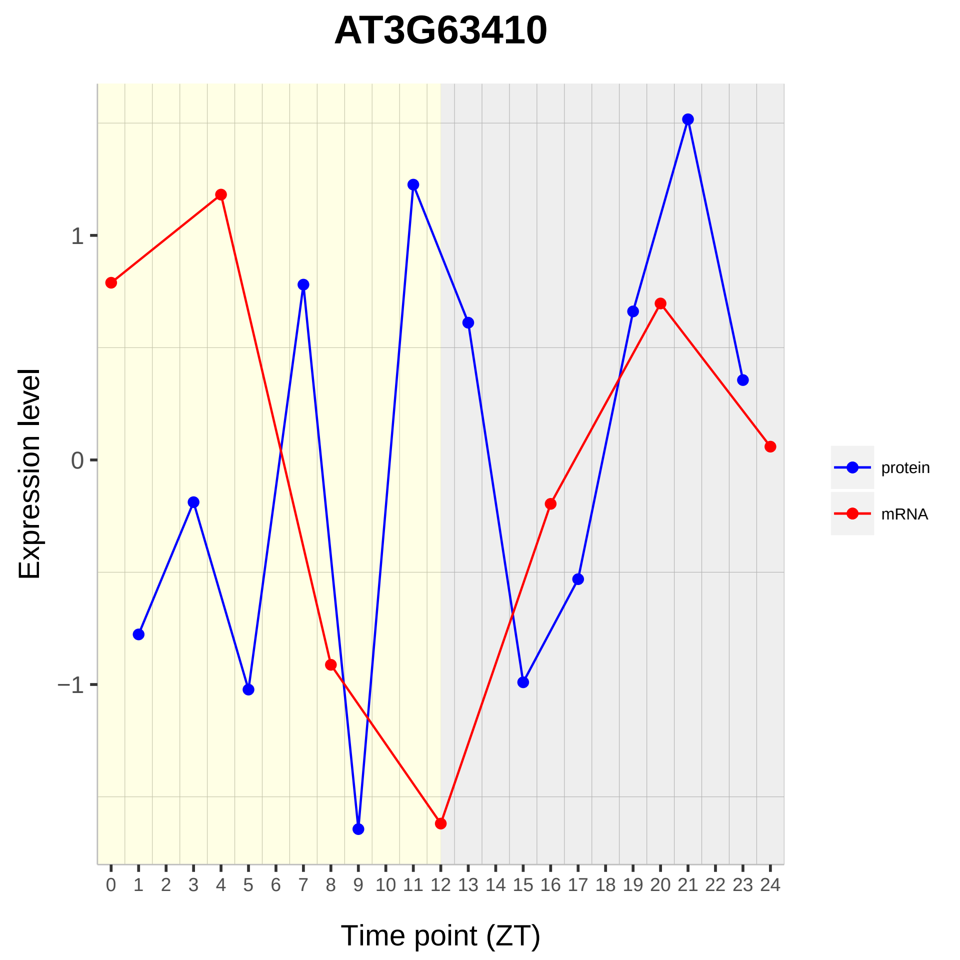

AT3G63410
Expression level
1
0
−1
Time point (ZT)
protein
mRNA

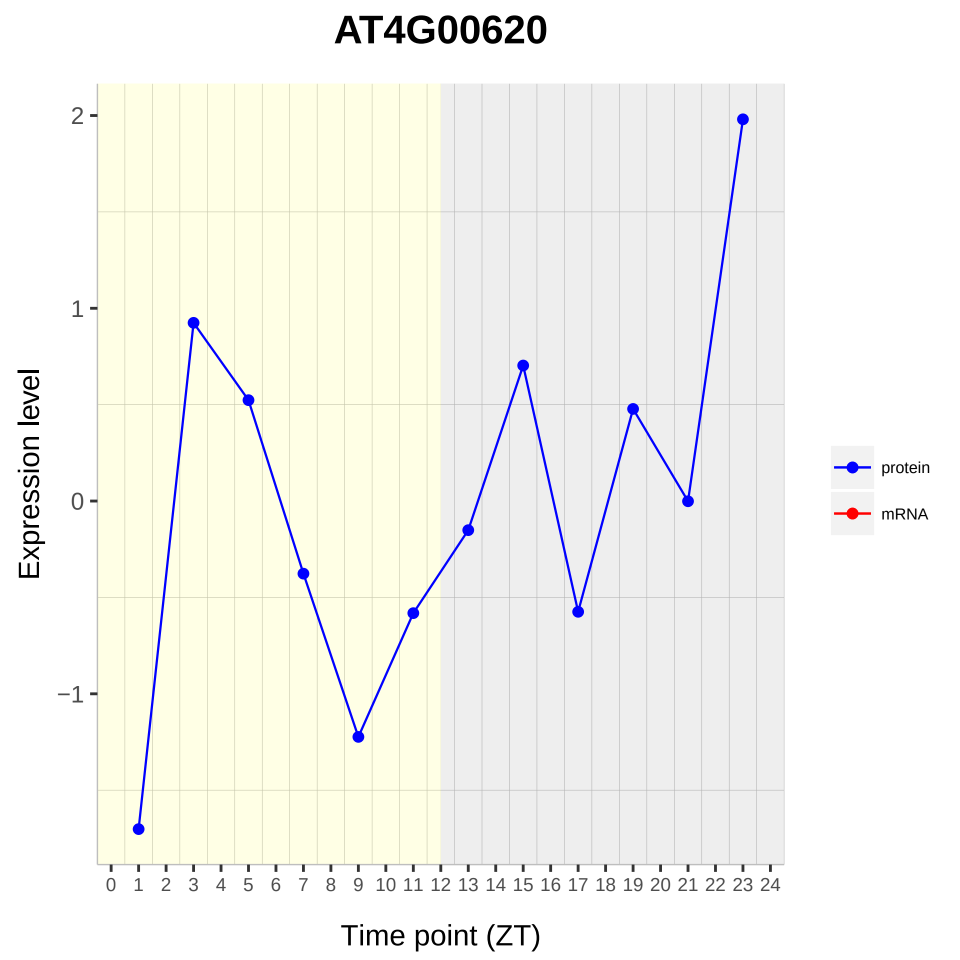
AT4G00620
Expression level
2
1
0
−1
0 1 2 3 4 5 6 7 8 9 10 11 12 13 14 15 16 17 18 19 20 21 22 23 24
Time point (ZT)
protein
mRNA

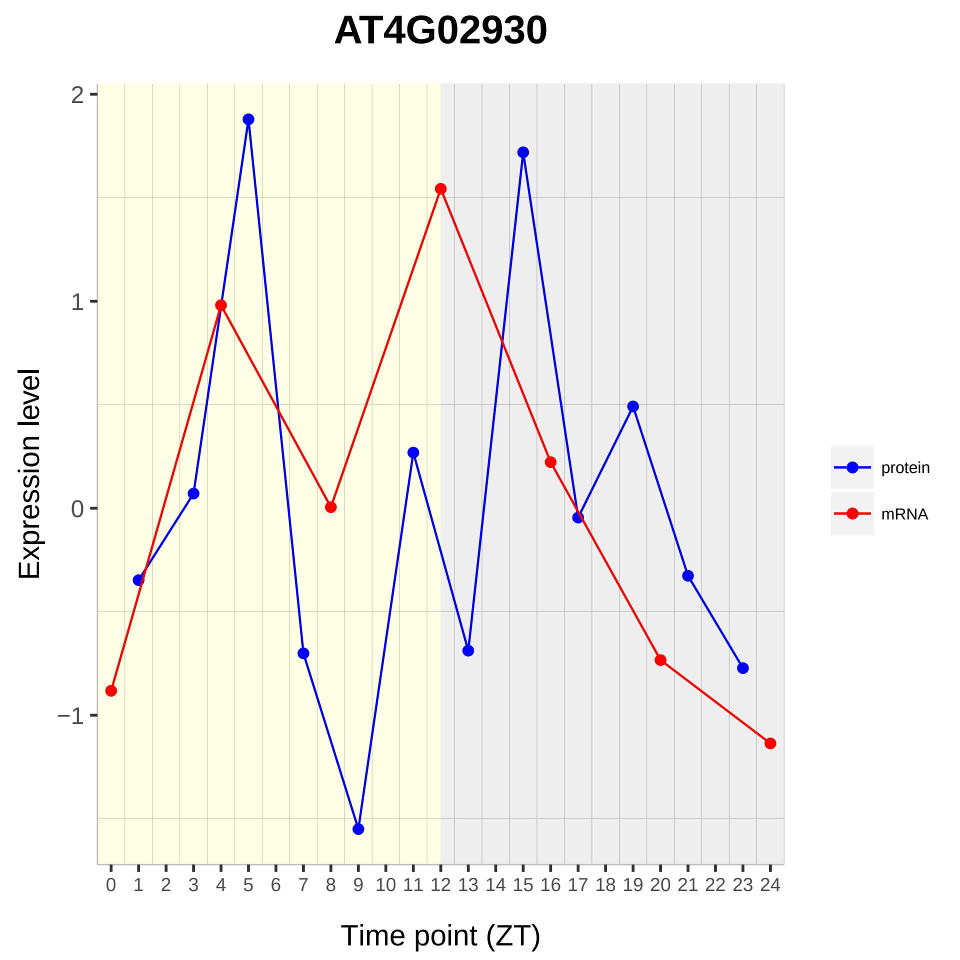

AT4G02930
Expression level
Time point (ZT)
protein
mRNA
2
1
0
−1

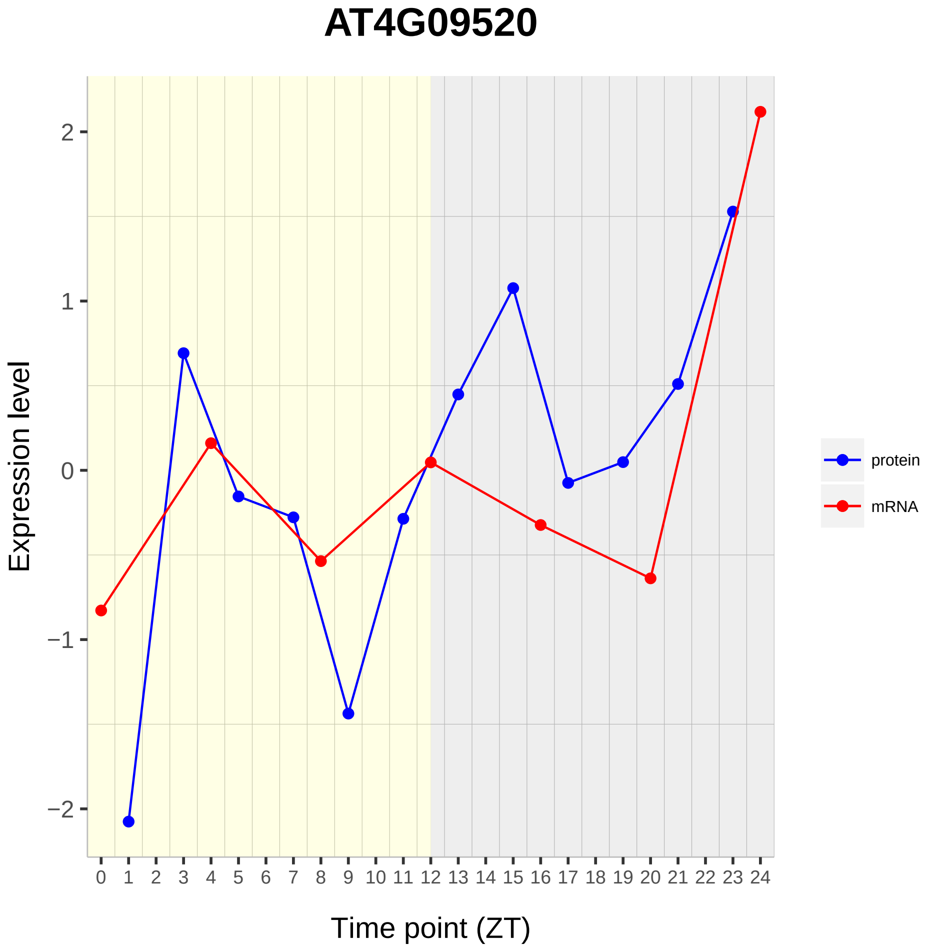
AT4G09520
Expression level
Time point (ZT)
2
1
0
−1
−2
protein
mRNA

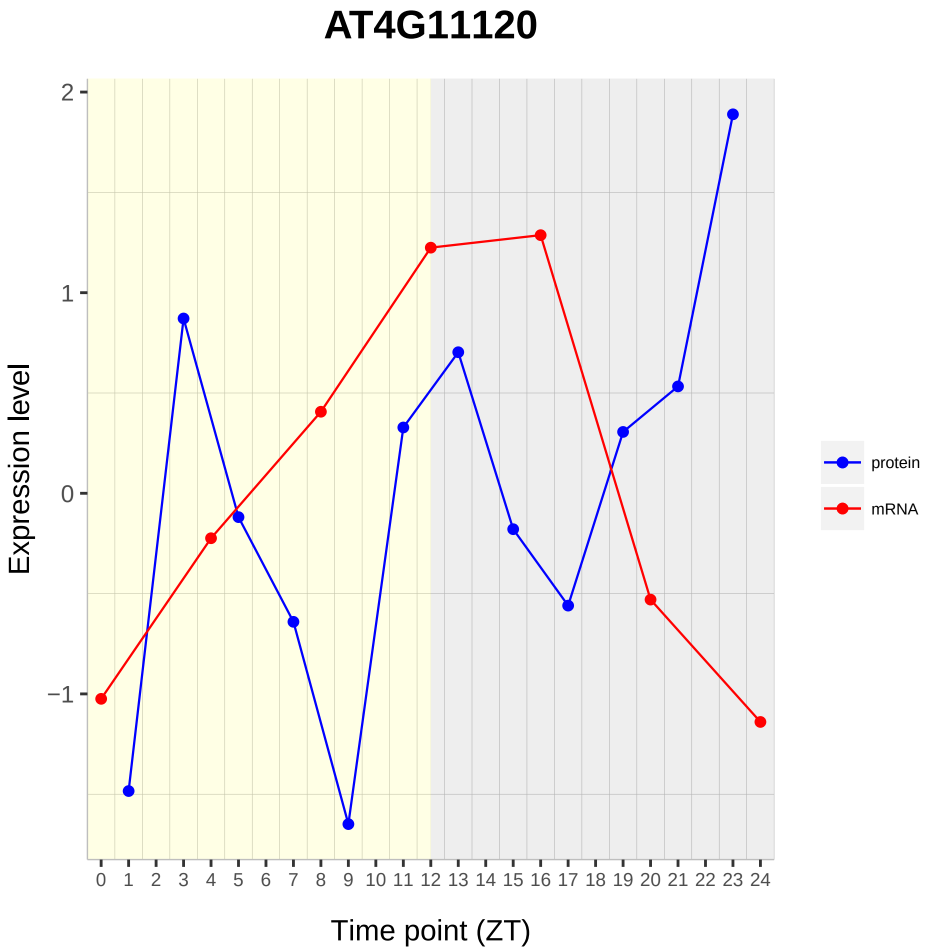
AT4G11120
Expression level
Time point (ZT)
2
1
0
−1
0 1 2 3 4 5 6 7 8 9 10 11 12 13 14 15 16 17 18 19 20 21 22 23 24
protein
mRNA

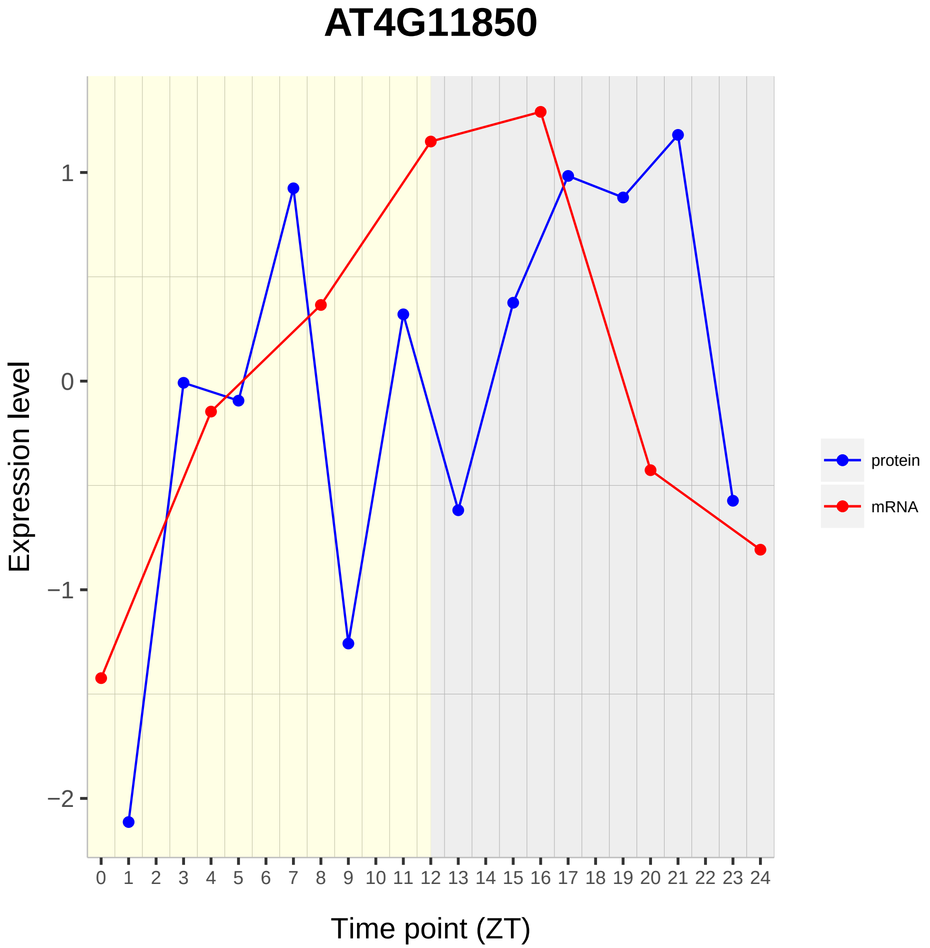

AT4G11850
Expression level
1
0
−1
−2
Time point (ZT)
protein
mRNA

# AT4G18465

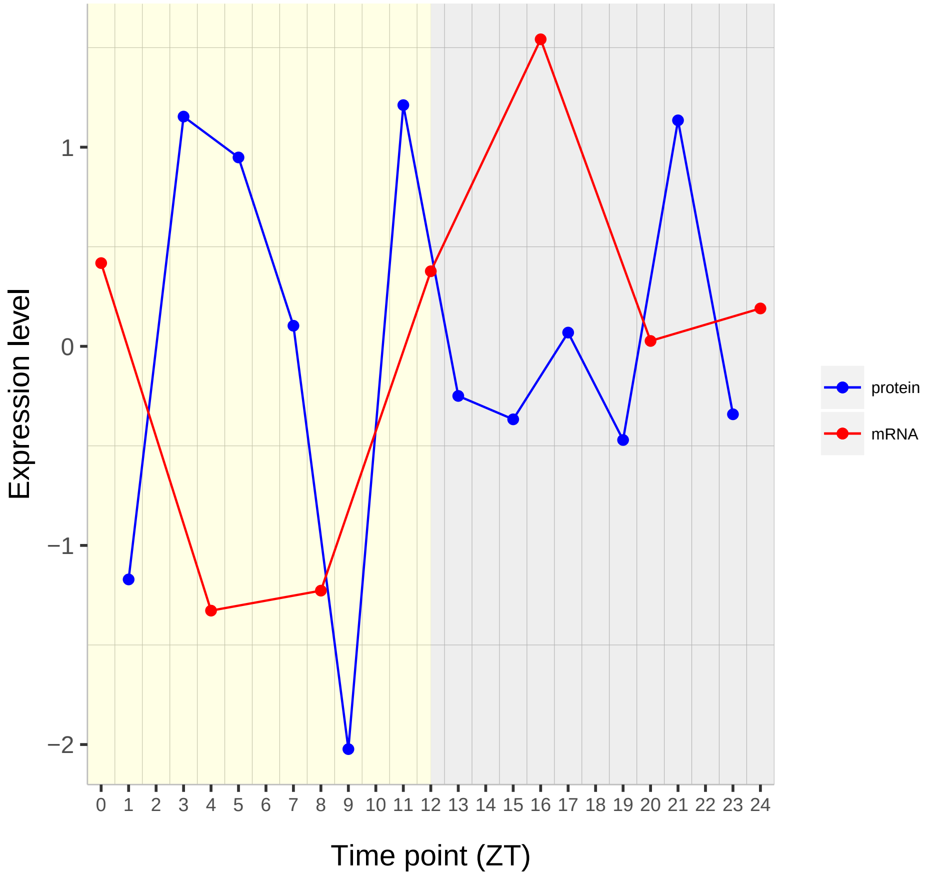

Expression level
1
0
−1
−2
Time point (ZT)
protein
mRNA

# AT4G33945

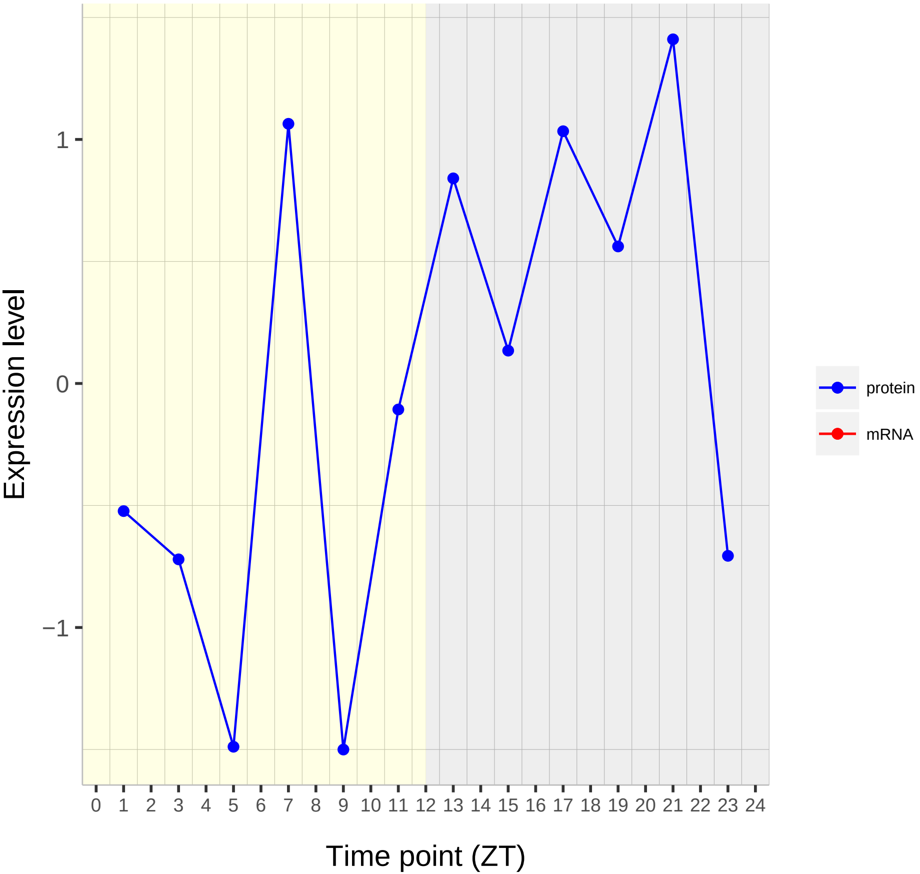

Expression level
1
0
−1
0 1 2 3 4 5 6 7 8 9 10 11 12 13 14 15 16 17 18 19 20 21 22 23 24
Time point (ZT)
protein
mRNA

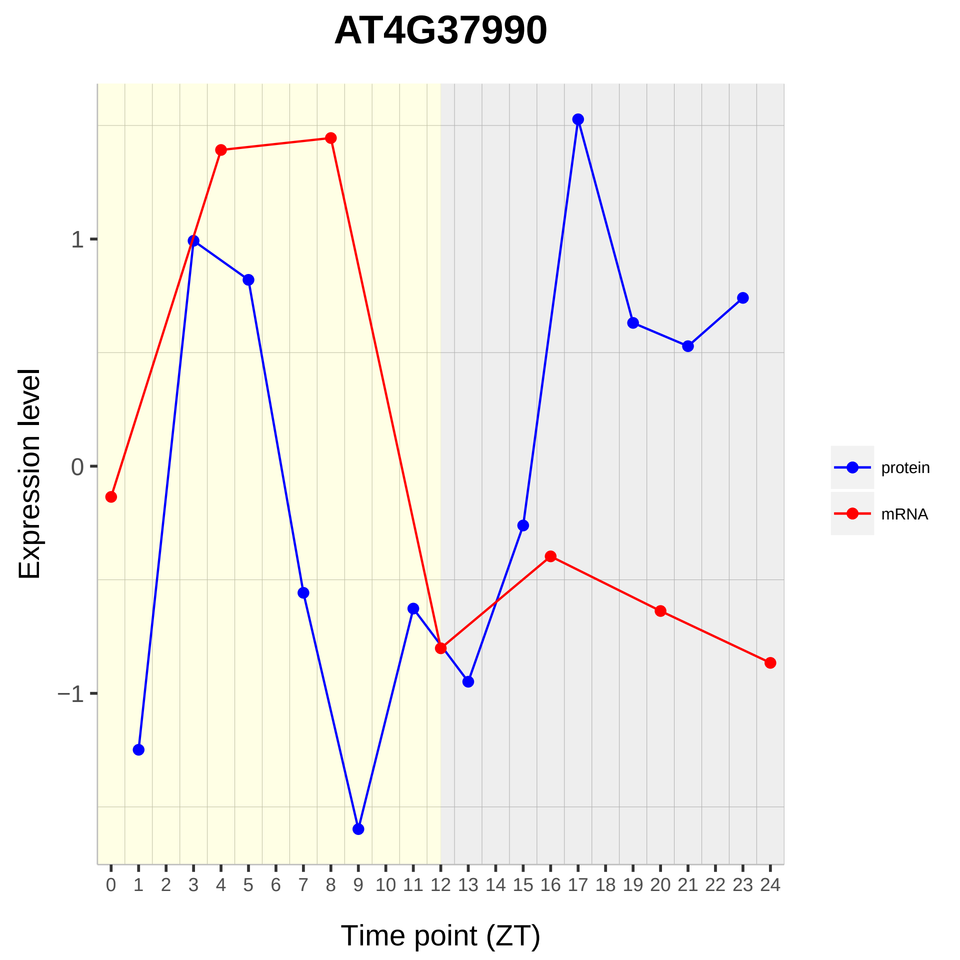

AT4G37990
Expression level
1
0
−1
Time point (ZT)
protein
mRNA

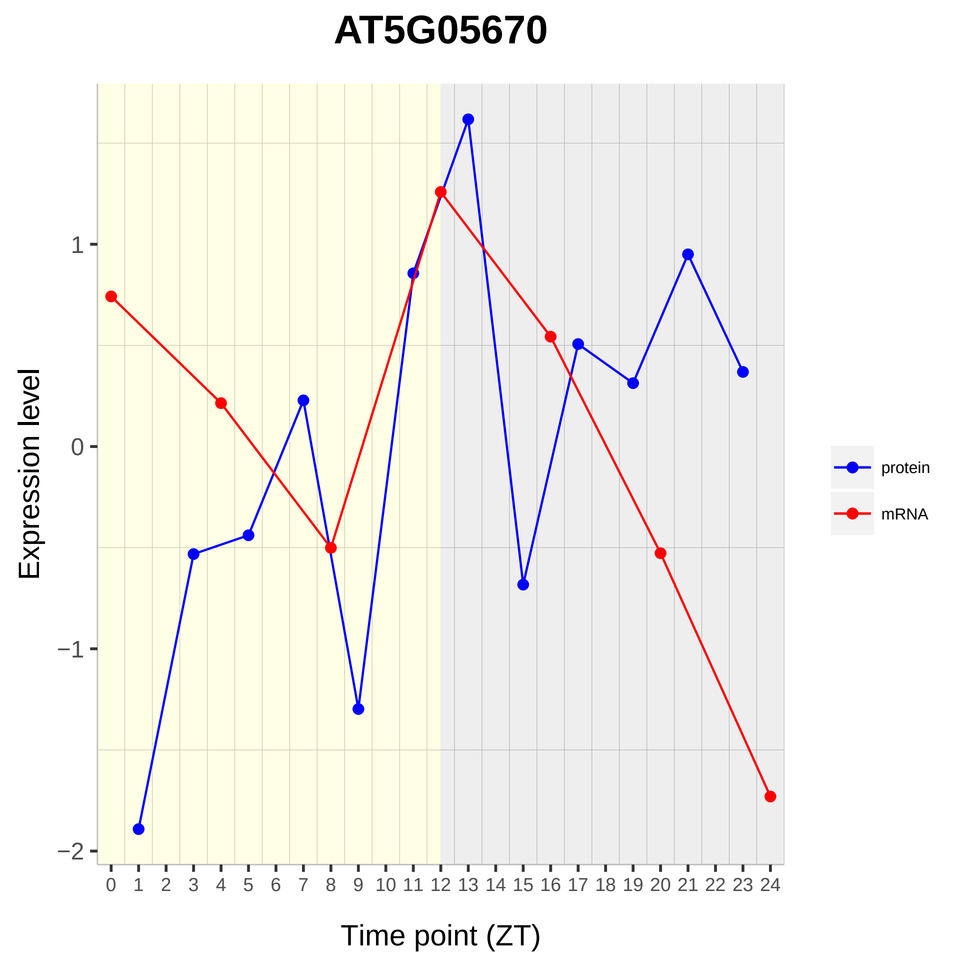

AT5G05670
Expression level
1
0
−1
−2
0 1 2 3 4 5 6 7 8 9 10 11 12 13 14 15 16 17 18 19 20 21 22 23 24
Time point (ZT)
protein
mRNA

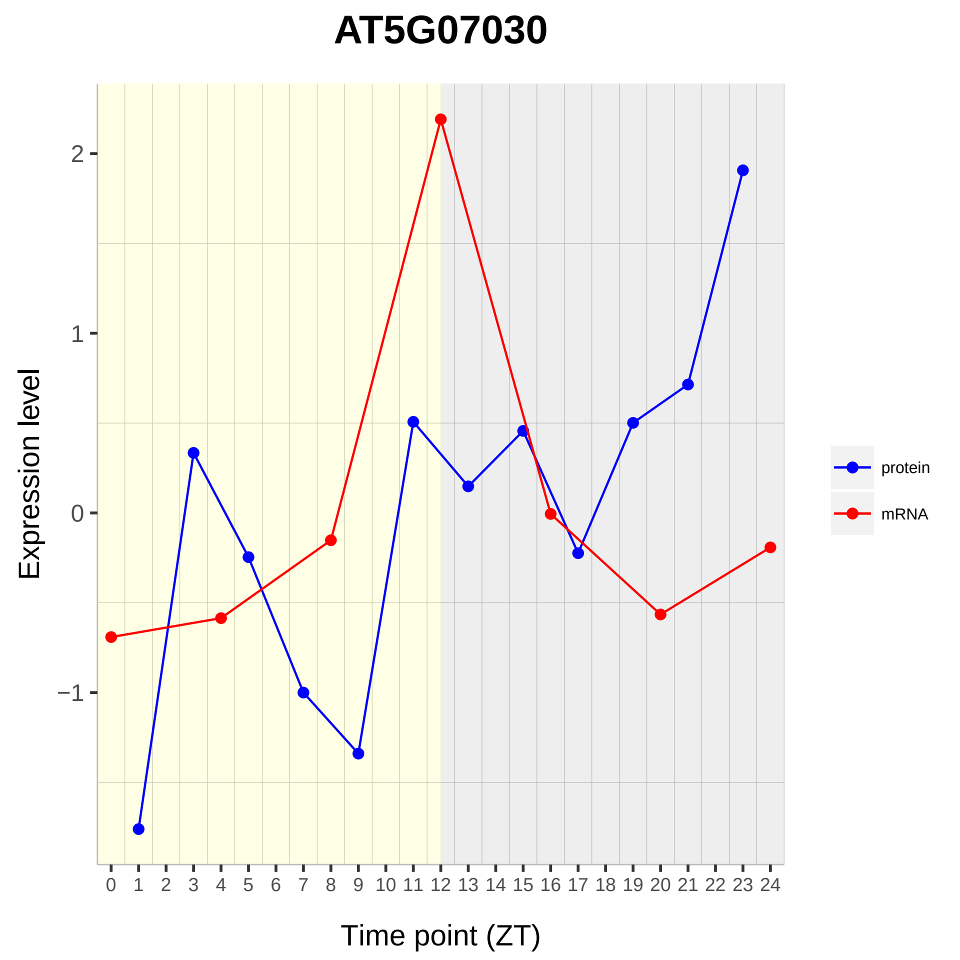

AT5G07030
Expression level
2
1
0
−1
Time point (ZT)
protein
mRNA

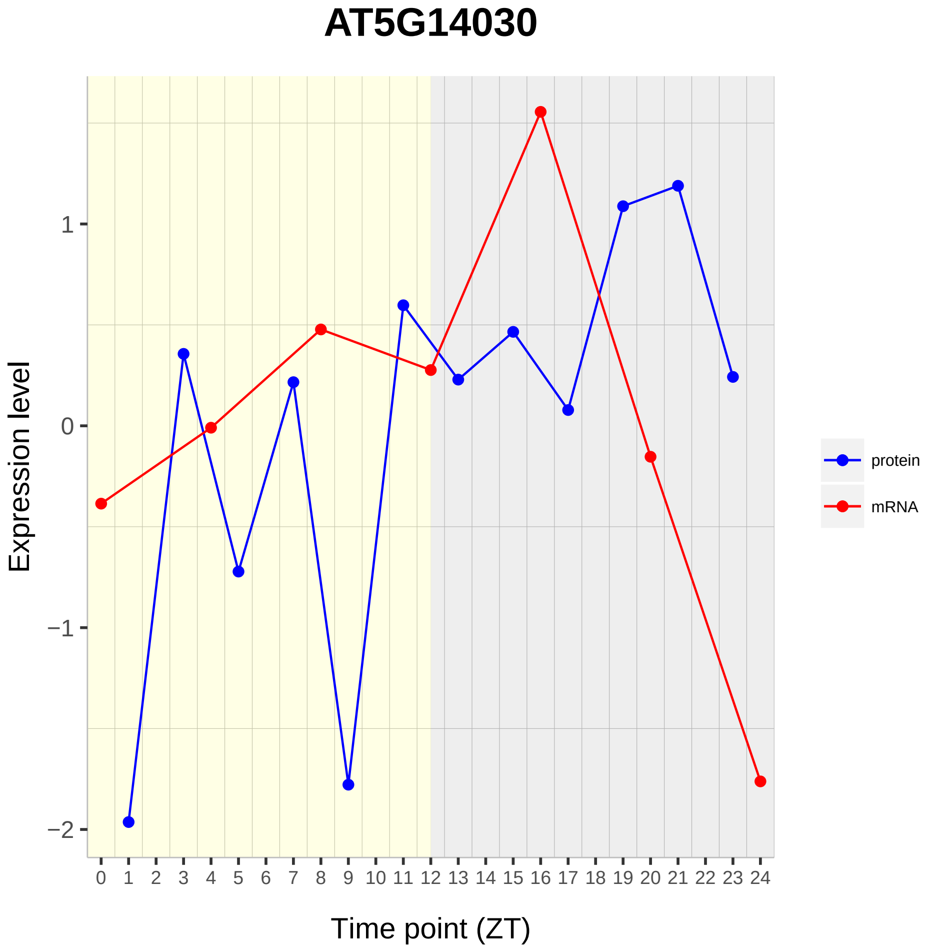
AT5G14030
Expression level
1
0
−1
−2
Time point (ZT)
protein
mRNA

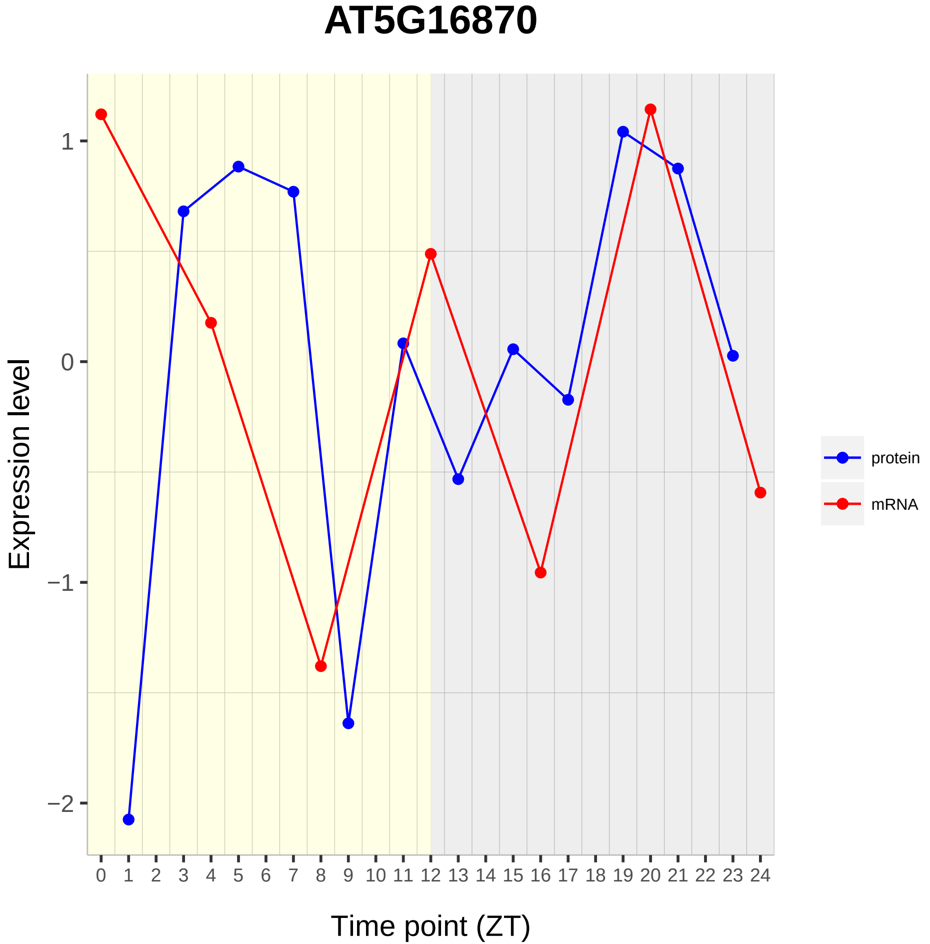
AT5G16870
Expression level
Time point (ZT)
1
0
−1
−2
protein
mRNA

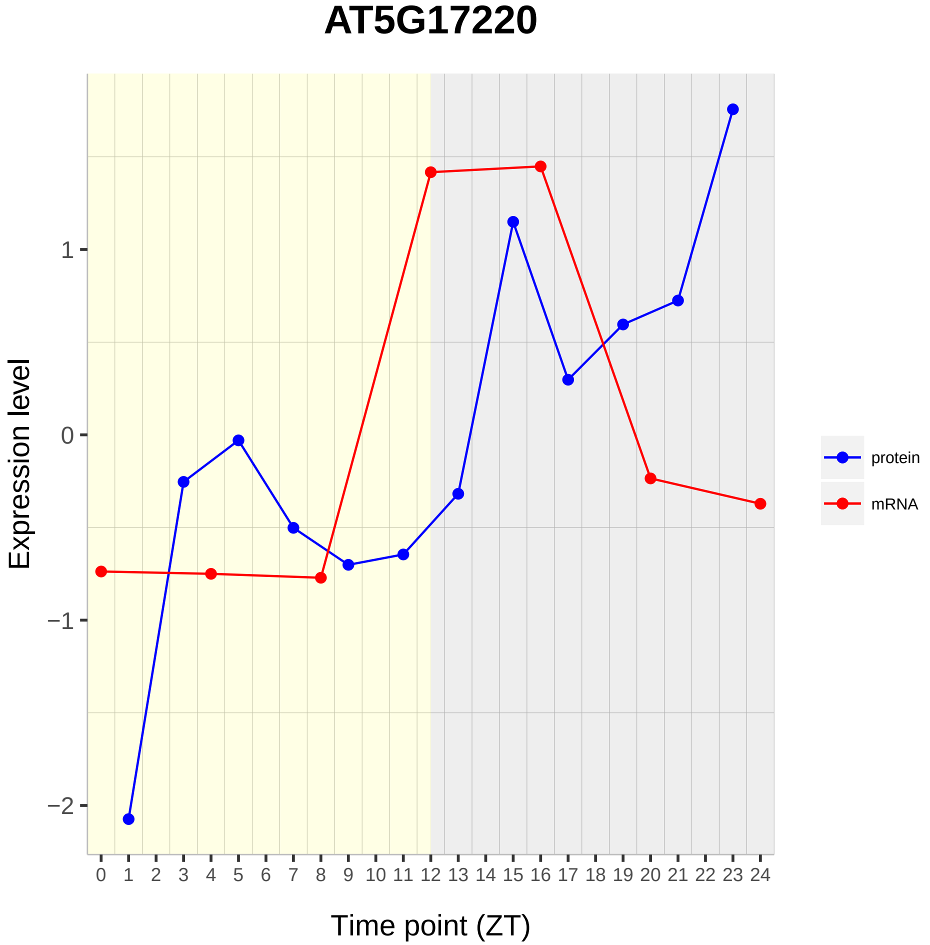
AT5G17220
Expression level
1
0
−1
−2
0 1 2 3 4 5 6 7 8 9 10 11 12 13 14 15 16 17 18 19 20 21 22 23 24
Time point (ZT)
protein
mRNA

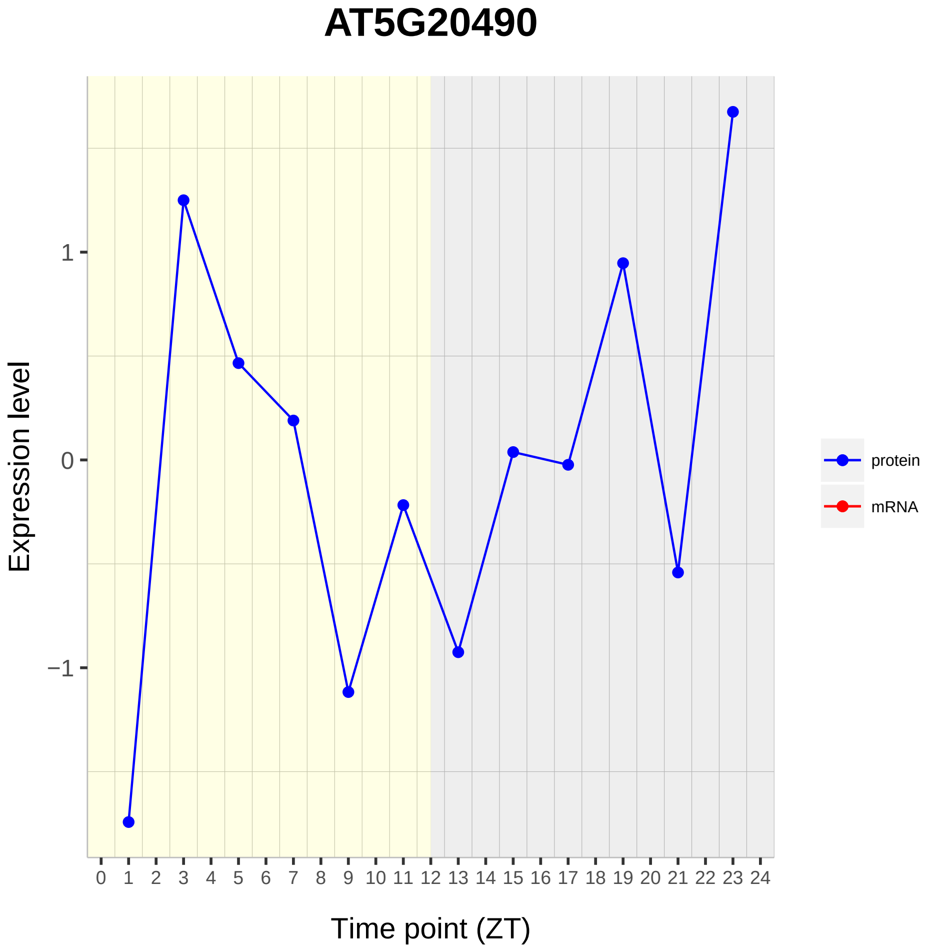
AT5G20490
Expression level
1
0
−1
0 1 2 3 4 5 6 7 8 9 10 11 12 13 14 15 16 17 18 19 20 21 22 23 24
Time point (ZT)
protein
mRNA

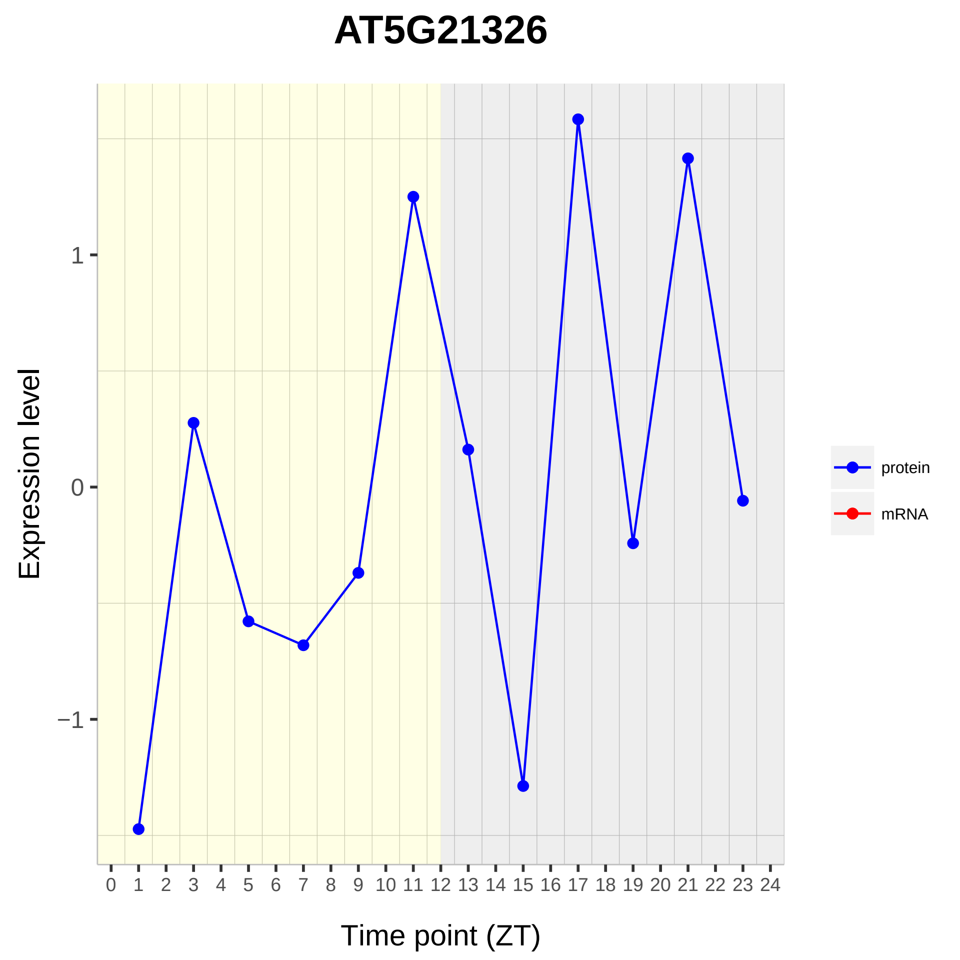

AT5G21326
Expression level
1
0
−1
0 1 2 3 4 5 6 7 8 9 10 11 12 13 14 15 16 17 18 19 20 21 22 23 24
Time point (ZT)
protein
mRNA

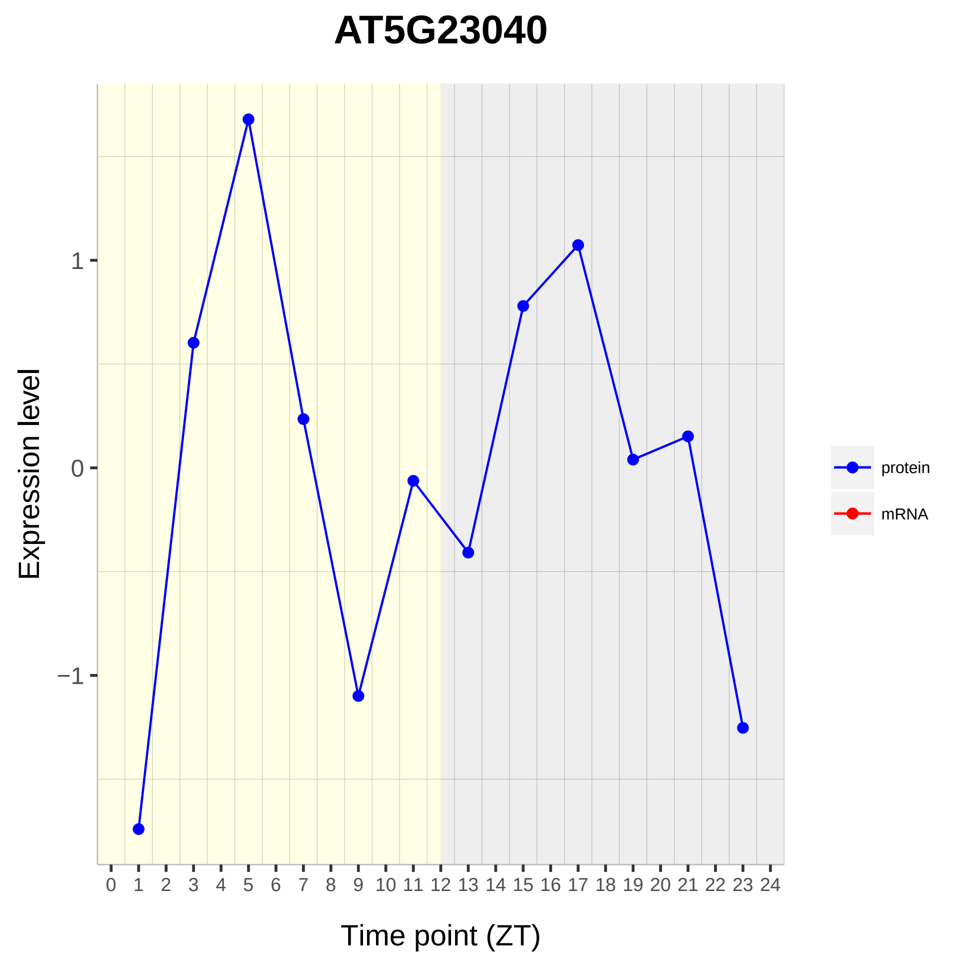

AT5G23040
Expression level
1
0
−1
0 1 2 3 4 5 6 7 8 9 10 11 12 13 14 15 16 17 18 19 20 21 22 23 24
Time point (ZT)
protein
mRNA

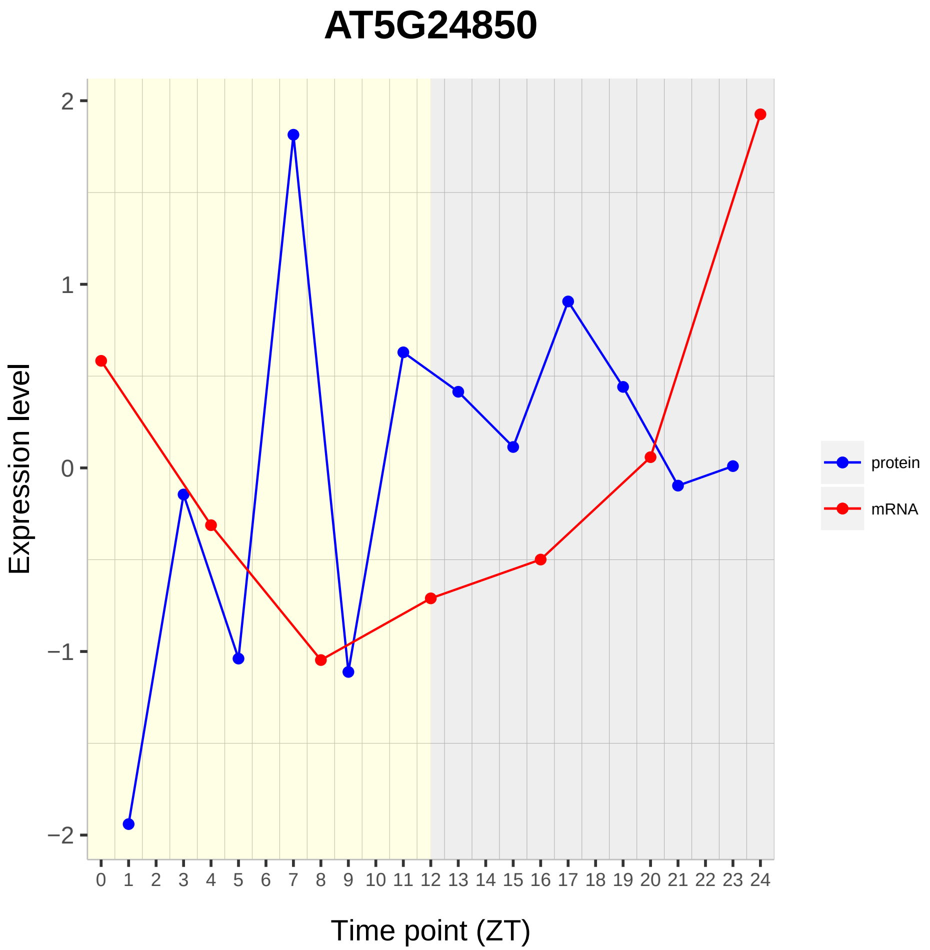
AT5G24850
Expression level
Time point (ZT)
2
1
0
−1
−2
0 1 2 3 4 5 6 7 8 9 10 11 12 13 14 15 16 17 18 19 20 21 22 23 24
protein
mRNA

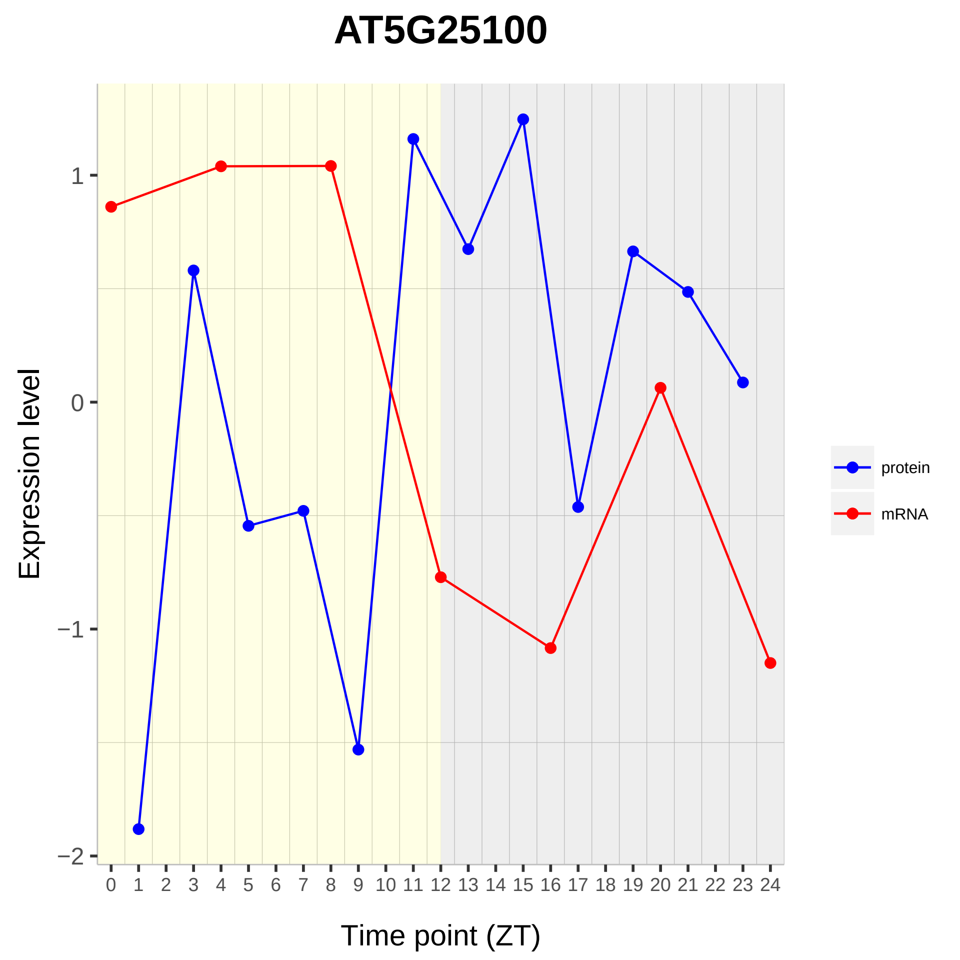

AT5G25100
Expression level
Time point (ZT)
1
0
−1
−2
0 1 2 3 4 5 6 7 8 9 10 11 12 13 14 15 16 17 18 19 20 21 22 23 24
protein
mRNA

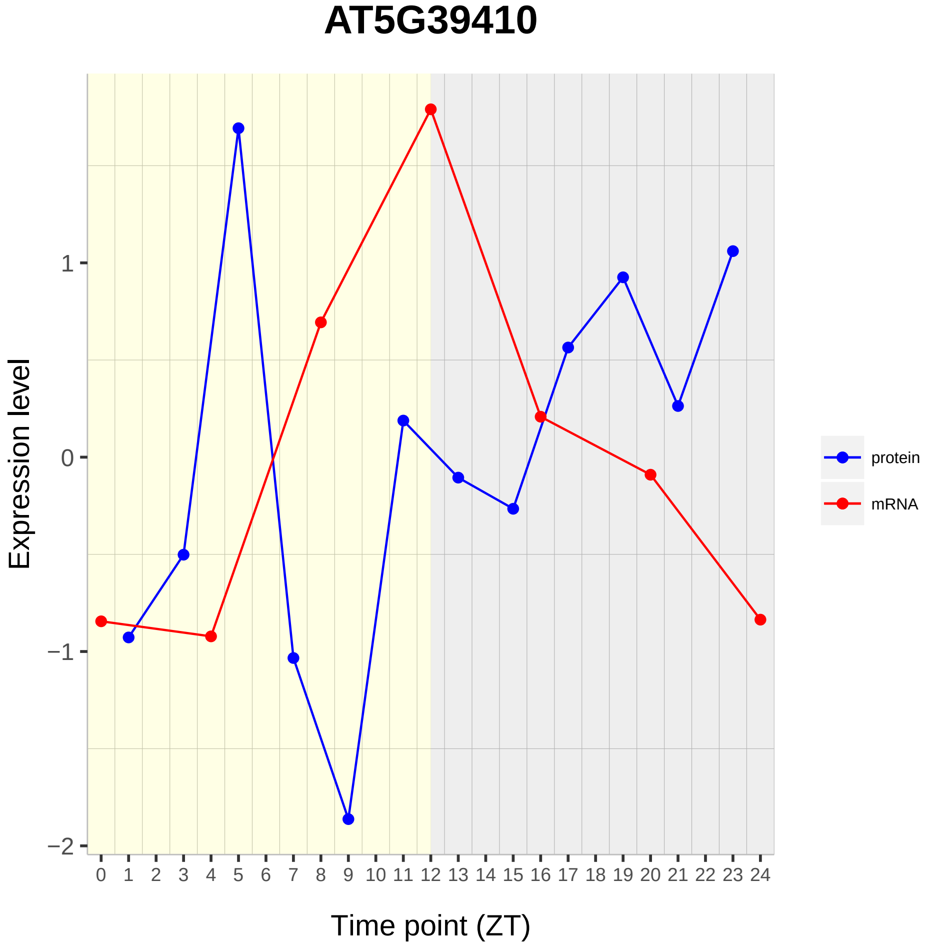

AT5G39410
Expression level
1
0
−1
−2
0 1 2 3 4 5 6 7 8 9 10 11 12 13 14 15 16 17 18 19 20 21 22 23 24
Time point (ZT)
protein
mRNA

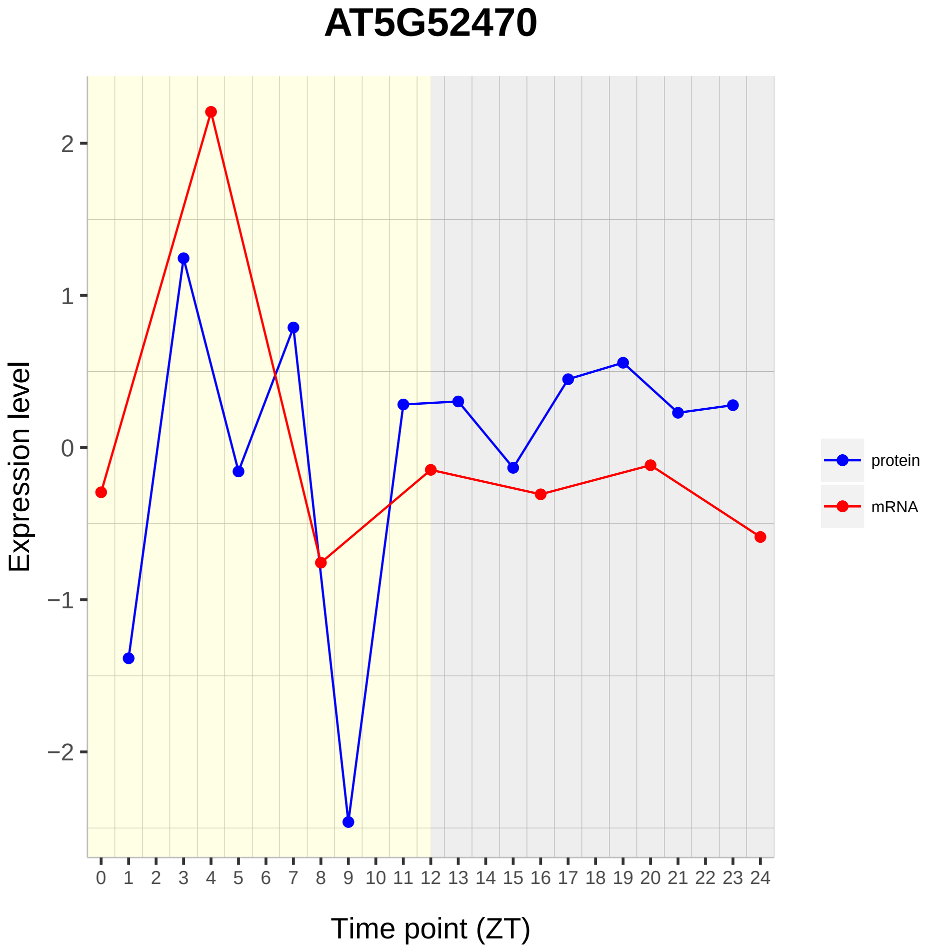
AT5G52470
Expression level
Time point (ZT)
2
1
0
−1
−2
0 1 2 3 4 5 6 7 8 9 10 11 12 13 14 15 16 17 18 19 20 21 22 23 24
protein
mRNA

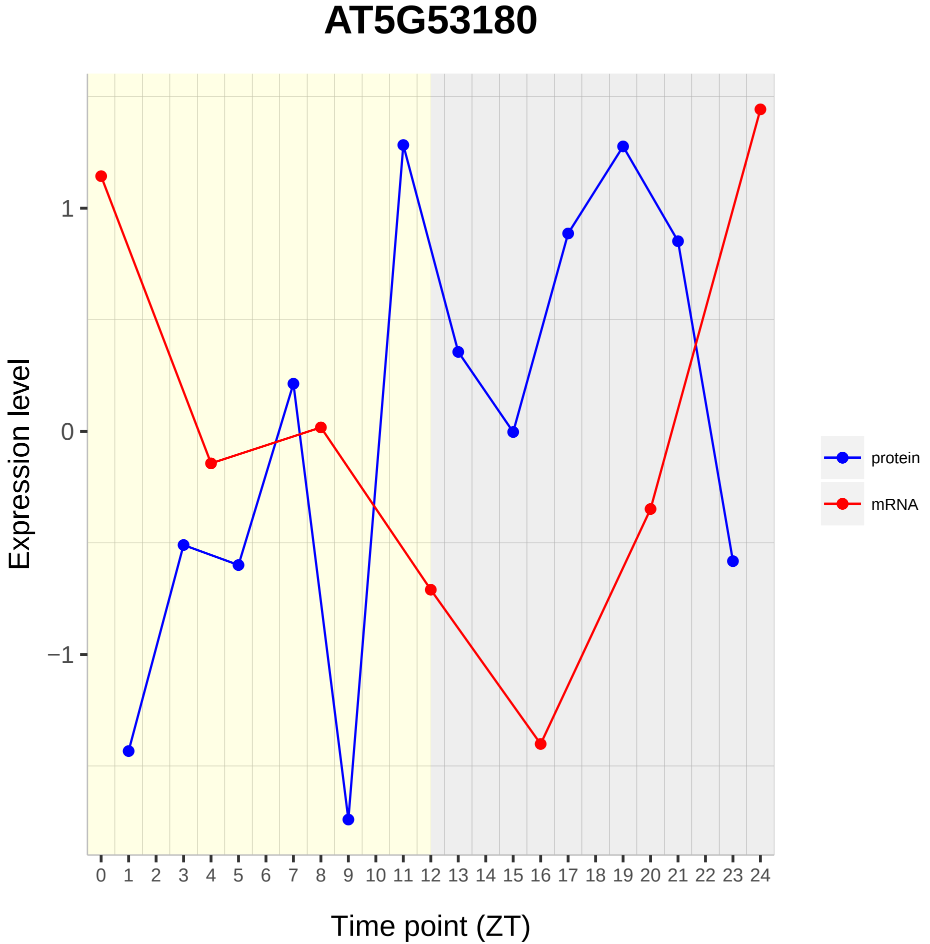

AT5G53180
Expression level
1
0
−1
Time point (ZT)
protein
mRNA

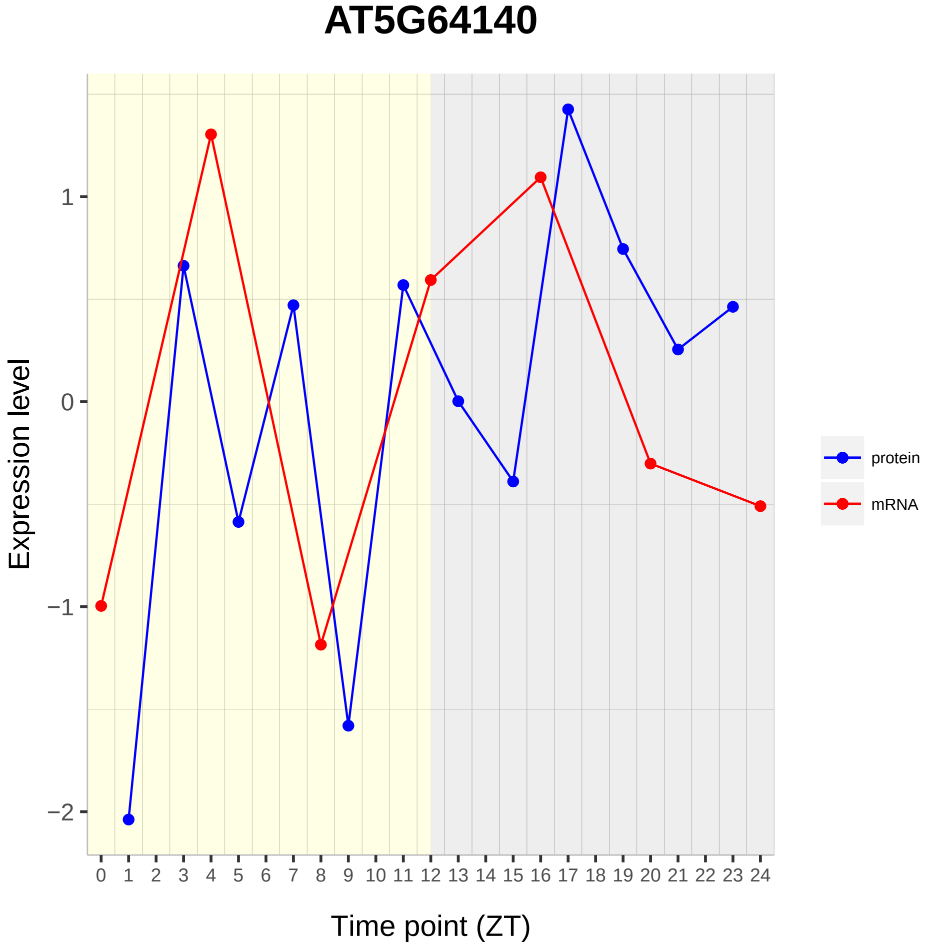

AT5G64140
Expression level
1
0
−1
−2
Time point (ZT)
protein
mRNA

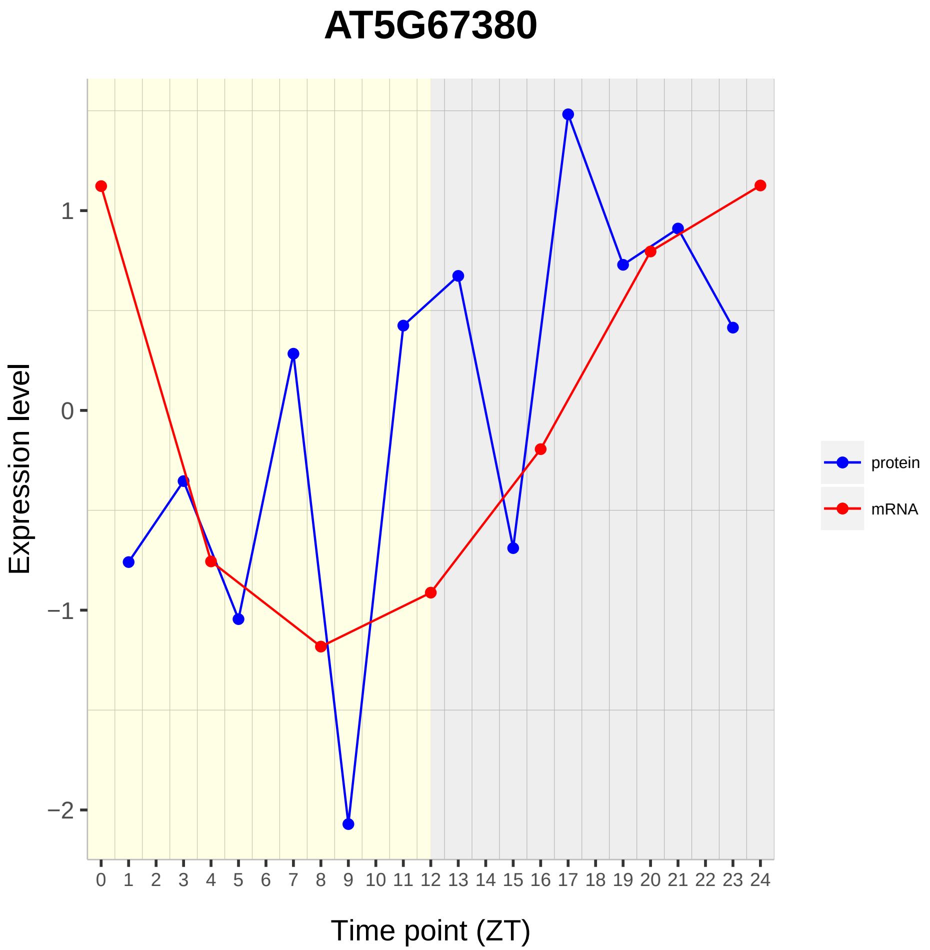

AT5G67380
Expression level
1
0
−1
−2
Time point (ZT)
protein
mRNA

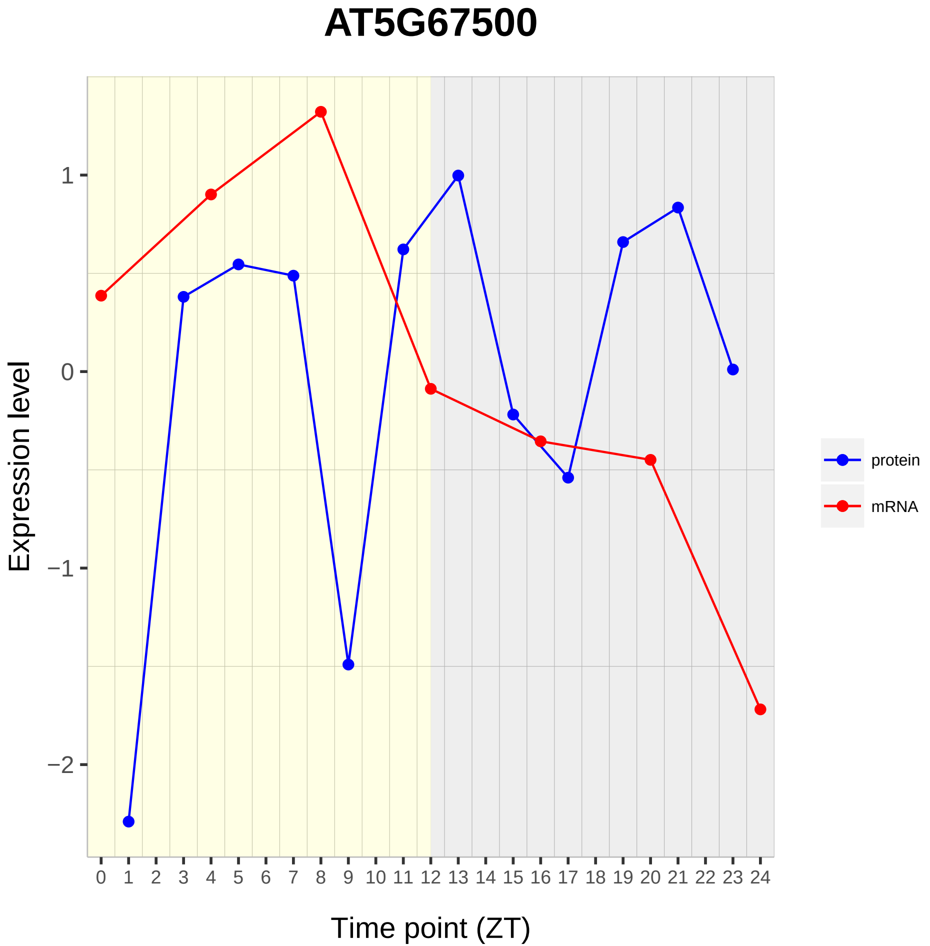

AT5G67500
Expression level
1
0
−1
−2
0 1 2 3 4 5 6 7 8 9 10 11 12 13 14 15 16 17 18 19 20 21 22 23 24
Time point (ZT)
protein
mRNA
