## Supplementary material for "Diurnal Dynamics of the Arabidopsis Rosette Proteome and Phosphoproteome": SuppData4

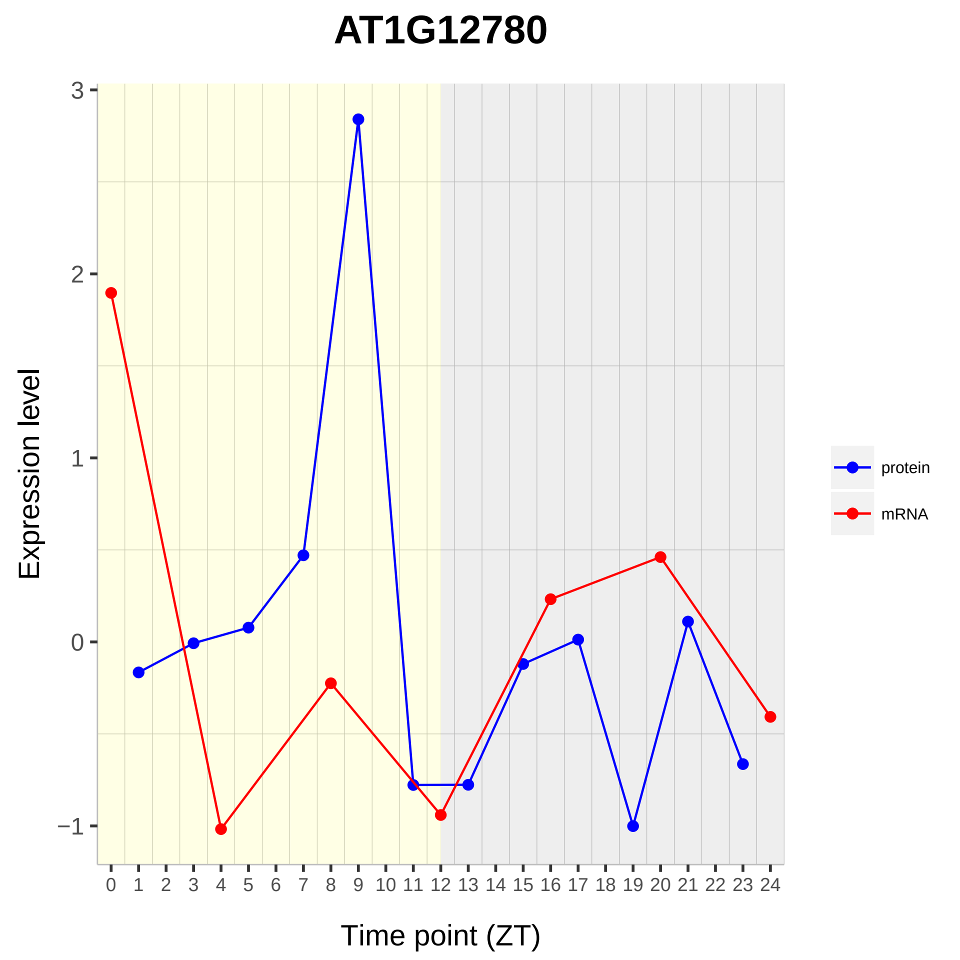

AT1G12780
Expression level
Time point (ZT)
3
2
1
0
−1
0 1 2 3 4 5 6 7 8 9 10 11 12 13 14 15 16 17 18 19 20 21 22 23 24
protein
mRNA

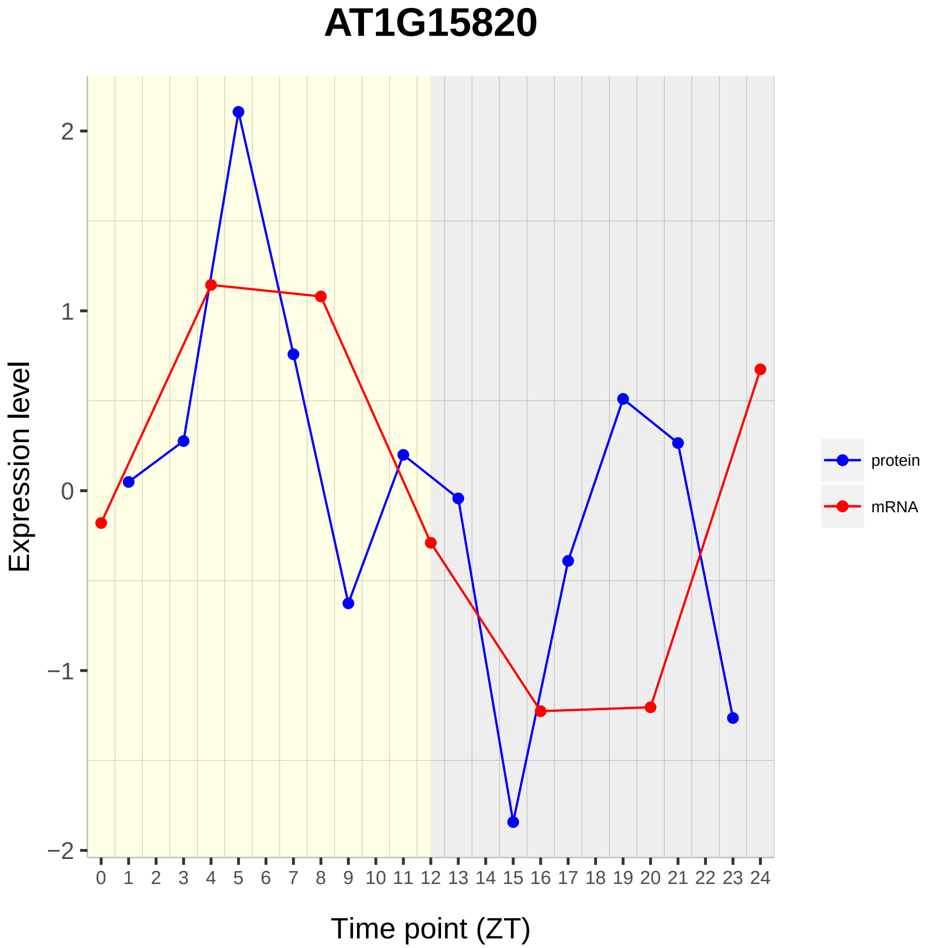

AT1G15820
Expression level
2
1
0
−1
−2
0 1 2 3 4 5 6 7 8 9 10 11 12 13 14 15 16 17 18 19 20 21 22 23 24
Time point (ZT)
protein
mRNA

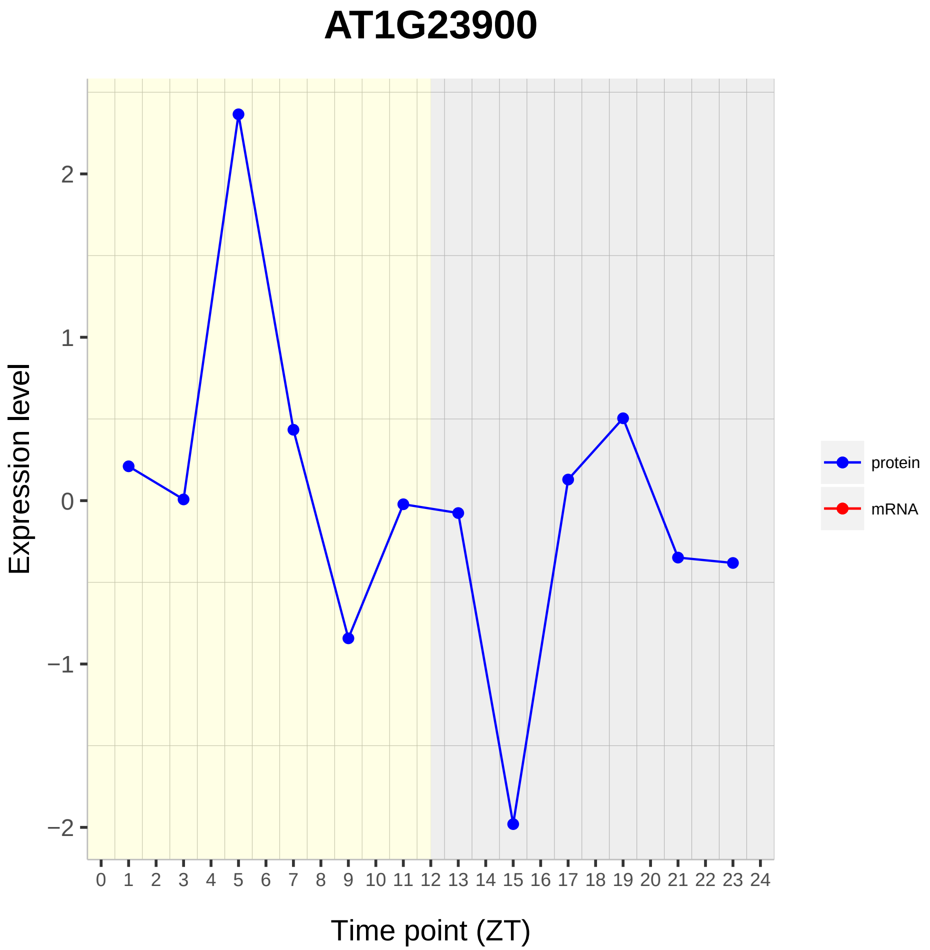
AT1G23900
Expression level
Time point (ZT)
2
1
0
−1
−2
0 1 2 3 4 5 6 7 8 9 10 11 12 13 14 15 16 17 18 19 20 21 22 23 24
protein
mRNA

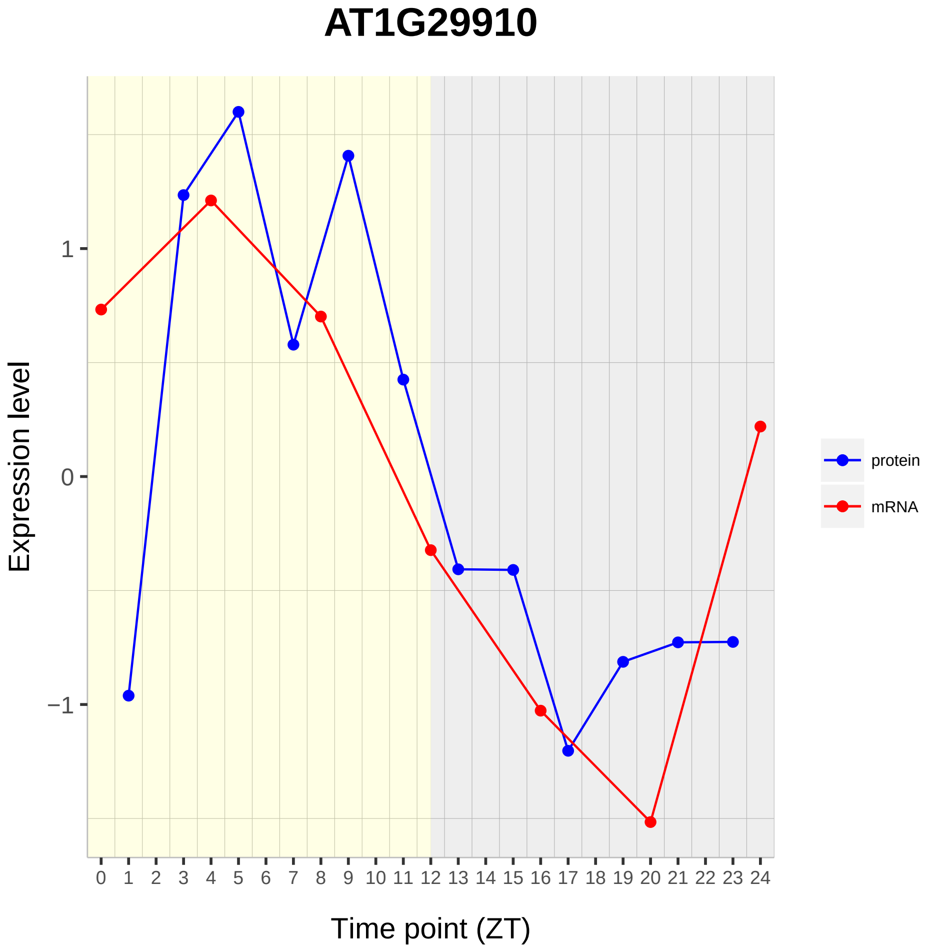

AT1G29910
Expression level
1
0
−1
0 1 2 3 4 5 6 7 8 9 10 11 12 13 14 15 16 17 18 19 20 21 22 23 24
Time point (ZT)
protein
mRNA

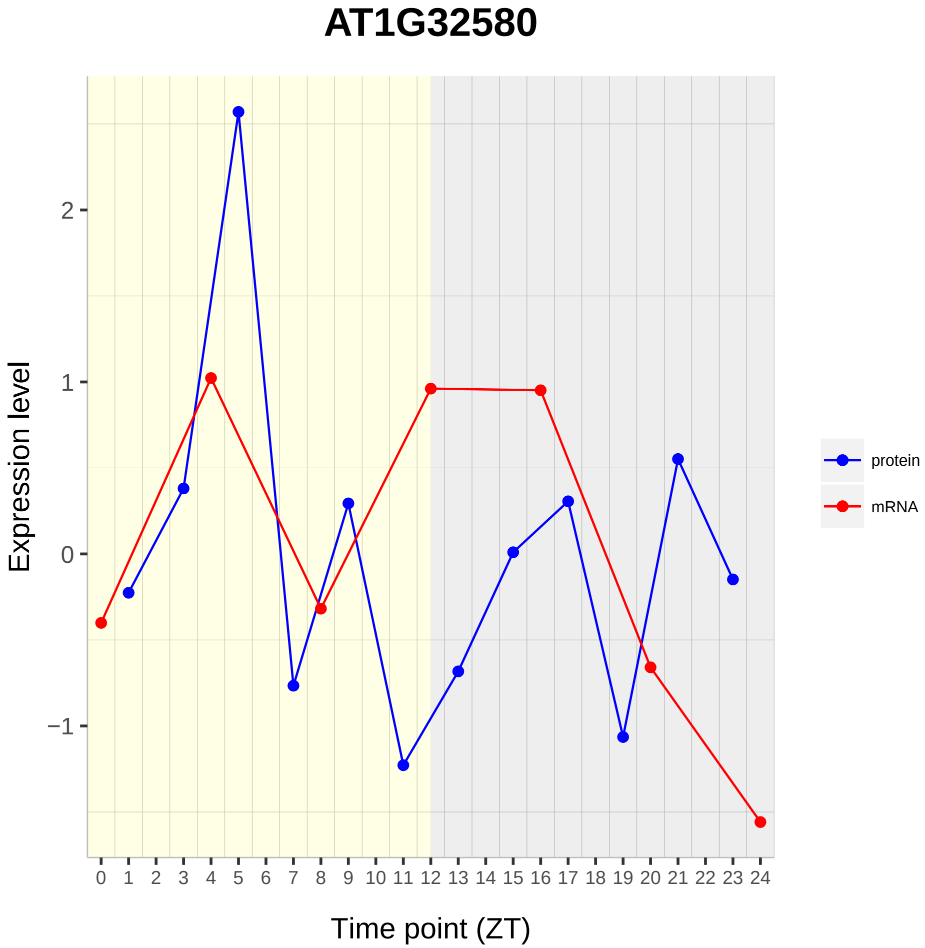
AT1G32580
Expression level
2
1
0
−1
0 1 2 3 4 5 6 7 8 9 10 11 12 13 14 15 16 17 18 19 20 21 22 23 24
Time point (ZT)
protein
mRNA

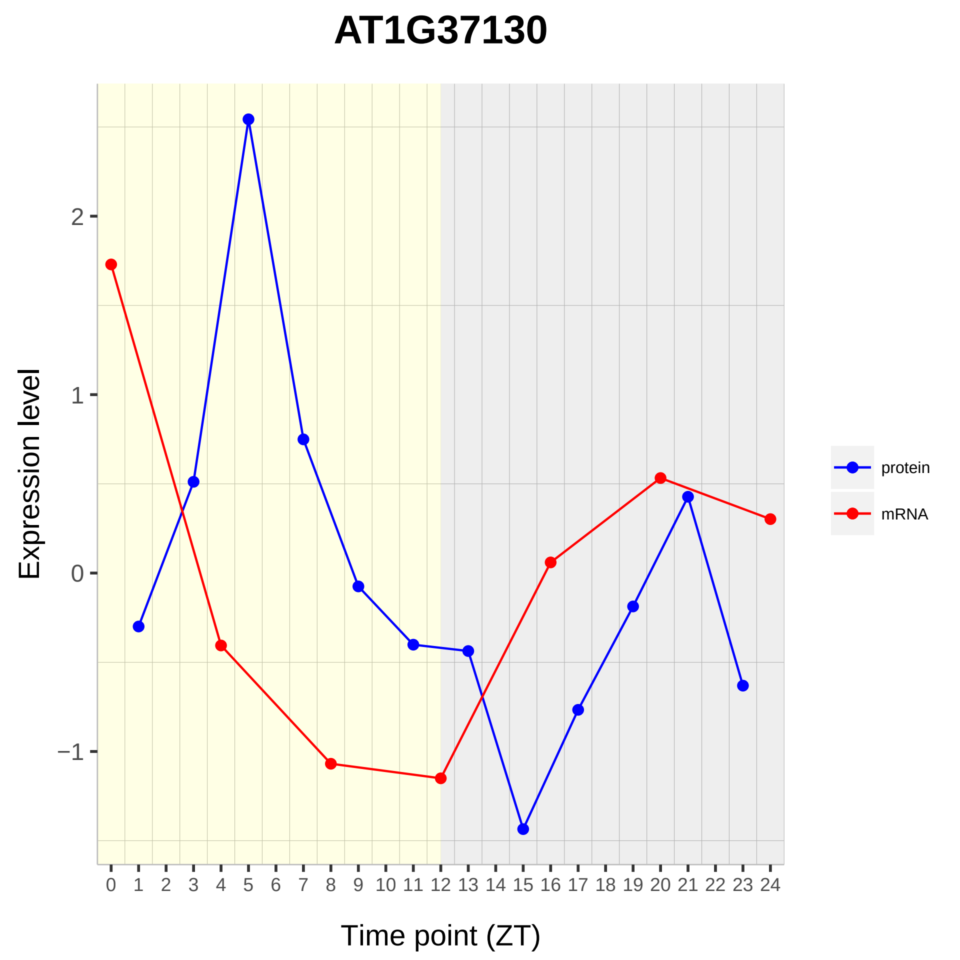

AT1G37130
Expression level
Time point (ZT)
protein
mRNA
2
1
0
−1
0 1 2 3 4 5 6 7 8 9 10 11 12 13 14 15 16 17 18 19 20 21 22 23 24

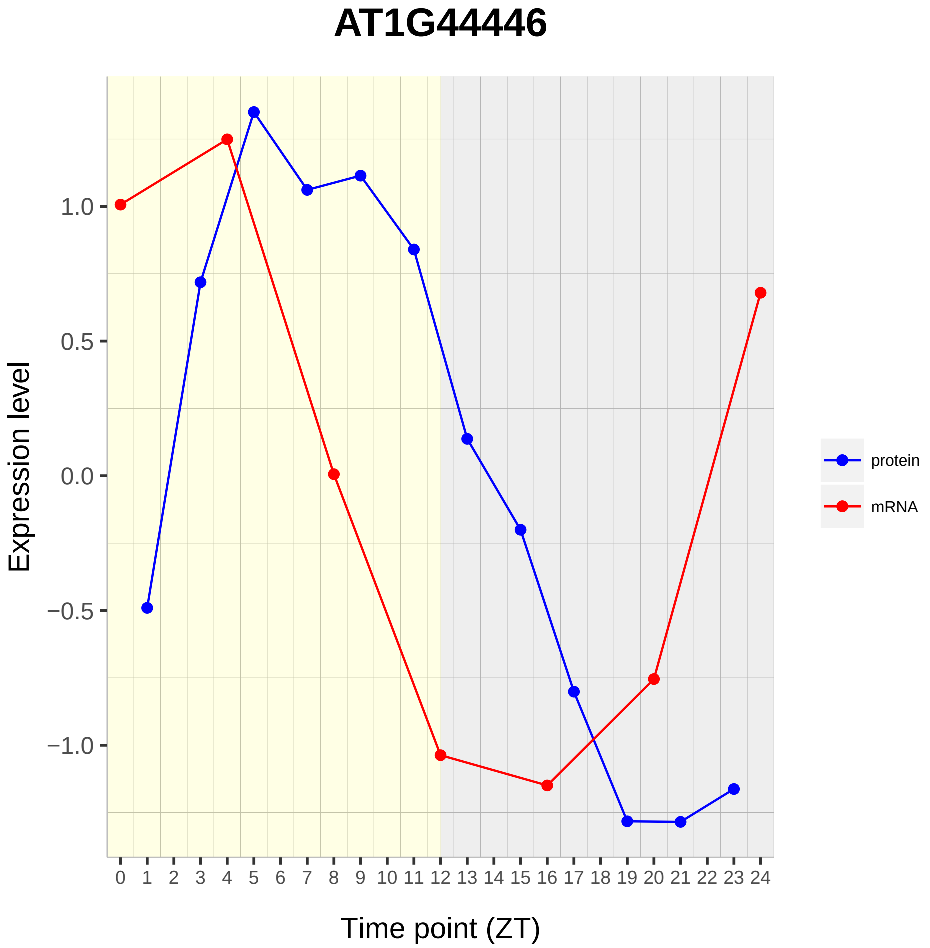

AT1G44446
Expression level
1.0
0.5
0.0
−0.5
−1.0
Time point (ZT)
0 1 2 3 4 5 6 7 8 9 10 11 12 13 14 15 16 17 18 19 20 21 22 23 24
protein
mRNA

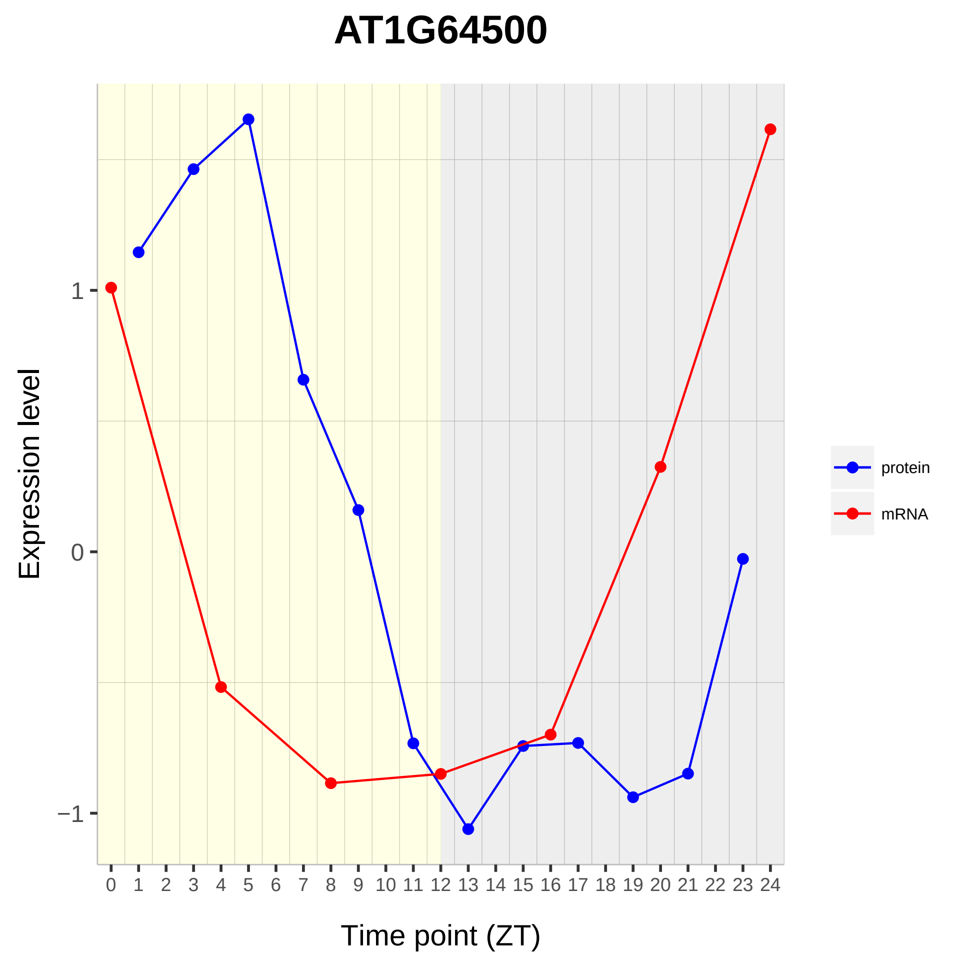
AT1G64500
Expression level
1
0
−1
0 1 2 3 4 5 6 7 8 9 10 11 12 13 14 15 16 17 18 19 20 21 22 23 24
Time point (ZT)
protein
mRNA

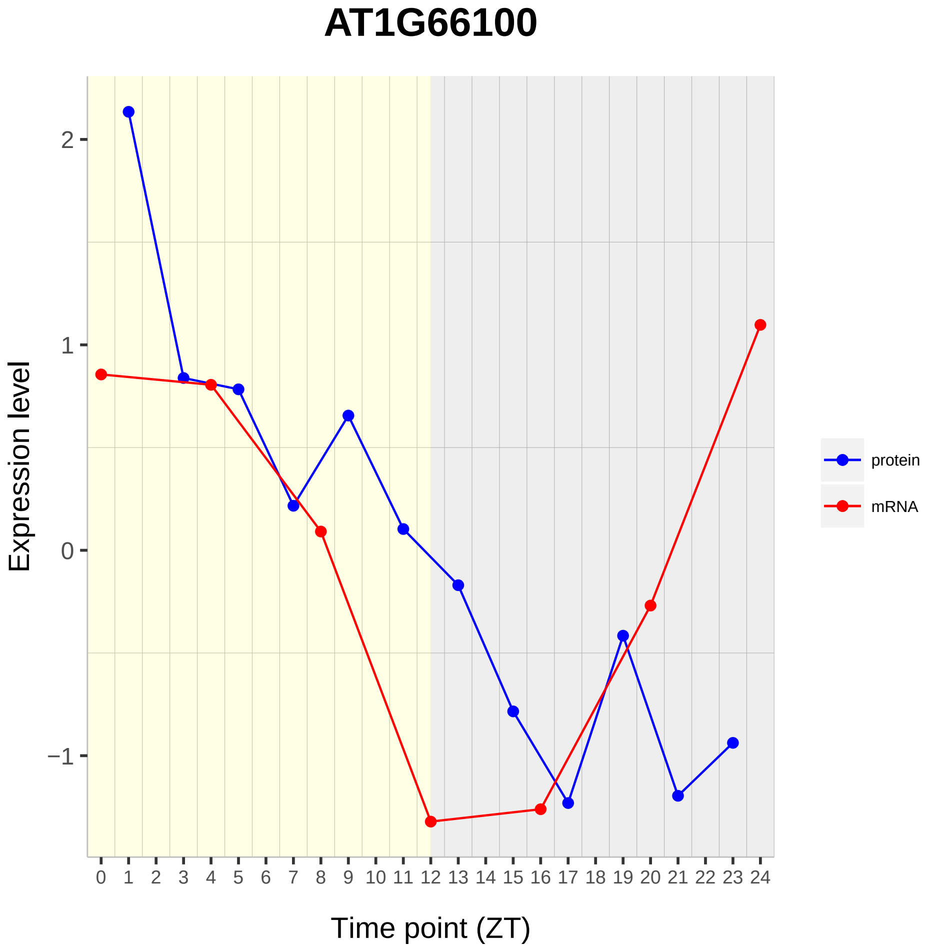

AT1G66100
Expression level
2
1
0
−1
0 1 2 3 4 5 6 7 8 9 10 11 12 13 14 15 16 17 18 19 20 21 22 23 24
Time point (ZT)
protein
mRNA

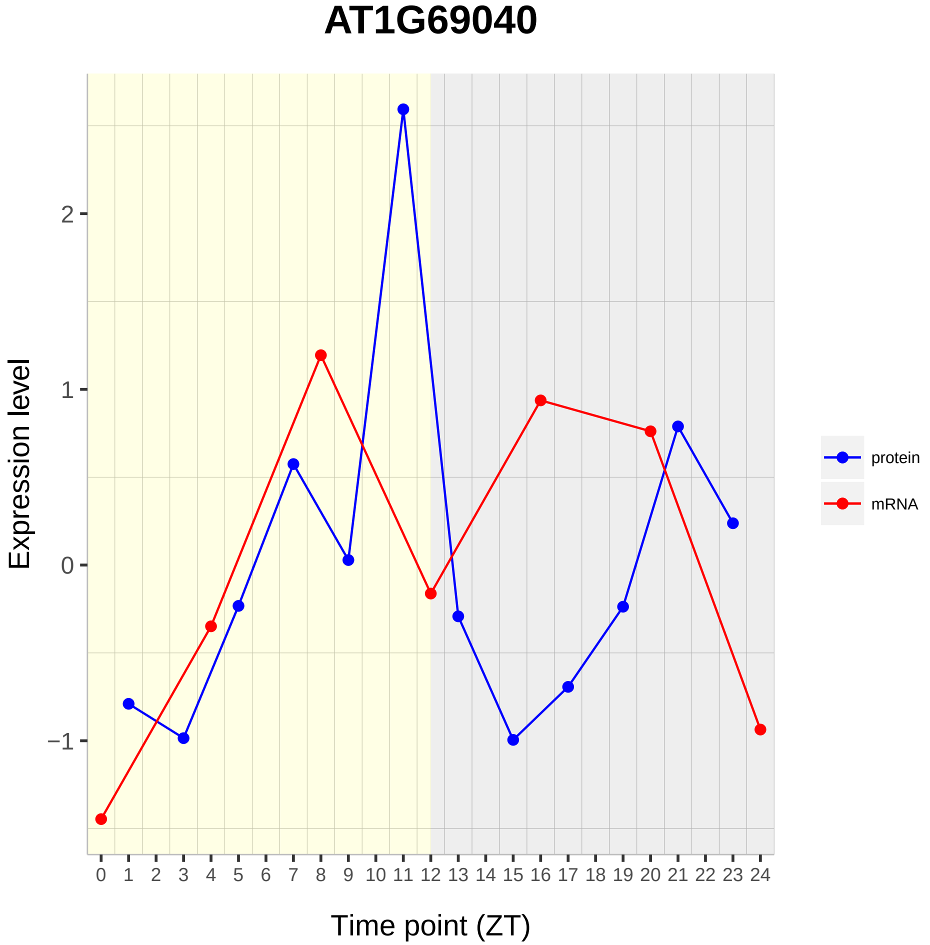

AT1G69040
Expression level
2
1
0
−1
0 1 2 3 4 5 6 7 8 9 10 11 12 13 14 15 16 17 18 19 20 21 22 23 24
Time point (ZT)
protein
mRNA

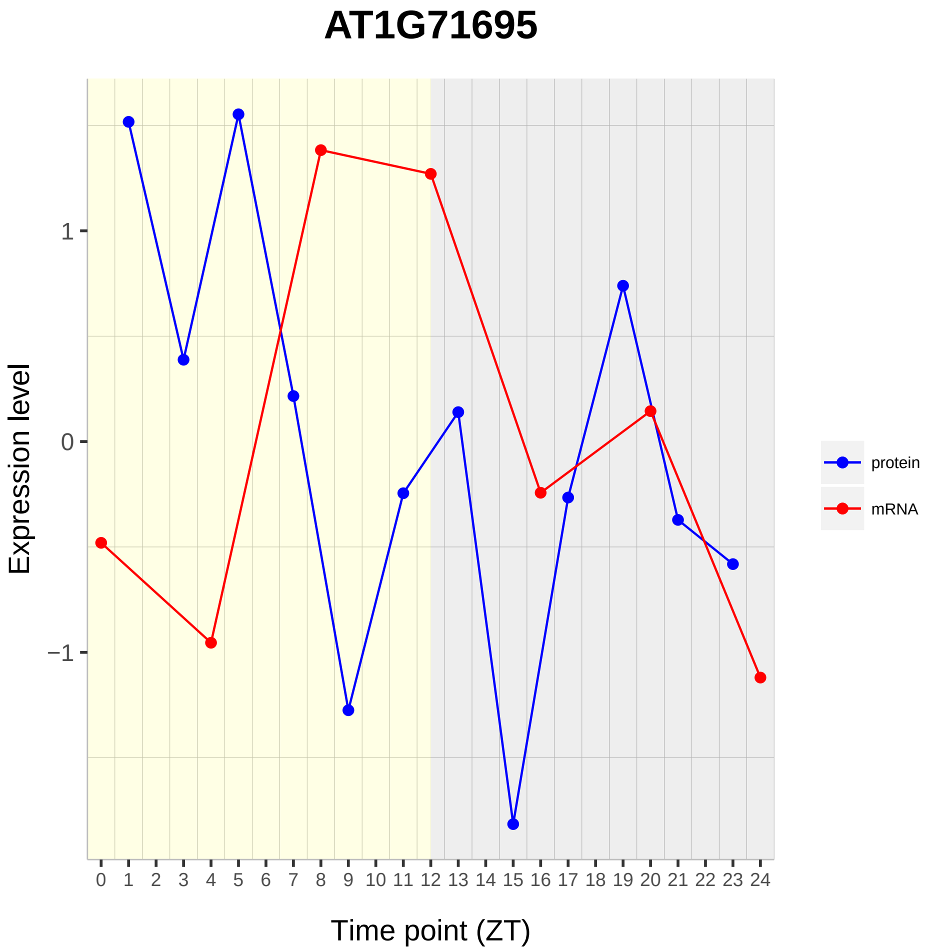

AT1G71695
Expression level
1
0
−1
0 1 2 3 4 5 6 7 8 9 10 11 12 13 14 15 16 17 18 19 20 21 22 23 24
Time point (ZT)
protein
mRNA

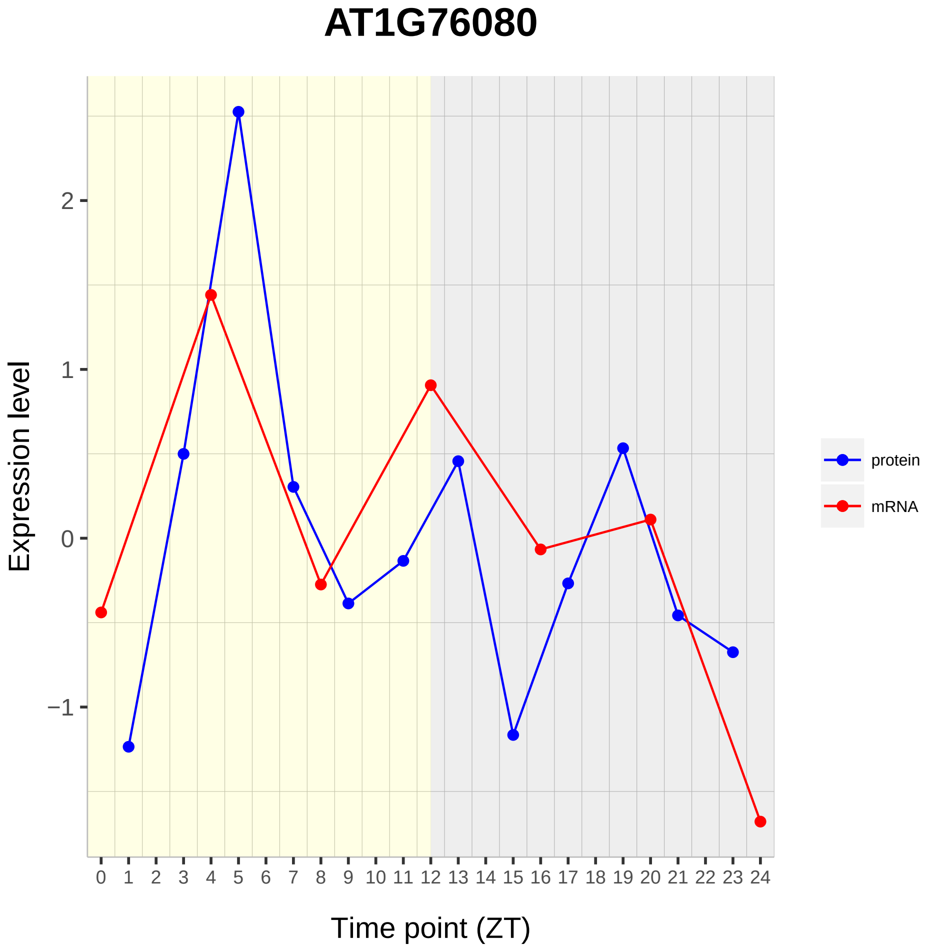

AT1G76080
Expression level
Time point (ZT)
2
1
0
−1
0 1 2 3 4 5 6 7 8 9 10 11 12 13 14 15 16 17 18 19 20 21 22 23 24
protein
mRNA

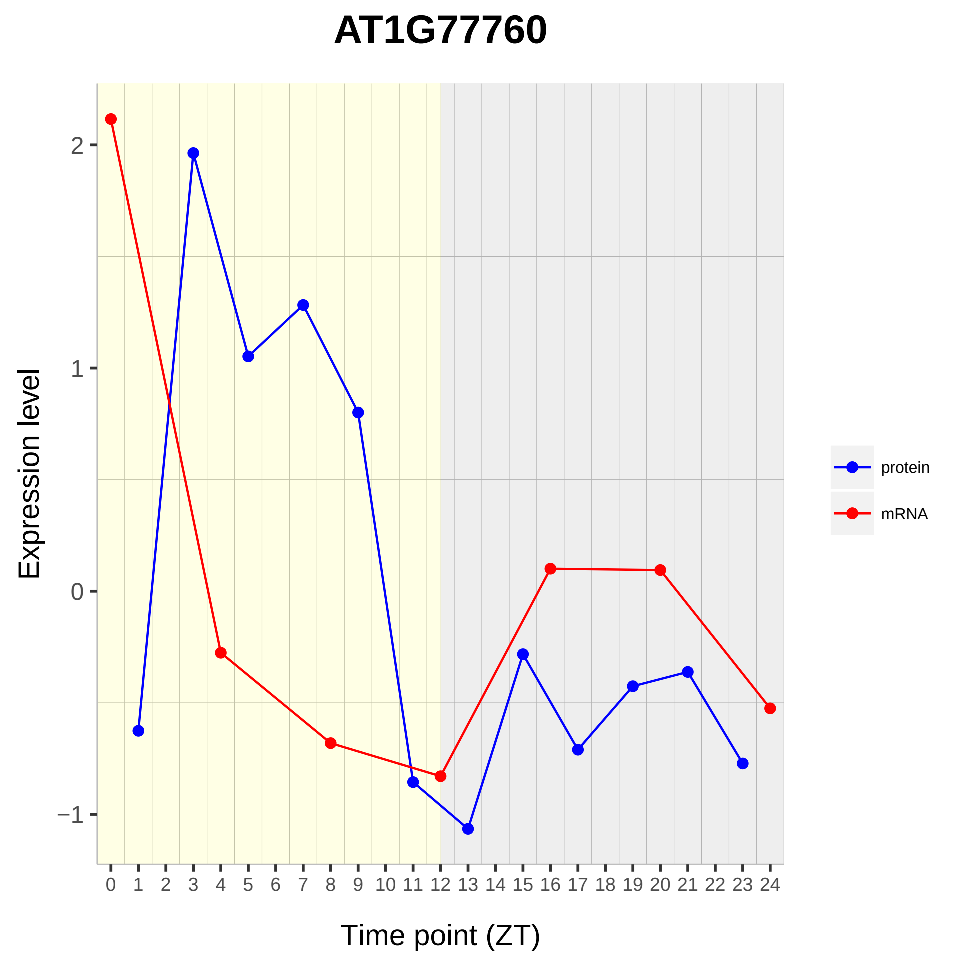

AT1G77760
Expression level
2
1
0
−1
0 1 2 3 4 5 6 7 8 9 10 11 12 13 14 15 16 17 18 19 20 21 22 23 24
Time point (ZT)
protein
mRNA

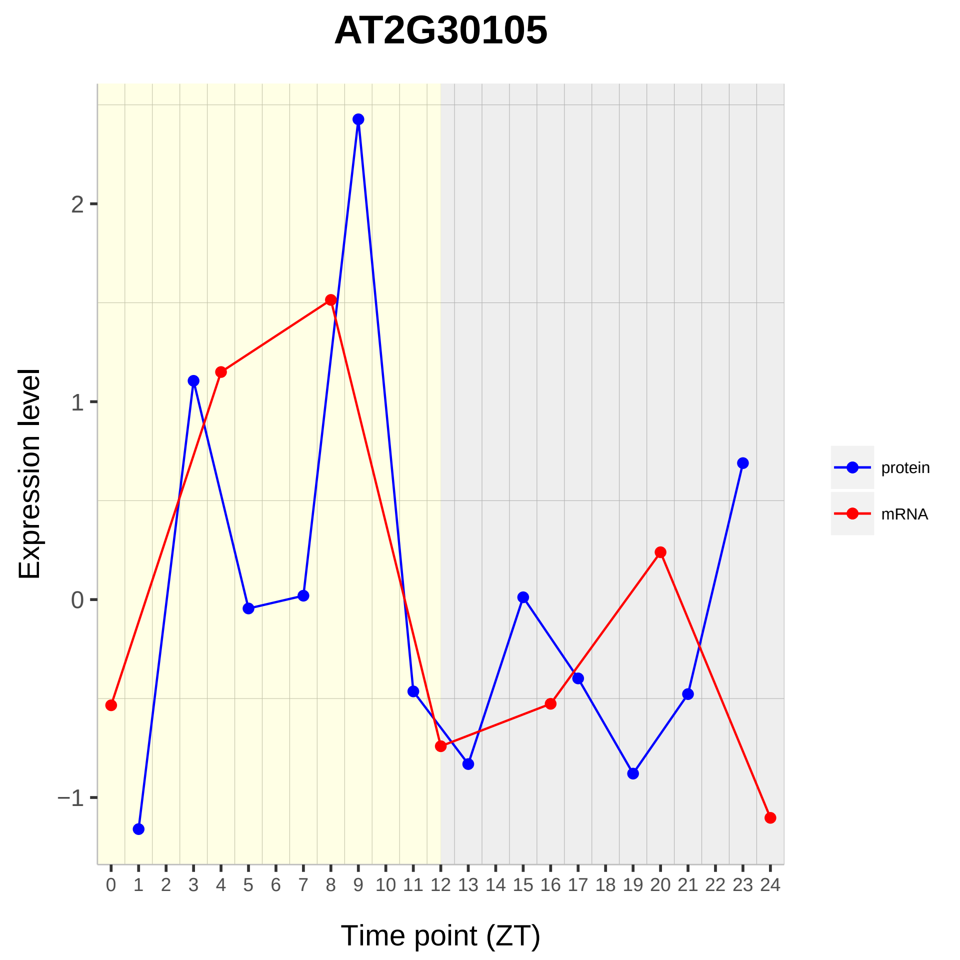

AT2G30105
Expression level
2
1
0
−1
Time point (ZT)
protein
mRNA

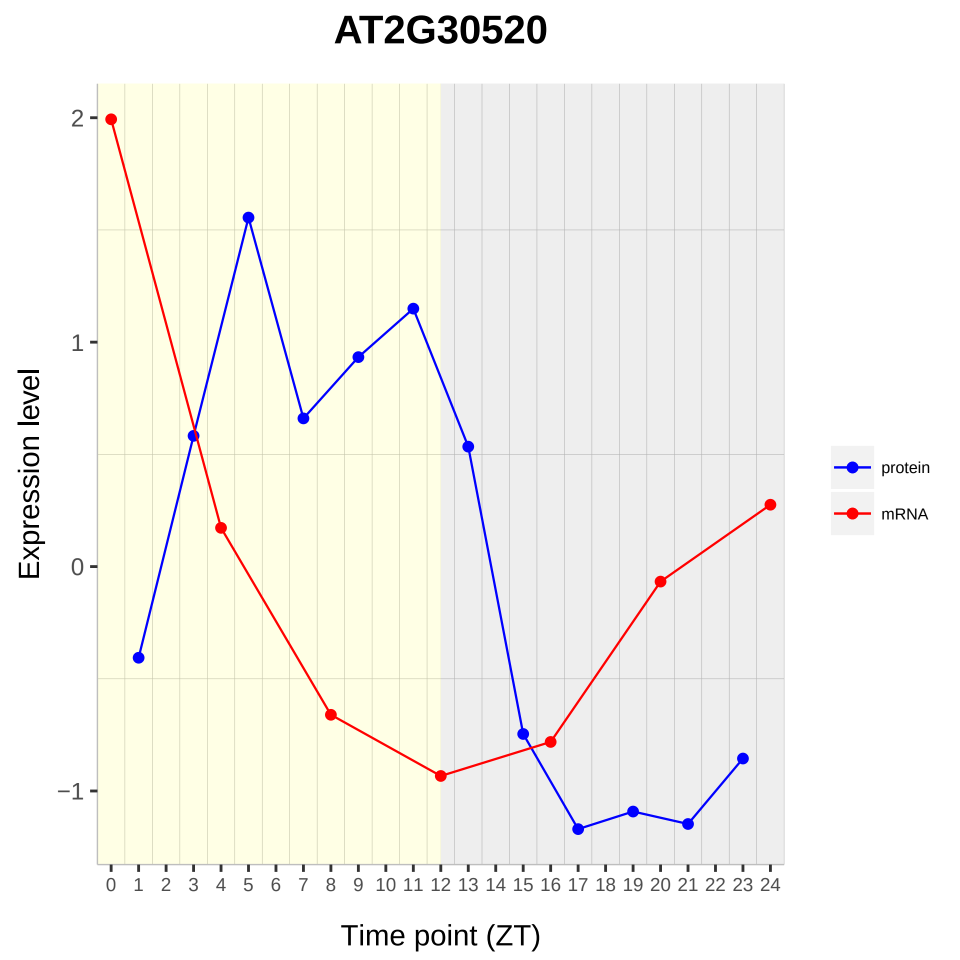

AT2G30520
Expression level
2
1
0
−1
Time point (ZT)
protein
mRNA

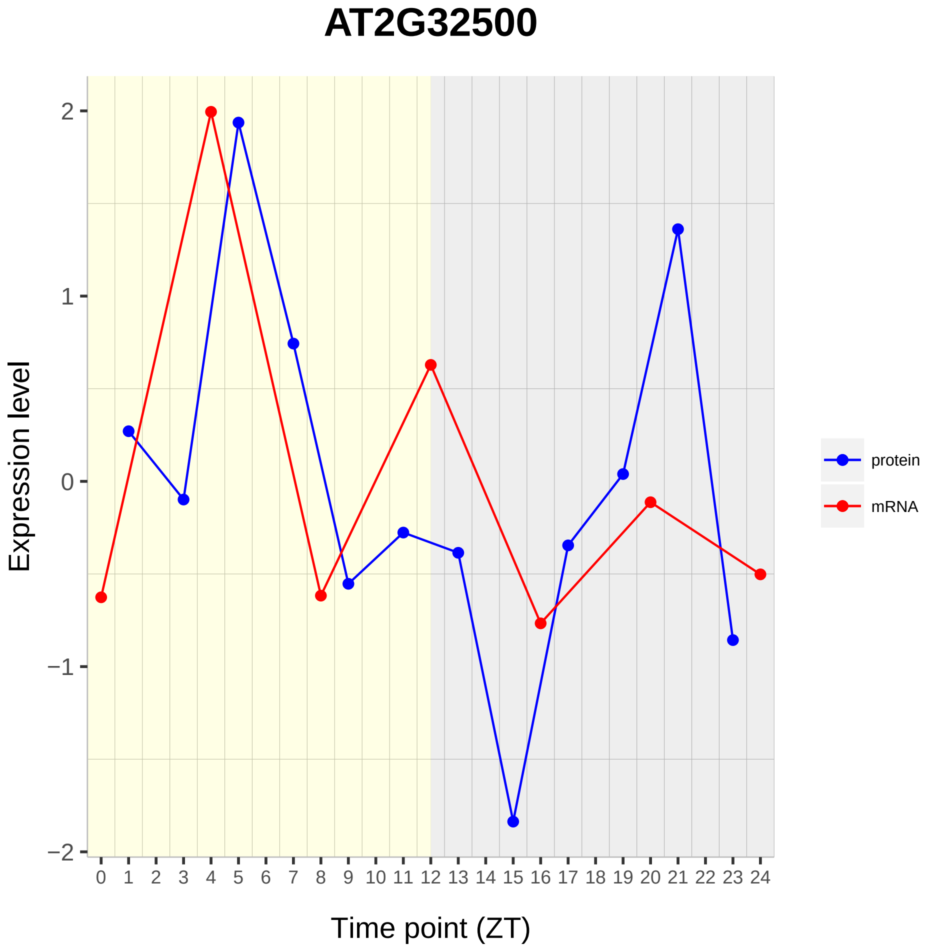

AT2G32500
Expression level
Time point (ZT)
2
1
0
−1
−2
0 1 2 3 4 5 6 7 8 9 10 11 12 13 14 15 16 17 18 19 20 21 22 23 24
protein
mRNA

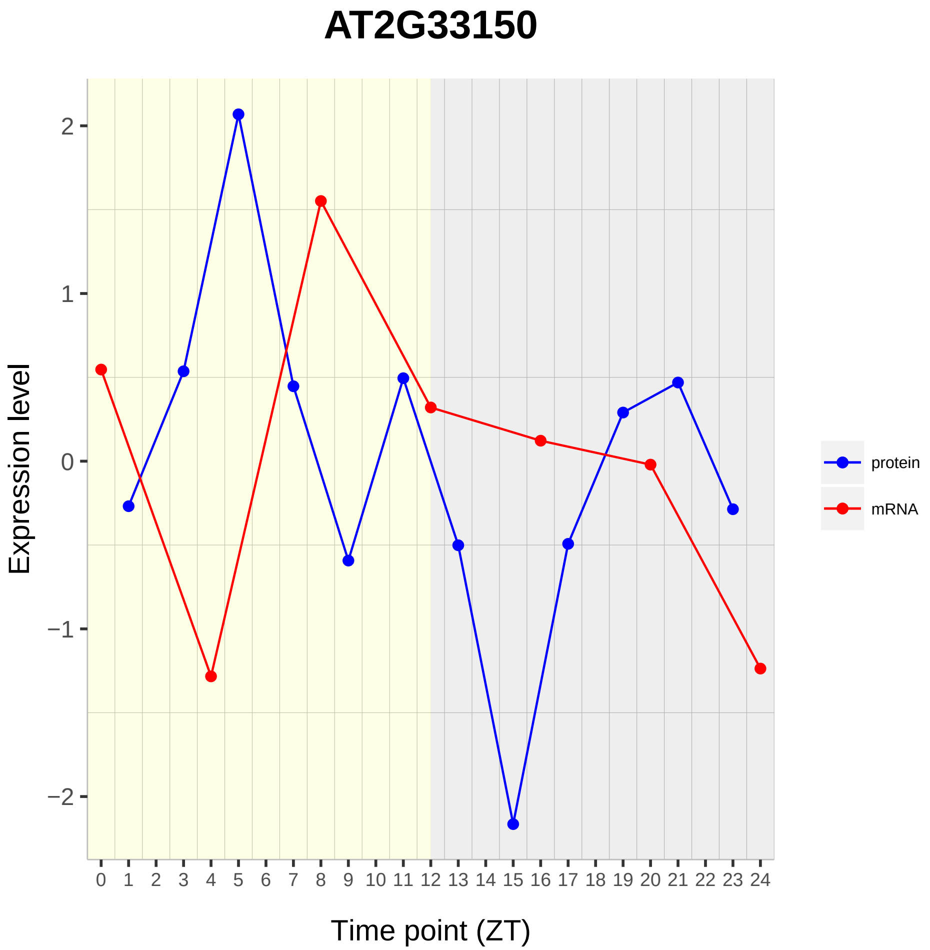

AT2G33150
Expression level
Time point (ZT)
2
1
0
−1
−2
0 1 2 3 4 5 6 7 8 9 10 11 12 13 14 15 16 17 18 19 20 21 22 23 24
protein
mRNA

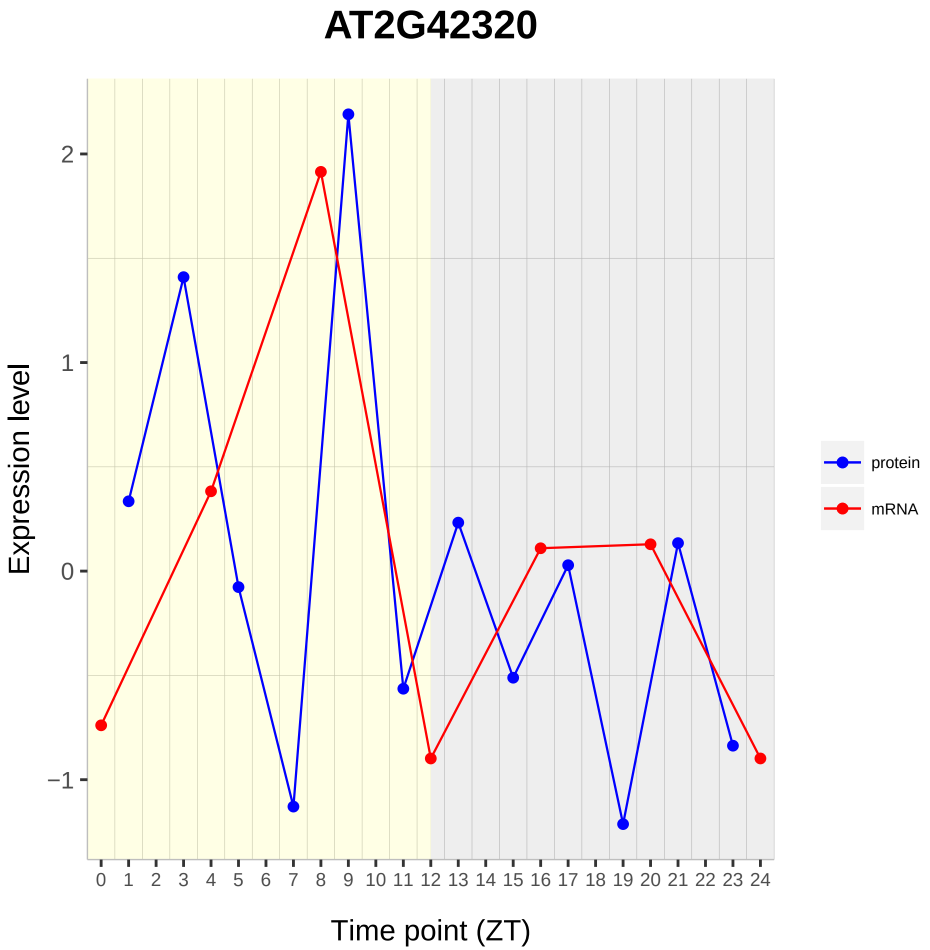

AT2G42320
Expression level
2
1
0
−1
0 1 2 3 4 5 6 7 8 9 10 11 12 13 14 15 16 17 18 19 20 21 22 23 24
Time point (ZT)
protein
mRNA

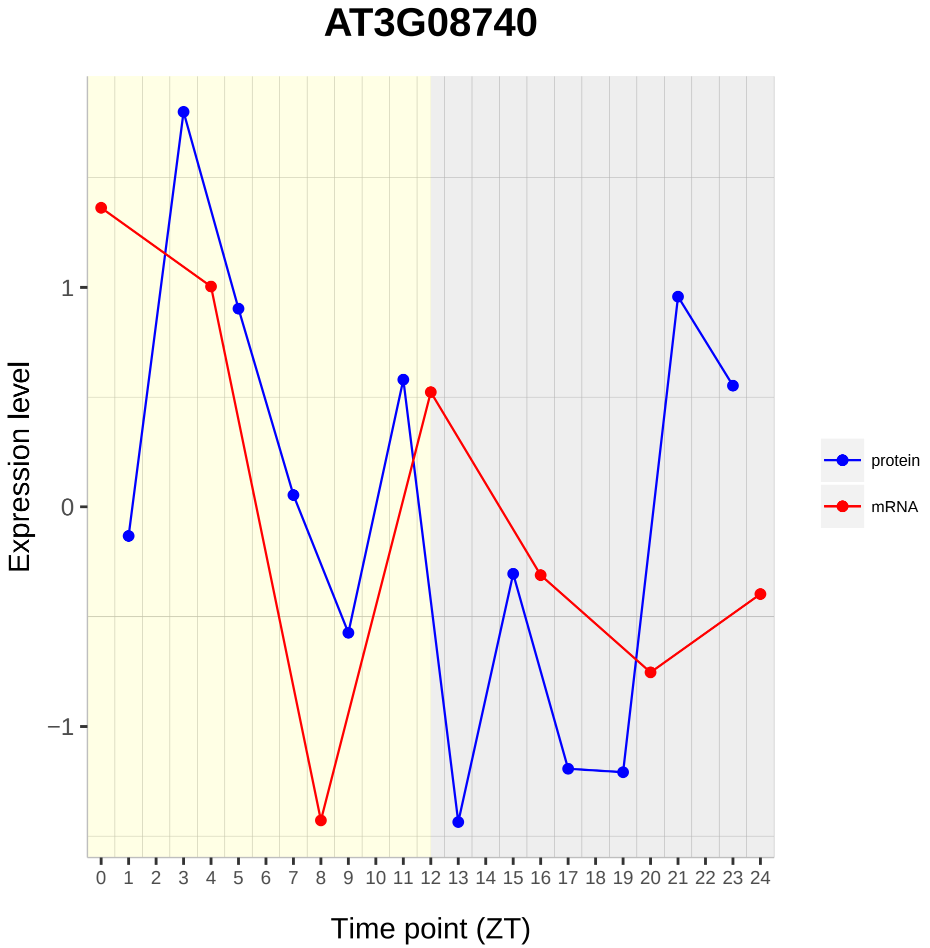
AT3G08740
Expression level
1
0
−1
0 1 2 3 4 5 6 7 8 9 10 11 12 13 14 15 16 17 18 19 20 21 22 23 24
Time point (ZT)
protein
mRNA

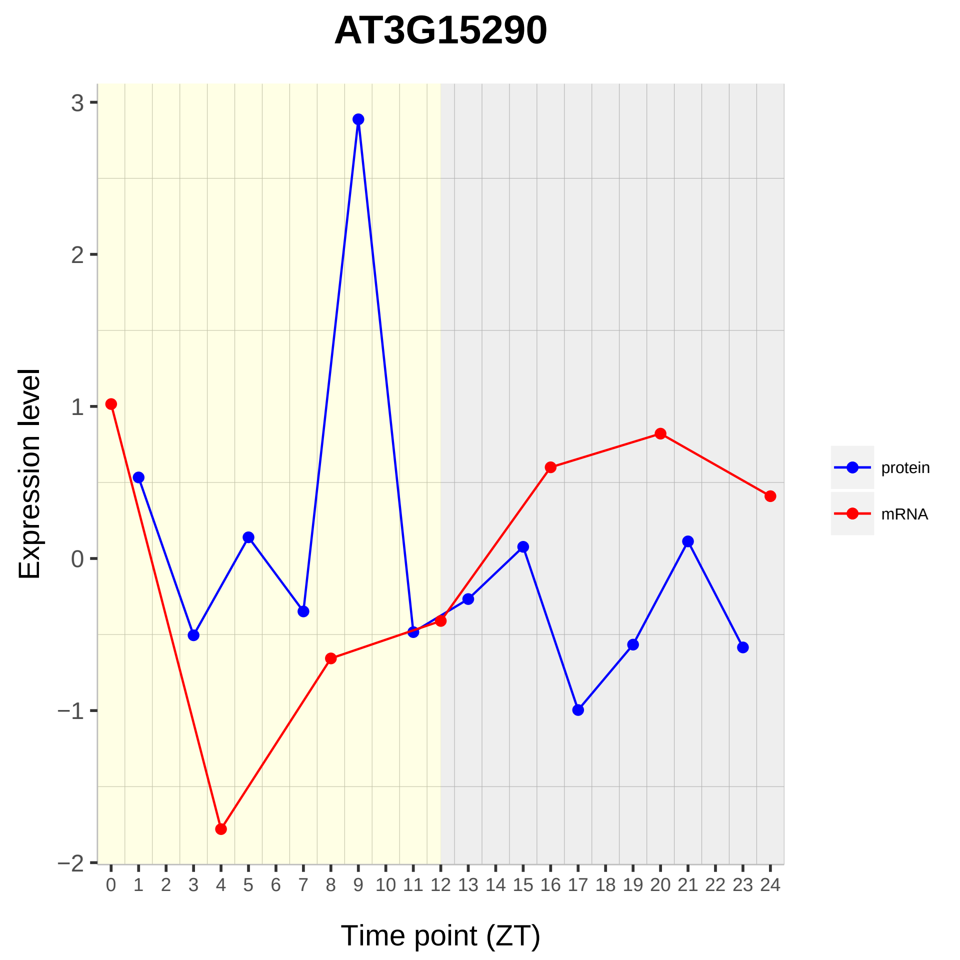

AT3G15290
Expression level
Time point (ZT)
3
2
1
0
−1
−2
0 1 2 3 4 5 6 7 8 9 10 11 12 13 14 15 16 17 18 19 20 21 22 23 24
protein
mRNA

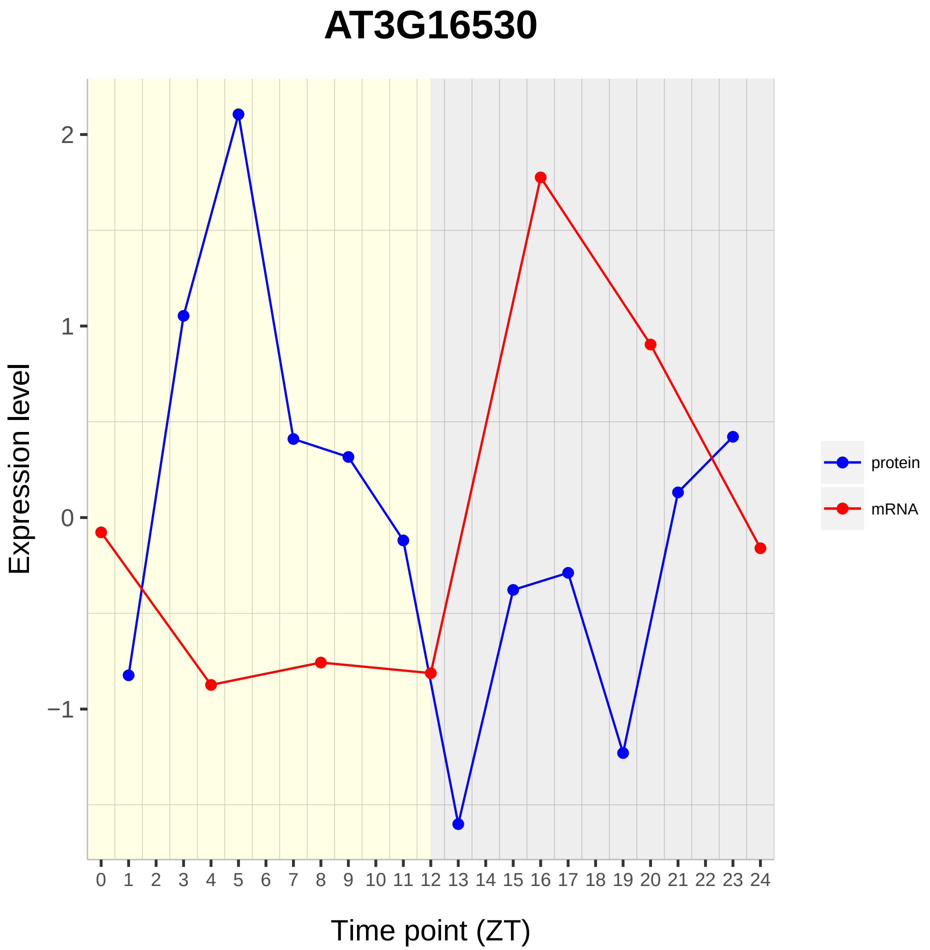

AT3G16530
Expression level
Time point (ZT)
2
1
0
−1
0 1 2 3 4 5 6 7 8 9 10 11 12 13 14 15 16 17 18 19 20 21 22 23 24
protein
mRNA

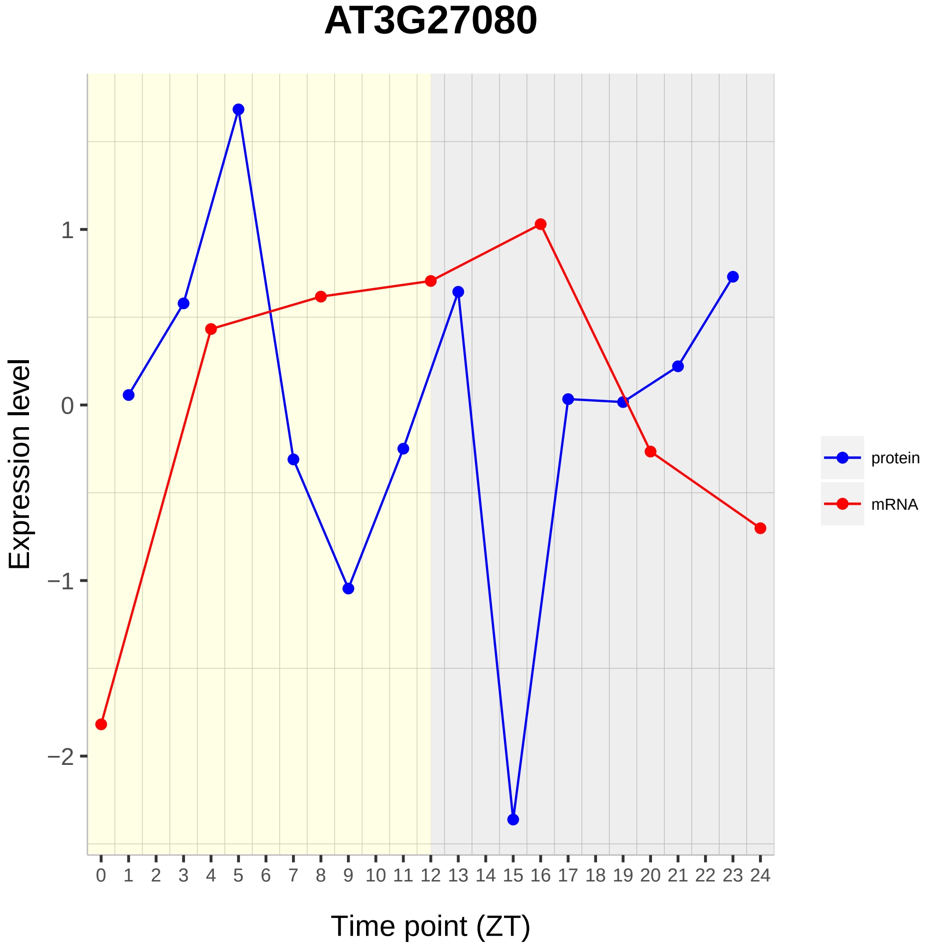

AT3G27080
Expression level
1
0
−1
−2
0 1 2 3 4 5 6 7 8 9 10 11 12 13 14 15 16 17 18 19 20 21 22 23 24
Time point (ZT)
protein
mRNA

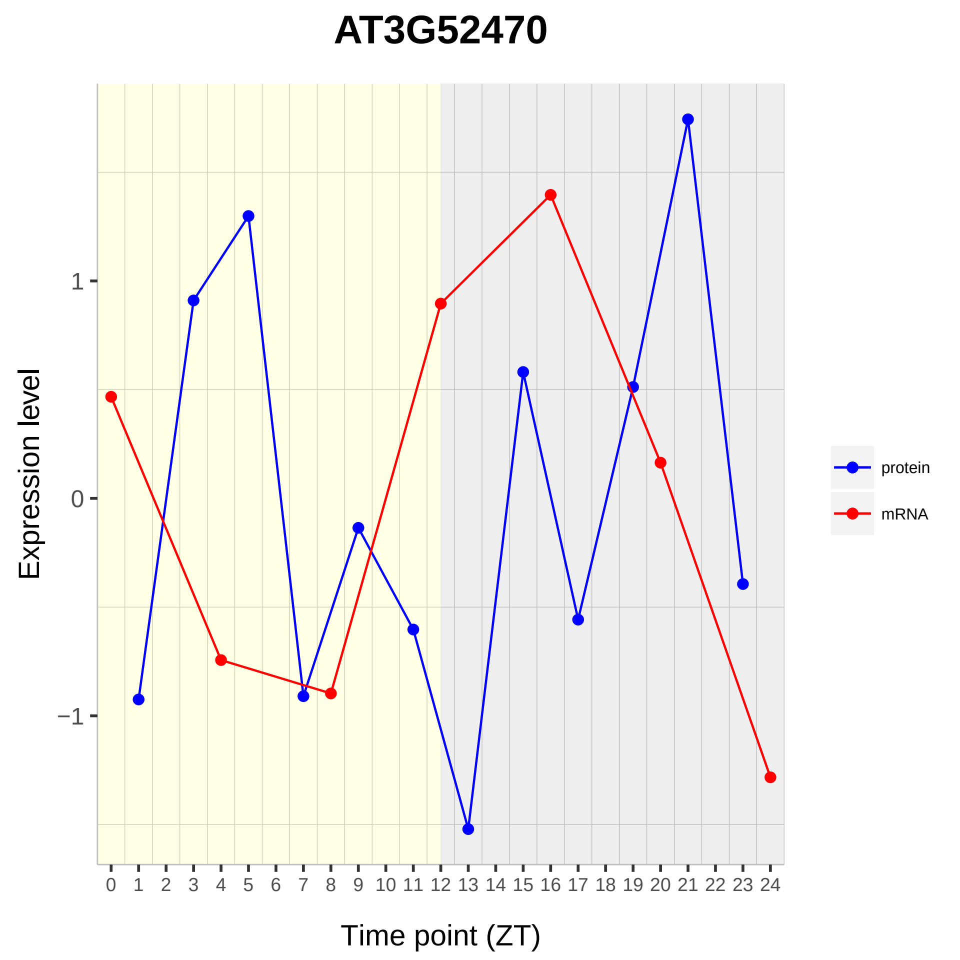

AT3G52470
Expression level
1
0
−1
Time point (ZT)
protein
mRNA

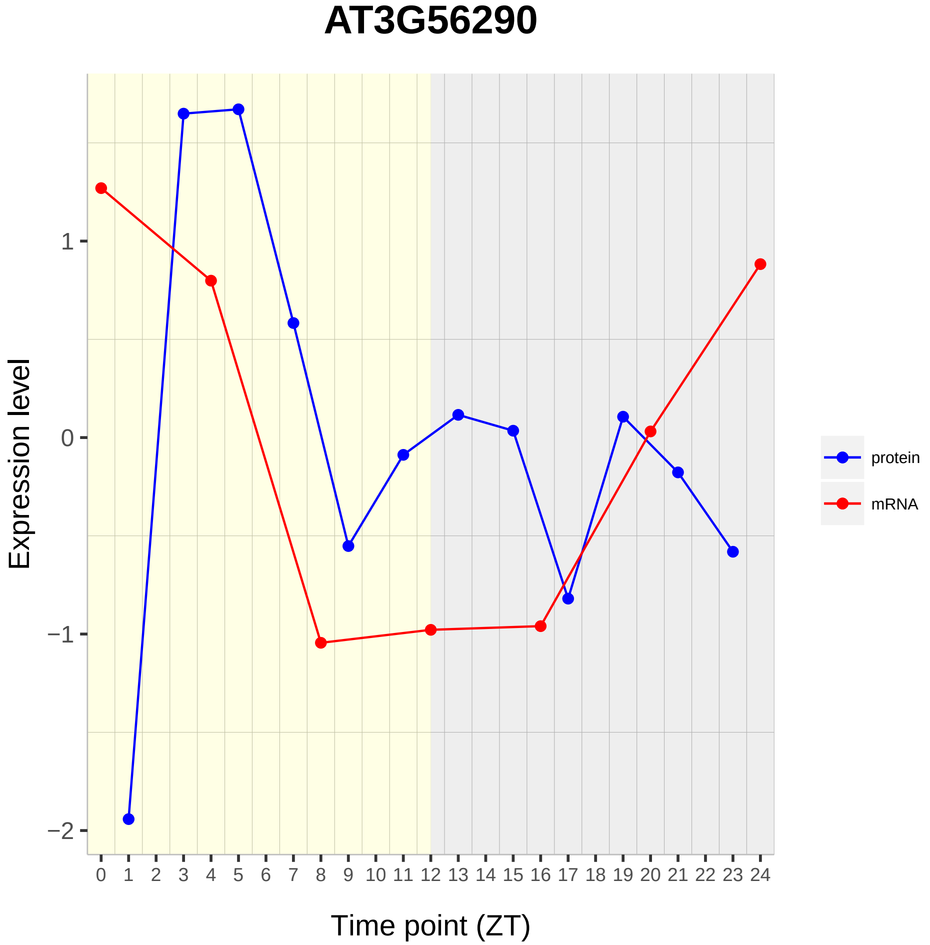
AT3G56290
Expression level
Time point (ZT)
1
0
−1
−2
0 1 2 3 4 5 6 7 8 9 10 11 12 13 14 15 16 17 18 19 20 21 22 23 24
protein
mRNA

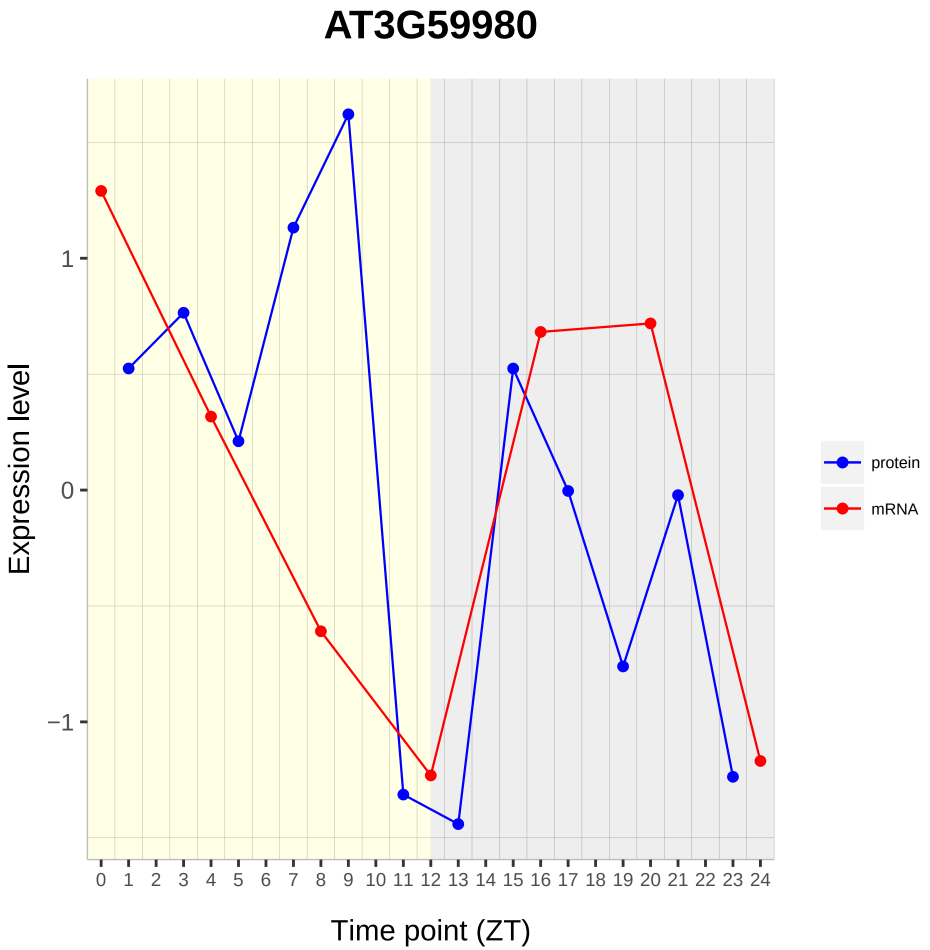

AT3G59980
Expression level
1
0
−1
0 1 2 3 4 5 6 7 8 9 10 11 12 13 14 15 16 17 18 19 20 21 22 23 24
Time point (ZT)
protein
mRNA

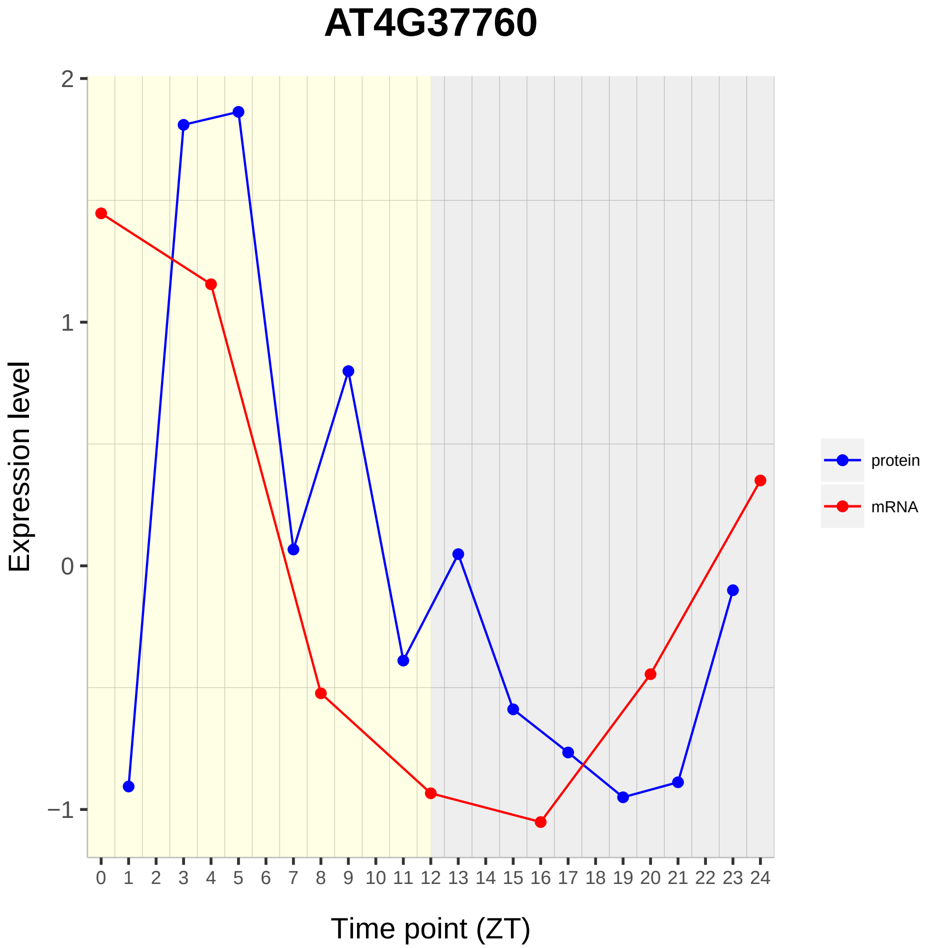
AT4G37760
Expression level
Time point (ZT)
2
1
0
−1
0 1 2 3 4 5 6 7 8 9 10 11 12 13 14 15 16 17 18 19 20 21 22 23 24
protein
mRNA

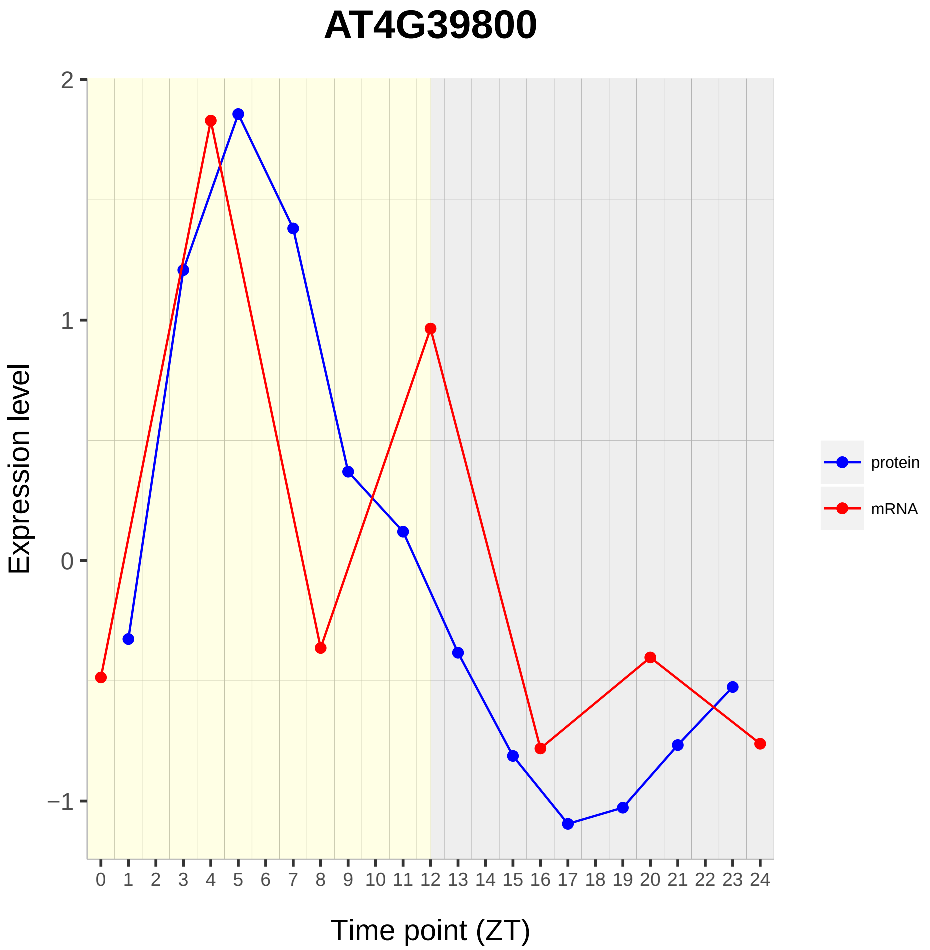
AT4G39800
Expression level
2
1
0
−1
0 1 2 3 4 5 6 7 8 9 10 11 12 13 14 15 16 17 18 19 20 21 22 23 24
Time point (ZT)
protein
mRNA

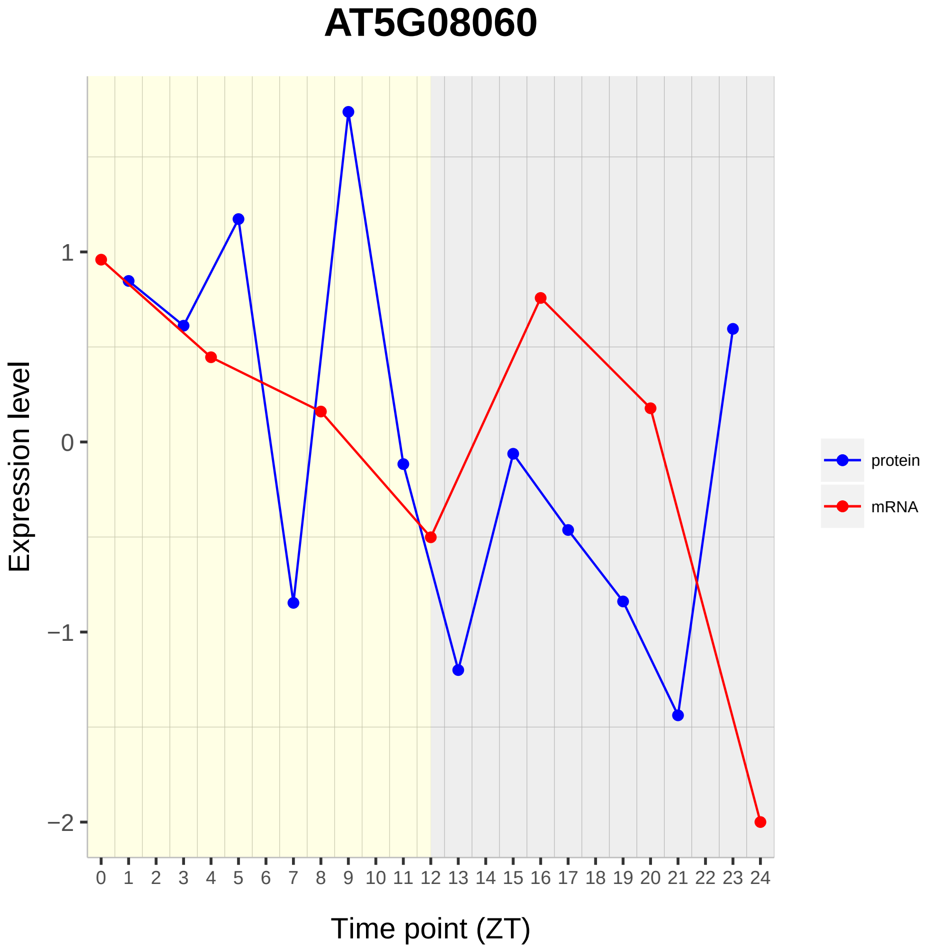

AT5G08060
Expression level
1
0
−1
−2
0 1 2 3 4 5 6 7 8 9 10 11 12 13 14 15 16 17 18 19 20 21 22 23 24
Time point (ZT)
protein
mRNA

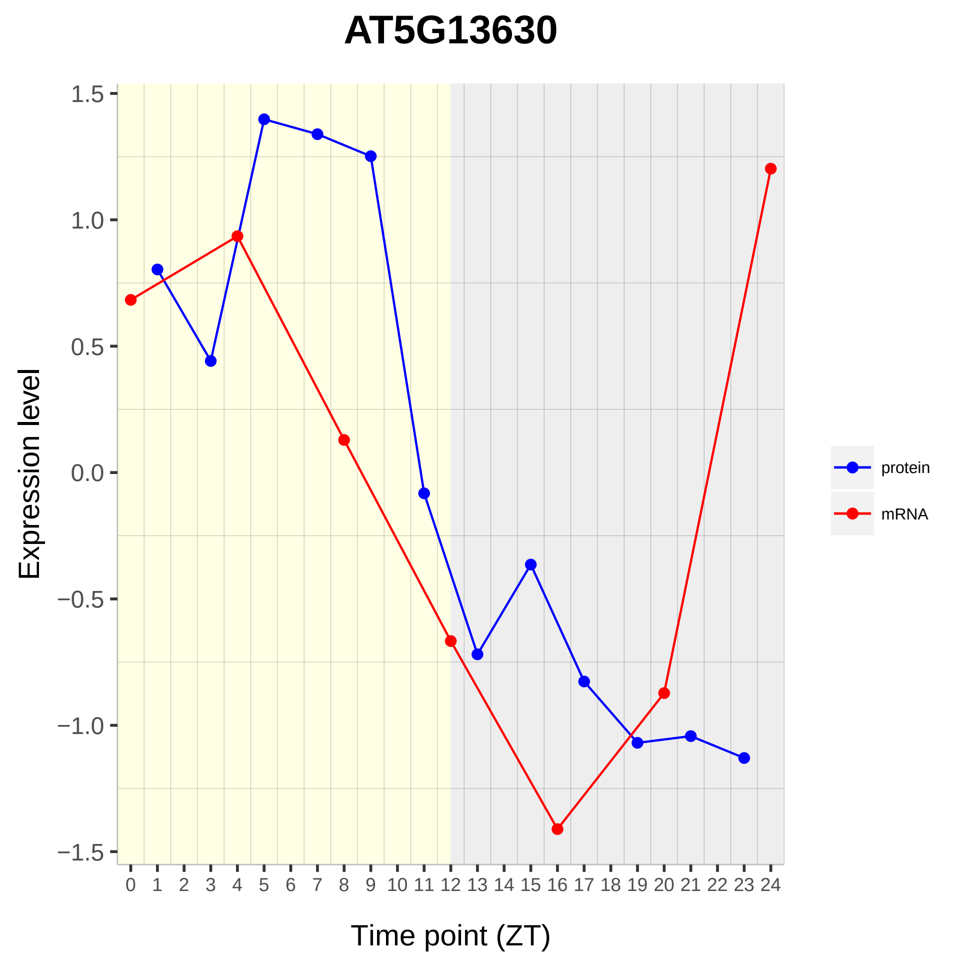
AT5G13630
Expression level
1.5
1.0
0.5
0.0
−0.5
−1.0
−1.5
0 1 2 3 4 5 6 7 8 9 10 11 12 13 14 15 16 17 18 19 20 21 22 23 24
Time point (ZT)
protein
mRNA

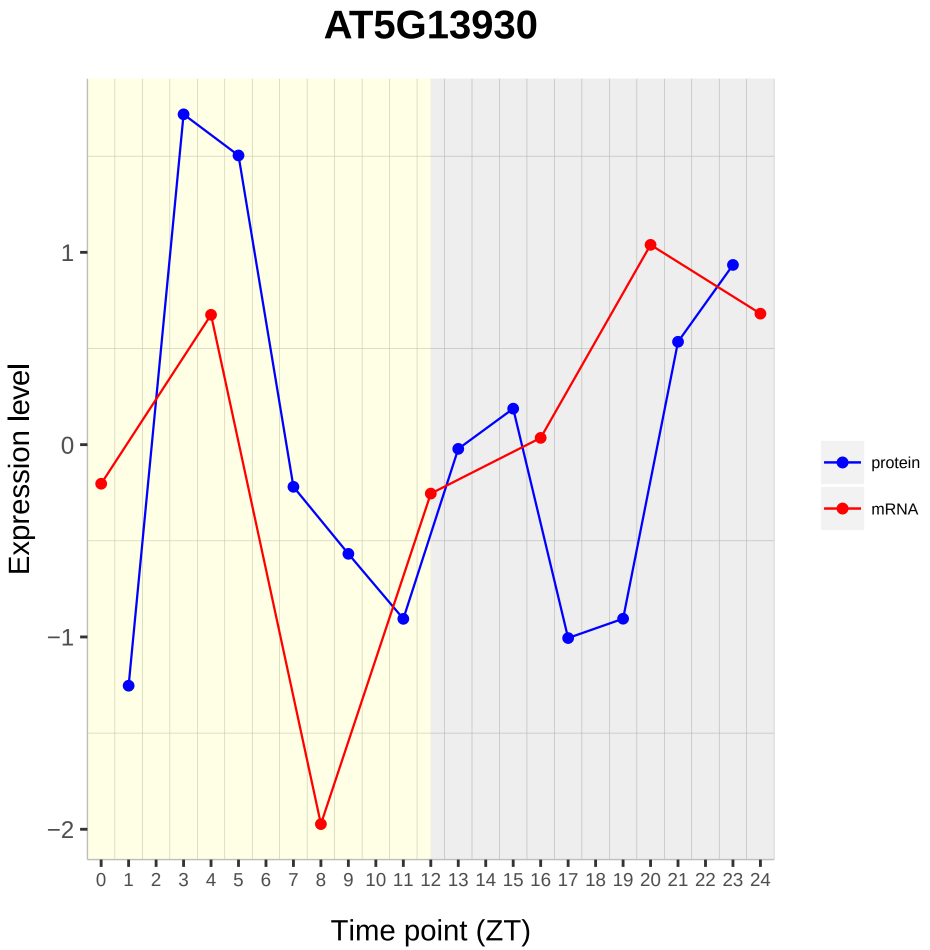

AT5G13930
Expression level
1
0
−1
−2
0 1 2 3 4 5 6 7 8 9 10 11 12 13 14 15 16 17 18 19 20 21 22 23 24
Time point (ZT)
protein
mRNA

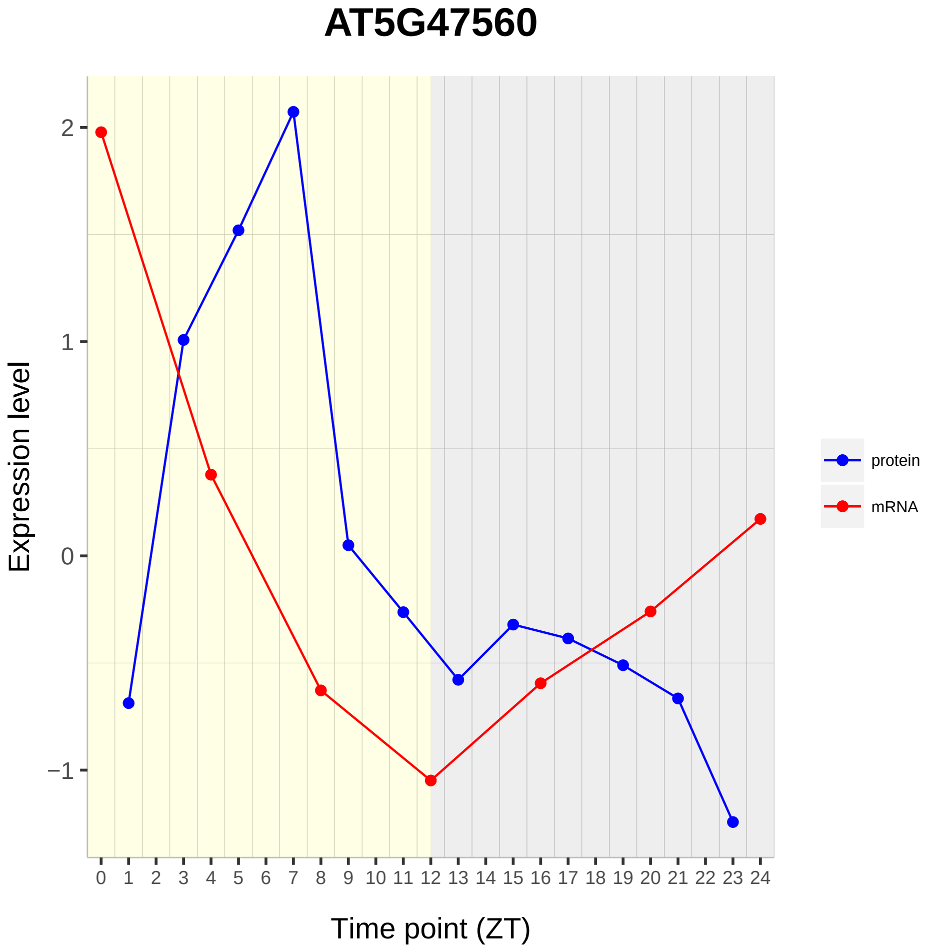
AT5G47560
Expression level
2
1
0
−1
Time point (ZT)
0 1 2 3 4 5 6 7 8 9 10 11 12 13 14 15 16 17 18 19 20 21 22 23 24
protein
mRNA

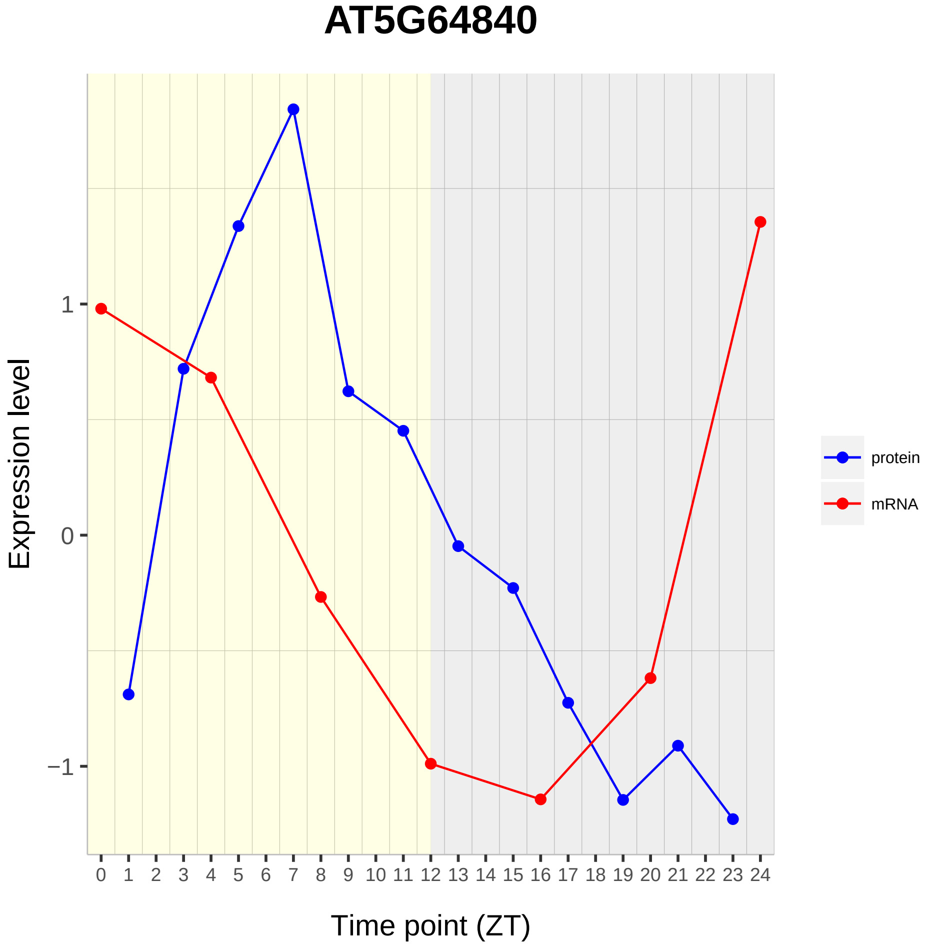

AT5G64840
Expression level
1
0
−1
0 1 2 3 4 5 6 7 8 9 10 11 12 13 14 15 16 17 18 19 20 21 22 23 24
Time point (ZT)
protein
mRNA
