## Supplementary material for "Diurnal Dynamics of the Arabidopsis Rosette Proteome and Phosphoproteome": SuppData6

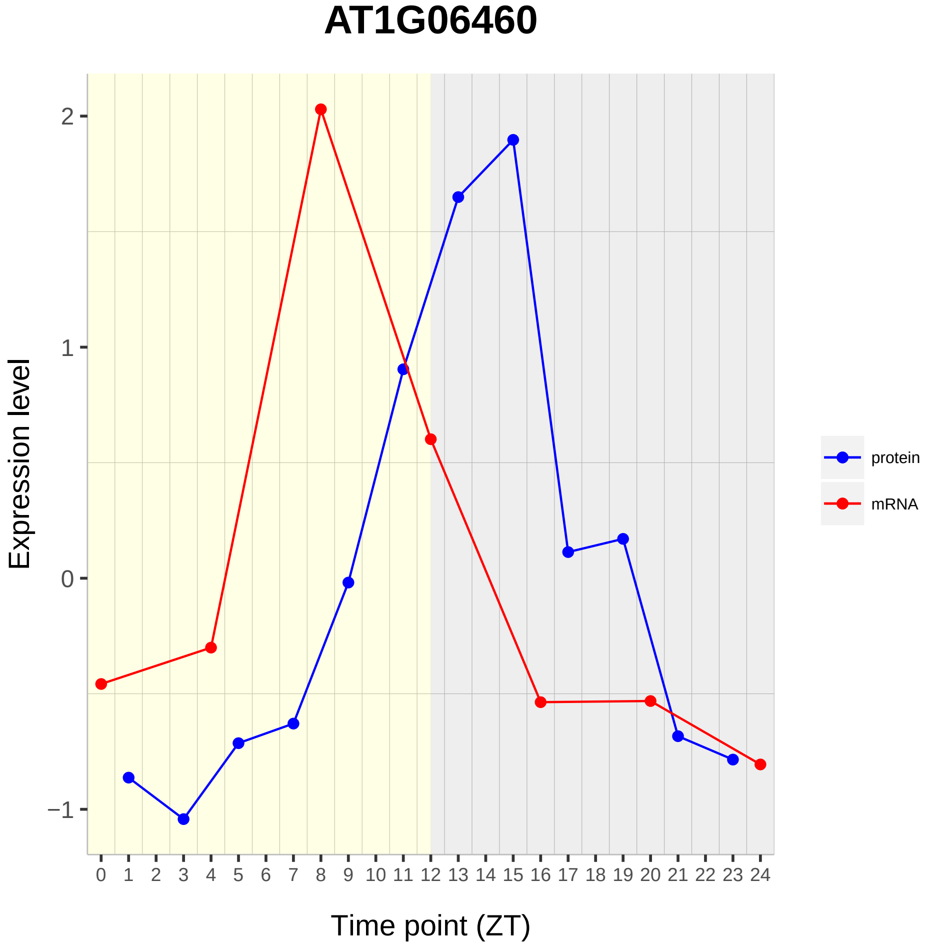
AT1G06460
Expression level
2
1
0
−1
0 1 2 3 4 5 6 7 8 9 10 11 12 13 14 15 16 17 18 19 20 21 22 23 24
Time point (ZT)
protein
mRNA

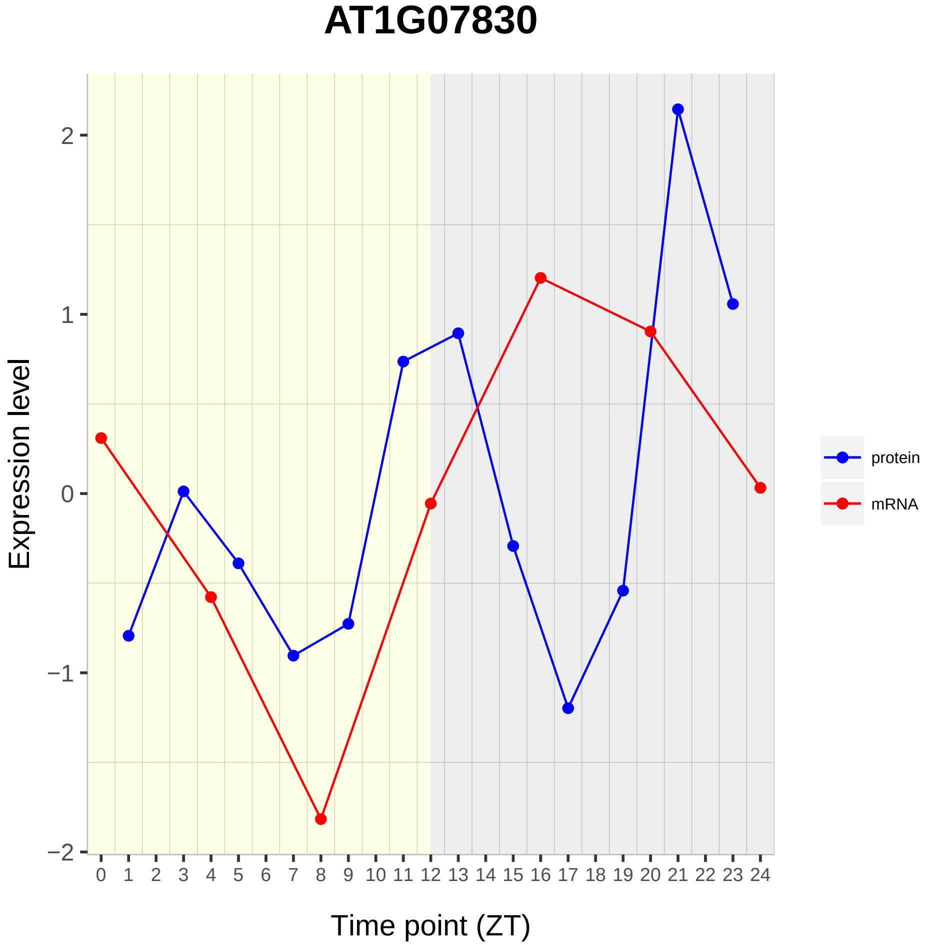

AT1G07830
Expression level
2
1
0
−1
−2
0 1 2 3 4 5 6 7 8 9 10 11 12 13 14 15 16 17 18 19 20 21 22 23 24
Time point (ZT)
protein
mRNA

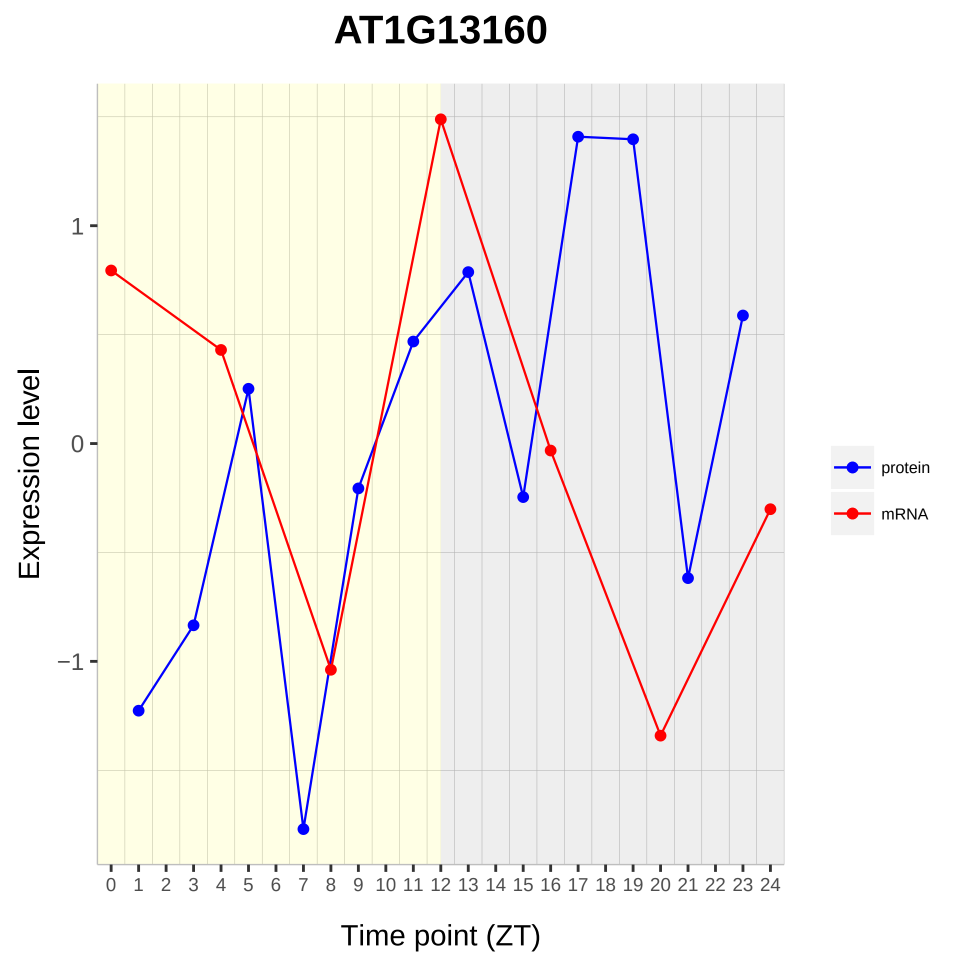
AT1G13160
Expression level
1
0
−1
0 1 2 3 4 5 6 7 8 9 10 11 12 13 14 15 16 17 18 19 20 21 22 23 24
Time point (ZT)
protein
mRNA

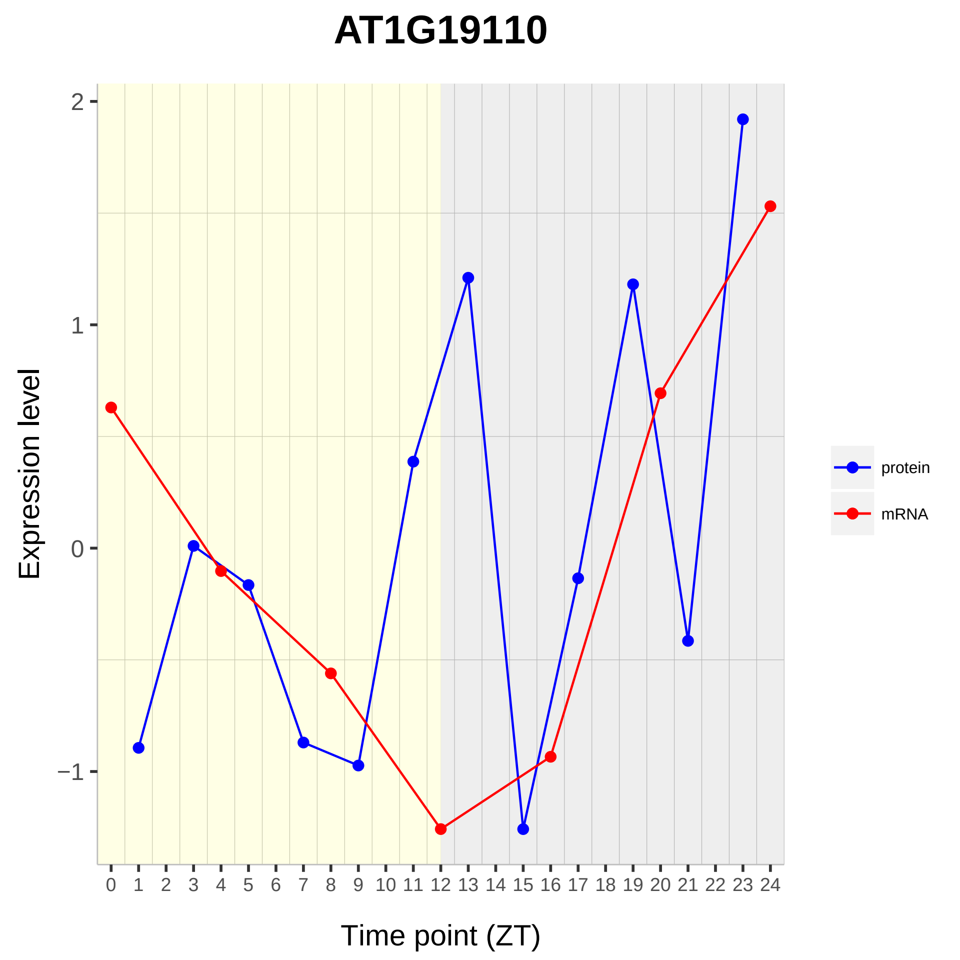

AT1G19110
Expression level
Time point (ZT)
2
1
0
−1
0 1 2 3 4 5 6 7 8 9 10 11 12 13 14 15 16 17 18 19 20 21 22 23 24
protein
mRNA

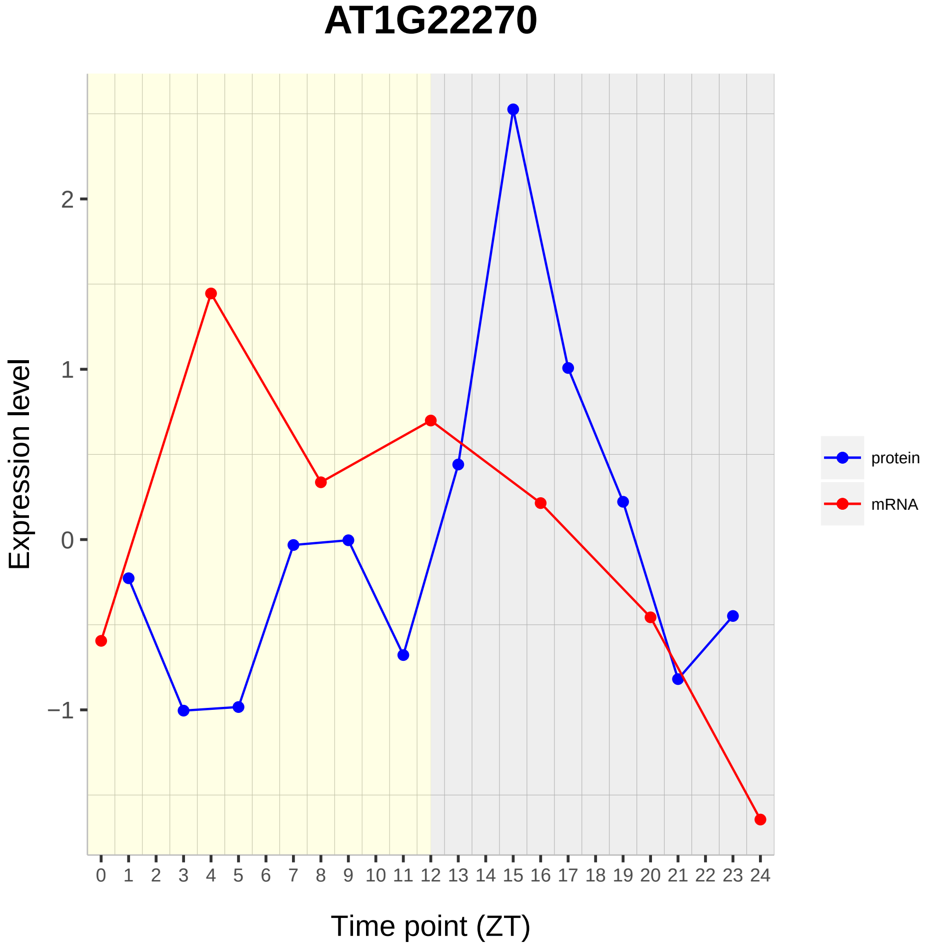
AT1G22270
Expression level
2
1
0
−1
0 1 2 3 4 5 6 7 8 9 10 11 12 13 14 15 16 17 18 19 20 21 22 23 24
Time point (ZT)
protein
mRNA

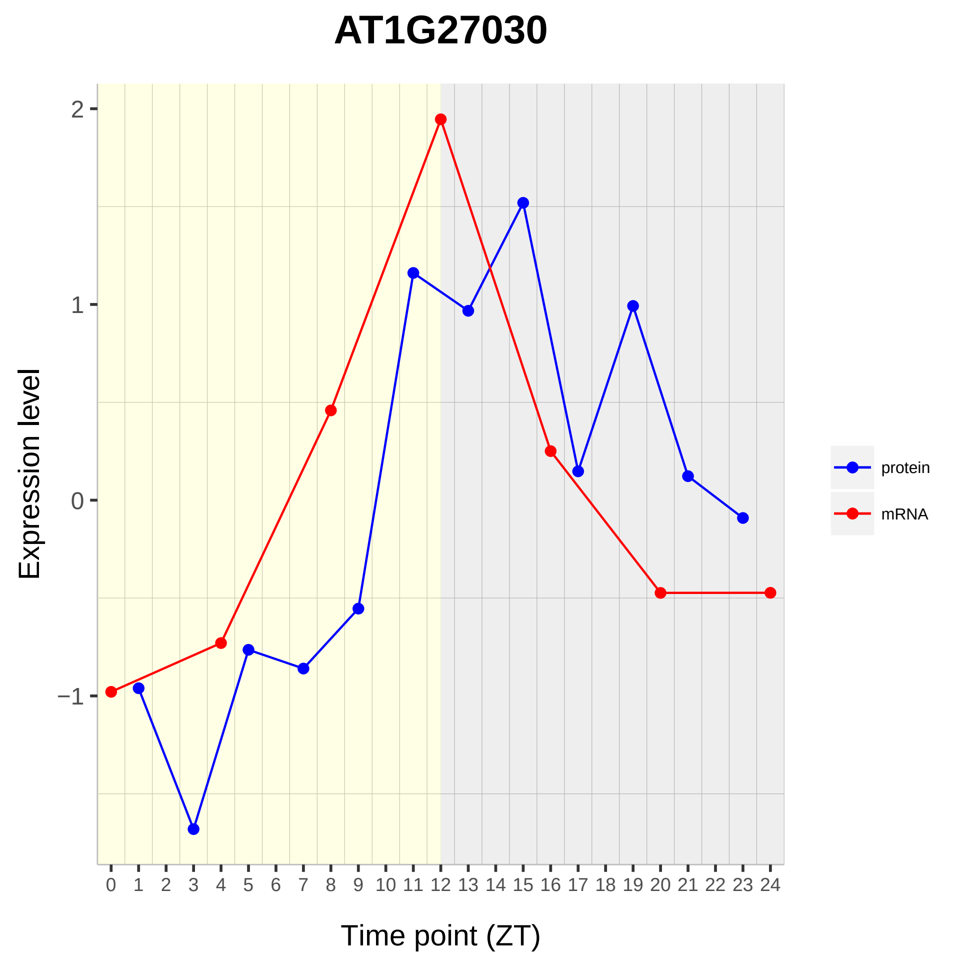

AT1G27030
Expression level
Time point (ZT)
2
1
0
−1
protein
mRNA

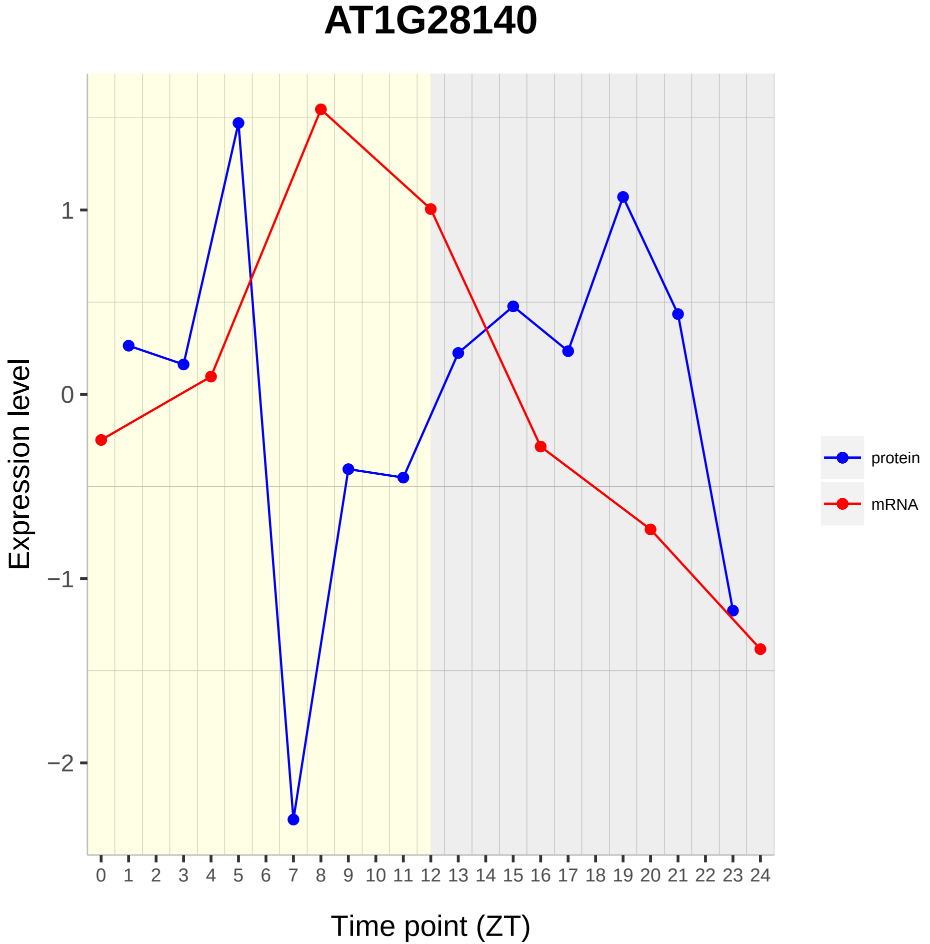
AT1G28140
Expression level
1
0
−1
−2
Time point (ZT)
protein
mRNA

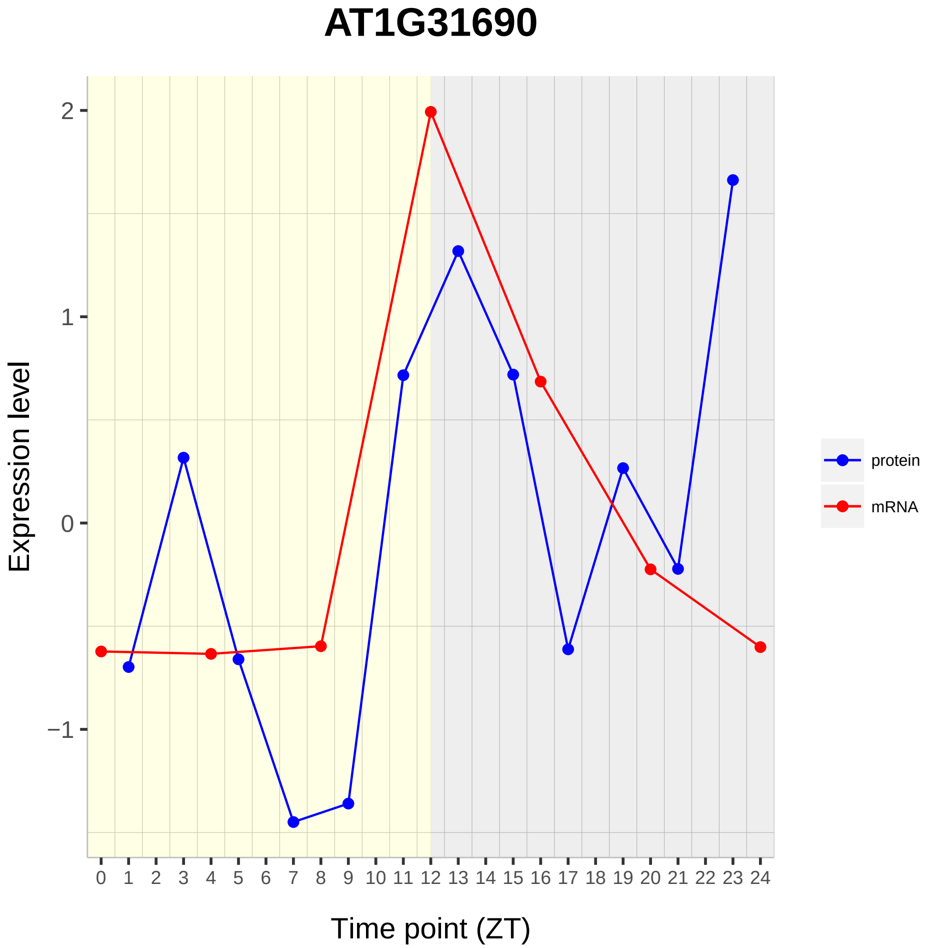

AT1G31690
Expression level
Time point (ZT)
2
1
0
−1
0 1 2 3 4 5 6 7 8 9 10 11 12 13 14 15 16 17 18 19 20 21 22 23 24
protein
mRNA

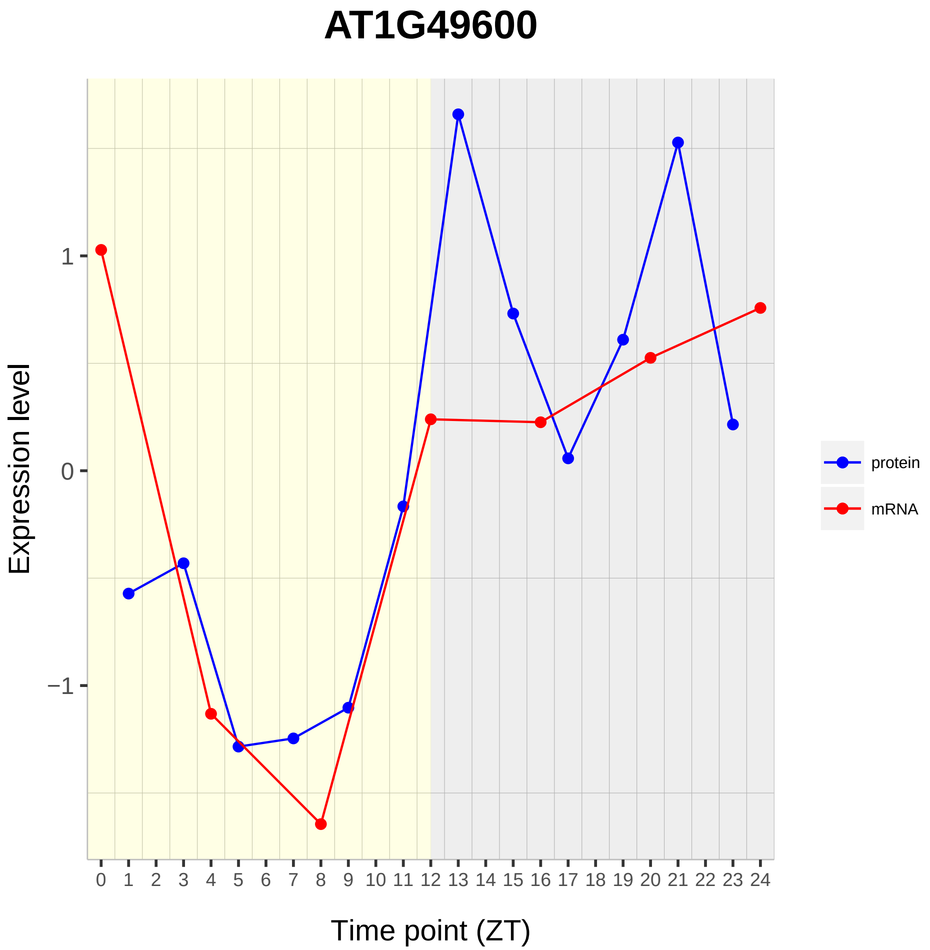

AT1G49600
Expression level
1
0
−1
0 1 2 3 4 5 6 7 8 9 10 11 12 13 14 15 16 17 18 19 20 21 22 23 24
Time point (ZT)
protein
mRNA

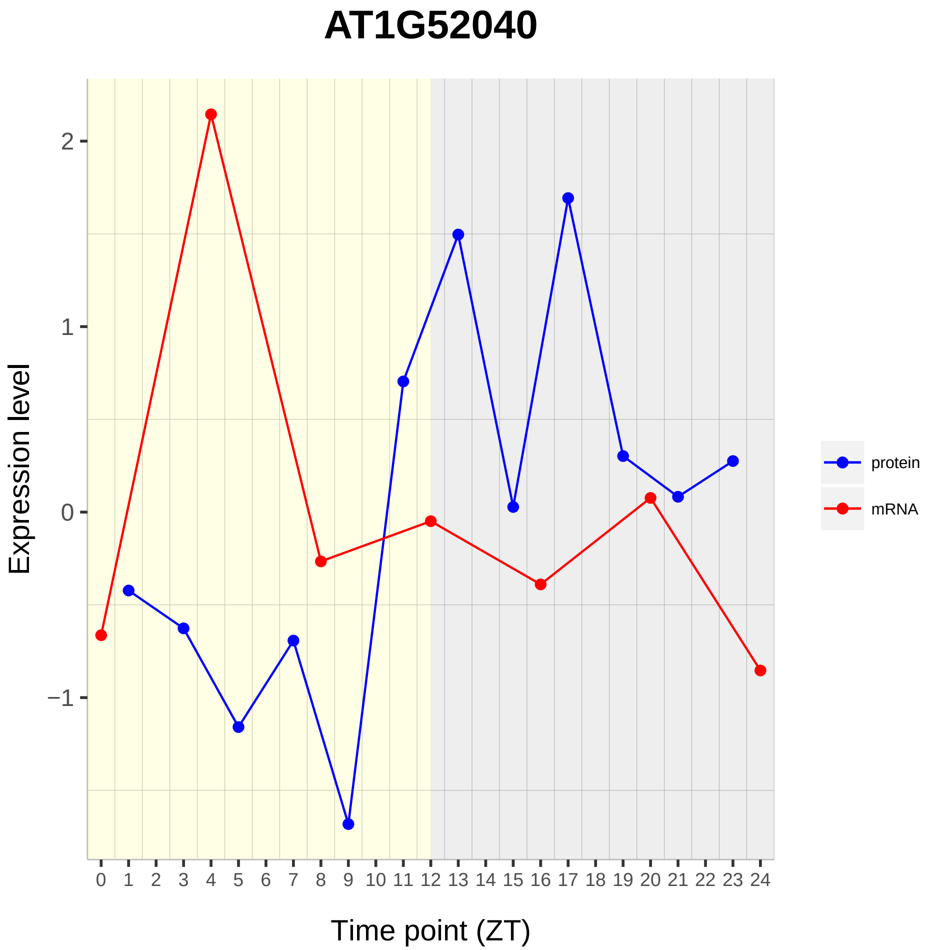

AT1G52040
Expression level
2
1
0
−1
0 1 2 3 4 5 6 7 8 9 10 11 12 13 14 15 16 17 18 19 20 21 22 23 24
Time point (ZT)
protein
mRNA

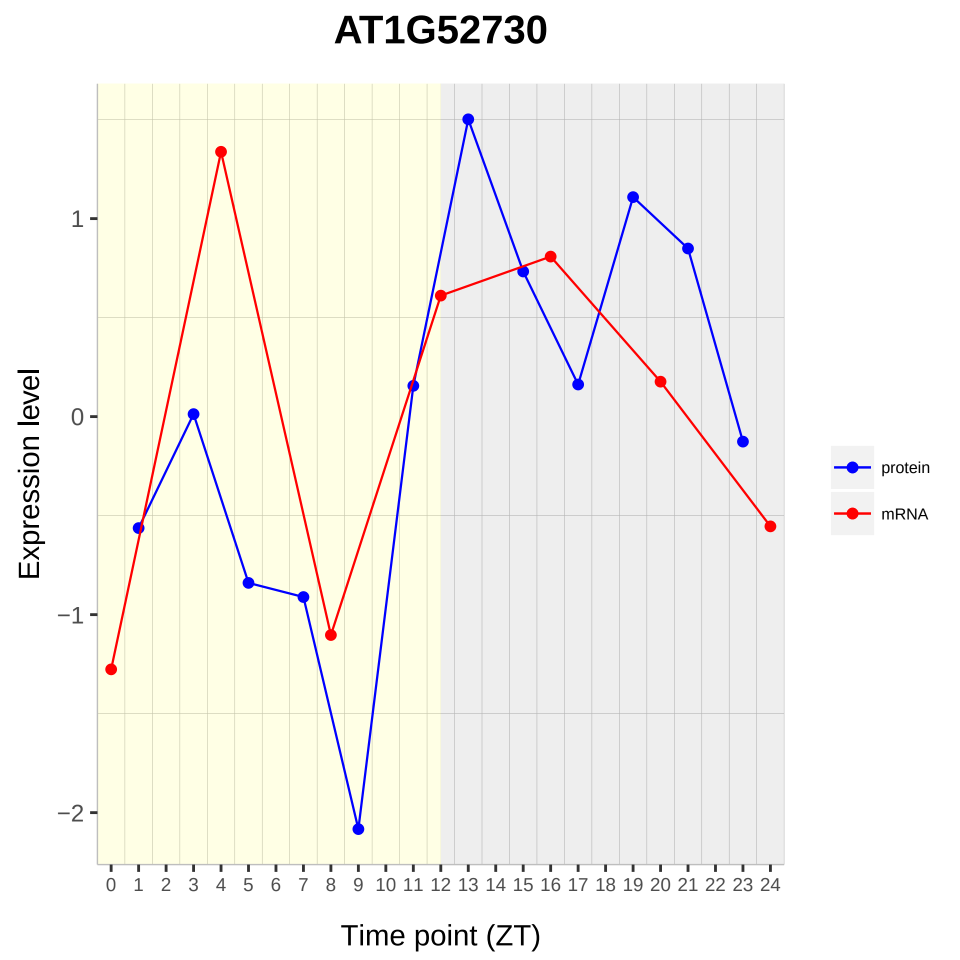
AT1G52730
Expression level
1
0
−1
−2
Time point (ZT)
protein
mRNA

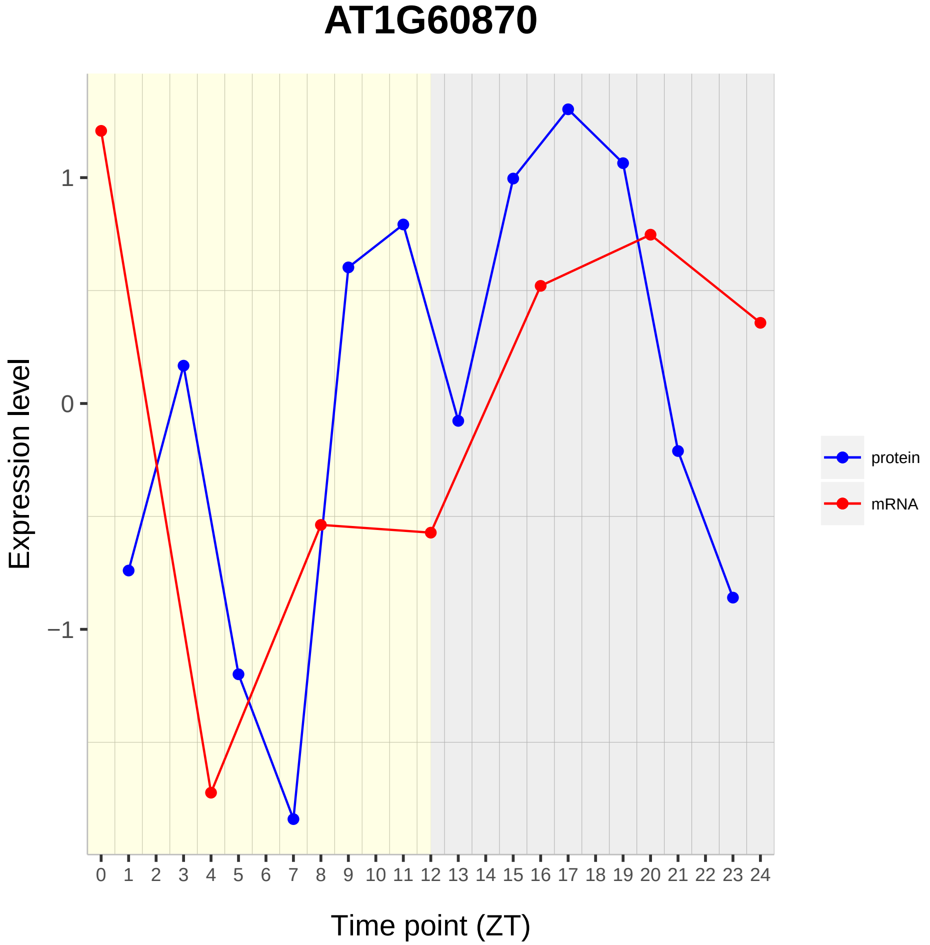
AT1G60870
Expression level
1
0
−1
0 1 2 3 4 5 6 7 8 9 10 11 12 13 14 15 16 17 18 19 20 21 22 23 24
Time point (ZT)
protein
mRNA

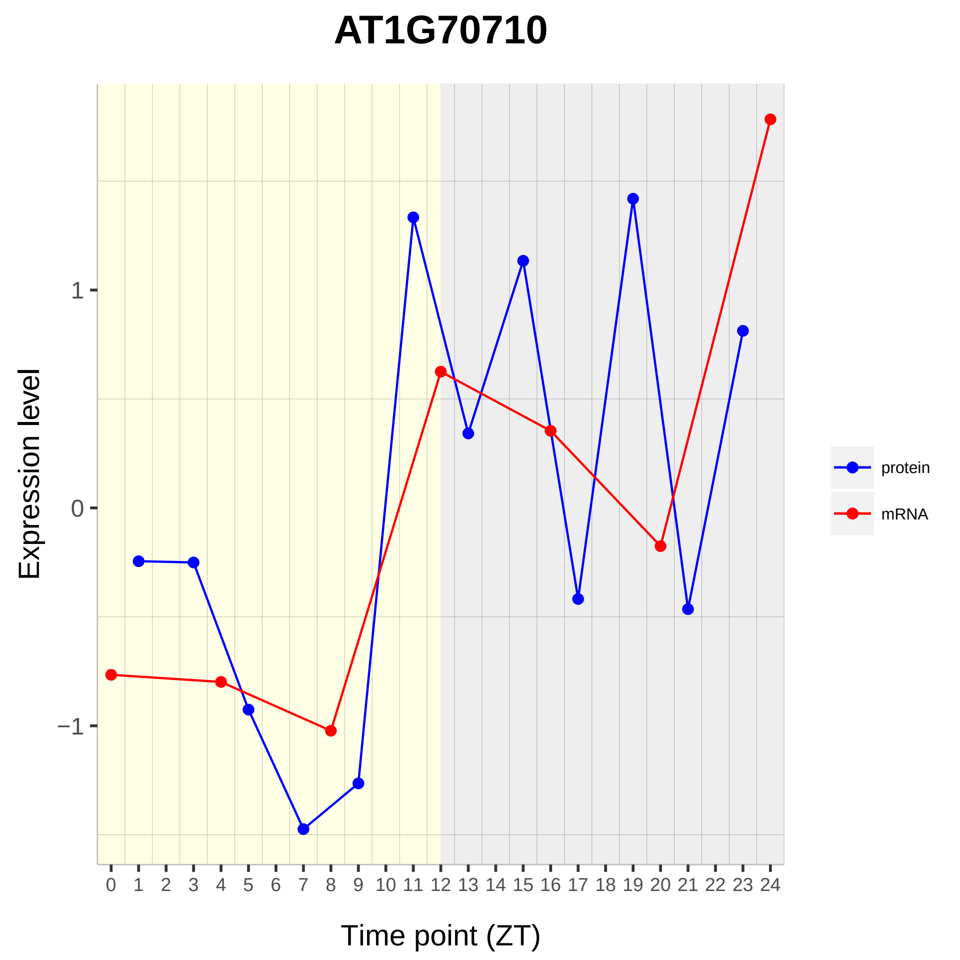
AT1G70710
Expression level
1
0
−1
Time point (ZT)
protein
mRNA

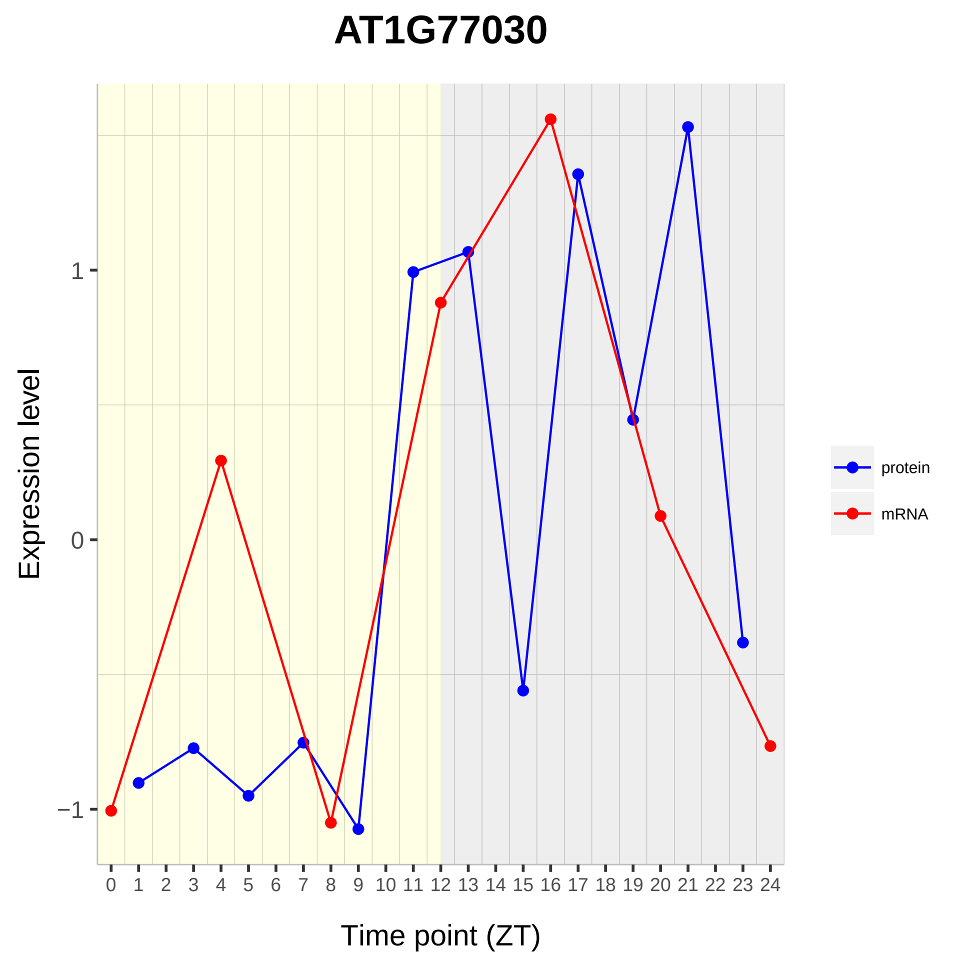

AT1G77030
Expression level
1
0
−1
0 1 2 3 4 5 6 7 8 9 10 11 12 13 14 15 16 17 18 19 20 21 22 23 24
Time point (ZT)
protein
mRNA

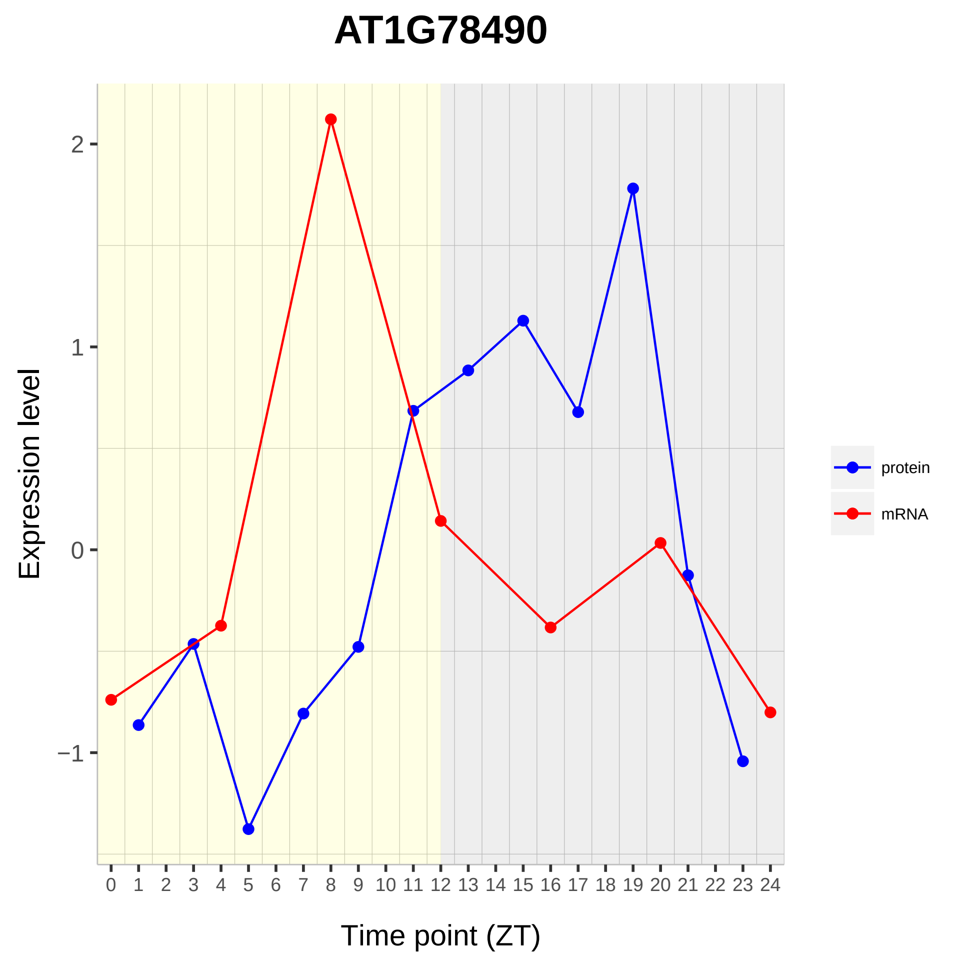

AT1G78490
Expression level
2
1
0
−1
Time point (ZT)
protein
mRNA

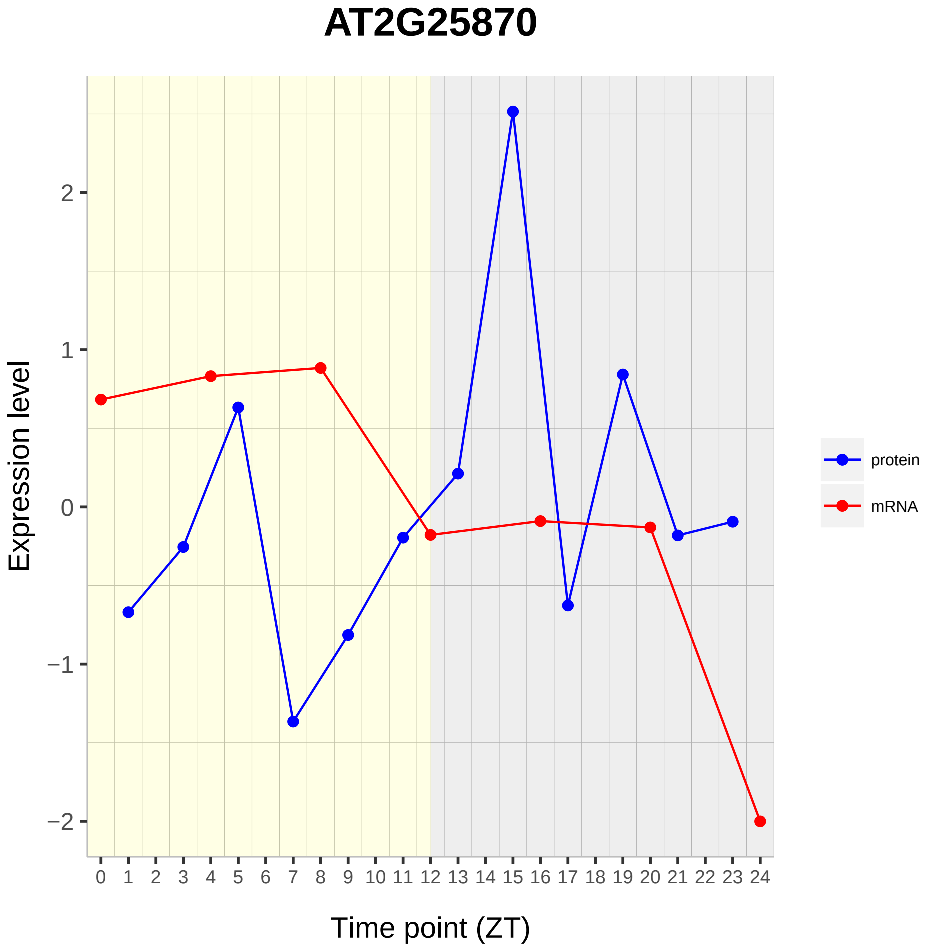

AT2G25870
Expression level
2
1
0
−1
−2
Time point (ZT)
protein
mRNA

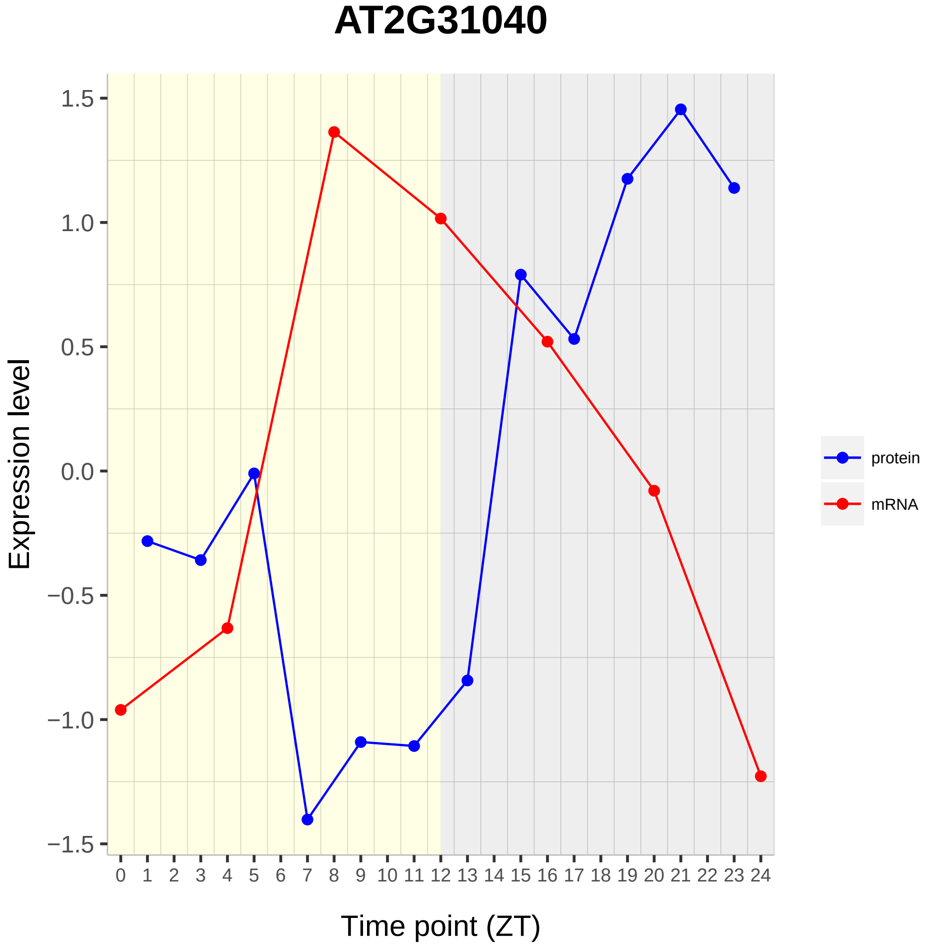
AT2G31040
Expression level
Time point (ZT)
1.5
1.0
0.5
0.0
−0.5
−1.0
−1.5
0 1 2 3 4 5 6 7 8 9 10 11 12 13 14 15 16 17 18 19 20 21 22 23 24
protein
mRNA

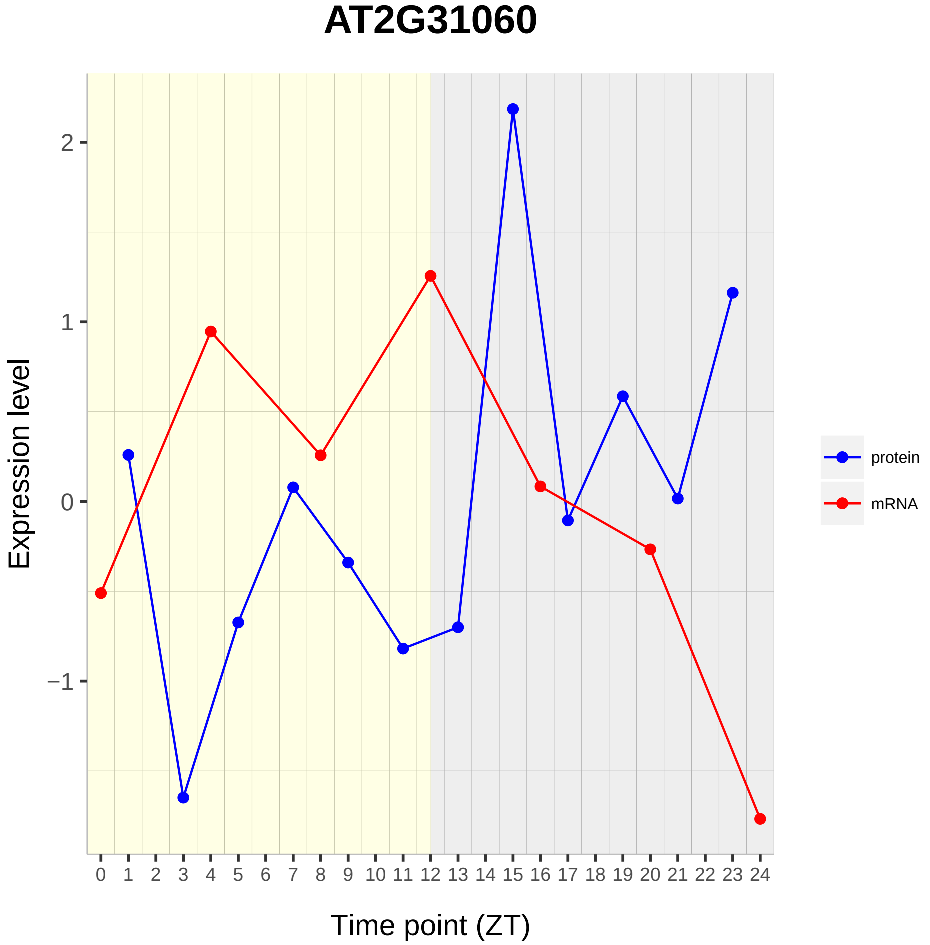
AT2G31060
Expression level
2
1
0
−1
0 1 2 3 4 5 6 7 8 9 10 11 12 13 14 15 16 17 18 19 20 21 22 23 24
Time point (ZT)
protein
mRNA

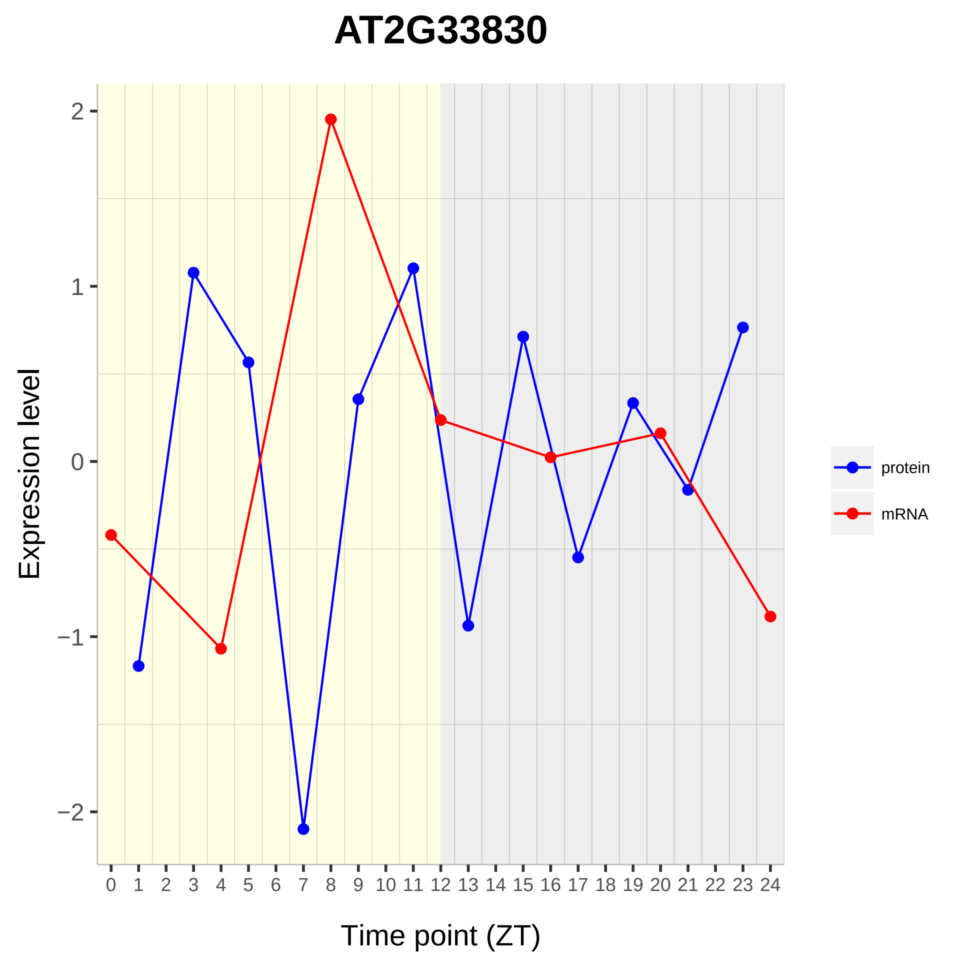
AT2G33830
Expression level
Time point (ZT)
2
1
0
−1
−2
0 1 2 3 4 5 6 7 8 9 10 11 12 13 14 15 16 17 18 19 20 21 22 23 24
protein
mRNA

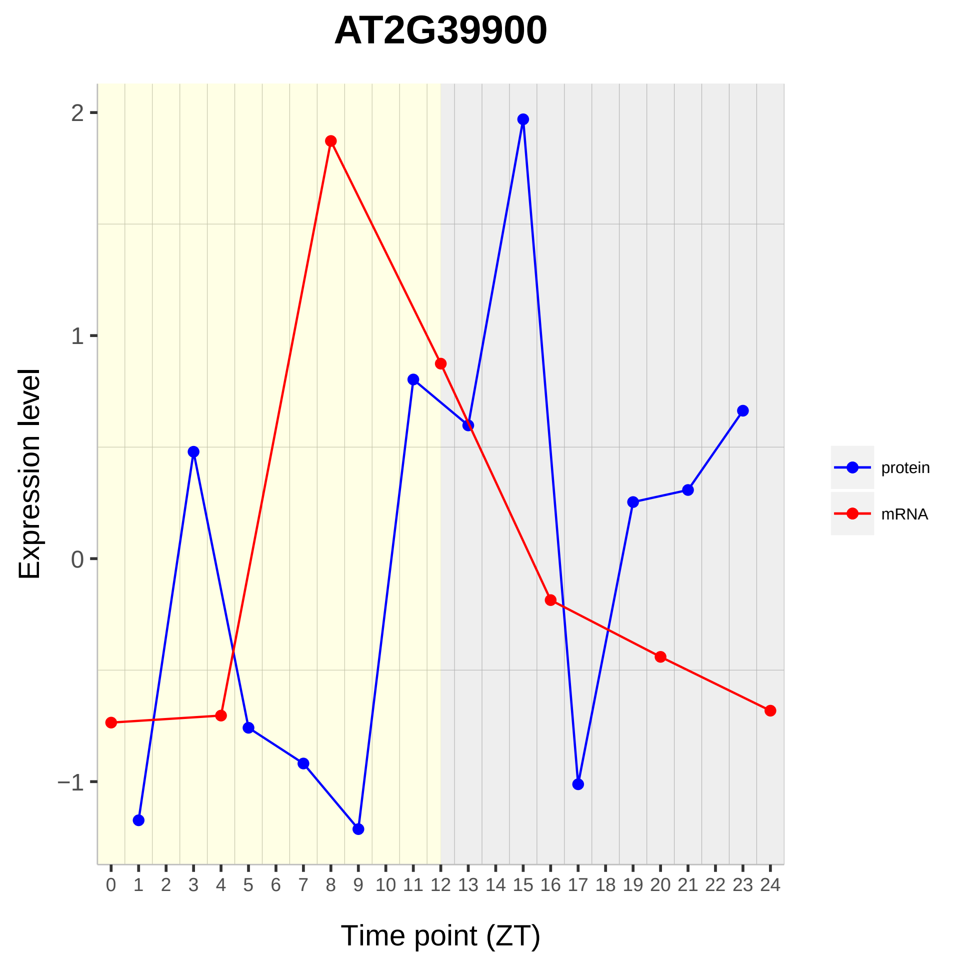

AT2G39900
Expression level
Time point (ZT)
2
1
0
−1
0 1 2 3 4 5 6 7 8 9 10 11 12 13 14 15 16 17 18 19 20 21 22 23 24
protein
mRNA

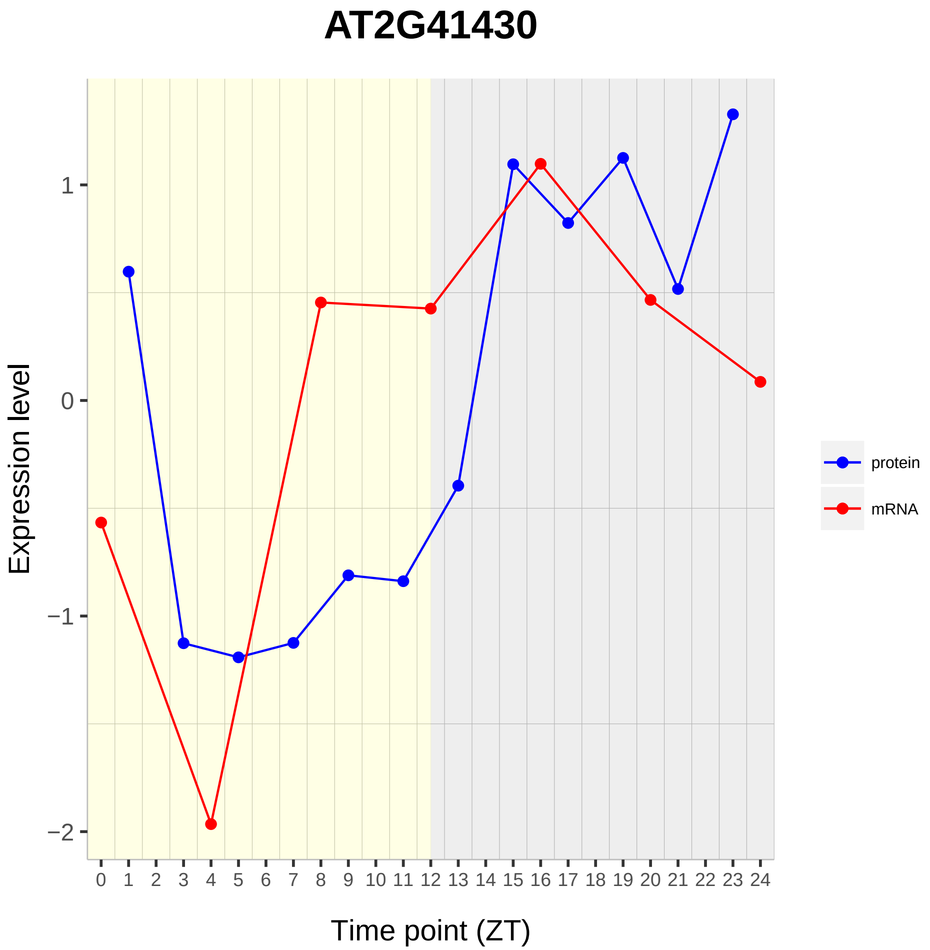

AT2G41430
Expression level
1
0
−1
−2
0 1 2 3 4 5 6 7 8 9 10 11 12 13 14 15 16 17 18 19 20 21 22 23 24
Time point (ZT)
protein
mRNA

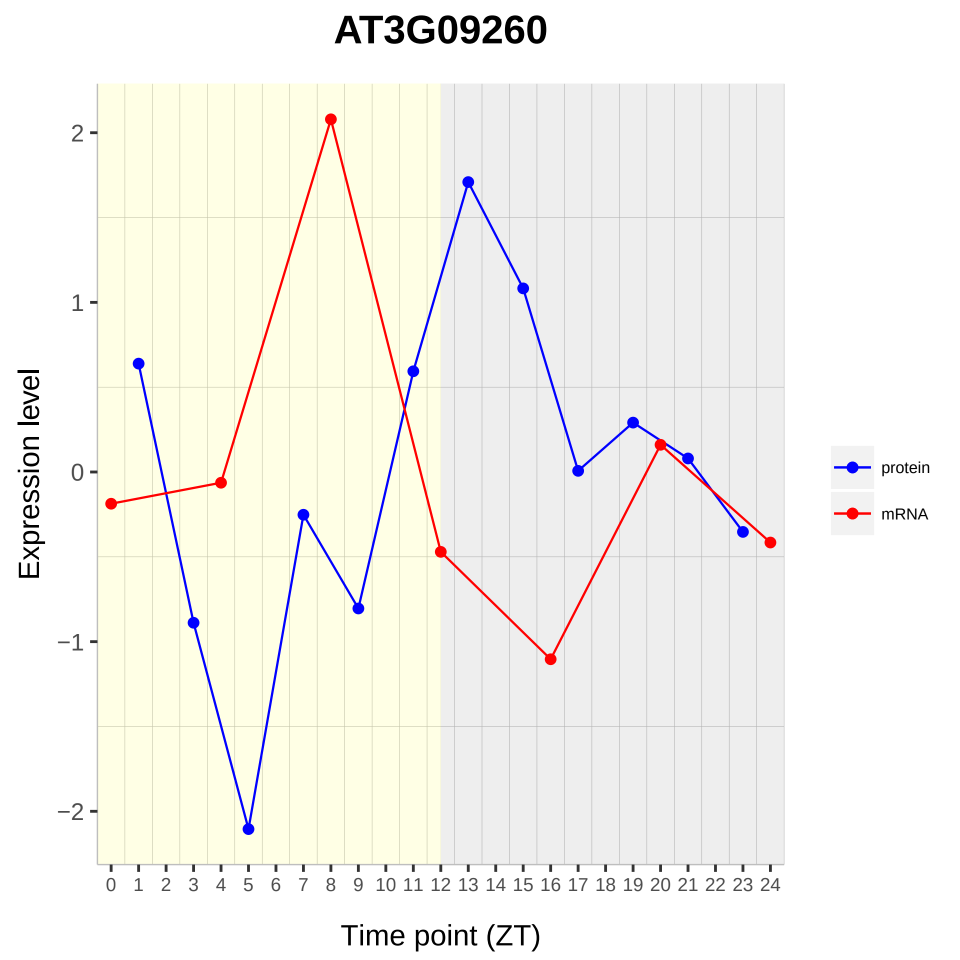

AT3G09260
Expression level
Time point (ZT)
protein
mRNA
2
1
0
−1
−2

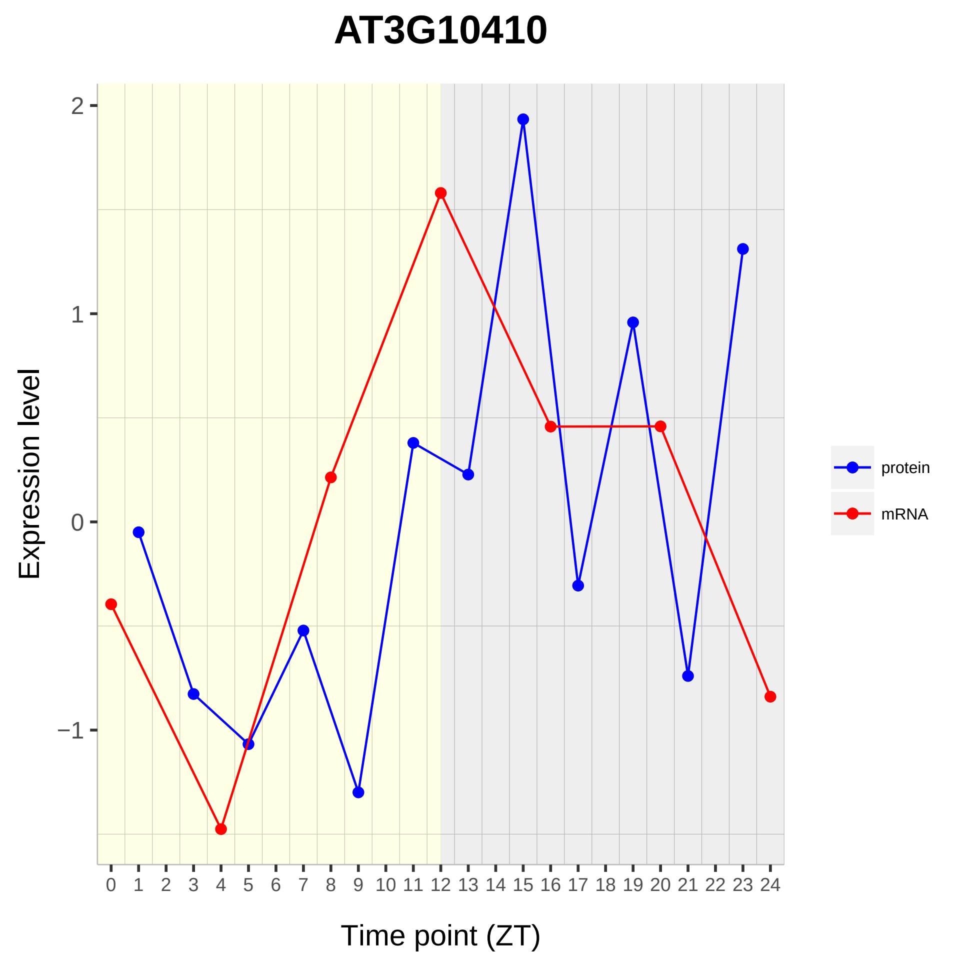

AT3G10410
Expression level
Time point (ZT)
2
1
0
−1
protein
mRNA

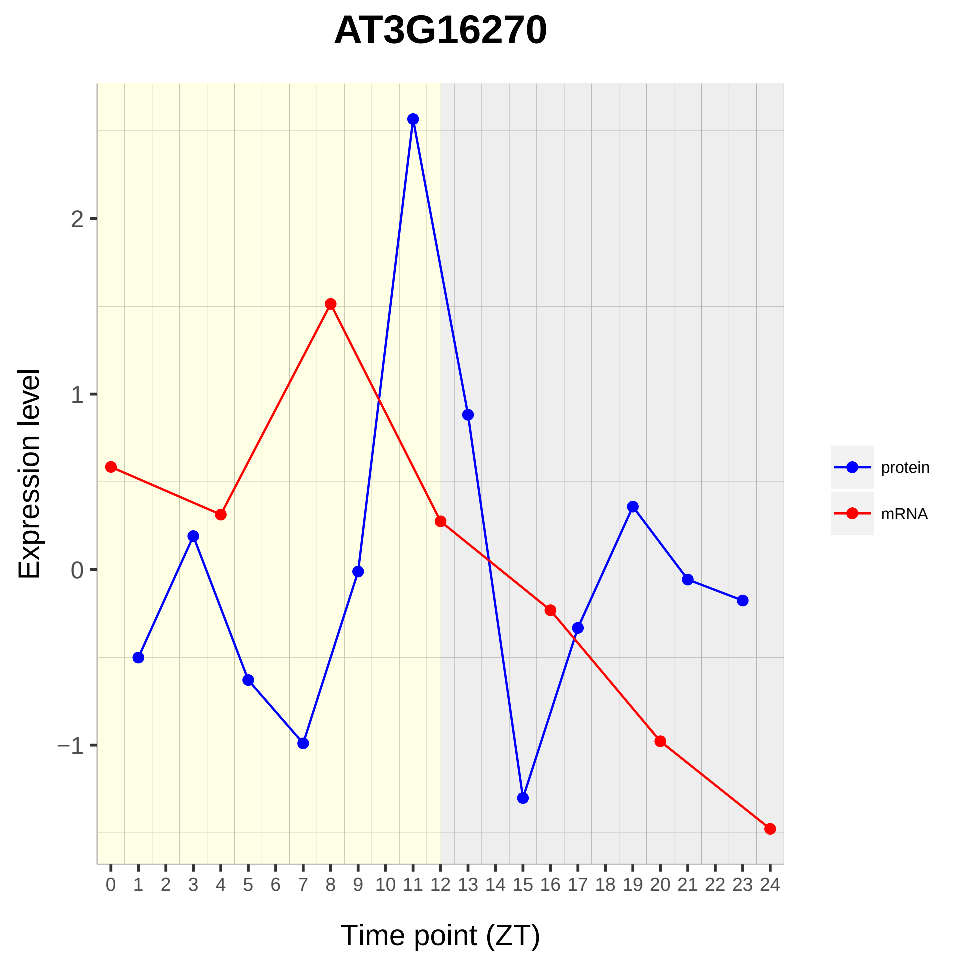

AT3G16270
Expression level
2
1
0
−1
0 1 2 3 4 5 6 7 8 9 10 11 12 13 14 15 16 17 18 19 20 21 22 23 24
Time point (ZT)
protein
mRNA

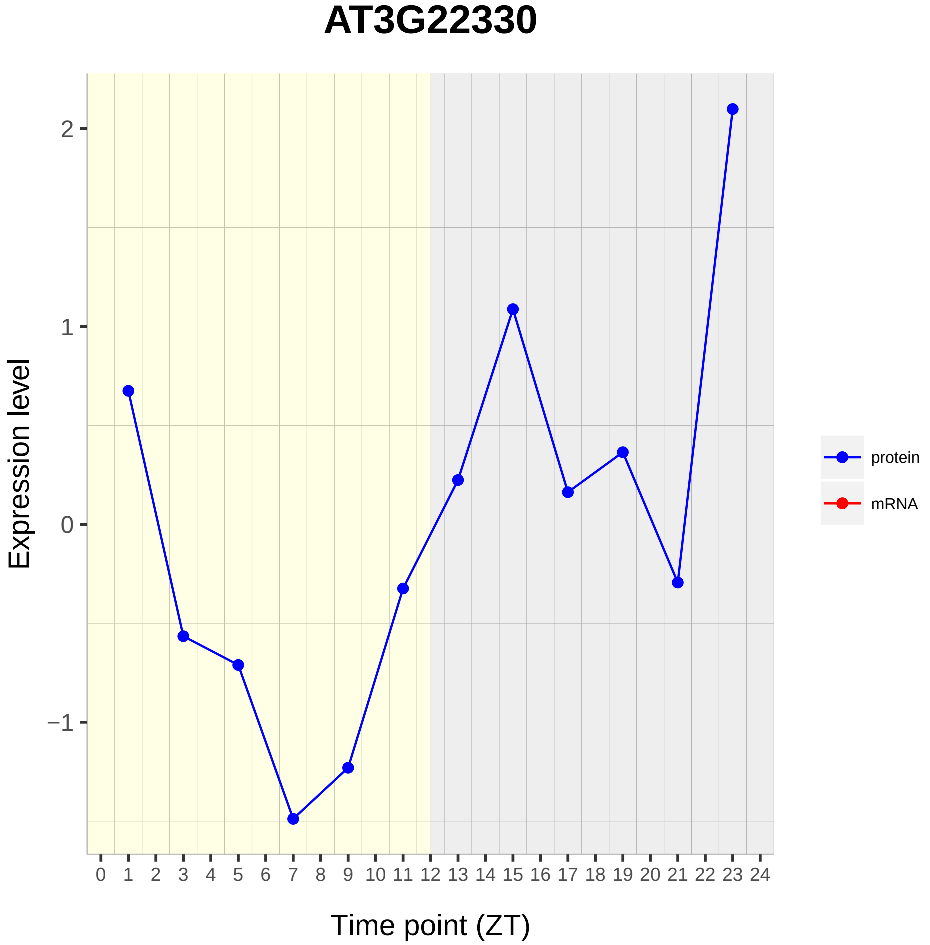

AT3G22330
Expression level
2
1
0
−1
Time point (ZT)
protein
mRNA

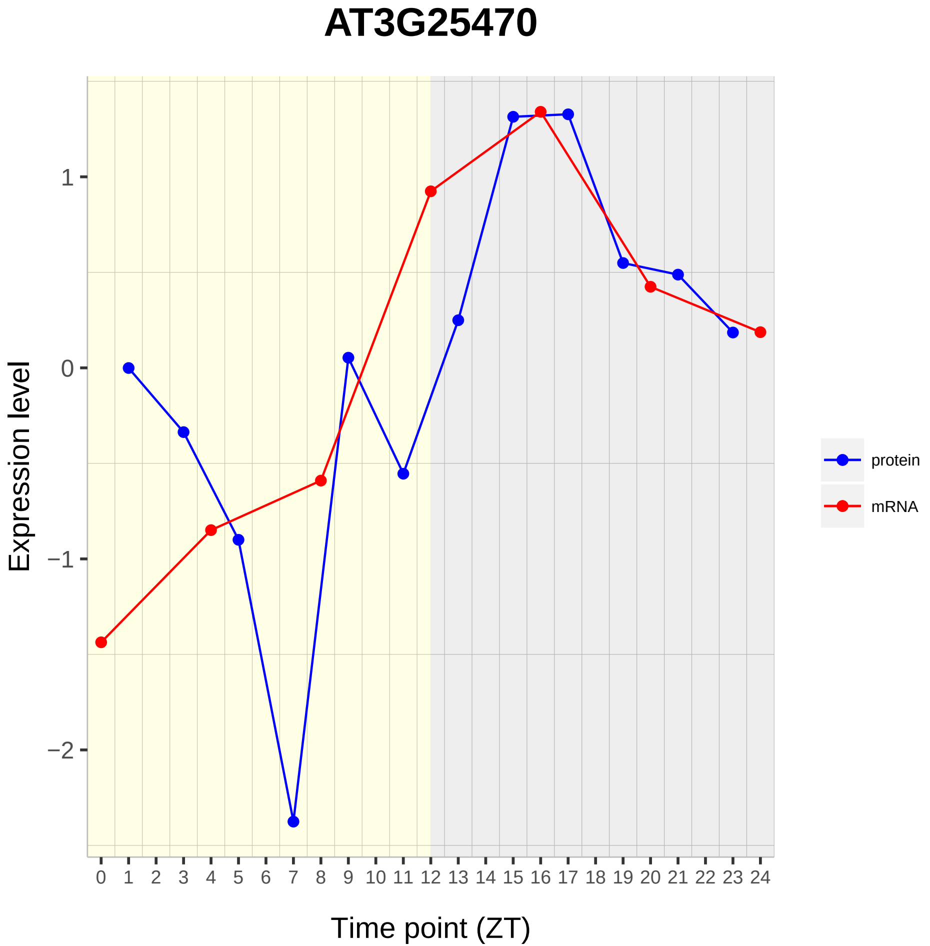

AT3G25470
Expression level
1
0
−1
−2
Time point (ZT)
protein
mRNA

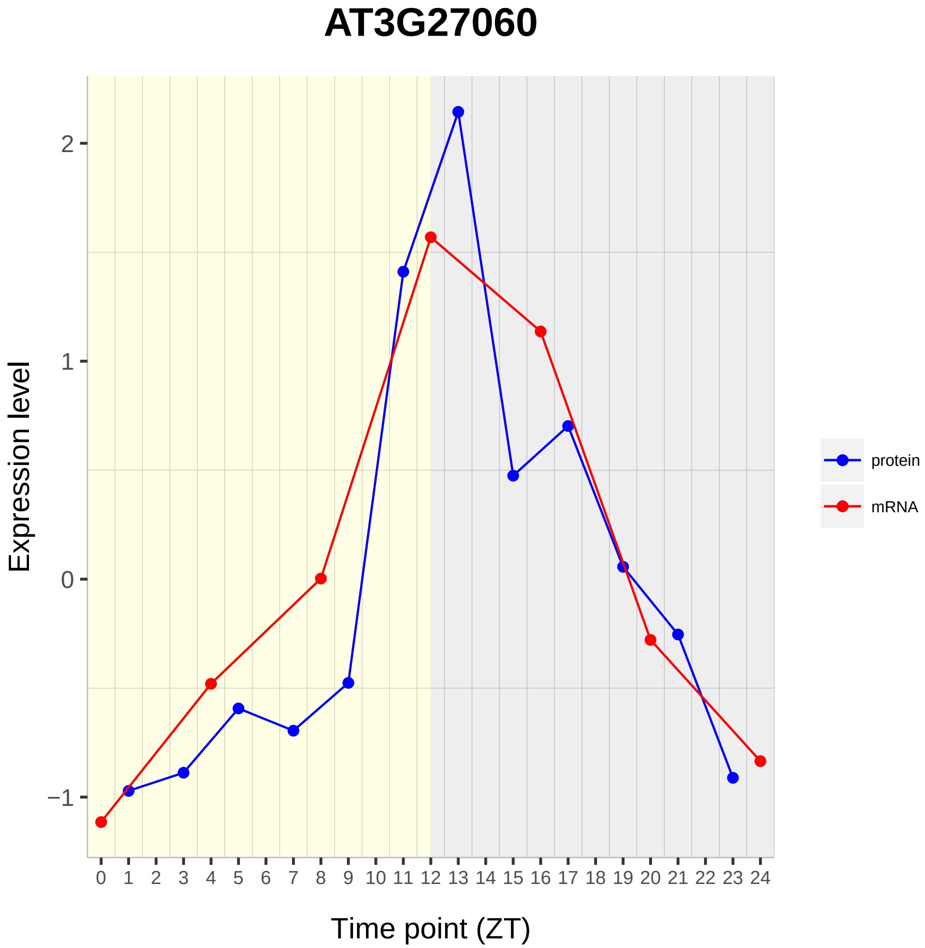

AT3G27060
Expression level
2
1
0
−1
Time point (ZT)
protein
mRNA

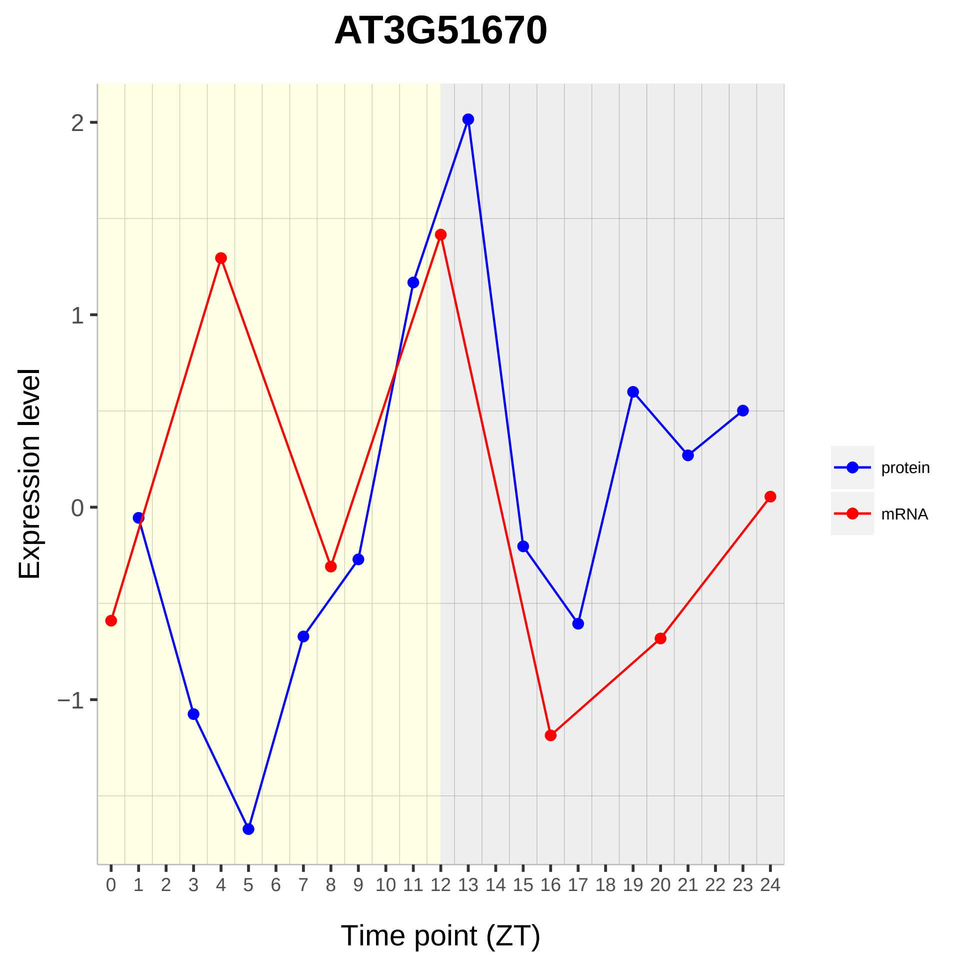

AT3G51670
Expression level
Time point (ZT)
2
1
0
−1
0 1 2 3 4 5 6 7 8 9 10 11 12 13 14 15 16 17 18 19 20 21 22 23 24
protein
mRNA

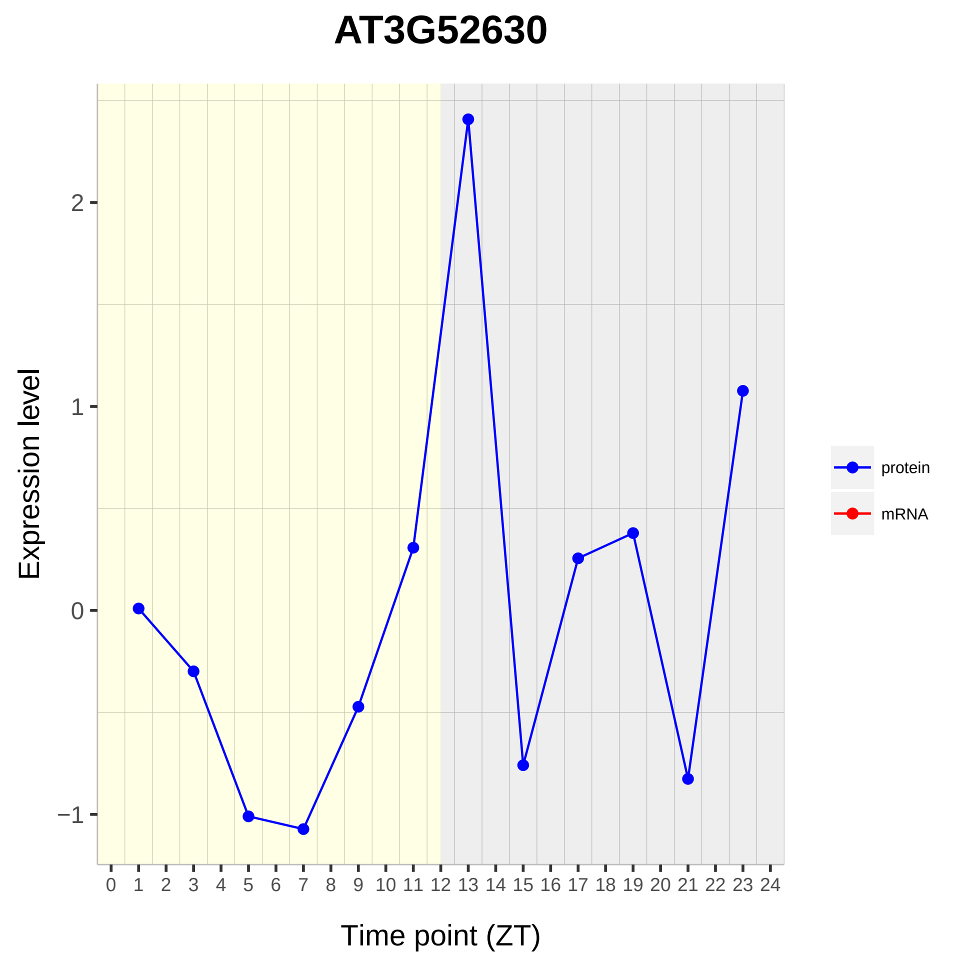

AT3G52630
Expression level
2
1
0
−1
0 1 2 3 4 5 6 7 8 9 10 11 12 13 14 15 16 17 18 19 20 21 22 23 24
Time point (ZT)
protein
mRNA

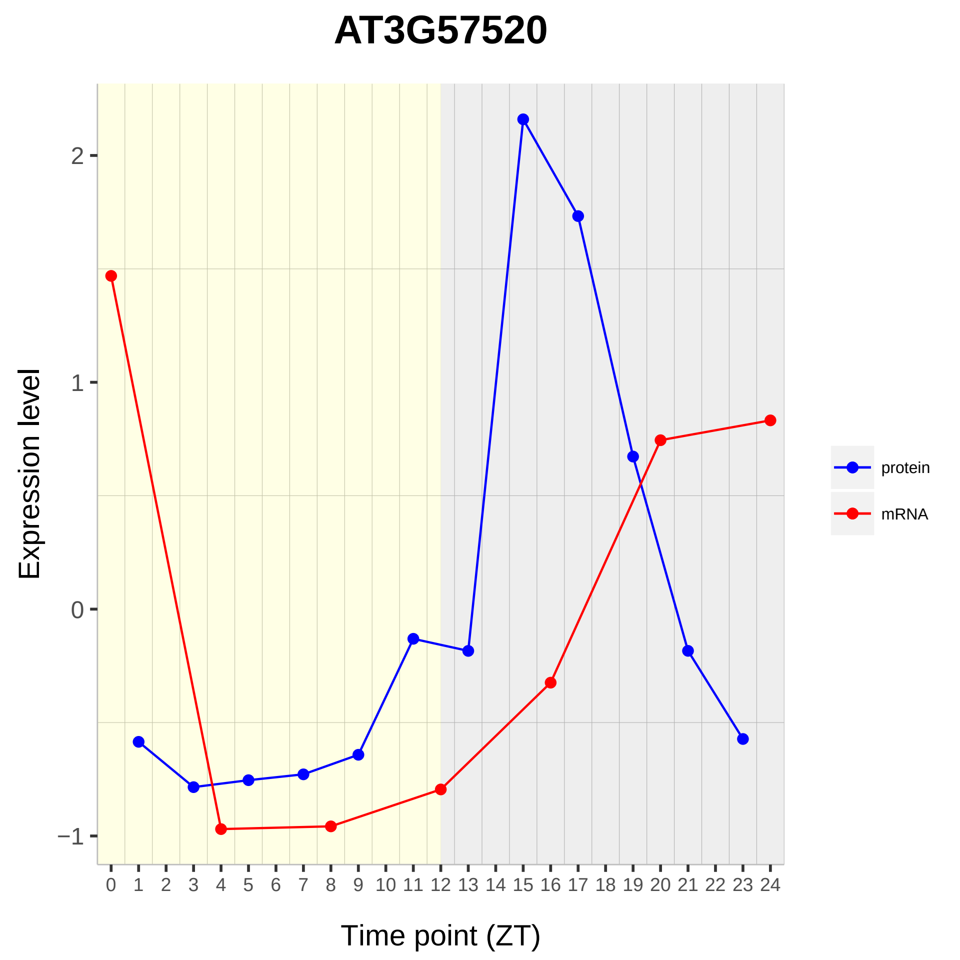
AT3G57520
Expression level
2
1
0
−1
0 1 2 3 4 5 6 7 8 9 10 11 12 13 14 15 16 17 18 19 20 21 22 23 24
Time point (ZT)
protein
mRNA

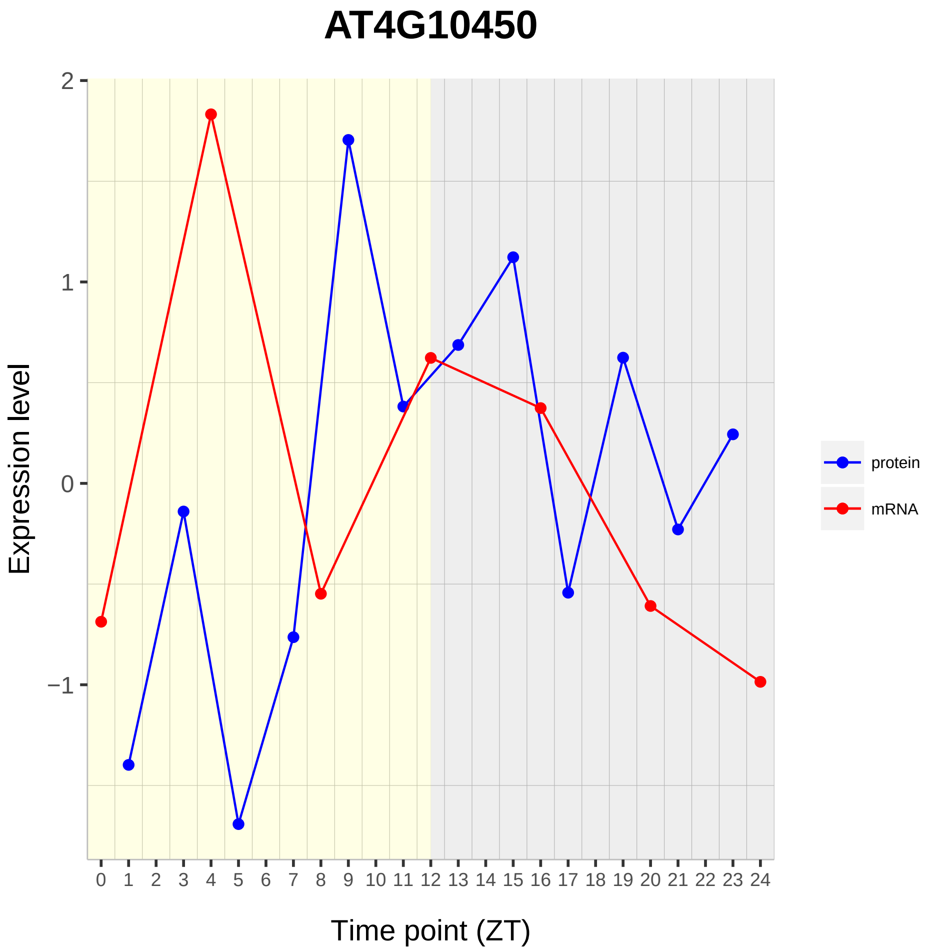
AT4G10450
Expression level
Time point (ZT)
protein
mRNA
2
1
0
−1

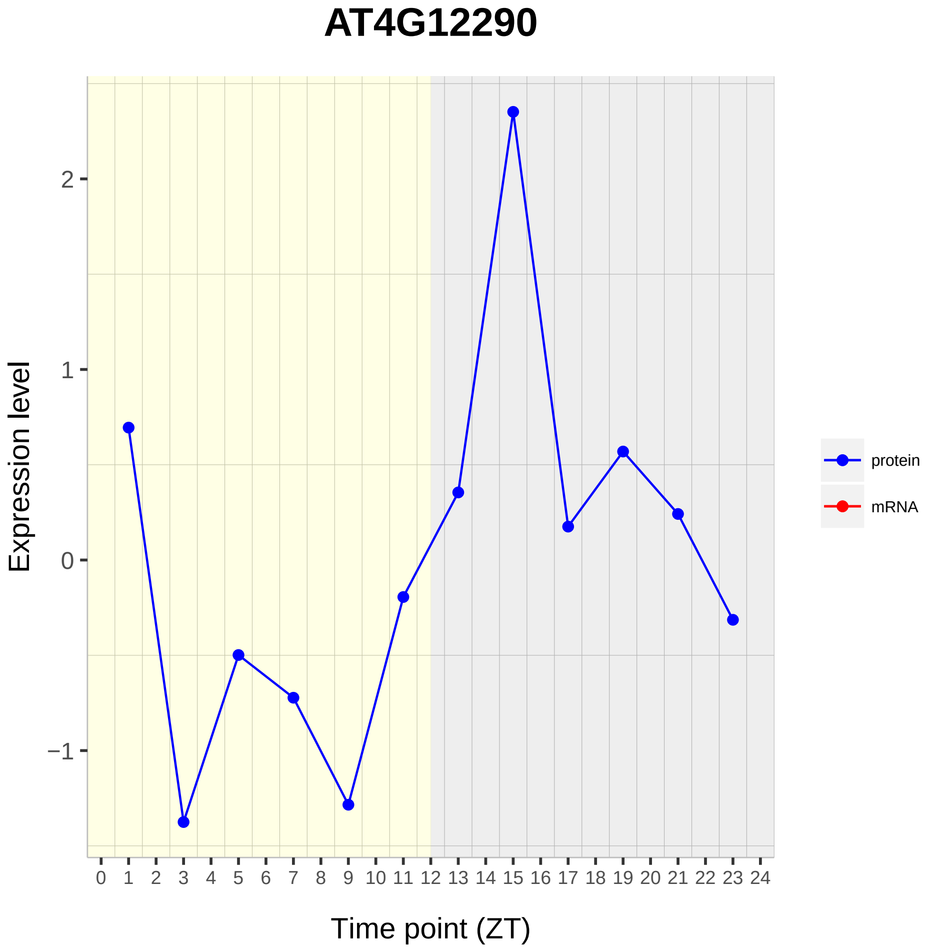
AT4G12290
Expression level
Time point (ZT)
protein
mRNA
2
1
0
−1
0 1 2 3 4 5 6 7 8 9 10 11 12 13 14 15 16 17 18 19 20 21 22 23 24

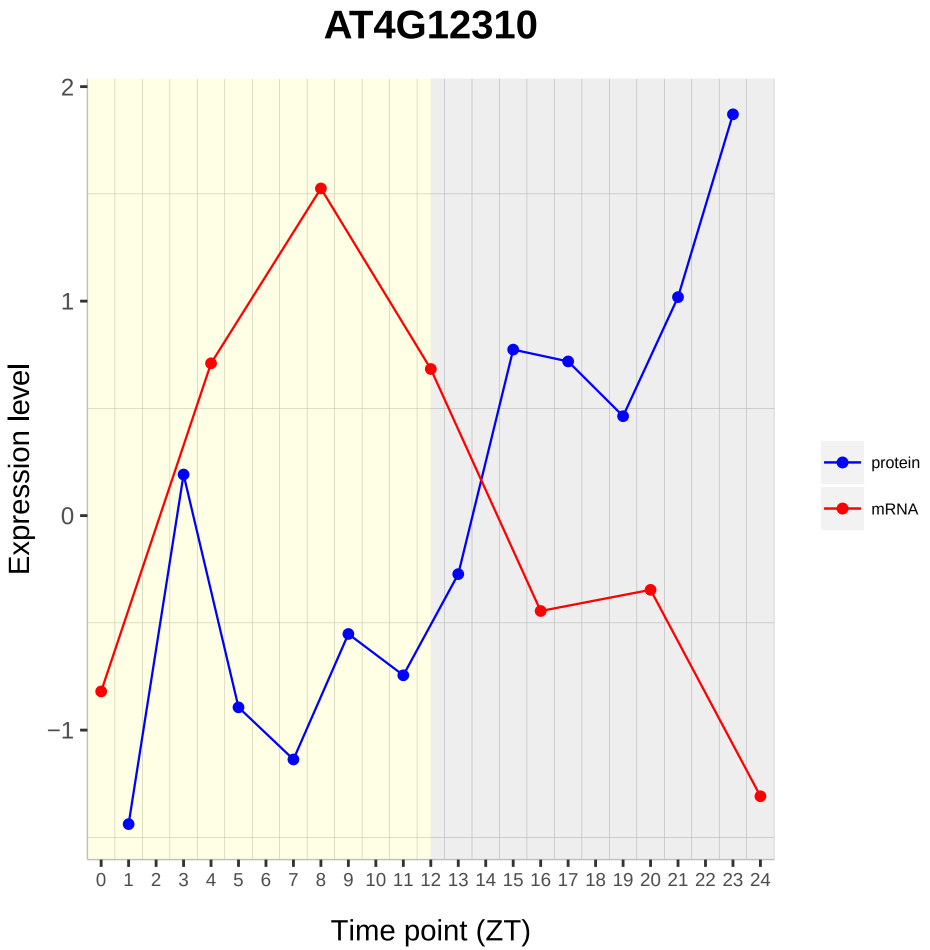

AT4G12310
Expression level
Time point (ZT)
2
1
0
−1
0 1 2 3 4 5 6 7 8 9 10 11 12 13 14 15 16 17 18 19 20 21 22 23 24
protein
mRNA

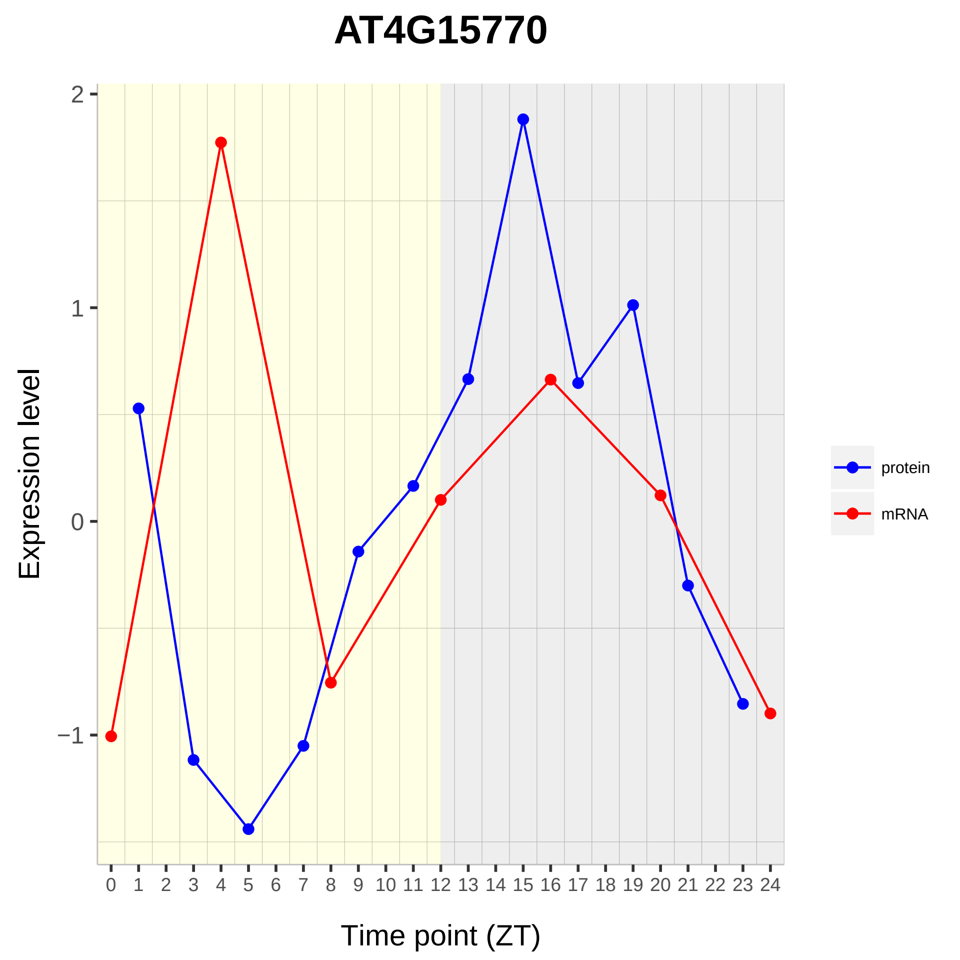
AT4G15770
Expression level
Time point (ZT)
protein
mRNA
2
1
0
−1
0 1 2 3 4 5 6 7 8 9 10 11 12 13 14 15 16 17 18 19 20 21 22 23 24

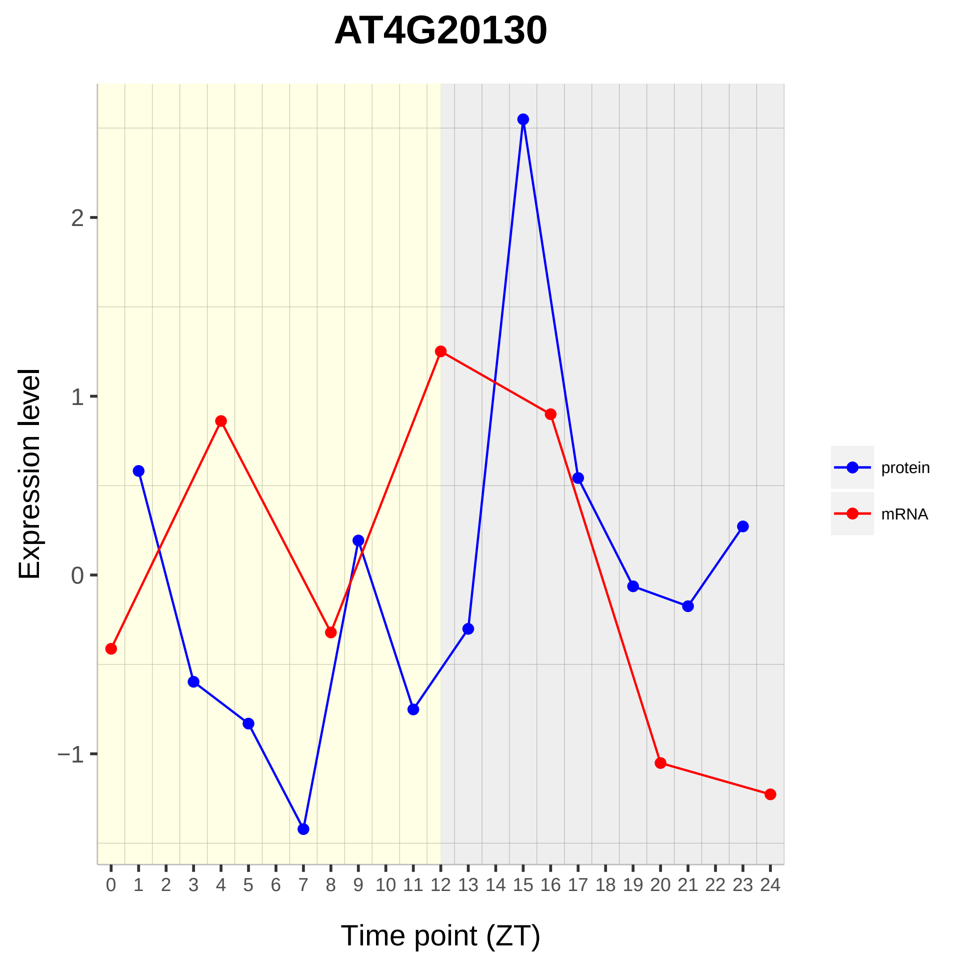
AT4G20130
Expression level
Time point (ZT)
protein
mRNA
2
1
0
−1
0 1 2 3 4 5 6 7 8 9 10 11 12 13 14 15 16 17 18 19 20 21 22 23 24

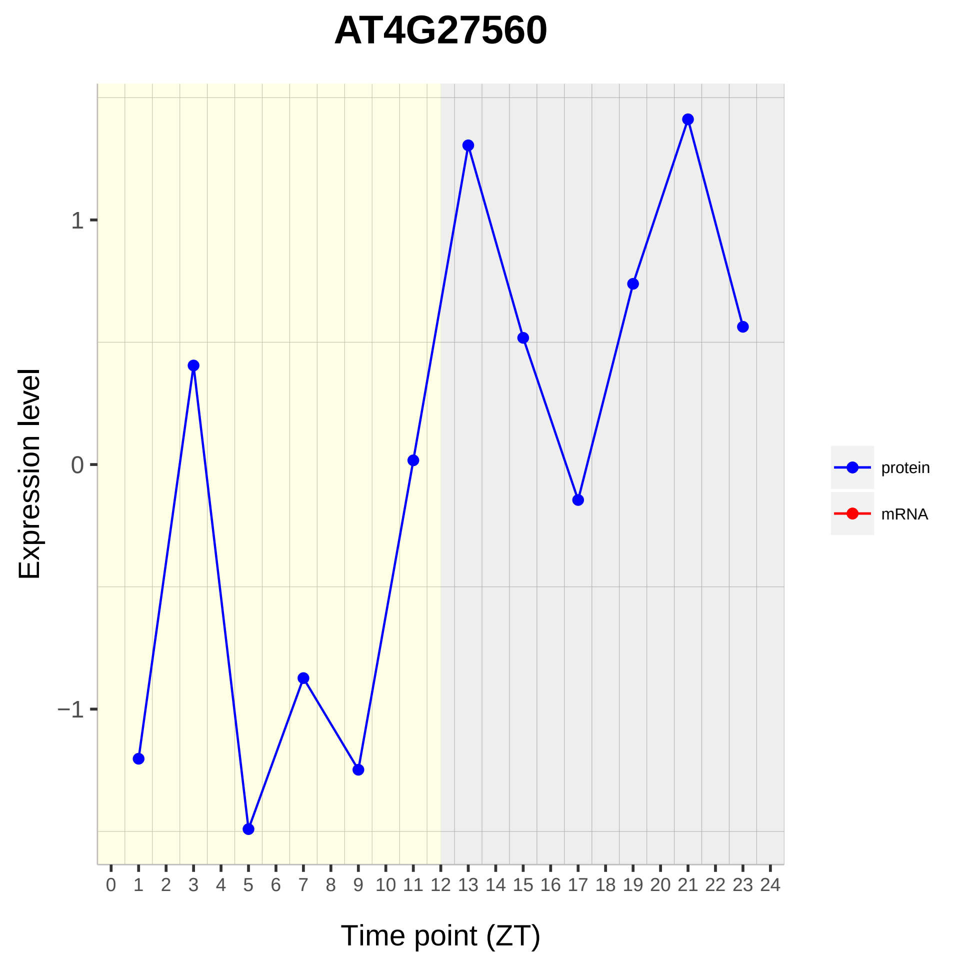
AT4G27560
Expression level
1
0
−1
0 1 2 3 4 5 6 7 8 9 10 11 12 13 14 15 16 17 18 19 20 21 22 23 24
Time point (ZT)
protein
mRNA

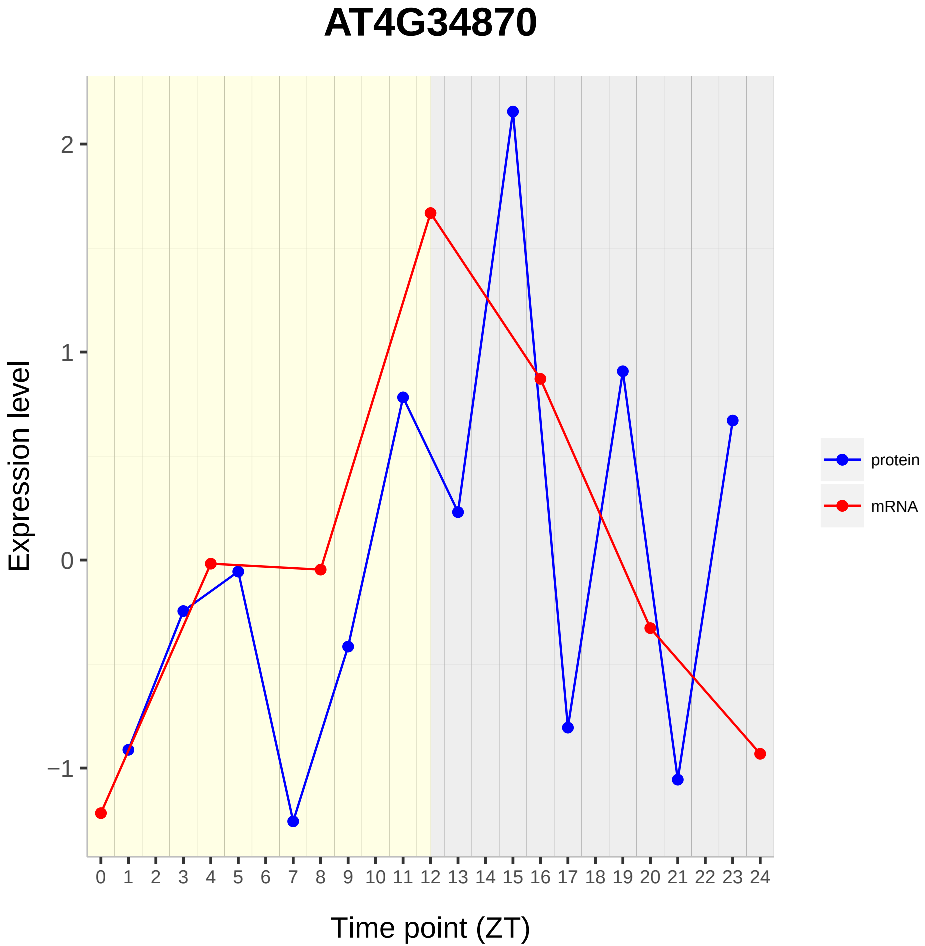

AT4G34870
Expression level
2
1
0
−1
0 1 2 3 4 5 6 7 8 9 10 11 12 13 14 15 16 17 18 19 20 21 22 23 24
Time point (ZT)
protein
mRNA

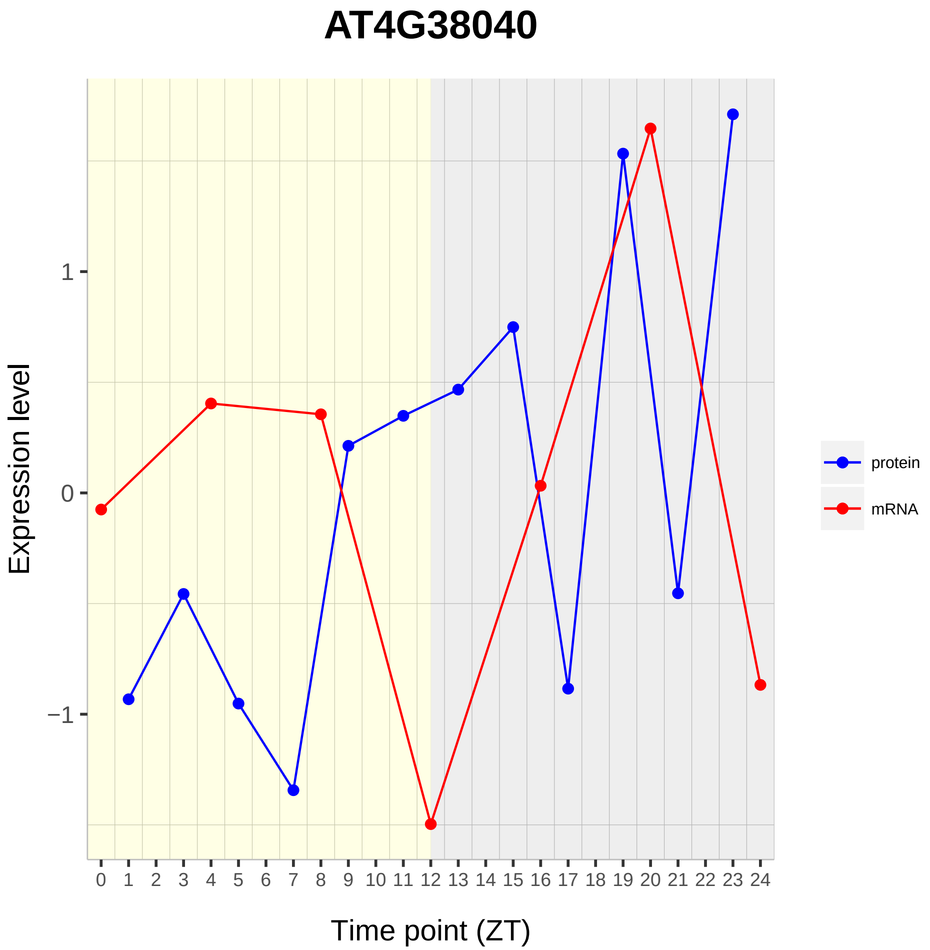

AT4G38040
Expression level
1
0
−1
0 1 2 3 4 5 6 7 8 9 10 11 12 13 14 15 16 17 18 19 20 21 22 23 24
Time point (ZT)
protein
mRNA

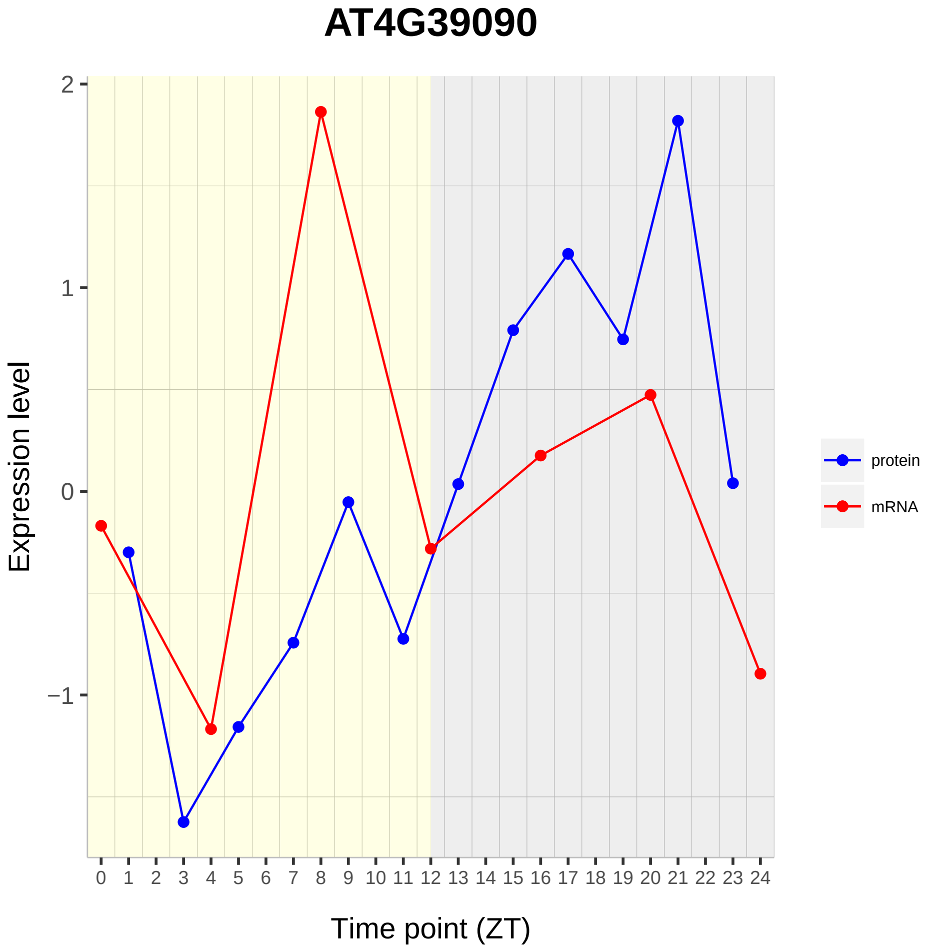

AT4G39090
Expression level
Time point (ZT)
2
1
0
−1
protein
mRNA

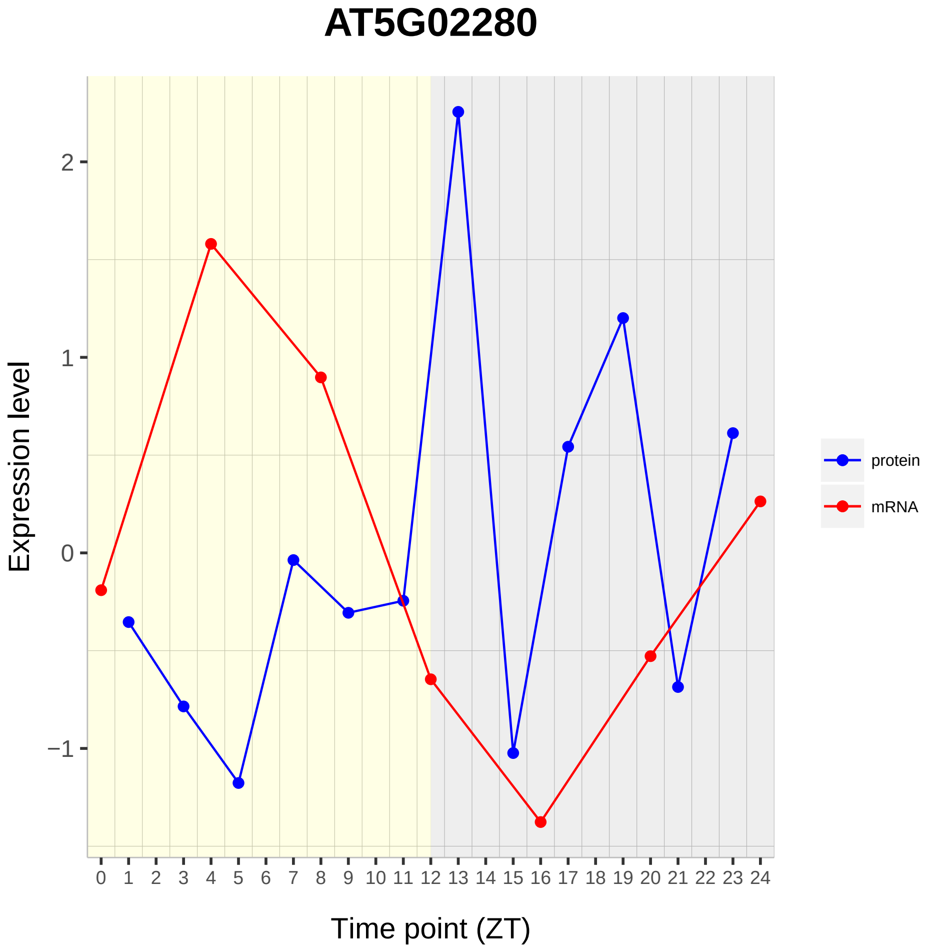

AT5G02280
Expression level
Time point (ZT)
2
1
0
−1
0 1 2 3 4 5 6 7 8 9 10 11 12 13 14 15 16 17 18 19 20 21 22 23 24
protein
mRNA

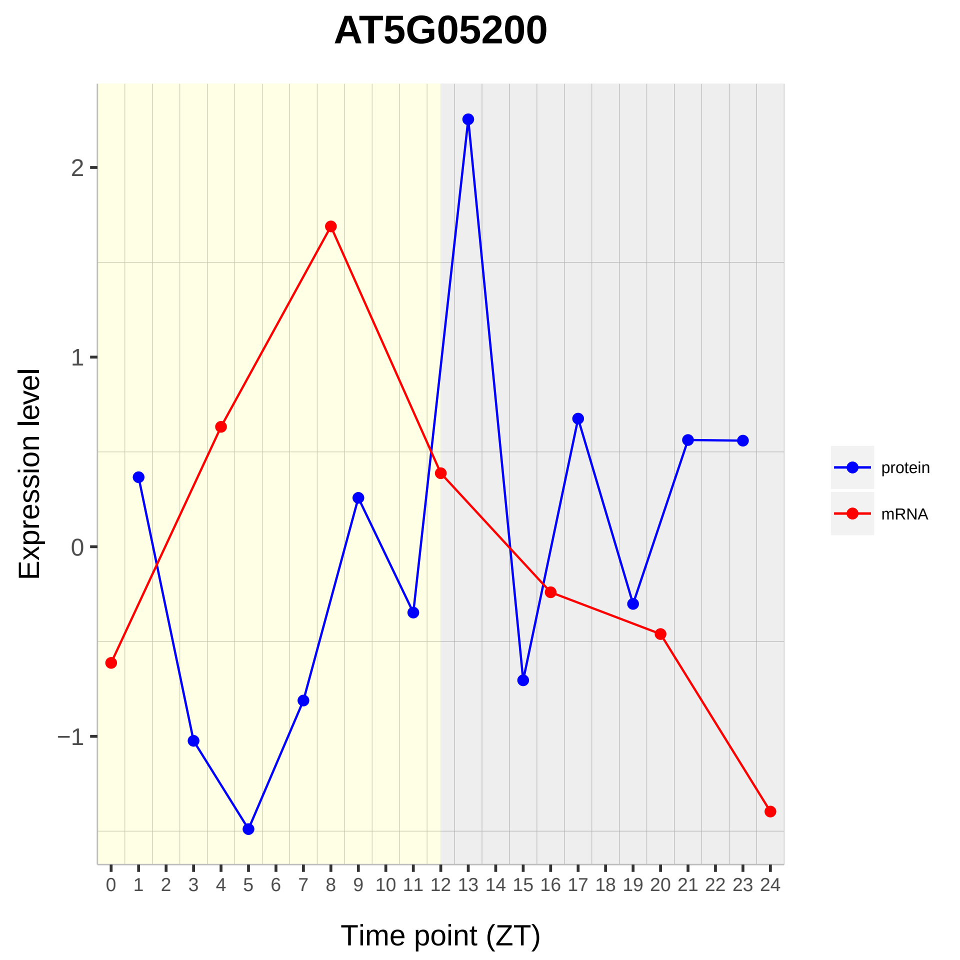

AT5G05200
Expression level
Time point (ZT)
2
1
0
−1
0 1 2 3 4 5 6 7 8 9 10 11 12 13 14 15 16 17 18 19 20 21 22 23 24
protein
mRNA

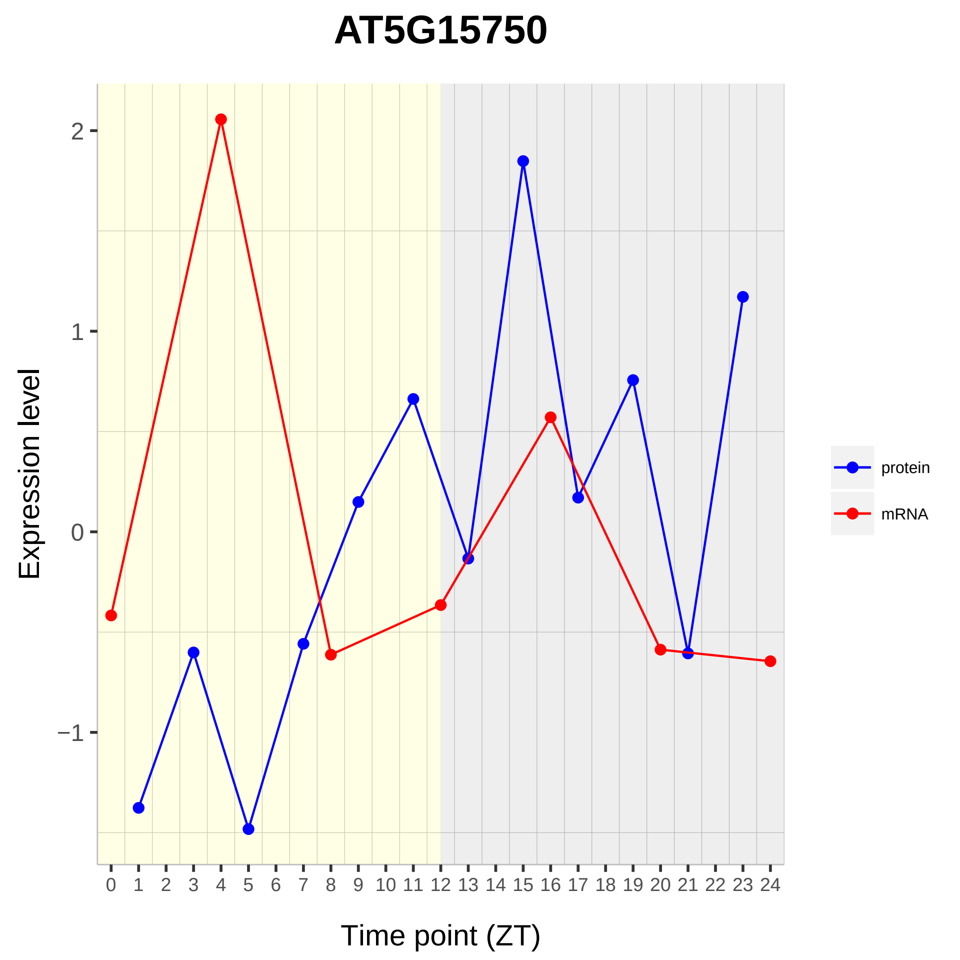

AT5G15750
Expression level
2
1
0
−1
Time point (ZT)
protein
mRNA

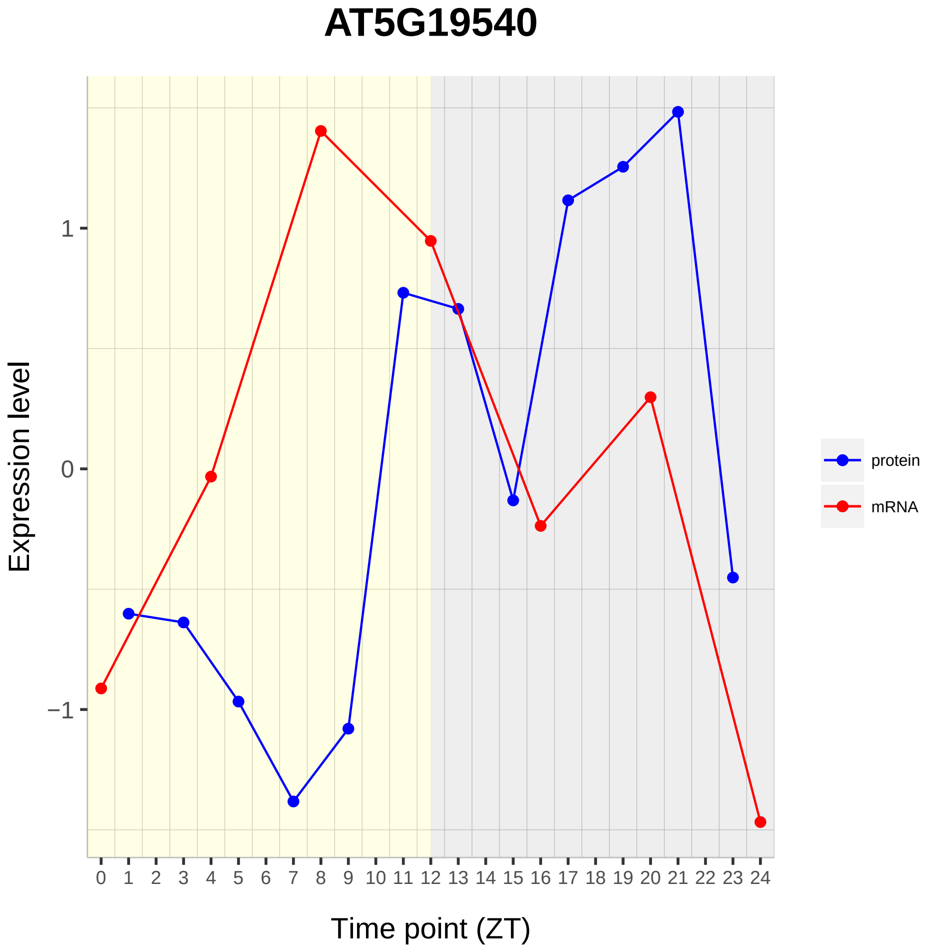
AT5G19540
Expression level
1
0
−1
0 1 2 3 4 5 6 7 8 9 10 11 12 13 14 15 16 17 18 19 20 21 22 23 24
Time point (ZT)
protein
mRNA

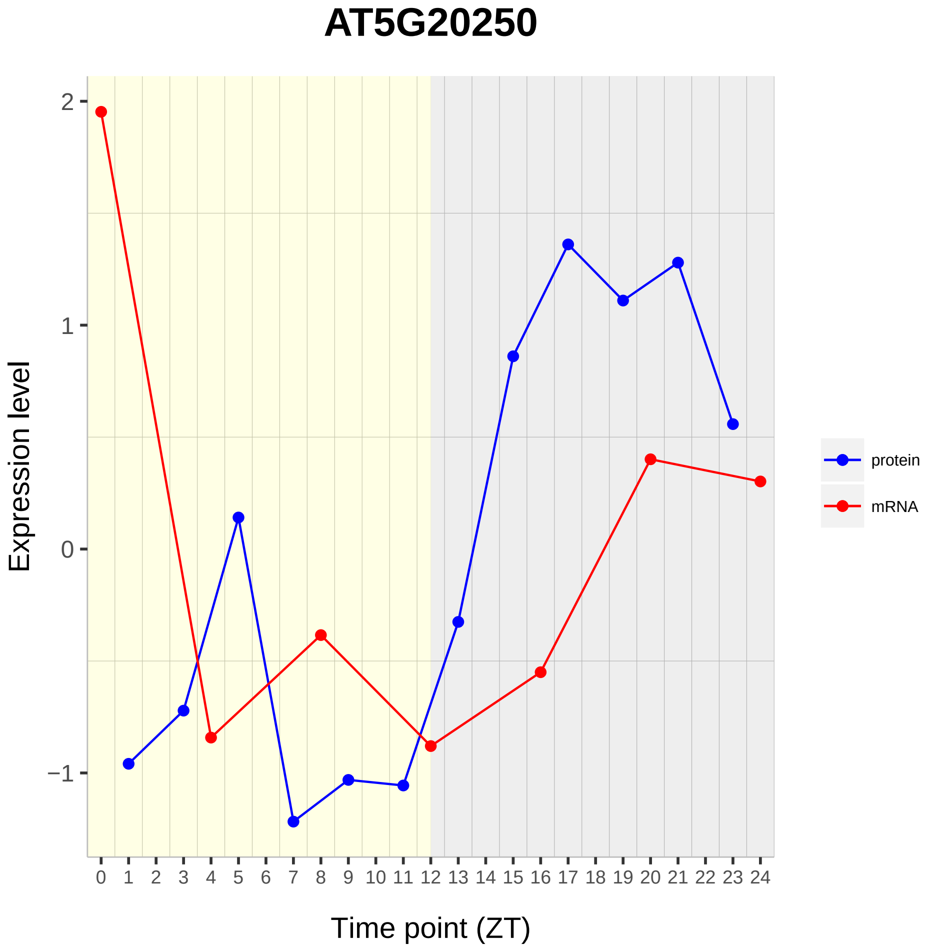

AT5G20250
Expression level
2
1
0
−1
Time point (ZT)
protein
mRNA

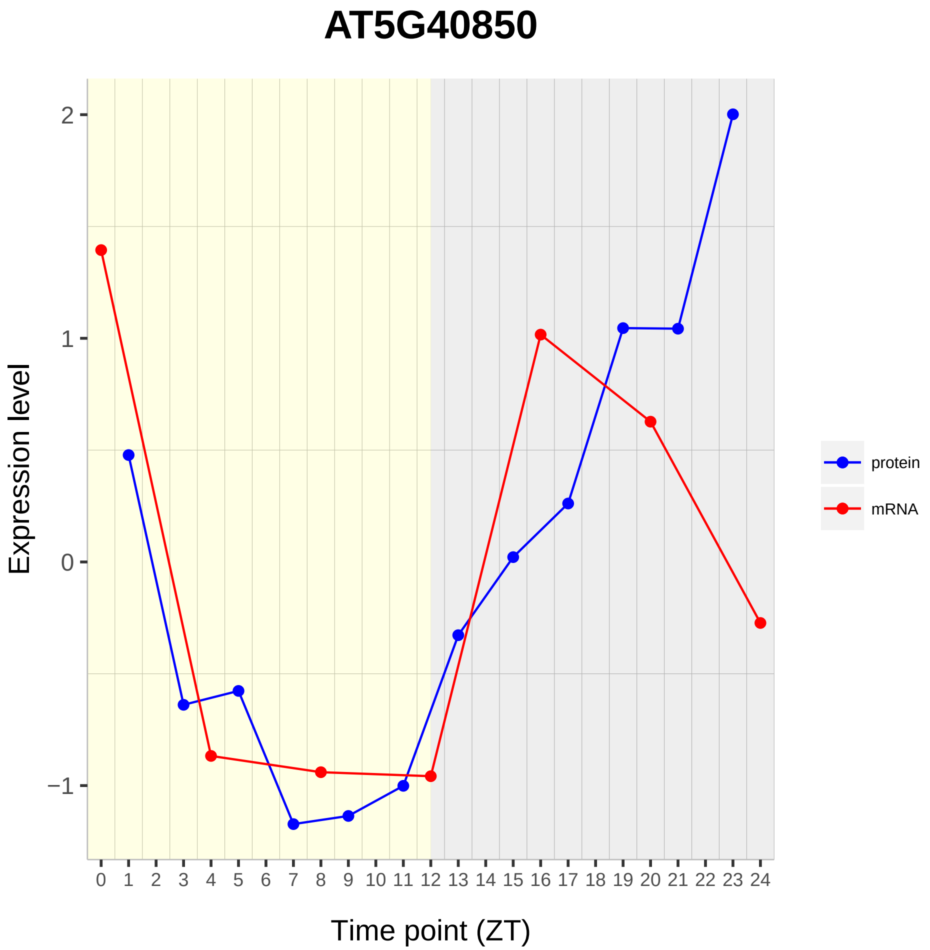

AT5G40850
Expression level
Time point (ZT)
2
1
0
−1
0 1 2 3 4 5 6 7 8 9 10 11 12 13 14 15 16 17 18 19 20 21 22 23 24
protein
mRNA

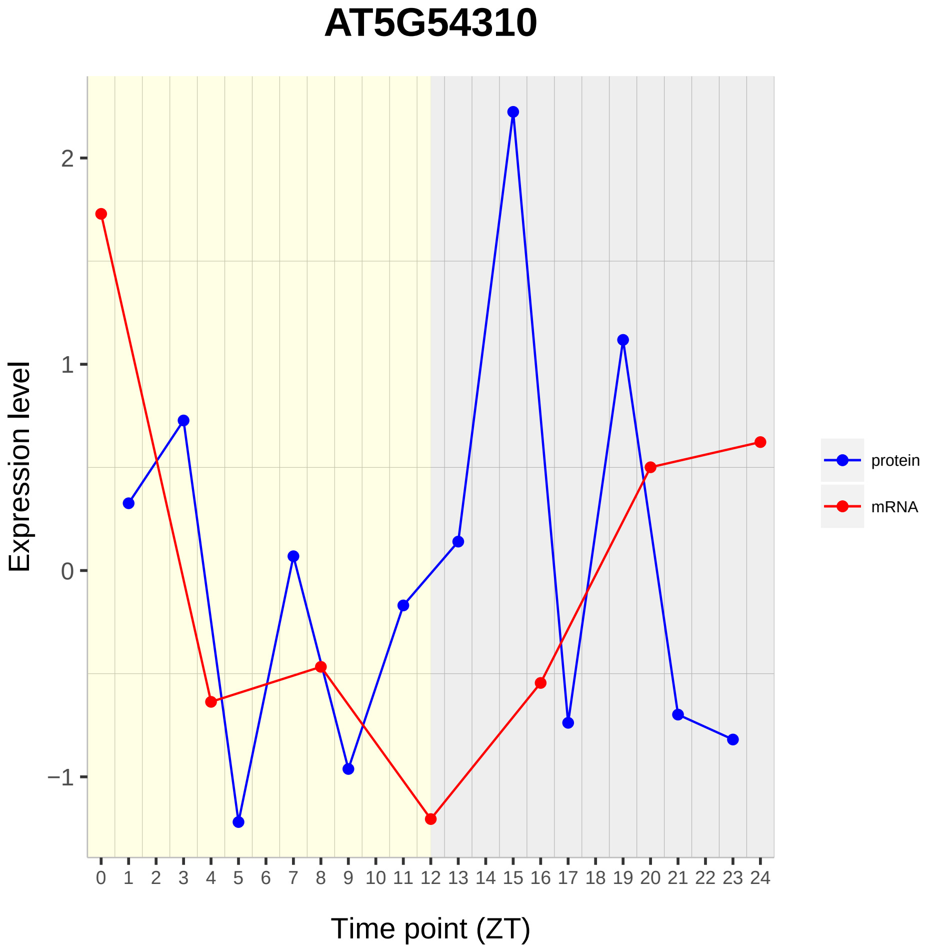
AT5G54310
Expression level
Time point (ZT)
protein
mRNA
2
1
0
−1
0 1 2 3 4 5 6 7 8 9 10 11 12 13 14 15 16 17 18 19 20 21 22 23 24

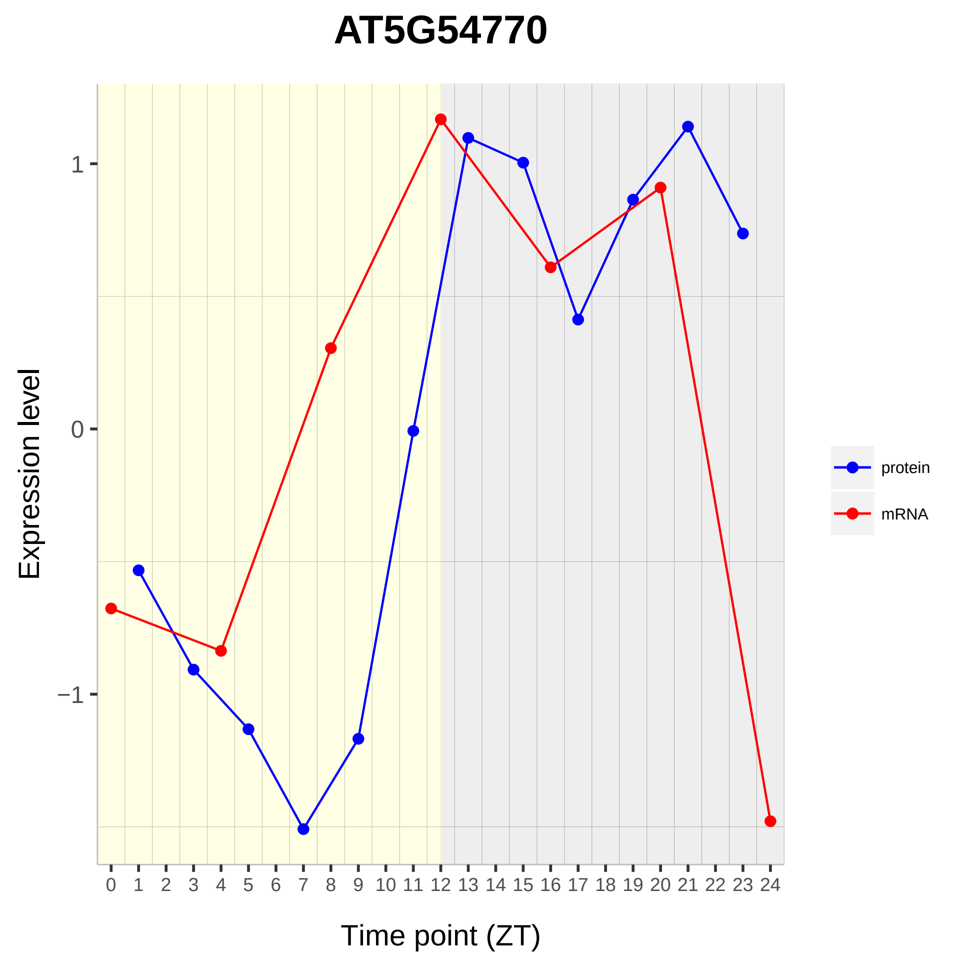

AT5G54770
Expression level
1
0
−1
Time point (ZT)
protein
mRNA

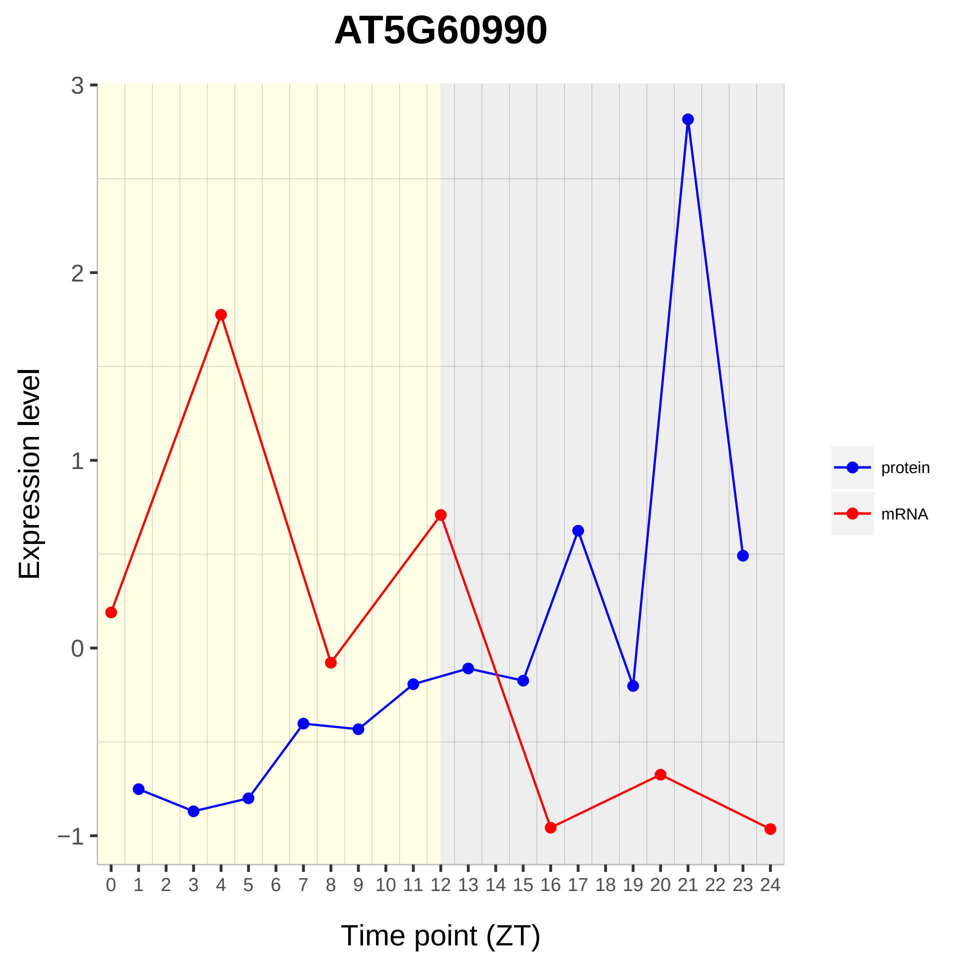
AT5G60990
Expression level
Time point (ZT)
3
2
1
0
−1
0 1 2 3 4 5 6 7 8 9 10 11 12 13 14 15 16 17 18 19 20 21 22 23 24
protein
mRNA

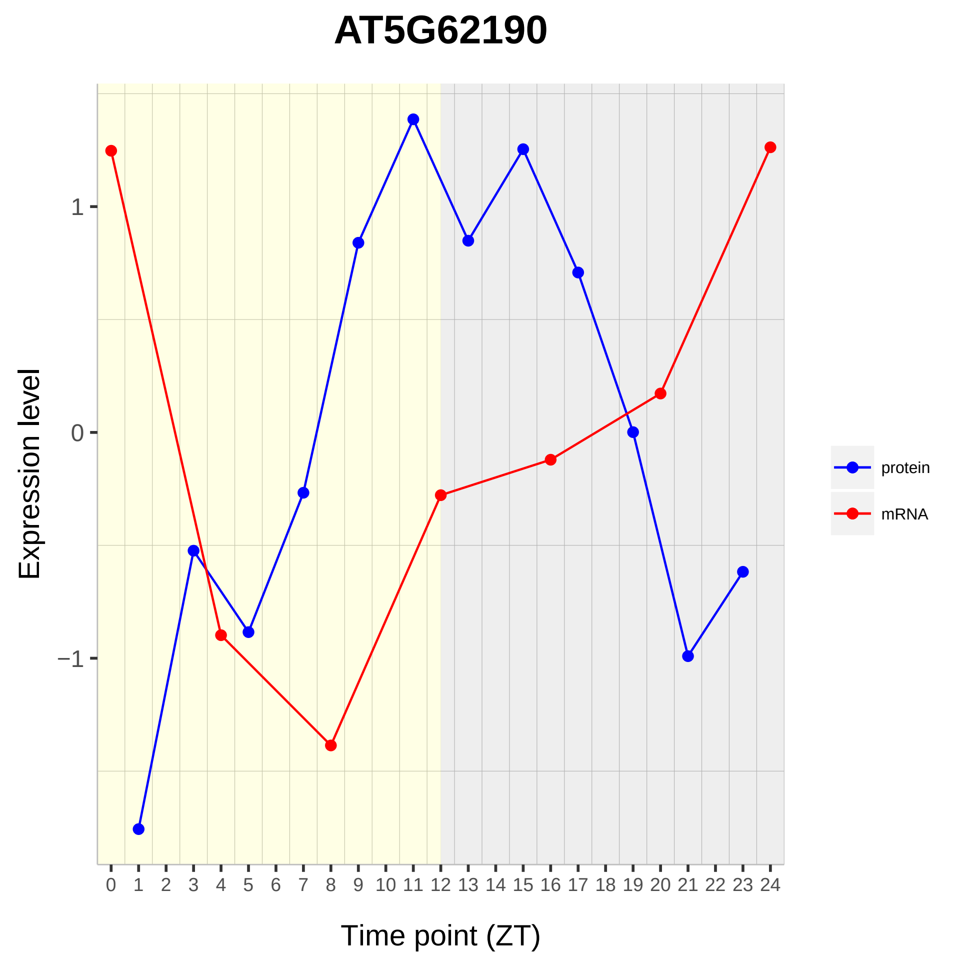

AT5G62190
Expression level
1
0
−1
0 1 2 3 4 5 6 7 8 9 10 11 12 13 14 15 16 17 18 19 20 21 22 23 24
Time point (ZT)
protein
mRNA
